# In vivo Antibody Painting for Next Generation Weight Loss Drugs

**DOI:** 10.1101/2024.08.22.609257

**Authors:** Katsushi Kitahara, Aurélie Rondon, Edward Miller, Howard H. Mak, Andrei Loas, Bradley L. Pentelute

**Affiliations:** Department of Chemistry, Massachusetts Institute of Technology, 77 Massachusetts Avenue, Cambridge, Massachusetts 02139, USA; Sumitomo Pharma Co., Ltd., 3-1-98 Kasugade-naka, Konohana-ku, Osaka 554-0022, Japan; The Koch Institute for Integrative Cancer Research, Massachusetts Institute of Technology, 500 Main Street, Cambridge, Massachusetts 02139, USA; Center for Environmental Health Sciences, Massachusetts Institute of Technology, 77 Massachusetts Avenue, Cambridge, Massachusetts 02139, USA; Broad Institute of MIT and Harvard, 415 Main Street, Cambridge, Massachusetts 02142, USA

**Author notes:** These authors contributed equally to this work (Katsushi Kitahara, Aurélie Rondon).

## Abstract

Peptide-based therapeutics are currently in great demand but often suffer from rapid clearance and accumulation in off-target tissues which continue to present barriers in their clinical translation. Here, we developed an electrophilic peptide for the attachment of therapeutics to native immunoglobulin (IgG) in vivo, enabling the bioorthogonal covalent linkage, or ‘painting’, of peptide drugs of choice to circulating IgGs directly in live animals. Native IgG painting with glucagon-like peptide-1 (GLP-1) results in sustained body weight loss and prolonged blood glucose management after one dose. Such technology might revolutionize the next generation of long-acting peptide-based medicines.

## Introduction

Peptide drugs are advantageous due to their high target selectivity, high efficacy, safety, and low cost (*1–3*). However, they also suffer from poor in vivo stability, short plasma half-life (a few minutes), and poor oral availability. Recent efforts in biotechnology focused on improving peptide pharmacokinetic (PK) and pharmacodynamic (PD) profiles using chemical modifications to diminish renal clearance (*4*), increase chemical stability (*5, 6*), and enhance bioavailability (*7*).

One of the most remarkable recent successes in the development of long-acting peptide drugs is exemplified by glucagon-like peptide-1 receptor agonists (GLP-1 RAs) (*8–10*). While native GLP-1 has a very short plasma half-life, of a few minutes, long-acting GLP-1 RAs have been proven efficient for the treatment of type II diabetes mellitus (T2DM) and obesity (*11*–*16*). GLP-1 RAs conjugation to neonatal Fc receptor (FcRn) binders, such as serum albumin or IgGs, leads to endosomal internalization, transportation and recycling to the blood (*17, 18*). Despite great achievements, the first generations of long-acting GLP-1 RAs are associated with high costs for IgG production and purification. New technologies that leverage native antibodies, obviate in vitro IgG production, and enable the assembly of the IgG-GLP-1 conjugate in vivo would greatly streamline the manufacturing process and reduce costs of next-generation long-acting GLP-1 RAs.

Here, we report a novel drug delivery platform to attach therapeutic payloads specifically to IgGs *directly in vivo*, aiming to extend the PK/PD of both existing and candidate drugs via FcRn binding. We termed our technology “in vivo antibody painting” as different lysine (Lys) residues from the Fc domain of native IgGs can be subsequently modified site-selectively to incorporate multiple drugs (Fig. 1). To achieve specific targeting of IgGs, we designed several electrophilic affinity peptides composed of three parts: (i) an Fc-binder peptide, (ii) a reactive electrophilic function, and (iii) a therapeutic payload. Through proximity-induced effect (*19*), the electrophilic payload is covalently conjugated to the heavy chain of the IgG Fc domain. The reaction is fully biocompatible, occurs at physiological pH and temperature (37 °C) without any catalyst, and does not require incorporation of any reactive handle to IgGs beforehand.

**Fig. 1:**
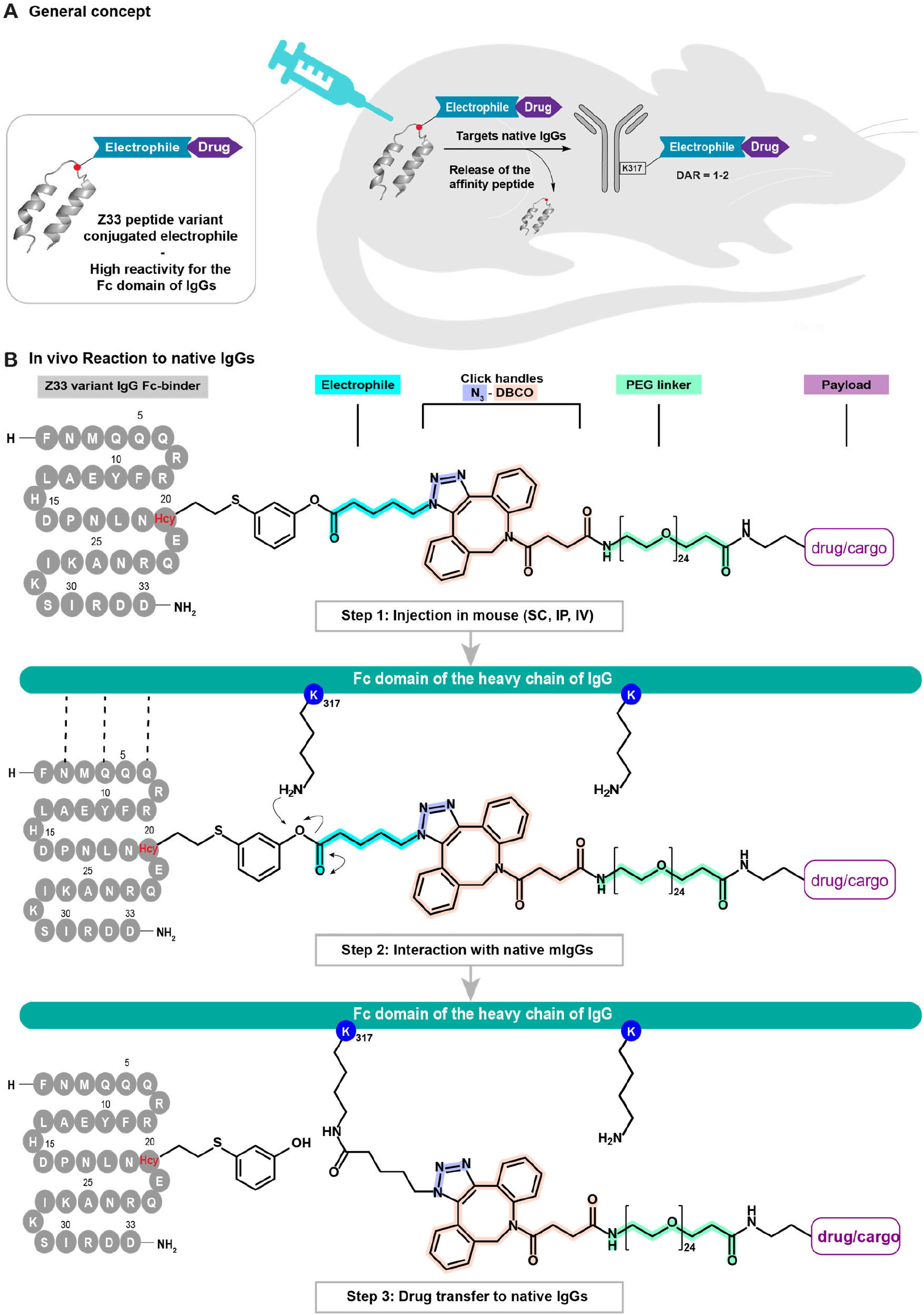
General overview of the in vivo IgG ‘painting’ technology. (A) Concept. (B) In vivo reaction to native IgGs. Z33 peptide variant conjugated to electrophile-drug moieties is administered in vivo. The peptide recognizes lysine-317 of the IgG Fc-domain heavy chain A enabling the transfer of its payload to IgGs through proximity effect. The reaction is fast (a few hours), biocompatible and covalent. The binding of IgGs to the neonatal Fc receptor (FcRn) the half-life is expected to extend and enhance drug efficacy. This technology is versatile and can be accomplished with a variety of cargoes: small molecules, therapeutic peptides, or even radionuclides. DAR: drug-antibody ratio; DBCO: dibenzyl-cyclooctyne; Hcy: homocysteine; IgG: immunoglobulin G; IP: intraperitoneal; IV: intravenous; PEG: polyethylene glycol; SC: subcutaneous.

Fc-binder affinity peptides were synthesized by Fmoc-based solid-phase peptide synthesis (Fmoc-SPPS) using our previously reported automated fast-flow peptide synthesis (AFPS) instrumentation (*20*). Their safety and duration of efficacy was assessed in two different mice models, WT lean Swiss and *Lep*^ob/ob^, the latter having spontaneous type II diabetes mellitus (T2DM) related obesity. Our results show promising outcomes for advancing the next generation of weight loss and T2DM drugs. Furthermore, we demonstrate the applicability of IgG painting to a variety of payloads (small molecules, peptides, and radionuclides), highlighting its versatility and potential for numerous applications. This work thereby represents a novel impactful insight for designing long-acting drugs and antibody-drug conjugates.

### Synthesis of Fc-reactive derivatives of the minimal Z-domain of protein A

The minimal Z-domain of protein A, a 33-mer peptide (Z33), was shown to bind the Fc fragment of IgG1 with an apparent dissociation constant (*K*_D_) of 43 nM (*21*). In a previous study, it was determined that specific amino acids in the sequence of the Z33 peptide have little to no influence on its binding to IgGs while other residues play a crucial role (*22*). The interaction between Glu-20 (E20) of protein A and Lys-317 (K317, (*23, 24*)) of the Fc domain of IgG1 was previously confirmed using co-crystal structure (Fig. S1). Selective targeting of the Fc fragment of native IgGs, using affinity peptides, was already successfully reported for manufacturing antibody-drug conjugates (ADCs), via the AJICAP® technology (*25, 26*). However, the first generation required redox treatment for reagent conjugation to thiol groups and was thereby associated to high risks of aggregation. While the second generation brought significant improvements, and enabled to maintain aggregation under 10%, IgG modification was still achieved following multi-steps chemical reactions, making the technology non-suitable for direct in vivo targeting (*27*). Following this evidence, our aim was to develop a novel drug delivery platform, derived from Z33 peptide, for targeting and modifying native IgGs directly in vivo, in one single step. Here, we expect to i) avoid aggregation and immunogenicity by using fully biocompatible reagents, ii) increase the half-life of peptide drugs, iii) administer lower doses.

We first designed and synthesized several variants of Z33 using our previously reported AFPS technology (Fig. S2A and Table S1) (*20*). Based on previous findings (*22*), we substituted E20 with either L-Cys (peptide **1**), or L-homocysteine (L-Hcy, peptides **2-6**) to introduce a handle for the site-specific conjugation of an electrophile function bearing a free azido click moiety. The azido electrophile was conjugated by S-arylation using Pd-oxidative addition complexes (Pd-OAC) (*28*), leading to the final electrophile-containing affinity peptide constructs (N_3_-**1-6**, Fig. S2B). N_3_-**1** was used as a negative control as the L-Pro-16 substitution with a D-Pro resulted in a loss of binding affinity (∼3 µM). N_3_-**3** was based on the same structure as N_3_-**2** (binding affinity ∼60 nM) except its N-terminus is capped using 8 repeated units of polyethylene glycol (PEG_8_).

### In vitro selective transfer of azido moieties onto human and mouse IgGs

Several electrophiles (**I, II, III, IV, V**) were synthesized, and their reactivity after 24 h was assessed in vitro, using hIgG1 trastuzumab (Tmab; 1 eq.) (Fig. S3). The highest reaction transfer was observed for electrophile **I**, with up to 99% of modifications incorporated into the heavy chain of Tmab after reaction with peptide **2** (20 eq.) compared to only 47% with electrophiles **II** or **III**. Through a sequencing experiment using tandem liquid chromatography-mass spectrometry (LC-MS/MS), we were able to identify that the electrophile moiety was covalently transferred to the Lys-317 side chain of the heavy chain of the Fc domain of Tmab (Fig. S4). Further investigations showed conversions of ∼17%, 99%, and 95%, after reaction with peptides N_3_-**1-3** bearing electrophile **I**, respectively, confirming the highest reactivity associated with N_3_-**2** (Table S2). In vitro reactivity of peptide N_3_-**2-I** (20 eq.) was then assessed in vitro for other IgG subtypes and species and showed the highest reactivity was achieved using hIgG1-2 and excess amount of N_3_-**2-I**, with 98% conversion on the heavy chain Fc domain associated to a high selectivity with less than 2% of the light chains modified with the electrophilic groups (Fig. S5 and Table S3).

Peptide N_3_-**2-I** (20 eq.) incubated with a mixture of Tmab (1 eq.) and RNase A (1 eq.) showed that only Tmab ended being decorated with azido moieties (DAR 1-2) (Fig. S6). In addition, incubation of N_3_-**1-6-I** in mouse sera confirmed the absence of binding to other biomolecules circulating in blood (Fig. S7A-D). The highest IgG conversion rate was achieved using N_3_-**2-I** and N_3_-**3-I**, with 33 ± 1% and 49 ± 3%, respectively, after 2 h of incubation, versus 10 ± 1% for the low binder N_3_-**1-I** Given these good conversion rates, N_3_-**2**,**3-I** were taken forward for investigations in WT Swiss mice (Fig. S7E-F). We calculated that at least 30 mg/kg of N_3_-**2**,**3** (∼280 nmol) would be required to modify ∼100% of the mIgGs circulating in blood (∼3 mg/mL of mIgGs), with a DAR of at least 1 and a conversion rate of 100% (*29, 30*). However, as N_3_-**2-3** only induced up to 50% of electrophile transfer, in vitro, we expected to modify only 50% of the mice native IgGs in vivo, at best, after a single dose of 30 mg/kg. The dose of 30 mg/kg of N_3_-**2-3-I** was well tolerated after either subcutaneous (SC) or intraperitoneal (IP) injection, no significant adverse effects being observed. N_3_-**2-I** and N_3_-**3-I** showed similar in vivo reactivity, with 33 ± 12% and 37 ± 4% conversion rate, respectively, after SC injection. Considering that the two heavy chain Lys-317 side chains of the Fc domain can be modified using N_3_-**2-3-I** reagents, the painting of mIgGs resulted in a distribution of DAR from 0 to 2. This methodology does not enable to quantify how many modifications are carried on mIgG; therefore, we assume the conversion rate quantified here is probably underestimated.

### In vitro selective transfer of GLP-1 conjugates onto mouse IgGs

Motivated by the encouraging results obtained with azido transfer, we sought to apply our IgG painting technology to larger biomolecular cargos, such as GLP-1 RAs for extending their PK. Three different GLP-1 backbones were synthesized using AFPS (Fig. S8), based on the sequences of Semaglutide (GLP-1**a**), Liraglutide (GLP-1**b**), and Tirzepatide (GLP-1**c**). GLP-1**a**,**b** were covalently conjugated to a dibenzyl cyclooctyne (DBCO) moiety on Lys-20 (Fig. S8-S12) then reacted with N_3_-**2-I** to obtain electrophile peptides **2a**,**b** through the strain-promoted azide-alkyne cycloaddition click reaction (SPAAC) (*31*). Electrophile peptide **2c** was obtained following a multiple step conjugation process (Fig. S8B). The conversion rate of peptides **2a-c** determined by Q-ToF LC-MS indicated that up to 41% of Tmab Fc domain heavy chains were successfully modified with GLP-1**a**, 26% for GLP-1**b**, and 20% with GLP-1**c** (Fig. S10-S12, and Table S4). Following these encouraging results, we decided to assess the effect of in vivo IgG painting with peptides **2a**,**3a** in vivo, in two different mice models.

### In vivo IgG painting is well tolerated in mice

Three different doses of the commercial Semaglutide (0.5 mg/kg (∼5 nmol), 3 mg/kg (∼30 nmol) or 10 mg/kg (∼100 nmol)) were first SC injected in WT mice to verify the tolerability for GLP-1 RAs (Table S5). In the other cohorts, **2a**,**3a** was SC administered following either 10 mg/kg (25 nmol), 30 mg/kg (75 nmol), or three injections of 10 mg/kg, 1 per week for 3 weeks (30 mg/kg total, 75 nmol total). As expected, hypoglycemia was observed starting 0.5 mg/kg of Semaglutide and became critical for the highest dose suggesting 10 mg/kg as the maximum tolerated dose. None of the mice receiving SC injections of **2a** and **3a** showed hypoglycemia or behavioral change, confirming the safety of IgG painting with GLP-1 RAs.

In *Lep*^ob/ob^ obese mice, two routes were assessed for the10 mg/kg dose: SC and IP (Table S6). For Semaglutide, both routes did significantly impact the mice behavior and induce hypoglycemia. In the **2a** and **3a** cohorts, the SC route was better tolerated with no significant toxicity while IP induced some behavioral changes. We thereby pursued our investigations about the PD of **2a** and **3a** following SC injection of 10 mg/kg, in both mice models.

### The pharmacodynamic profile of GLP-1 is significantly improved after in vivo IgG painting in lean mice

A dose-response using the commercial Semaglutide (0.5 mg/kg (∼5 nmol), 3 mg/kg (∼30 nmol), or 10 mg/kg (∼100 nmol), SC) allowed us to validate the WT Swiss mice model as suitable for measuring blood glucose change in addition to body weight loss (Fig. S13). Then, female WT mice received one single SC injection of either 10 mg/kg (∼100 nmol) of Semaglutide, or 10 mg/kg (∼25 nmol) of **2a** and **3a** (Fig. 2A,B). A significant 4-5% body weight loss was observed at 24 h post injection, sustained for 10 days (**2a**) or 15 days (**3a**) (Fig. 2C).

**Figure 2:**
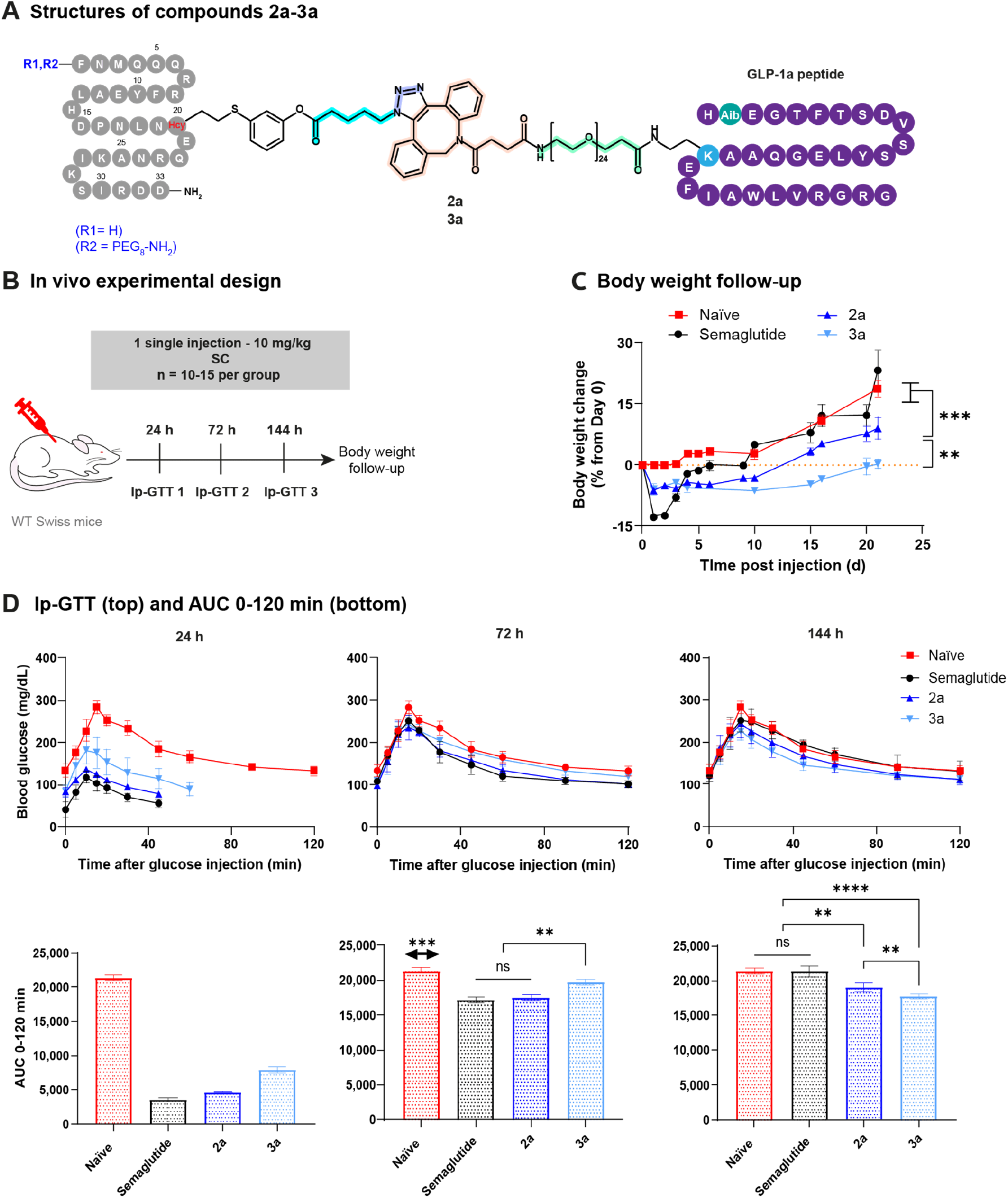
In vivo IgG painting extends GLP-1a pharmacodynamic profile in WT Swiss mice. (A) Structures of the synthesized compounds **2a-3a**. (B) In vivo experimental design: female WT Swiss mice were SC injected with 10 mg/kg (∼100 nmol) of Semaglutide or with 10 mg/kg (∼25 nmol) of **2a** or **3a**. (C) Body weight change from Day 0. (D) Ip-GTT curves (top) and AUC (bottom) at 24 h, 72 h, and 144 h post drug injection. Statistical analysis was performed using two-way ANOVA (C), multiparametric T-tests (C), or non-parametric T-tests (D): *P<0.05, **P<0.01, ***P<0.001, ****P<0.0001. AUC: area under the curve; GTT: glucose-tolerance tests; IP: intraperitoneal; SC: subcutaneous.

Intraperitoneal glucose tolerance tests (Ip-GTT) demonstrated blood glucose decrease for all the GLP-1 analogs compared to the naïve mice, after 24 h of injection (Fig. 2D). After 6 days, Semaglutide showed no more ability to reduce blood glucose in the WT mice (AUC_0-120 min_ = 21,282 ± 795 vs 21,354 ± 431 for the naïve) while **2a** and **3a** were still exerting significant activation of GLP-1R (AUC_0-120min_ = 19,004 ± 650 and 17,744 ± 366, respectively). In addition, a stacking dose of three injections of 10 mg/kg each (∼75 nmol total), following one injection per week for three weeks induced similar efficacy as one single dose of 10 mg/kg, while a single dose of 30 mg/kg of **2a** was significantly more efficient than a 10 mg/kg dose (P<0.0001), at both 24 h and 72 h p.i, but presented no benefit at the 144-h time-point (Fig. S14).

### In vivo IgG painting with GLP-1a sustains body weight loss and improves blood glucose management in obese *Lep*^ob/ob^ mice for at least 10 days after SC injection

Both males and females were used for this efficacy study to avoid sex-related biases. One single injection of 10 mg/kg (∼170 nmol) of Semaglutide, or 10 mg/kg (∼45 nmol) of **2a**,**3a** was SC in *Lep*^ob/ob^ mice (Fig. 3A,B). Significant body weight loss (∼3.6 ± 0.8 and 4.3 ± 1.4%) was observed for 10 days after **2a** or **3a** injections, respectively, with slower body weight intake up to Day 21 (Fig. 3C). Semaglutide induced about 8.3 ± 0.5% of body weight loss for 3 days, then the intake went progressively up until 21 days p.i.

**Figure 3:**
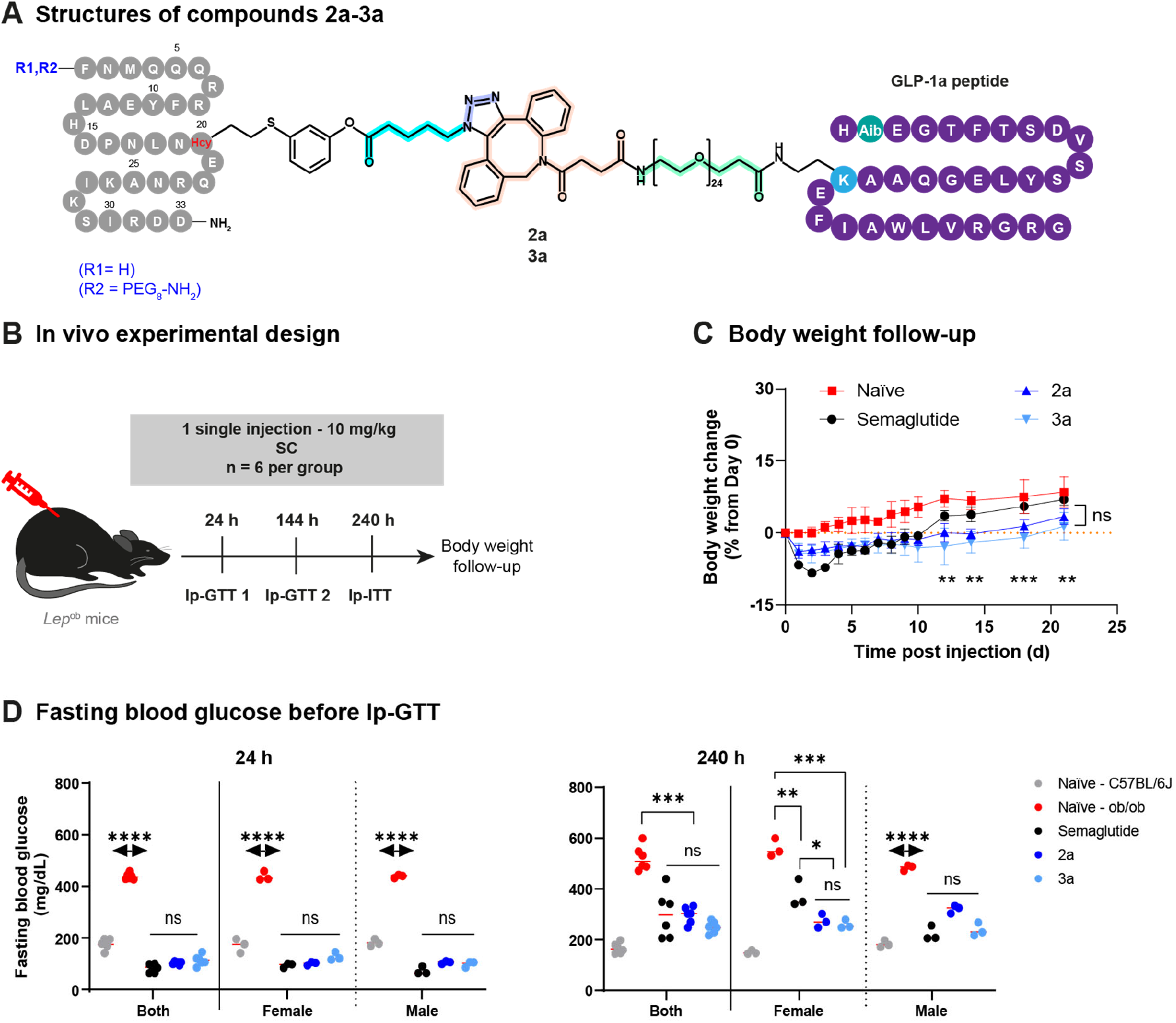
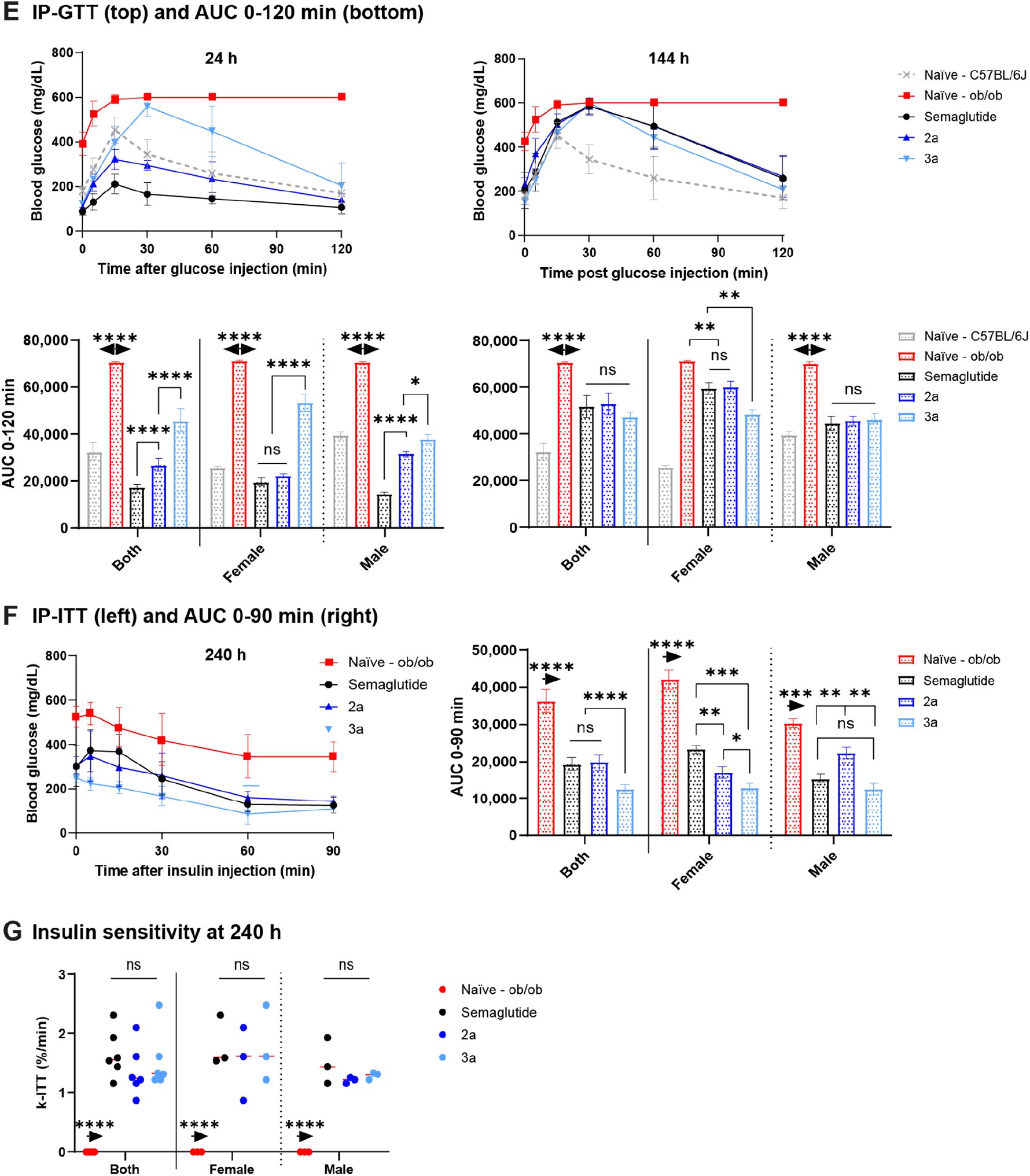
In vivo IgG painting with 2a,2b sustains body weight loss and exerts extended blood glucose control in obese *Lep*^ob/ob^ mice. (A) Structure of compounds **2a-3a**. (B) Experimental design: *Lep*^ob/ob^ obese mice (n = 6 per compound, split as 3 males and 3 females) were SC injected with 10 mg/kg (∼170 nmol) of Semaglutide or 10 mg/kg (∼45 nmol) of **2a**,**3a**. Non-obese C57BL/6J mice (n = 6, 3 males and 3 females) have also been included in the study as an additional naïve control to obtain the baseline values for healthy subjects. (C) Body weight change from Day 0. (D) Blood glucose measured after 6 hours of fasting. (E) Ip-GTT curves (top) and AUC (bottom) at 24 h and 144 h p.i. (F) Ip-ITT curve (left) and AUC 0-90 min (right) at 240 h p.i. (G) Plasma glucose disappearance rate (k-ITT) determined at 240 h p.i. Statistical analysis was performed using two-way ANOVA (C), multiparametric T-tests (C), or non-parametric T-tests (D-G): *P<0.05, **P<0.01, ***P<0.001, ****P<0.0001. AUC: area under the curve; GTT: glucose tolerance test; IP: intraperitoneal; ITT: insulin tolerance test; p.i: post injection, SC: subcutaneous.

*Lep*^ob/ob^ are insulin resistant as confirmed by the blood glucose levels measured for the naïve, with 440 ± 9 mg/dL vs 178 ± 14 mg/dL for the non-obese C57BL/6J mice, both sexes combined (Fig. S15B). The cohorts were fasted for 5-6 hours then challenged with glucose injections (2g/kg) at 24 and 72 h p.i (Fig. S15D, left), and with insulin injection (2.5 IU/kg) at day 10 p.i. Semaglutide, **2a** and **3a** were efficient up to 10 days p.i., enabling to down-regulate blood glucose levels significantly, in both sexes combined (Fig. 3D-E). Moreover, the response to insulin observed after 10 days highlighted that **2a**,**3a** induced similar effect than Semaglutide, **3a** even showing a slightly better efficacy in females (P<0.001) (Fig. 3F). The improvement of the insulin sensitivity status was confirmed using the plasma glucose disappearance rate (k-ITT) (Fig. 3G). In *Lep*^ob/ob^ mice, in vivo IgG painting with GLP-1 analogs demonstrated similar efficacy than Semaglutide, after SC injection, even though **2a**,**3a** contain almost 4 times less GLP-1 peptide compared to the Semaglutide drug.

### In vivo IgG painting with GLP-1a improved metabolic health in *Lep*^ob/ob^ mice after IP administration as well

The IP route was assessed in male and female *Lep*^ob/ob^ mice (Fig. S16A-B). Body weight loss after IP **2a**,**3a** was comparable to the SC cohorts (P<0.0001) (Fig. S16C). However, IP Semaglutide sustained body weight loss up to 7 days p.i, then followed a similar pattern as **2a**,**3a**. Fasting blood glucose levels were maintained in the normal range for up to 6 days (Fig. S16D and Fig. S15D, right). After 10 days, Semaglutide showed no more efficacy while **2a**, and **3a** still enabled to lower blood glucose (P<0.0001 and P<0.001, respectively). IP seems to induce a faster clearance of Semaglutide while increasing the efficiency of **2a**,**3a** (Fig. S16E). After 10 days, **2a** was the most effective for reducing blood glucose in response to insulin, in both sexes, highlighting that the presence of the PEG in **3a** reduced the affinity peptide efficiency to bind IgGs after IP injection (Fig. S16F). All the GLP-1 analogs enabled to reverse insulin resistance in *Lep*^ob/ob^ mice (P<0.0001) (Fig. S16G).

### IgG painting with radionuclides highlights the payload versatility

We hypothesized that our in vivo IgG painting approach can be used as a general drug delivery platform suitable for numerous kinds of payloads. Radionuclides are among the most challenging payloads but are particularly useful for in vivo detection and quantification. N_3_-**1-3-I** was radiolabeled using zirconium-89 and the fate of [^89^Zr]Zr-**1-3** was assessed in female WT mice, after either intravenous (IV), SC, or IP injection of 1.2-1.5 MBq (Fig. 4A-B and Fig. S17-S19). In mice, the highest amounts of FcRn receptors were found in the following tissues: small intestine, spleen, large intestine, kidney, liver and lungs (*32, 33*). As functional FcRn binds, transports, and recycles IgGs, after the IV injection of [^89^Zr]Zr-**1-3**, the resulting [^89^Zr]Zr-IgGs were most likely expected to be located in those very same tissues.

**Figure 4:**
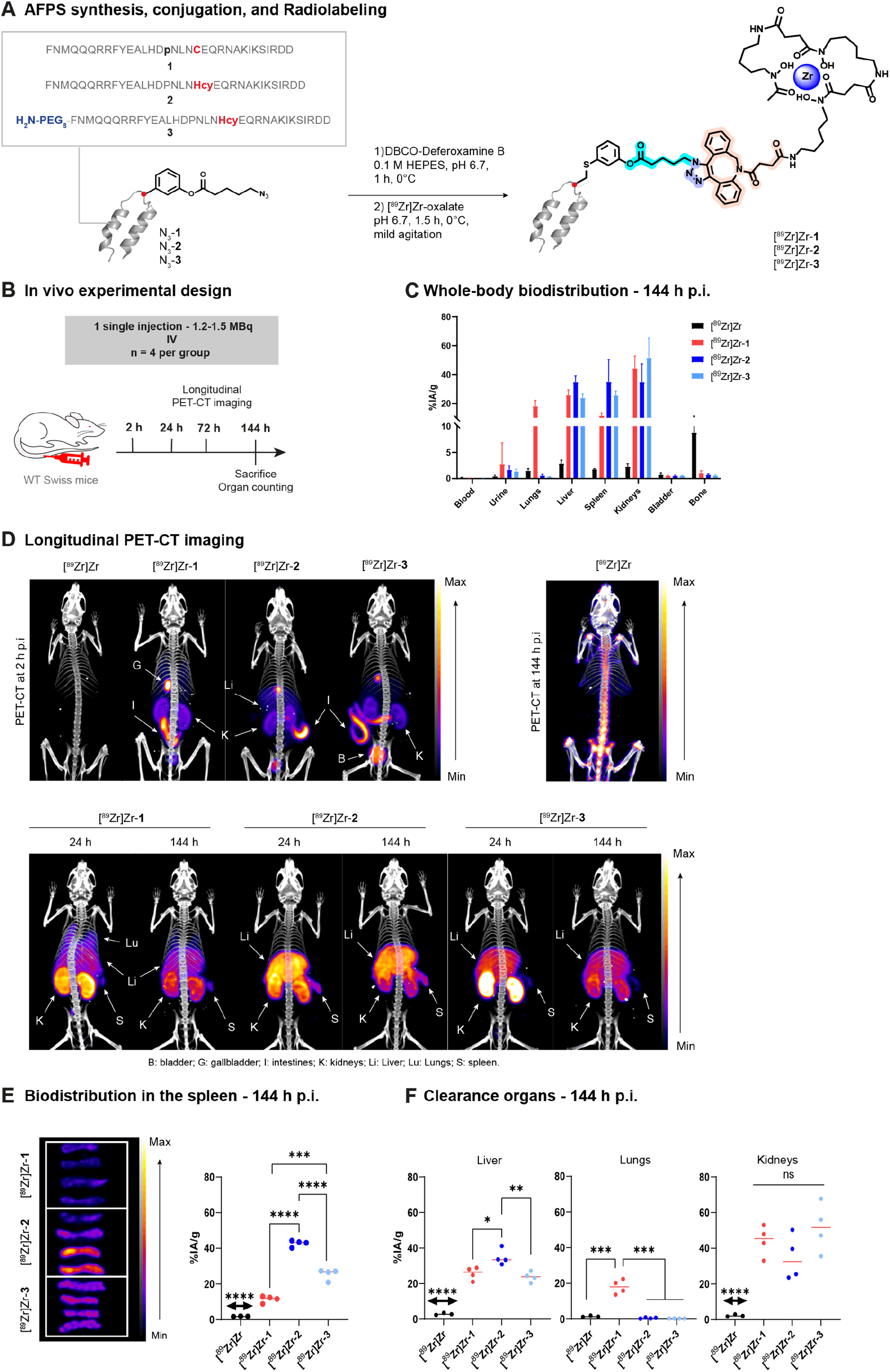
Payload versatility highlighted by in vivo radionuclide transfer to native mIgGs in WT Swiss mice. (A) Radiolabeling of peptides **1-3**. (B) Experimental design: 1.2-1.5 MBq of [^89^Zr]Zr-**1-3** was administered IV in female WT Swiss mice (n = 4 mice per compound). Longitudinal PET-CT imaging was performed over 6 days, and organs were harvested for quantification. (C) Whole-body biodistribution of [^89^Zr]Zr-**1-3** at 144 h p.i. (D) Representative longitudinal PET-CT imaging at 2 h (top), 24 h (bottom), and 144 h (bottom) p.i. Scale (top): min = 1.0e+05 kBq/mL, max = 3.0e+06 kBq/mL; (bottom): min = 2.7e+04 kBq/mL, max = 4.0e+05 kBq/mL. The same thresholds have been applied for the PET of the free [^89^Zr]Zr used as control. (E) PET-CT of all spleens and uptake at 144 h p.i. Scale: min = 1.31e+04 kBq/mL, max = 3.73e+05 kBq/mL. (F) Comparison of the %IA/g in the liver, lungs, and kidneys, at 144 h p.i. Statistical analysis was performed using non-parametric T-tests: *P< 0.05, **P<0.01, ***P<0.001, ****P<0.0001. %IA/g: percent of injected activity per gram of tissue; IV: intravenous; PET-CT: positron emission tomography-computed tomography scan; p.i.: post injection.

Longitudinal PET-CT imaging and ex vivo quantifications in organs confirmed the uptake of IV injected [^89^Zr]Zr-**1-3** in organs rich in FcRn, for up to 6 days p.i. (Fig. 4, Fig. S17, and Table S7). De-chelation was neglectable over the 6-days experiment period, confirming that the uptake quantified in the tissues is specific (Fig. S17D) (*34*). The SC route appeared to be less efficient than the IV, as the uptake measured in tissues is neglectable after 24 h, except for the kidneys and the liver (Fig. S18 and Table S8-S9). Longitudinal PET-CT also indicated that most of the signal was stuck at the injection point for 10 days, suggesting that SC is less suitable for radionuclide transfer using the in vivo IgG painting approach than IV injection (data not shown). IP injection showed better suitability than SC, with a persistent uptake organs rich in FcRn after injection of [^89^Zr]Zr-**2**(Fig. S19-S20 and Table S10).

Altogether, these results indicate that in vivo IgG painting using electrophile peptide **2**,**3**-**I** may be impactful for the development of long-acting drugs by proposing a broad platform freed from any cumbersome in vitro antibody engineering.

### Expanding the chemistry towards multi-drug painting and ADC engineering

To enable the conjugation of two or more different drugs on one single IgG, we have explored two different strategies: (i) the site-specific modification of two different lysine side chains using different electrophilic warheads, (ii) the use of a bivalent electrophile. First, we synthesized 5 novel Z33 variants (Tables S1). For peptides **7** and **8**, the L-Cys or L-Hcy residue substituted Met-3, then were conjugated to azido electrophiles **VI** and **VII** (Fig. S21-S24). Tmab sequencing indicated that azido conjugation occurred predominantly (∼93.63%) on Lys-248. Variants **9-11** carry a substitution of either Arg-31, or Asp-32 for the incorporation of electrophile **VII**. About 91% of Tmab heavy chain was successfully modified with two or more azido moieties while 86% of the light chain remained unaffected, highlighting the possibility to incorporate two different drug payloads via two steps serial reactions.

Multi-drug conjugation was then achieved through the linkage of peptides **2** and **3** to **VIII**, a dendronized electrophile bearing both an azido and a Tz moiety (Fig. S25). N_3_-Tz-**2**,**3 VIII** was conjugated to DBCO-PEG_24_-GLP-1**a**, then to TCO-DFO and radiolabeled with Zr-89, leading to [^89^Zr]Zr-**12a**,**13a**. After SC injection in WT female mice (1.2-1.5 MBq), a significant uptake was observed in the gut and in lymph nodes, confirming the reaction to mIgGs.

It is important to note that our novel drug delivery system can also be used in vitro, for engineering traditional antibody-drug conjugates (ADC), in a fast, site-selective, and highly reproducible manner (Fig. S26A-B). Here we attached monomethyl auristatin E (MMAE) to Tmab, modifying up to 55% of the heavy chains of the Fc domain of Tmab with a DAR of 2 and a 99% selectivity. Hereby, our results highlight that this new drug delivery platform may be used for a wide variety of applications.

## Conclusion

Fusion to FcRn binder molecules for PK extension purposes was explored in the past two decades and has already led to the approval of several drugs (*18, 35*). Applied to GLP-1 analogs, such approaches have provided T2DM patients an efficient therapeutic option and recently, FDA approval was obtained for weight loss applications (*36, 37*). A recent report on AMG-133 indicated that the fusion of two GLP-1 moieties on an anti-GIP IgG enabled body-weight loss in db/db mice after 24 h and persistent up to 216 h post IP injection of 2 mg/kg. In the same study, blood glucose was reduced up to 144 h post injection (*15*). In addition, in humans, AMG-133 was safe and tolerable besides an increase of amylase and lipase, possessed a half-life of 14-16 days with peak of duration reached 4-7 days post SC injection, and maintained body weight loss up to 120 days after the last dose. These data suggest that an antibody-GLP-1 conjugate may be effective at a twice monthly dosing regimen rather than the once-weekly Semaglutide, with a significant lower toxicity. The manufacturing of antibodies is however time consuming, and among the most-costly steps in the entire production pipeline. Therefore, the novel in vivo antibody painting technology we developed here proposes an efficient, straightforward, and cheaper alternative.

We report the synthesis of 13 different Fc-binder peptides, derived from the structure of Z33 (*21*). Every compound was obtained with high purity and good yield, as a result of AFPS synthesis technology (*20*). Seven electrophilic groups were synthesized, conjugated to the Fc-binder peptides to evaluate their transfer reactivity towards the heavy chain Fc domain of Tmab, enabling us to determine the best Fc-binder peptide/electrophile pair for moving forward to in vivo experiments. We successfully demonstrated the ability to transfer a cargo from a peptide to a native IgG while circulating in blood, in two hours. To the best of our knowledge, such a transformation has not been reported before.

The proof-of-concept of in vivo IgG painting with GLP-1 analogs was performed in WT mice and *Lep*^ob/ob^ (*38*) to evaluate the safety, the PK/PD extension, and the efficacy on obesity and TD2M. In both models we successfully demonstrated a sustained body weight loss for 10 to 15 days, associated with blood glucose decrease for 144 h post SC injection. Moreover, in the obesity model, a better response to insulin was also measured after 10 days following either SC or IP injection. While the outcome was similar, slight sex-related differences were observed for the treatment response among the male and female obese mice, a pattern observed also for GLP-1 agonists in humans (*39*). Overall, in vivo IgG painting with GLP-1 achieved either similar or better results than the commercial Semaglutide, at a dose 4 times lower (*40, 41*).

Finally, we highlight that in vivo IgG painting can achieve site-selective modification of either one or two lysine side chains, from both sides of the heavy chain Fc domain depending on the electrophile used, making the conjugation of two different drugs possible on the same antibody. The development of long-acting GLP-1 analogs is currently moving towards the combination of two or more incretin co-agonists to boost the effect on T2DM and weight loss (*16, 42*). While the engineering of bispecific antibodies is challenging, our IgG painting platform, which can be performed either in vitro or in vivo, introduces a facile modality to access multi-drug antibody conjugates in the body.

## Acknowledgements

We thank Jacob Rodriguez and Heemal Dhanjee for their discussion on peptide synthesis, purification, and modification. The authors also acknowledge the Koch Institute for Integrative Cancer Research at MIT for technical support, specifically the Preclinical Modeling, Imaging and Testing Core and its director Virginia Spanoudaki.

