## Supplementary Materials for "In vivo Antibody Painting for Next Generation Weight Loss Drugs"

*for*

**Table of Contents**

**Material and Methods:** ... 5

**1. General Information** ... 5

**1.1. Materials** ... 5

**1.2. Nuclear Magnetic Resonance Spectroscopy (NMR)** ... 5

**1.3. Liquid Chromatography-Mass Spectrometry (LC-MS)** ... 6

**1.3.1. LC-MS Using Quadrupole Time-of-Flight (Q-ToF)** ... 6

**1.3.2. LC-MS Using Single Quadrupole Mass Spectrometry** ... 7

**1.3.3. Nano-liquid chromatography-tandem mass spectrometry (nLC-MS/MS)** ... 7

**1.4. Ultra High-Performance Liquid Chromatography (UHPLC)** ... 9

**1.5. Automated Liquid Chromatography Purification** ... 9

**1.6. Sodium Dodecyl Sulfate-polyacrylamide Gel Electrophoresis (SDS-PAGE)** ... 9

**1.7. Determination of the Reaction Conversions of IgGs** ... 9

**2. Chemical Synthesis of IgG Binding Peptide Z33 Analogs** ... 10

**2.1. Automated Flow-based Peptide Synthesis System** ... 10

**2.2. Synthesis of Z33 Variants Composed of Natural Amino Acids** ... 10

**2.3. Synthesis of Z33 variants containing unnatural amino acids** ... 10

**2.4. Capping of the N-terminus of Z33 Variants.** ... 11

**2.5. Cleavage of Z33 Variants from Resin** ... 11

**2.6. Purification of the Z33 Variants from Resin** ... 11

**3. Synthesis of Small Molecule Reagents** ... 12

**3.1. Synthesis of Electrophiles with Allyl Halides** ... 12

**3.2. Synthesis of Palladium Oxidative Addition Complexes** ... 18

**4. Synthesis of Electrophile-attached Z33 Reagents** ... 21

**4.2. Click handle transfer to IgGs using electrophile-Z33 variants.** ... 23

**4.2.1. Bioconjugation to Trastuzumab (IgG1)** ... 23

**4.2.2. Bioconjugation to other IgGs (human IgG1,2,4 & mouse IgG1)** ... 24

**4.3. Determination of the Modification Sites** ... 24

**4.4. Expanding the Chemistry with Other Z33 Variants** ... 25

**4.4.1. Using the Electrophile-attached Z33M3C/Hcy Reagents** ... 25

**4.4.2. Using the Electrophile-attached Z33R31C/Hcy and D32C/Hcy Reagents** ... 25

**4.5. Dual Modification of the Trastuzumab Heavy Chain by Stepwise Reactions** ... 26

**4.6. Trastuzumab Modification in the Presence of RNase A** ... 26

**4.7. Antibody-drug conjugate (ADC) preparation with MMAE transfer reagent** ... 27

**5. Animal Experiments** ... 27

**5.1. Mouse IgG Painting with Azido Moieties** ... 28

**5.1.1. Enzyme-linked Immunosorbent Assay (ELISA)** ... 28

### Material and Methods:

### 1. General Information

#### 1.1. Materials

Fmoc-L-Ala-OH, Fmoc-L-Arg(Pbf)-OH, Fmoc-L-Asn(Trt)-OH, Fmoc-L-Asp-(OtBu)-OH, Fmoc-L-Cys(Trt)-OH, Fmoc-L-Gln(Trt)-OH, Fmoc-L-Glu(OtBu)-OH, Fmoc-L-Gly-OH, Fmoc-L-His(Trt)-OH, Fmoc-L-Ile-OH, Fmoc-L-Leu-OH, Fmoc-L-Lys(Boc)-OH, Fmoc-L-Met-OH, Fmoc-L-Phe-OH, Fmoc-L-Pro-OH, Fmoc-L-Ser(tBu)-OH, Fmoc-L-Thr(tBu)-OH, Fmoc-L-Trp(Boc)-OH, Fmoc-L-Tyr(tBu)-OH, and Fmoc-L-Val-OH were purchased from Novabiochem, Millipore Sigma, or Chem-Impex Inc. Fmoc-alpha-methylalanine (Fmoc-Aib-OH) and Fmoc-L-HomoCys(Trt)-OH (Fmoc-Hcy-OH) were purchased from Combi-Blocks Inc. 5-Azidopentanoic acid was purchased from ChemPep®. 1-(9*H*-Fluoren-9-yl)-3-oxo-2,7,10,13,16,19,22,25,28-nonaoxa-4-azahentriacontan-31-oic acid (Fmoc-PEG8-$CO_2H$) was purchased from AmBeed. *O*-(7-azabenzotriazol-1-yl)-*N*,*N*,*N'*,*N'*-tetramethyluronium hexafluorophosphate (HATU), and (7-azabenzotriazol-1yloxy)tripyrrolidinophosphonium hexafluorophosphate (PyAOP) were purchased from P3 Biosystems. H-Rink Amide-ChemMatrix resin was purchased from PCAS BioMatrix Inc. Polyethylene filter paper for peptide synthesis (0.60 mm thick, pore size 7-12 μm) was purchased from Interstate Specialty Products. *N*,*N*-Dimethylformamide (OmniSolv® for Biosynthesis, DMF, stored with an AldraAmine trapping packet), diethyl ether (≥99.0% stabilized with BHT, $Et_2O$), acetonitrile (≥ 99.9%, $CH_3CN$), dichloromethane (≥ 99.8% stabilized with amylene, $CH_2Cl_2$), 1,4-dioxan (≥ 99.5% stabilized by BHT), 2-propanol (99.9%), 2-methyl-tetrahydrofuran (≥ 99.5% stabilized with BHT, 2-Me-THF), dimethylsulfide (≥ 99.5%, BioUltra for molecular biology, DMSO), and pentane (98% reagent grade) were obtained from Millipore-Sigma. Ethyl acetate (≥99.5%), hexane (≥98.5%, mixture of isomers), methanol (≥99.8%), and acetone (≥ 99.5%) were purchased from VWR. The water used in all reactions involving proteins, in the preparation of buffers, and in the preparation of mobile phases for purification was obtained via filtration of deionized water through a Millipore Sigma Milli-Q Ultrapure Water System. AldraAmine trapping packets and piperidine were purchased from Millipore Sigma. Amine-free DMF refers to DMF stored over AldraAmine trapping packets for a minimum of 24 hours prior to use. LC/MS grade water, acetonitrile, and formic acid were purchased from Thermo Fisher Scientific Inc and were used for liquid chromatography-mass spectrometry (LC-MS). All small compounds were purchased from TCI America, Millipore Sigma, AmBeed, Matrix Scientific, or Combi-Blocks in purities ≥95%, and were used without further purification (unless otherwise noted). PBS buffer (1X and 10X) were purchased from CORNING®. Reactions monitored by analytical thin-layer chromatography (TLC) were carried out using glass-backed plates pre-coated with silica gel impregnated with a fluorescent indicator (254 nm). All deuterium solvents were purchased from Cambridge Isotope Laboratories, Inc. C18-ZipTips (0.6 μL) were purchased from Millipore Sigma. Zeba spin desalting columns were purchased from Thermo Fisher Scientific Inc. Trastuzumab and its biosimilar were purchased from Syd Labs Inc., MedChemExpress LLC., and Bio X Cell.

#### 1.2. Nuclear Magnetic Resonance Spectroscopy (NMR)

NMR spectra were recorded on a Bruker Avance III HD 400 MHz spectrometer. All $^1H$ NMR chemical shifts are expressed in parts per million (ppm, δ scale) and are referenced to the residual proton in the NMR solvent ($CDCl_3$-d3: 7.26, DMSO-d6: 2.50). All $^{13}C$ spectra recorded are proton decoupled. The $^{13}C$ NMR chemical shifts are expressed in part per million (ppm, δ scale) and are referenced to the carbon resonance of the NMR solvent ($CDCl_3$-d3: 77.16, DMSO-d6: 39.52). $^1$H NMR spectroscopic data are reported as follows: a chemical shift in ppm (multiplicity, coupling constants J (Hz), integration intensity, assigned number of protons in molecule). The multiplicities are abbreviated with s (singlet), br. s (broad singlet), d (doublet), t (triplet), q (quartet), and m (multiplet). In the case of combined multiplicities, the multiplicity with the larger coupling constant is stated first. The chemical shift of all signals is reported as the center of the resonance range, except in the case of multiplets, which are reported as ranges in chemical shift. All raw fid files were processed, and the spectra analyzed using the program MestReNova 14.2.3 from Mestrelab Research S. L. Copies of the $^1$H and $^{13}$C spectra can be found at the end of this document (unless otherwise noted).

**1.3. Liquid Chromatography-Mass Spectrometry (LC-MS)**

**1.3.1. LC-MS Using Quadrupole Time-of-Flight (Q-ToF)**

LC-MS chromatograms and associated mass spectra were acquired using an Agilent Technologies 6550 Q-ToF LC-MS system. Solvent compositions are 0.1% formic acid in $H_2O$ (solvent A) and 0.1% formic acid in acetonitrile (solvent B). For methods A, B, and C, a calibration solution containing m/z at 922.0098 constantly flowed through the system. The following LC-MS methods were used:

Q-ToF LC-MS Method A

LC conditions: Zorbax 300SB C3 column: 2.1 × 150 mm, 5 µm, column temperature: 40 °C, gradient: 0-1 min 1% B, 1-6 min 1-61% B, flow rate: 0.8 mL/min. MS conditions: positive electrospray ionization (ESI) extended dynamic mode in mass range 300–3000 m/z, temperature of drying gas = 350 °C, flow rate of drying gas = 11 L/min, pressure of nebulizer gas = 60 psi, the capillary, fragmentor, and octapole voltages were set at 4000, 175, and 750 V, respectively.

Q-ToF LC-MS Method B

LC conditions: Zorbax 300SB C3 column: 2.1 × 150 mm, 5 µm, column temperature: 40 °C, gradient: 0-1 min 1% B, 1-7 min 1-91% B, flow rate: 0.7 mL/min. MS conditions: positive electrospray ionization (ESI) extended dynamic mode in mass range 300–3000 m/z, temperature of drying gas = 350 °C, flow rate of drying gas = 11 L/min, pressure of nebulizer gas = 60 psi, the capillary, fragmentor, and octapole voltages were set at 4000, 175, and 750 V, respectively.

Q-ToF LC-MS Method C

LC conditions: Zorbax 300SB C3 column: 2.1 × 150 mm, 5 µm, column temperature: 40 °C, gradient: 0-1 min 1% B, 1-6 min 1-41% B, flow rate: 0.8 mL/min. MS conditions: positive electrospray ionization (ESI) extended dynamic mode in mass range 300–3000 m/z, temperature of drying gas = 350 °C, flow rate of drying gas = 11 L/min, pressure of nebulizer gas = 60 psi, the capillary, fragmentor, and octapole voltages were set at 4000, 175, and 750 V, respectively.

LC-MS Method D

LC conditions: Aeris WIDEPORE C4 200 column: 2.1 × 150 mm, 3.6 µm, column temperature: 40 °C, gradient: 0-2 min 1% B, 2-8 min 1-91% B, 8-10 min 91-95% B, flow rate: 0.3 mL/min. MS conditions: positive electrospray ionization (ESI) extended dynamic mode in mass range

100–1700 m/z, temperature of drying gas = 200 °C, flow rate of drying gas = 14 L/min, pressure of nebulizer gas = 55 psi, the capillary, fragmentor, and octapole voltages were set at 3500, 175, and 750 V, respectively.

LC-MS Method E

LC conditions: Aeris WIDEPORE C4 200 column: 2.1 × 150 mm, 3.6 µm, column temperature: 40 °C, gradient: 0-2 min 1% B, 2-7 min 1-61% B, 7-8 min 61-95% B, flow rate: 0.3 mL/min. MS conditions: positive electrospray ionization (ESI) extended dynamic mode in mass range 100–1700 m/z, temperature of drying gas = 200 °C, flow rate of drying gas = 14 L/min, pressure of nebulizer gas = 55 psi, the capillary, fragmentor, and octapole voltages were set at 3500, 175, and 750 V, respectively.

Data were processed using Agilent MassHunter Workstation Qualitative Analysis Version B.06.00 Software or Agilent MassHunter BioConfirm Software B.10.00. Deconvoluted masses of proteins were obtained using a maximum entropy algorithm. Unless otherwise depicted, the following parameters were used for deconvolution: Mass range is given in the experimental section for each peptide; Mass step was set to 1.00 Daltons; Baseline was set to Subtract baseline, Baseline factor as 7.00. The y-axis of all chromatograms shown in the supplemental figures represents the total ion current (TIC), and the inset of the mass spectrum corresponds to the deconvolution of the entire protein including peaks.

#### 1.3.2. LC-MS Using Single Quadrupole Mass Spectrometry

Mass spectra were obtained on an Agilent 6125B mass spectrometer attached to an Agilent 1260 Infinity LC. Solvent compositions are 0.1% formic acid in $H_2O$ (solvent A) and 0.1% formic acid in acetonitrile (solvent B). The following LC-MS method was used:

LC conditions: Poroshell 120 SB C18: 2.1 × 50 mm, 2.7 µm, column temperature: 40 °C, gradient: 0-1 min 10% B, 1-5 min 10-100% B, 5-6 min 100% B, 6-7 min 100-10% B, flow rate: 0.4 mL/min. MS conditions: positive electrospray ionization (ESI) extended dynamic mode in mass range 100–1500 m/z, temperature of drying gas = 350 °C, flow rate of drying gas = 13 L/min, pressure of nebulizer gas = 35 psi, the capillary, fragmentor, and octapole voltages were set at 4000, 70, and 650 V, respectively.

#### 1.3.3. Nano-liquid chromatography-tandem mass spectrometry (nLC-MS/MS)

Analysis was performed on an EASY-nLC 1200 nano-liquid chromatography system connected to an Orbitrap Fusion Lumos Tribrid Mass Spectrometer or to an Orbitrap Fusion Eclipse Tribrid Mass Spectrometer (Thermo Fisher Scientific). Samples were run on a PepMap RSLC C18 column (C18, 50 µm x 15 cm, 2 µm, 100 Å, Thermo Fisher Scientific, P/N ES901). An Acclaim PepMap 100 Trap column (C18, 75 µm x 2 cm, 3 µm, 100 Å, Thermo Fisher Scientific, P/N 164946) was used for desalting. Solvent compositions are 0.1% formic acid in $H_2O$ (solvent A) and 0.1% formic acid in 80% acetonitrile and 19.9% $H_2O$ (solvent B). The following conditions were used for each sample measurement:

nLC-MS/MS Method A

LC conditions: column temperature: 40 °C, gradient: 0-30 min 30-95% B, 30-40 min 95% B, flow rate: 300 nL/min. MS conditions: positive ion spray voltage was set to 2200 V. Orbitrap detection was used for primary MS, with the following parameters: Application mode = peptide, cycle time = 3 s, resolution = 120000, mass range = normal, scan range = 250-2000 m/z, RF

Lens = 30%, AGC target = standard, maximum injection time mode = auto, 1 microscan, data type = profile, polarity = positive. The following filters were applied for MS2 precursor selection: monoisotopic peak determination = peptide, filter type = threshold, Intensity threshold = 5.0e5, include charge state = 2-6, mass list type = m/z, target mass*, mass tolerance = 25 ppm. Fragmentation was induced by higher-energy collisional dissociation (HCD) and electron-transfer dissociation with higher-energy collision (EThcD). Specifications HCD: isolation mode = quadrupole, isolation window = 1.2 m/z, isolation offset = 0.6 m/z, activation type = HCD, collision energy mode = fixed (30%), detection type = orbitrap, resolution = 30000, mass range = normal, first mass = 120 m/z, AGC target = standard, maximum injection time mode = dynamic, 1 microscan, data type = profile. Specifications EThcD: isolation mode = quadrupole, isolation window = 1.2 m/z, isolation offset = 0.6 m/z, activation type = ETD, use calibrated charge-dependent ETD parameters, ETD supplemental activation = EThcD, SA collision energy = 30 %, detection = orbitrap, resolution = 30000, mass range = normal, first mass = 120 m/z, normalized AGC target = standard, maximum injection time = dynamic, 1 microscans, data type = profile.

* A list of target masses was created by calculating the intrinsic masses of the measured samples.

nLC-MS/MS Method B

LC conditions: column temperature: 40 °C, gradient: 0-30 min 1-10% B, 30-120 min 10-81% B, 120-125 min 81-90% B, 125-135 min 90% B, flow rate: 300 nL/min. MS conditions: positive ion spray voltage was set to 2200 V, and negative ion spray voltage was set to 600 V. Orbitrap detection was used for primary MS, with the following parameters: Application mode = peptide, cycle time = 3 s, resolution = 120000, mass range = normal, scan range = 200-1400 m/z, RF Lens = 30%, AGC target = custom, normalized AGC target (%) = 250, maximum injection time mode = auto, 1 microscan, data type = profile, polarity = positive. The following filters were applied for MS2 precursor selection: monoisotopic peak determination = peptide, exclude after 4 times, If occurs within 30 s, exclusion duration = 30 s, mass tolerance = 10 ppm, exclude isotope, include charge state = 2-10, filter type = threshold, Intensity threshold = 5.0e4, precursor selection range = 200-1400, Fragmentation was induced by collision-induced dissociation (CID), higher-energy collisional dissociation (HCD), and electron-transfer dissociation with higher-energy collision (EThcD). Specifications CID: isolation mode = quadrupole, isolation window = 1.3 m/z, activation type = CID, collision energy mode = fixed (30%), CID activation time = 10 ms, activation Q = 0.25, detection type = orbitrap, resolution = 30000, mass range = normal, AGC target = custom, normalized AGC target = 40%, maximum injection time mode = auto, 1 microscan, data type = centroid. Specifications HCD: isolation mode = quadrupole, isolation window = 1.3 m/z, activation type = HCD, detection type = orbitrap, resolution = 30000, mass range = normal, scan range mode = auto, AGC target = custom, normalized AGC target = 40%, maximum injection time mode = auto, 1 microscan, data type = centroid. Specifications EThcD: isolation mode = quadrupole, isolation window = 1.3 m/z, activation type = ETD, use calibrated charge-dependent ETD parameters, ETD supplemental activation = EThcD, SA collision energy = 25 %, detection = orbitrap, resolution = 30000, mass range = normal, scan range mode = auto, normalized AGC target = 40%, normalized AGC target = custom, maximum injection time = auto, 1 microscans, data type = profile.

#### 1.4. Ultra High-Performance Liquid Chromatography (UHPLC)

The samples were analyzed using an Agilent Technologies 1290 Infinity II LC system which was computer-controlled through Agilent ChemStation software. The following methods were used:

UHPLC Method (for peptide/protein analysis excluding IgG)

Solvent compositions used in the UHPLC are 0.1% TFA in $H_2O$ (solvent A) and 0.1% TFA in acetonitrile (solvent B). ACQUITY UPLC Protein BEH C4 column: 2.1 × 50 mm, 1.7 μm (Waters, P/N 186004495), ACQUITY UPLC Protein BEH C4 VanGurde pre-column: 2.1 × 5 mm, 1.7 μm (Waters, P/N 186004623), column temperature: 27 °C, gradient: 0-3 min 1% B, 3-13 min 1-61% B, flow rate: 0.5 mL/min.

#### 1.5. Automated Liquid Chromatography Purification

A Biotage® Selekt automated flash chromatography system was used for the purification of small molecules and peptides. The columns, solvents, and solvent gradients employed for the purification of individual compounds are written in the synthesis section of each compound.

#### 1.6. Sodium Dodecyl Sulfate-polyacrylamide Gel Electrophoresis (SDS-PAGE)

Bolt 4-12% Bis-Tris Plus 1.0 mm × 15 well or 1.0 mm × 10 well plates (Invitrogen), Mini Gel Tank (Invitrogen) along with the PowerPac HC (BIO-RAD) were used for SDS-PAGE analysis. SeeBlue® Plus2 standard (Invitrogen) was used as the molecular weight standard. Gels were run using Bolt MOPS SDS Running Buffer (1X, Invitrogen) under the conditions of 135 V for 55 min. After electrophoresis, the running buffer was discarded, and the gel was placed in deionized water. Subsequently, the water with gel was heated in a microwave for 1 min and 30 s. The heated water was replaced, and the gel was thoroughly rinsed. Gel staining was performed for 10 min using SimpleBlue SafeStain (Invitrogen). The stain was removed following staining, and the gel was immersed in deionized water. The tray with gel and deionized water was placed on a shaker overnight to remove the stain from the gels. Then, ChemDoc MP Imaging System (BIO-RAD) was used to analyze the stained gels.

#### 1.7. Determination of the Reaction Conversions of IgGs

The conversion rate for each reaction was calculated by one of the following methods.

Determination by LC-MS

Reported yields based on LC-MS spectra were determined by extracting the total ion current (TIC) spectra of all protein-containing species in the chromatogram utilizing Agilent MassHunter 6.0 software with the BioConfirm package. These extracted chromatograms were deconvoluted utilizing a maximum entropy algorithm and abundance of each species determined using total ion count.

Determination by SDS-PAGE

Band densitometry was calculated using ImageJ software (https://imagej.nih.gov/ij/). Densitometry-based yields were calculated based on the product: starting material protein ratio in lanes and standardized based on molecular weight. An example is shown below.

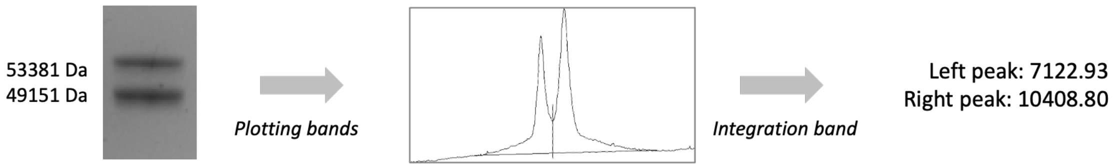

After plotting bands and integration using ImageJ software, the value of each band was obtained as 7122.93 and 10408.80, respectively.

Starting material: right peak standardization (MW 49151) 10408.80/49151 = 0.21177

Modified product: left peak standardization (MW 53381) 7122.93/53381 = 0.13344

Thus, the determined conversion (%) corresponds to Standardized Product / (Standardized Total): 0.13344 / (0.21177 + 0.13344) *100 = 39%

### 2. Chemical Synthesis of IgG Binding Peptide Z33 Analogs

#### 2.1. Automated Flow-based Peptide Synthesis System

The synthesis of the IgG binding peptide Z33 analogs were performed using the Automated fast-flow solid phase peptide synthesis (AFPS) system assembled in the Pentelute lab (*20*, *21*). The synthesis conditions are summarized in the following table.

| Parameter | Conditions |
|---|---|
| temperature | 85-90 °C in reactor, 60 °C in 5' activation loop (for C and H), 90 °C in 10' activation loop (for all other amino acids) |
| Flow Rate | 40 mL/min |
| Coupling Step | 0.40 M amino acids stocks in DMF<br>0.38 M activator stocks in DMF<br>Coupling conditions: HATU (13 pump strokes) except S and A with HATU (26 pump strokes) and C, H, N, Q, V, R, T with PyAOP (26 pump strokes) |
| Deprotection Step | 40% piperidine in DMF with 2% formic acid (13 pump strokes) |
| Washing Steps | DMF (40 pump strokes) |

#### 2.2. Synthesis of Z33 Variants Composed of Natural Amino Acids

A plastic fritted syringe (6 mL) was equipped with polyethylene filter paper, and ChemMatrix® H-Rink Amide resin (0.49 mmol/g, 150 mg) was loaded onto it. The syringe assembly was then placed on a manifold. The resin was swollen in DMF, and subsequently, a homogeneous resin slurry was prepared by repeatedly drawing back 500 µL of the slurry using a 1 mL pipette. Next, the syringe was set to the AFPS system for peptide synthesis. After completion of the synthesis, the syringe was transferred from the AFPS system to the manifold and washed three times with $CH_2Cl_2$. To minimize methionine oxidation in the peptide sequence, drying of the resin was not conducted using air flow on the manifold but rather using nitrogen flow. Specifically, the syringe was capped with a rubber septum, and a needle connected to a nitrogen flow line was inserted into the septum for 15 min. The resulting peptide-bound dry resin was subjected to subsequent steps of cleavage and purification.

#### 2.3. Synthesis of Z33 variants containing unnatural amino acids

The peptide was synthesized using the method described in **Section 2.2** from the C-terminus of the sequence to the position of the unnatural amino acid. The resin in a plastic fritted syringe was transferred from the AFPS system to a manifold. Fmoc-protected unnatural amino acid (10 equivalents for the peptide on the resin) in HATU/DMF (0.38 M, 9.5 equivalents) solution and DIEA (15 equivalents) were added to the resin and allowed to react for 50 min at room temperature. The reaction solution was stirred with a spatula every 10 min. After the reaction, the reaction solution was removed, and the resin in the syringe was washed three times with 5 mL of DMF. Then, 3 mL of 40% piperidine/DMF solution (+2% formic acid) was added and allowed to react for 10 min. The reaction solution was removed, and 3 mL of 40% piperidine/DMF solution (+2% formic acid) was added again and allowed to react for 10 min. The reaction solution was removed, and the resin was washed three times with 4 mL of DMF. Addition of the subsequent natural amino acids to the peptide chain was again performed according to the procedure described in **Section 2.2**.

**2.4. Capping of the N-terminus of Z33 Variants.**

After synthesis of the peptide by AFPS, the fritted syringe with resin was placed on a manifold. A HATU/DMF solution (0.38 M, 9.5 equivalents) of the carboxylic acid corresponding to the acyl group (10 equivalents for the peptide on the resin) and DIEA (15 equivalents) were added and allowed to react for 50 min at room temperature. During that time, the reaction solution was stirred with a spatula every 10 min. After the reaction, the reaction solution was removed, and the resin in the syringe was washed three times with 10 mL of DMF. If the introduced acyl group contained the Fmoc protecting group, 3 mL of 40% piperidine/DMF solution (+2% formic acid) was added and allowed to react for 10 min. The reaction solution was removed, and 3 mL of 40% piperidine/DMF solution (+2% formic acid) was added again and allowed to react for 10 min. The reaction solution was removed and washed three times with 4 mL of DMF and three times with $CH_2Cl_2$. To minimize methionine oxidation in the peptide sequence, the resin was not dried using airflow on the manifold but rather using nitrogen flow. Specifically, the syringe was capped with a rubber septum, and a needle connected to a nitrogen flow line was inserted into the septum for 15 min. The resulting peptide-bound dry resin was subjected to subsequent steps of cleavage and purification.

**2.5. Cleavage of Z33 Variants from Resin**

The peptide-bound resin was transferred to a 15 mL centrifuge tube. Subsequently, a 7.5 mL peptide cleavage solution (a mixture of TFA, water, EDT, and TIPS in a ratio of 94:2.5:2.5:1 by volume) was added to the centrifuge tube, and the reaction was allowed to proceed at room temperature for 2 hours on the shaker. Afterward, 5 mL of a plastic fritted syringe was placed on top of a 50 mL centrifuge tube, and the reaction solution containing the resin was filtered. The resin inside the syringe was washed twice with 3 mL of TFA. To the resulting solution, diethyl ether pre-cooled at −80 °C was added until a total volume of 45 mL was reached. The centrifuge tube was then capped, and the reaction solution was vortexed for 5 s, followed by centrifugation at 3220 rcf (relative centrifugal force) for 4 min. Subsequently, the supernatant was removed by decanting. The obtained crude peptide was dissolved in 10 mL of 50:50 water and acetonitrile solution (+0.1% TFA), followed by freezing the solution with liquid nitrogen and subjecting it to lyophilization. When the peptide contained the azido moiety, another peptide cleavage solution (a mixture of TFA, water, thioanisole, and TIPS in a ratio of 94:2.5:2.5:1 by volume) was used instead of the solution described above.

**2.6. Purification of the Z33 Variants from Resin**

The crude Z33 analog was dissolved in a 95:5 water-acetonitrile solution and loaded onto 12 g of Biotage® Sfär C18 column. The gradient used for purification was as follows: 3 CV (column volume) 5% B, 1 CV 5-10% B, 30 CV 10-40% B. Fractions containing the desired product were collected in 50 mL centrifuge tubes, and the solution was frozen using liquid nitrogen. The target Z33 variant was obtained as white solids by removing the solvent through lyophilization.

| Code | N-term | Sequence | C-term | Calculated Mass (Da) | Observed Mass (Da) |
|---|---|---|---|---|---|
| **1** | Free | FNMQQQRRFYEALHDpNLNCEQRNAKIKSIRDD | Amide | 4078.6 | 4078.6 |
| **2** | Free | FNMQQQRRFYEALHDPNLNHcyEQRNAKIKSIRDD | Amide | 4092.6 | 4093.1 |
| **3** | 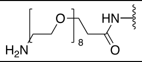 | FNMQQQRRFYEALHDPNLNHcyEQRNAKIKSIRDD | Amide | 4516.1 | 4516.2 |
| **4** | Free | FNMQQQRRFYEALHDPNLNHcyEQRNAKIKSIRDd | Amide | 4092.6 | 4092.5 |
| **5** | Free | fNMQQQRRFYEALHDPNLNHcyEQRNAKIKSIRDd | Amide | 4092.6 | 4092.5 |
| **6** | Free | fNMQQQRRFYEALHDPNLNHcyEQRNAKIKSIRdd | Amide | 4092.6 | 4092.6 |
| **BB-1** | Free | FNMQQQRRFYEALHDPNLNCEQRNAKIKSIRDD | Amide | 4078.6 | 4078.5 |
| **BB-2** | Free | FNCQQQRRFYEALHDPNLNEEQRNAKIKSIRDD | Amide | 4076.5 | 4077.1 |
| **BB-3** | Free | FNHcyQQQRRFYEALHDPNLNEEQRNAKIKSIRDD | Amide | 4090.5 | 4090.7 |
| **BB-4** | Free | FNMQQQRRFYEALHDPNLNEEQRNAKIKSICDD | Amide | 4051.5 | 4052.3 |
| **BB-5** | Free | FNMQQQRRFYEALHDPNLNEEQRNAKIKSIRCD | Amide | 4092.6 | 4092.7 |
| **BB-6** | Free | FNMQQQRRFYEALHDPNLNEEQRNAKIKSIRHcyD | Amide | 4106.6 | 4106.5 |

### 3. Synthesis of Small Molecule Reagents

#### 3.1. Synthesis of Electrophiles with Allyl Halides

*Synthesis of* ***BB-7***

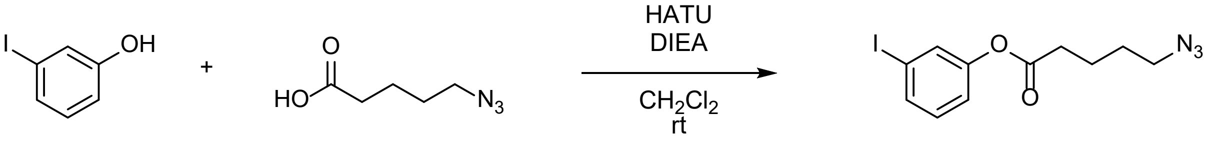

**BB-7**

A 100 mL round-bottom flask equipped with a stir bar was charged with 3-iodophenol (400 mg, 1.82 mmol), followed by the addition of 8 mL of $CH_2Cl_2$ to provide a clear solution. Next, 5-azidopentanoic acid (200 mg, 1.40 mmol), DIEA (0.973 µL, 5.59 mmol), and HATU (797 mg, 2.10 mmol) were added, and the resulting mixture was stirred at rt for 48 hours. The resulting mixture was quenched with saturated $NaHCO_3$ solution (100 mL) and partitioned between $CH_2Cl_2$ (200 mL) and water (100 mL). The organic phase was collected, and the aqueous phase was extracted with $CH_2Cl_2$ (100 mL) twice. The combined organic extract was washed with brine (50 mL), dried over sodium sulfate, filtered, and concentrated under reduced pressure. The obtained residue was purified by column chromatography (using Biotage® Sfär Silica 60 µm 25 g column, 95:5 to 3:1 hexane:ethyl acetate v/v) and dried under reduced pressure, yielding the desired compound as a clear oil (269 mg, 78%).

$^1$H NMR (400 MHz, $CDCl_3$): δ 7.57 (dt, J = 7.5, 1.5 Hz, 1H), 7.46 (t, J = 1.8 Hz, 1H), 7.15 – 7.03 (m, 2H), 3.35 (t, J = 6.6 Hz, 2H), 2.60 (t, J = 7.3 Hz, 2H), 1.90 – 1.78 (m, 2H), 1.79 – 1.66 (m, 2H).

$^{13}C$ NMR (101 MHz, $CDCl_3$) δ 171.27, 150.95, 135.04, 130.81, 130.79, 121.19, 93.62, 51.08, 33.69, 2

HRMS (DART) *m/z* calcd for $C_{11}H_{13}O_2N_3I$ ($[M+H]^+$) 346.0047, found 346.0052.

*Synthesis of* ***BB-8***

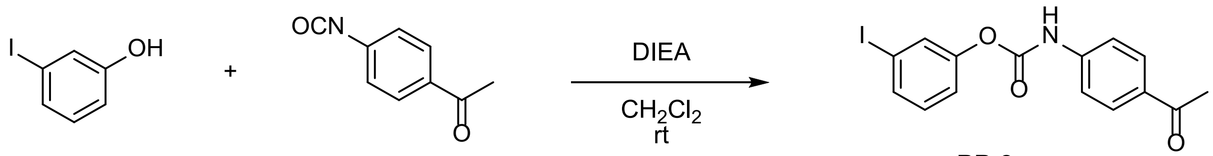

**BB-8**

A 50 mL round-bottom flask equipped with a stir bar was charged with 3-iodophenol (324 mg, 1.47 mmol), followed by the addition of 7 mL of $CH_2Cl_2$ to provide a clear solution. Next, 4-acetylphenyl isocyanate (250 mg, 1.55 mmol) and DIEA (330 µL, 1.86 mmol) were added, and the resulting mixture was stirred at rt for 1 hour. The resulting mixture was then treated with diethyl ether (4 mL) and subjected to sonication for approximately 30 s. The reaction solution was filtered using a Büchner funnel, and the resulting solid was washed with dichloromethane (1 mL). Finally, the solid was dried under reduced pressure, yielding the desired compound as a white solid (241 mg, 43%).

$^1H$ NMR (400 MHz, DMSO): δ 10.68 (s, 1H), 7.95 (d, J = 8.8 Hz, 2H), 7.72 – 7.59 (m, 4H), 7.34 – 7.20 (m, 2H), 2.53 (s, 3H).

*Synthesis of* ***BB-9***

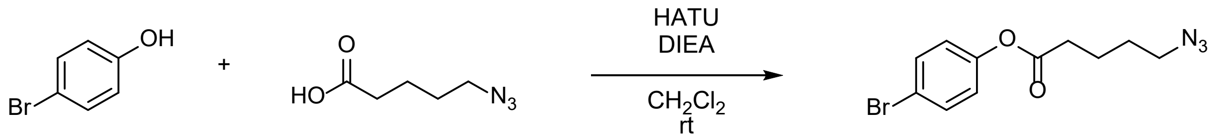

**BB-9**

A 100 mL round-bottom flask equipped with a stir bar was charged with 4-bromophenol (314 mg, 1.82 mmol), followed by the addition of 8 mL of dioxane to provide a clear solution. Next, 5-azidopentanoic acid (200 mg, 1.40 mmol), DIEA (0.973 µL, 5.59 mmol), and HATU (797 mg, 2.10 mmol) were added, and the resulting mixture was stirred at rt for 48 hours. The resulting mixture was quenched with saturated $NaHCO_3$ solution (100 mL) and partitioned between $CH_2Cl_2$ (200 mL) and water (100 mL). The organic phase was collected, and the aqueous phase was extracted with $CH_2Cl_2$ (100 mL) twice. The combined organic extract was washed with brine (50 mL), dried over sodium sulfate, filtered, and concentrated under reduced pressure. The obtained residue was purified by column chromatography (using Biotage® Sfär Silica 60 µm 25 g column, 95:5 to 3:1 hexane:ethyl acetate) and dried under reduced pressure, yielding the desired compound as a clear oil (338 mg, 81%).

$^1H$ NMR (400 MHz, $CDCl_3$): δ 7.49 (d, J = 8.8 Hz, 2H), 6.97 (d, J = 8.8 Hz, 2H), 3.35 (t, J = 6.6 Hz, 2H), 2.60 (t, J = 7.3 Hz, 2H), 1.90 – 1.78 (m, 2H), 1.77 – 1.67 (m, 2H).

$^{13}C$ NMR (101 MHz, $CDCl_3$) δ 171.41, 149.74, 132.59, 123.44, 119.02, 51.13, 33.78, 28.34, 22.13.

HRMS (DART) *m/z* calcd for $C_{11}H_{13}O_2N_3Br$ ($[M+H]^+$) 298.0186, found 298.0183.

*Synthesis of* ***BB-10***

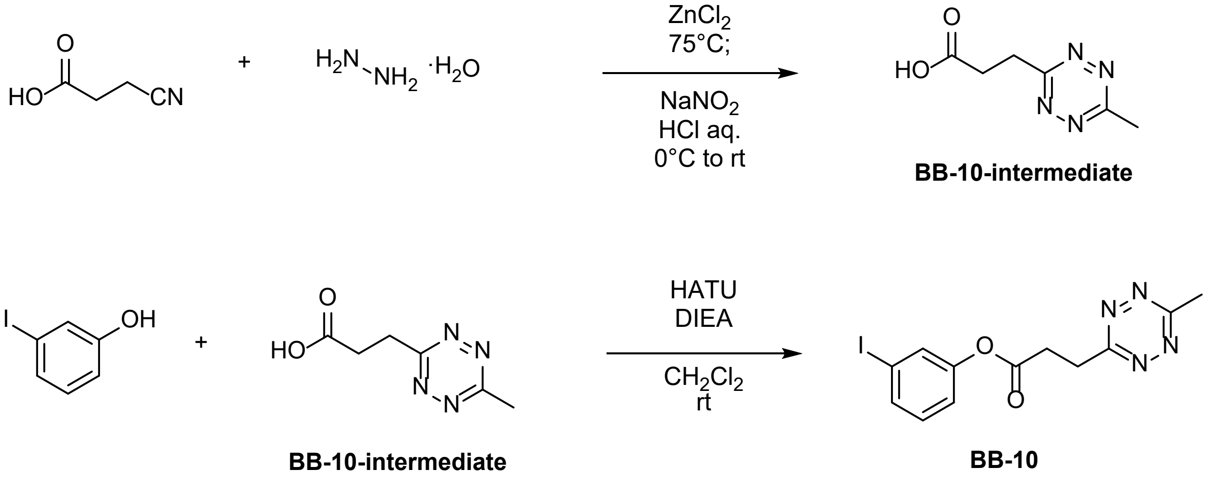

The **BB-10-intermediate** was synthesized as previously described (WO2014065860). The 1H NMR spectra of the obtained material were identical to those reported in the literature.

A 20 mL scintillation glass vial equipped with a stir bar was charged with 3-iodophenol (45.4 mg, 0.309 mmol), followed by the addition of 2 mL of $CH_2Cl_2$ to provide a clear solution. Next, **BB-10-intermediate** (40.0 mg, 0.238 mmol), DIEA (104 µL, 0.952 mmol), and HATU (90.5 mg, 0.357 mmol) were added, and the resulting mixture was stirred at rt for 12 hours. The resulting mixture was directly purified by column chromatography (using Biotage® Sfär Silica 60 µm 10 g column, 80:20 to 1:1 hexane:ethyl acetate) and dried under reduced pressure, yielding the desired compound as a pink oil (50 mg, 57%).

$^1H$ NMR (400 MHz, $CDCl_3$): δ 7.57 (ddd, J = 6.0, 3.1, 1.6 Hz, 1H), 7.50 – 7.44 (m, 1H), 7.14 – 7.05 (m, 2H), 3.75 (t, J = 6.9 Hz, 2H), 3.31 (t, J = 6.9 Hz, 2H), 3.06 (s, 3H).

*Synthesis of* ***BB-11***

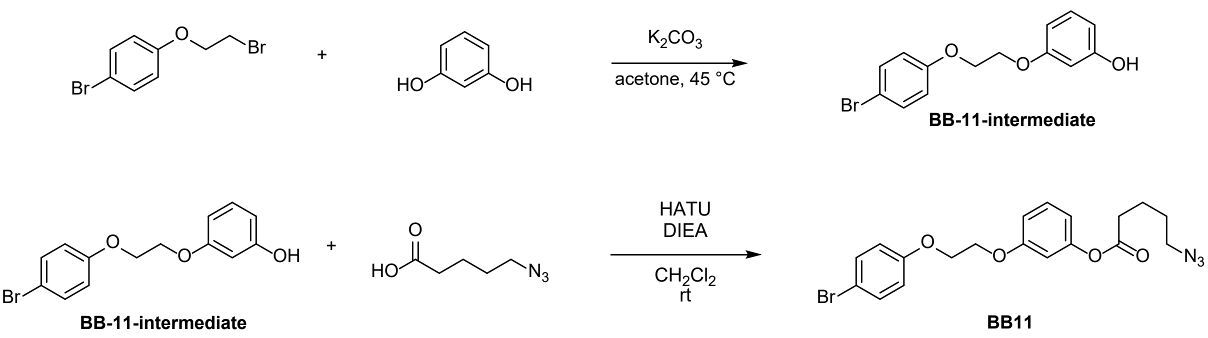

A 50 mL round-bottom flask equipped with a stir bar was charged with 1-bromo-4-(2-bromoethoxy)benzene (600 mg, 2.14 mmol), followed by the addition of 10 mL of acetone to provide a clear solution. Next, resorcinol (1.18 g, 10.7 mmol) and $K_2CO_3$ (1.48 g, 10.7 mmol) were added, and the resulting mixture was stirred at 45 °C for 12 hours. The resulting mixture was cooled to room temperature, followed by filtration using a Büchner funnel. The collected solid was then washed once with acetone (5 mL), and the resulting solution was concentrated under reduced pressure. The obtained residue was purified by column chromatography (using Biotage® Sfär Silica 60 µm 25 g column, 95:5 to 3:1 hexane:ethyl acetate) and dried under reduced pressure, yielding the **BB-11-intermediate** as a clear oil (415 mg, 63%).

$^{1}$H NMR (400 MHz, $CDCl_3$): δ 7.39 (d, J = 9.0 Hz, 2H), 7.14 (t, J = 8.5 Hz, 1H), 6.83 (d, J = 9.0 Hz, 2H), 6.57 – 6.49 (m, 1H), 6.47 – 6.43 (m, 2H), 4.71 (s, 1H), 4.28 (s, 4H).

$^{13}$C NMR (101 MHz, $CDCl_3$) δ 160.05, 157.90, 156.85, 132.46, 130.38, 129.66, 116.68, 113.49, 108.43, 108.06, 107.22, 102.49, 66.89, 66.57.

A 20 mL scintillation glass vial equipped with a stir bar was charged with **BB-11-intermediate** (150 mg, 0.485 mmol), followed by the addition of 2 mL of $CH_2Cl_2$ to provide a clear solution. Next, 5-azidopentanoic acid (63.1 mg, 0.441 mmol), DIEA (300 µL, 1.76 mmol), and HATU (252 mg, 0.662 mmol) were added, and the resulting mixture was stirred at rt for 12 hours. The resulting mixture was directly purified by column chromatography (using Biotage® Sfär Silica 60 µm 10 g column, 80:20 to 1:1 hexane:ethyl acetate) and dried under reduced pressure, yielding the desired compound as a pale pink solid (116 mg, 61%).

$^{1}$H NMR (400 MHz, $CDCl_3$): δ 7.39 (d, J = 9.0 Hz, 2H), 7.31 – 7.26 (m, 1H), 6.87 – 6.78 (m, 3H), 6.74 – 6.66 (m, 2H), 4.29 (s, 4H), 3.35 (t, J = 6.7 Hz, 2H), 2.60 (t, J = 7.2 Hz, 2H), 1.91 – 1.79 (m, 2H), 1.78 – 1.67 (m, 2H).

*Synthesis of **BB-12***

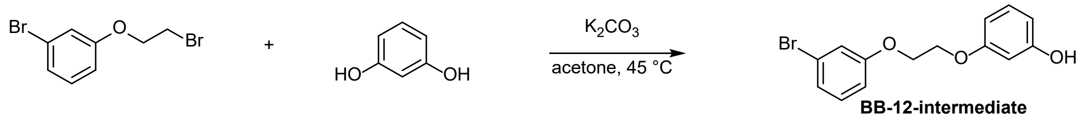

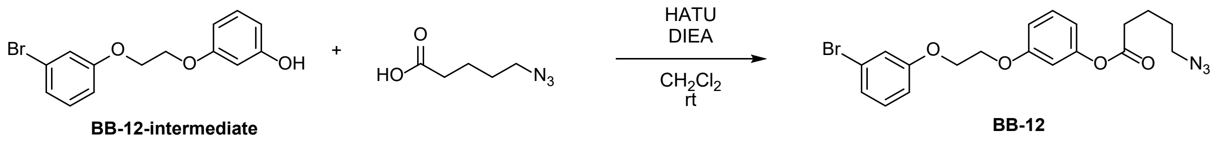

A 50 mL round-bottom flask equipped with a stir bar was charged with 1-bromo-3-(2-bromoethoxy)benzene (600 mg, 2.14 mmol), followed by the addition of 10 mL of acetone to provide a clear solution. Next, resorcinol (1.18 g, 10.7 mmol) and $K_2CO_3$ (1.48 g, 10.7 mmol) were added, and the resulting mixture was stirred at 45 °C for 12 hours. The resulting mixture was cooled to room temperature, followed by filtration using a Büchner funnel. The collected solid was then washed once with acetone (5 mL), and the resulting solution was concentrated under reduced pressure. The obtained residue was purified by column chromatography (using Biotage® Sfär Silica 60 µm 25 g column, 95:5 to 3:1 hexane:ethyl acetate) and dried under reduced pressure, yielding **BB-12-intermediate** as a clear oil (415 mg, 63%).

$^{1}$H NMR (400 MHz, $CDCl_3$): δ 7.20 – 7.07 (m, 4H), 6.91 – 6.86 (m, 1H), 6.53 (ddd, J = 8.3, 2.3, 1.0 Hz, 1H), 6.48 – 6.42 (m, 2H), 4.72 (s, 1H), 4.29 (s, 4H).

$^{13}C$ NMR (101 MHz, $CDCl_3$) δ 160.01, 159.52, 156.86, 130.72, 130.37, 124.38, 122.95, 118.13, 113.89, 108.44, 107.18, 102.49, 66.85, 66.50.

HRMS (ESI) *m/z* calcd for $C_{14}H_{14}O_3Br$ ($[M+H]^+$) 309.0126, found 309.0133.

A 20 mL scintillation glass vial equipped with a stir bar was charged with **BB-12-intermediate** (75 mg, 0.242 mmol), followed by the addition of 2 mL of $CH_2Cl_2$ to provide a clear solution. Next, 5-azidopentanoic acid (52.0 mg, 0.363 mmol), DIEA (158 µL, 0.968 mmol), and HATU (184 mg, 0.484 mmol) were added, and the resulting mixture was stirred at rt for 12 hours. The resulting mixture was directly purified by column chromatography (using Biotage® Sfär Silica 60 µm 10 g column, 80:20 to 1:1 hexane:ethyl acetate) and dried under reduced pressure, yielding the desired compound as a pale pink solid (74 mg, 70%).

$^1H$ NMR (400 MHz, $CDCl_3$): δ 7.29 (t, J = 8.1 Hz, 1H), 7.20 – 7.09 (m, 3H), 6.92 – 6.79 (m, 2H), 6.75 – 6.66 (m, 2H), 3.35 (t, J = 6.7 Hz, 2H), 2.60 (t, J = 7.2 Hz, 2H), 1.91 – 1.79 (m, 2H), 1.78 – 1.68 (m, 2H).

$^{13}C$ NMR (101 MHz, $CDCl_3$) δ 171.51, 159.40, 159.35, 151.58, 130.62, 129.95, 124.28, 122.83, 117.98, 114.29, 113.73, 112.28, 108.44, 66.64, 66.57, 51.06, 33.75, 28.26, 22.10.

HRMS (ESI) *m/z* calcd for $C_{19}H_{21}O_4N_3Br$ ($[M+H]^+$) 434.0715, found 434.0673.

*Synthesis of* ***BB-13***

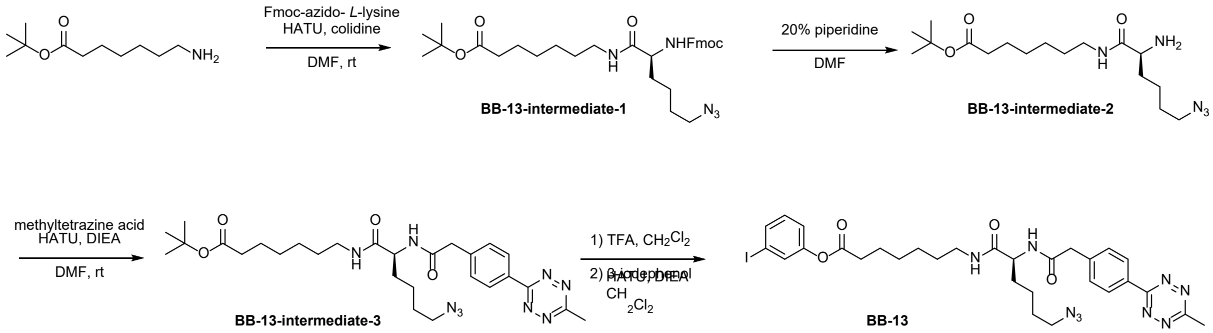

A 50 mL round-bottom flask equipped with a stir bar was charged with 7-amino-heptanoic acid *t*-butyl ester (346 mg, 1.72 mmol), followed by the addition of 9 mL of DMF to provide a clear solution. Next, Fmoc-L-azidolysine (746 mg, 1.89 mmol), 2,4,6-colidine (0.909 µL, 6.84 mmol), and HATU (781 mg, 2.06 mmol) were added, and the resulting mixture was stirred at rt for 12 hours. The resulting mixture was quenched with saturated $NaHCO_3$ solution (100 mL) and partitioned between ethyl acetate (200 mL) and water (100 mL). The organic phase was collected, and the aqueous phase was extracted with ethyl acetate (100 mL) twice. The combined organic extract was washed with saturated ammonium chloride solution (100 mL) twice and brine (50 mL), dried over sodium sulfate, filtered, and concentrated under reduced pressure. The obtained residue was purified by column chromatography (using Biotage® Sfär Silica 60um 25g column, 95:5 to 2:1 hexane:ethyl acetate) and dried under reduced pressure, yielding **BB-13-intermediate-1** as a clear oil (868 mg, 87%).

$^{1}H$ NMR (400 MHz, $CDCl_3$): δ 7.77 (d, J = 7.5 Hz, 2H), 7.58 (d, J = 7.5 Hz, 2H), 7.41 (t, J = 7.5 Hz, 2H), 7.32 (t, J = 7.4 Hz, 2H), 5.90 (s, 1H), 5.32 (d, J = 8.3 Hz, 1H), 4.42 (s, 2H), 4.21 (t, J = 6.7 Hz, 1H), 3.34 – 3.17 (m, 4H), 2.19 (t, J = 7.4 Hz, 2H), 1.57 (s, 8H), 1.43 (s, 9H), 1.43 – 1.40 (m, 2H), 1.31 (s, 4H).

$^{13}C$ NMR (101 MHz, $CDCl_3$) δ 173.28, 171.35, 156.33, 143.85, 141.45, 127.89, 127.21, 125.13, 120.14, 80.19, 67.13, 54.94, 51.26, 47.28, 39.62, 35.52, 32.38, 29.38, 28.69, 28.63, 28.24, 26.60, 24.99, 22.80.

HRMS (ESI) *m/z* calcd for $C_{32}H_{44}O_5N_5$ ($[M+H]^+$) 578.3342, found 578.3315.

A 50 mL round-bottom flask equipped with a stir bar was charged with **BB-13-intermediate-1** (400 mg, 0.69 mmol), followed by the addition of piperidine in DMF (3.4 mL, 20 v/v%) to provide a clear solution. The resulting mixture was stirred at rt for 2 hours. The resulting mixture was diluted with 20 mL of toluene and concentrated under reduced pressure. To remove DMF, 20 mL of toluene was added to the reaction mixture, and the solvent was subsequently evaporated under reduced pressure three times. The obtained residue was purified by column chromatography (using Biotage® Sfär Silica 60 µm 25 g column, 100:0 to 95:5 $CH_2Cl_2$ + 0.5 v/v% trimethylamine:MeOH) and dried under reduced pressure, yielding **BB-13-intermediate-2** as a clear oil (162 mg, 66%).

$^{1}H$ NMR (400 MHz, $CDCl_3$): δ 7.30 (s, 1H), 3.43 (dd, J = 7.7, 4.8 Hz, 1H), 3.34 – 3.19 (m, 4H), 2.20 (t, J = 7.5 Hz, 2H), 2.11 – 1.69 (m, 4H), 1.69 – 1.44 (m, 8H), 1.44 (s, 9H), 1.33 (p, J = 3.5 Hz, 4H).

$^{13}C$ NMR (101 MHz, $CDCl_3$) δ 173.68, 173.41, 80.22, 54.89, 51.30, 39.22, 35.58, 34.12, 29.46, 28.79, 28.75, 28.24, 26.71, 25.05, 22.94.

HRMS (ESI) *m/z* calcd for $C_{17}H_{34}O_3N_5$ ($[M+H]^+$) 356.2662, found 356.2644.

A 20 mL scintillation glass vial equipped with a stir bar was charged with **BB-13-intermediate-2** (93 mg, 0.261 mmol), followed by the addition of 2 mL of $CH_2Cl_2$ to provide a clear solution. Next, methyltetrazine acid (50 mg, 0.217 mmol), DIEA (154 µL, 0.868 mmol), and HATU (124 mg, 0.326 mmol) were added, and the resulting mixture was stirred at rt for 12 hours. The resulting mixture was directly purified by column chromatography (using Biotage® Sfär Silica 60 µm 10 g column, 80:20 to 1:1 hexane:ethyl acetate) and dried under reduced pressure, yielding **BB-13-intermediate-3** as a pale pink solid (101 mg, 82%).

$^{1}H$ NMR (400 MHz, $CDCl_3$): δ 8.58 (d, J = 8.4 Hz, 2H), 7.50 (d, J = 8.2 Hz, 2H), 6.19 (d, J = 8.0 Hz, 1H), 5.98 (s, 1H), 4.36 (q, J = 7.2 Hz, 1H), 3.68 (s, 2H), 3.23 (dq, J = 7.3, 4.9 Hz, 4H), 3.10 (s, 3H), 2.19 (t, J = 7.4 Hz, 2H), 1.90 – 1.77 (m, 1H), 1.66 – 1.41 (m, 7H), 1.44 (s, 9H), 1.40 – 1.24 (m, 6H).

$^{13}C$ NMR (101 MHz, $CDCl_3$) δ 173.24, 171.27, 170.37, 167.39, 163.89, 139.63, 130.95, 130.26, 128.43, 80.18, 53.21, 51.21, 43.46, 39.62, 35.49, 32.22, 29.32, 28.70, 28.63, 28.24, 26.62, 24.98, 22.76, 21.28.

HRMS (ESI) *m/z* calcd for $C_{28}H_{42}O_4N_9$ ($[M+H]^+$) 568.3360, found 568.3334.

A 10 mL round-bottom flask equipped with a stir bar was charged with **BB-13-intermediate-3** (30 mg, 0.053 mmol), followed by the addition of dichloromethane (2 mL) to provide a clear solution. Next, TFA (0.4 mL) was added, and the resulting mixture was stirred at rt for 3 hours. The reaction mixture was concentrated under reduced pressure, and the resulting residue was used in the next step without further purification.

Dichloromethane (1 mL) was added to a 10 mL round-bottom flask containing the residue obtained from the previous reaction, resulting in a clear solution. Next, 3-iodophenol (17.5 mg, 0.080 mmol), DIEA (55.0 µL, 0.318 mmol), and HATU (34.3 mg, 0.090 mmol) were added, and the resulting mixture was stirred at rt for 12 hours. The resulting mixture was directly purified by column chromatography (using Biotage® Sfär Silica 60 µm 10 g column, 80:20 to 1:1 hexane:ethyl acetate v/v) and dried under reduced pressure, yielding **BB-13** as a pale pink solid (35.9 mg, 95%).

$^{1}$H NMR (400 MHz, $CDCl_3$) δ 8.58 (d, J = 8.1 Hz, 2H), 7.56 (d, J = 7.5 Hz, 1H), 7.53 – 7.42 (m, 3H), 7.14 – 7.02 (m, 2H), 6.15 (d, J = 8.0 Hz, 1H), 5.97 (t, J = 6.0 Hz, 1H), 4.35 (q, J = 7.2 Hz, 1H), 3.67 (s, 2H), 3.23 (dq, J = 7.0, 4.5 Hz, 4H), 3.09 (s, 3H), 2.80 (s, 1H), 2.53 (t, J = 7.4 Hz, 2H), 1.84 (dq, J = 14.2, 7.3 Hz, 1H), 1.72 (p, J = 7.3 Hz, 2H), 1.65 – 1.55 (m, 2H), 1.56 – 1.45 (m, 3H), 1.47 – 1.23 (m, 6H).

$^{13}$C NMR (101 MHz, $CDCl_3$) δ 171.84, 171.25, 170.38, 167.43, 163.90, 151.08, 139.55, 135.04, 131.03, 130.90, 130.82, 130.27, 128.50, 121.30, 93.65, 53.26, 51.22, 43.53, 39.57, 38.75, 34.18, 32.11, 29.79, 29.34, 28.70, 28.62, 26.56, 24.73, 22.79, 21.30.

HRMS (ESI) *m/z* calcd for $C_{30}H_{37}O_4N_9I$ ($[M+H]^+$) 714.2013, found 714.2008.

### 3.2. Synthesis of Palladium Oxidative Addition Complexes

*Synthesis of [(cod)Pd(CH₂TMS)₂ ] (cod = 1,5 cyclooctadiene)*

This compound was synthesized as previously described (*28*). The $^{1}$H and $^{13}$C NMR spectra of the obtained material were identical to those reported in the literature.

*Synthesis of sodium 2'-dicyclohexylphosphino-2,6-dimethoxy-1,1'-biphenyl-3-sulfonate hydrate (sSPhos)*

This compound was synthesized as previously described (*43*). The $^{1}$H and $^{13}$C NMR spectra of the obtained material were identical to those reported in the literature.

*Synthesis of **BB-14***

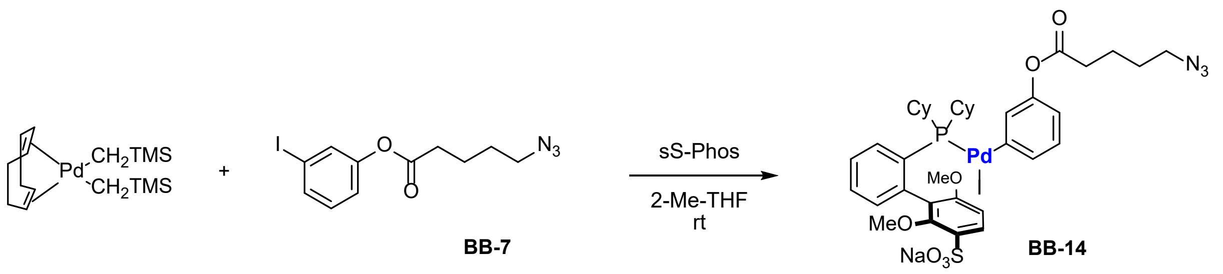

A 5 mL scintillation glass vial equipped with a stir bar was charged with **BB-7** (58.8 mg, 0.171 mmol) and sSPhos (82.8 mg, 0.162 mmol), and 2-methyltetrahydrofuran (0.51 mL). The reaction solution was sonicated for 1 min until it became completely clear, and then the [(cod)Pd($CH_2TMS$)$_2$ ] (60.0 mg, 0.154 mmol) was added, and the resulting mixture was stirred at rt for 2 hours. After the reaction time, pentane (1 mL) was added to the reaction mixture. The vial containing the white suspension was placed in a centrifuge and centrifuged at 3220 rcf for 2 min. The cap was removed, and the supernatant was decanted. The stir bar was removed, and the solid was resuspended in 2-methyltetrahydrofuran (1.0 mL) and pentane (1.0 mL). The vial was capped and sonicated for 1 min to obtain a homogeneous suspension. The vial was then centrifuged again at 3220 rcf for 2 min. The cap was removed, and the supernatant was decanted. This sonication/centrifugation/decanting procedure was repeated twice. The resulting beige solid was dried under reduced pressure to give the desired crude product as a light brown solid (111 mg, 75%). The obtained crude product was used in the following reaction without further purification.

MS (m/z): $C_{37}H_{47}N_3O_7PPdS^+$ ($[M\text{-}I\text{-}Na+H]^+$) 814.2, found 814.1.

*Synthesis of* ***BB-15***

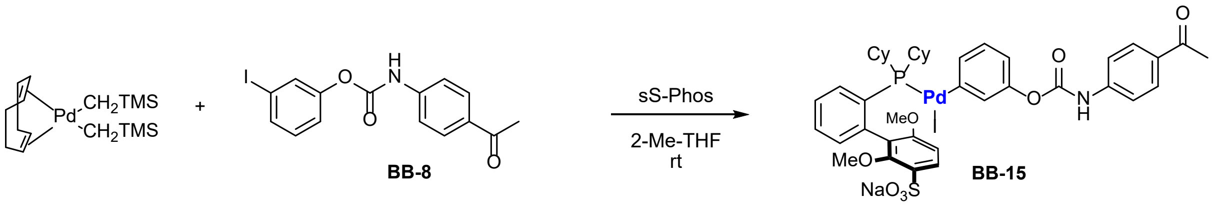

This compound was synthesized following the general procedure described in the section on the synthesis of **BB-14**. **BB-8** (16.3 mg, 0.043 mmol), sSPhos (20.7 mg, 0.040 mmol), [(cod)Pd($CH_2TMS$)$_2$ ] (15.0 mg, 0.039 mmol), and 2-methyltetrahydrofuran (0.5 mL) were used, and **BB-15** was obtained as a light brown solid (39 mg, quantitative).

MS (m/z): $C_{41}H_{47}NO_8PPdS^+$ ($[M\text{-}I\text{-}Na+H]^+$) 850.2, found 850.1.

*Synthesis of* ***BB-16***

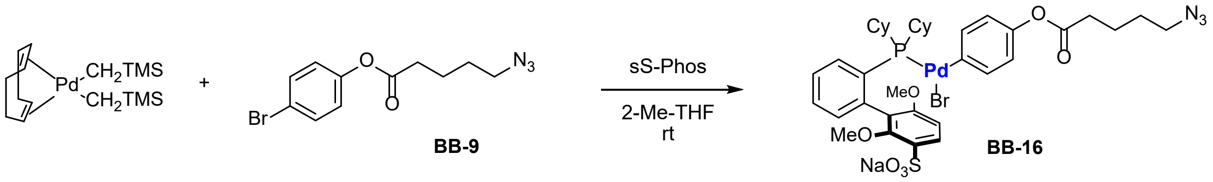

This compound was synthesized following the general procedure described in the section on the synthesis of **BB-14**. **BB-9** (12.7 mg, 0.043 mmol), sSPhos (20.7 mg, 0.040 mmol), [(cod)Pd(CH2TMS)2 ] (15.0 mg, 0.039 mmol), and 2-methyltetrahydrofuran (0.5 mL) were used, and **BB-16** was obtained as a light brown solid (34 mg, 96%).

MS (m/z): $C_{37}H_{47}NO_7PPdS^+$ ($[M\text{-}N_2\text{-}Br\text{-}Na+H]^+$) 786.2, found 786.1.

*Synthesis of* ***BB-17***

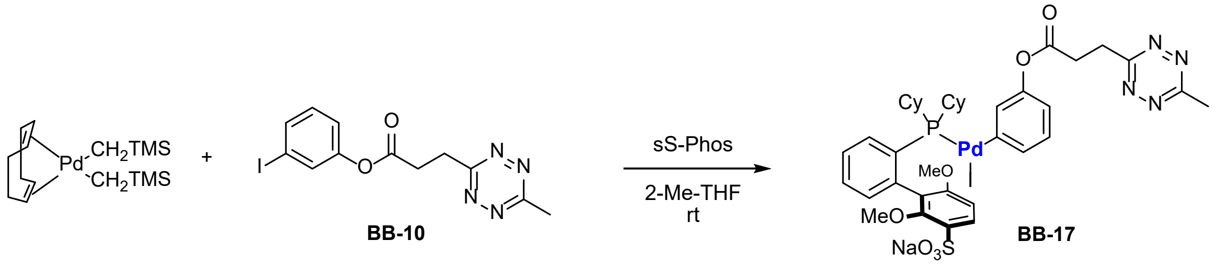

This compound was synthesized following the general procedure described in the section on the synthesis of **BB-14**. **BB-10** (15.8 mg, 0.043 mmol), sSPhos (20.7 mg, 0.040 mmol), [(cod)Pd($CH_2TMS$)$_2$ ] (15.0 mg, 0.039 mmol), and 2-methyltetrahydrofuran (0.5 mL) were used, and **BB-17** was obtained as a light brown solid (21 mg, 55%).

MS (m/z): $C_{38}H_{46}N_4O_7PPdS^+$ ([M-I-Na+H]$^+$) 839.2, found 839.1.

*Synthesis of* ***BB-18***

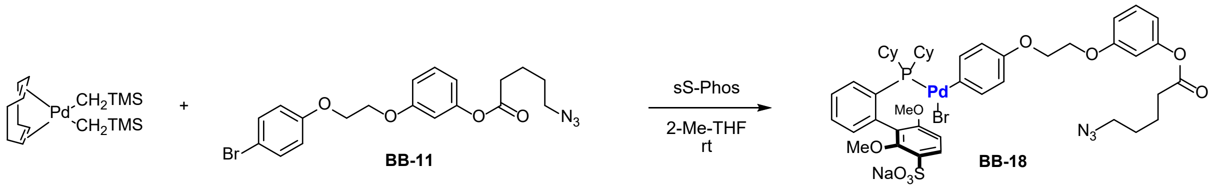

This compound was synthesized following the general procedure described in the section on the synthesis of **BB-14**. **BB-11** (18.5 mg, 0.043 mmol), sSPhos (20.7 mg, 0.040 mmol), [(cod)Pd($CH_2TMS$)$_2$ ] (15.0 mg, 0.039 mmol), and 2-methyltetrahydrofuran (0.5 mL) were used, and **BB-18** was obtained as a light brown solid (32 mg, 80%).

MS (m/z): $C_{45}H_{55}N_3O_9PPdS^+$ ([M-Br-Na+H]$^+$) 950.0, found 950.1.

*Synthesis of* ***BB-19***

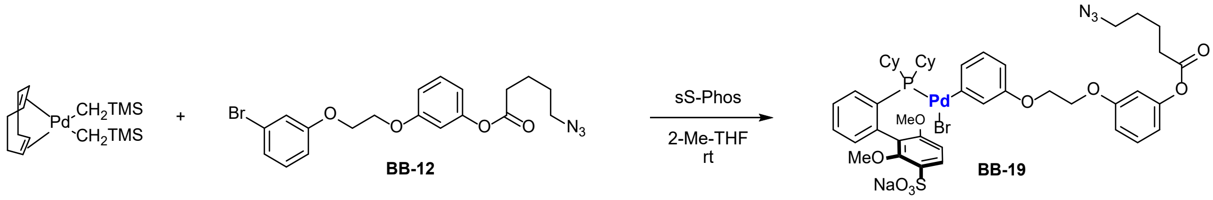

This compound was synthesized following the general procedure described in the section on the synthesis of **BB-14**. **BB-12** (17.0 mg, 0.039 mmol), sSPhos (19.0 mg, 0.037 mmol), [(cod)Pd($CH_2TMS$)$_2$ ] (13.8 mg, 0.035 mmol), and 2-methyltetrahydrofuran (0.5 mL) were used, and **BB-19** was obtained as a light brown solid (26 mg, 71%).

MS (m/z): $C_{45}H_{55}N_3O_9PPdS^+$ ([M-Br-Na+H]$^+$) 950.0, found 950.1.

*Synthesis of* ***BB-20***

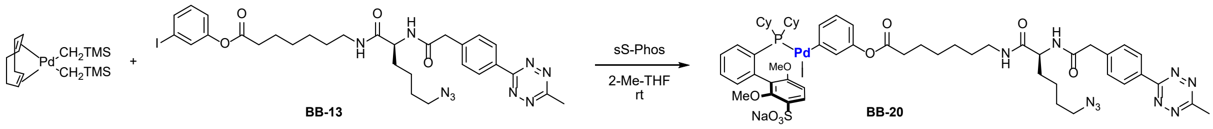

This compound was synthesized following the general procedure described in the section on the synthesis of **BB-14**. **BB-13** (15.0 mg, 0.021 mmol), sSPhos (10.3 mg, 0.020 mmol), [(cod)Pd($CH_2TMS)_2$ ] (7.5 mg, 0.019 mmol), and 2-methyltetrahydrofuran (0.4 mL) were used, and **BB-20** was obtained as a light brown solid (22 mg, 85%).

MS (m/z): $C_{56}H_{71}N_9O_9PPdS^+$ ([M-I-Na+H]$^+$) 1182.0, found 1182.0.

### 4. Synthesis of Electrophile-attached Z33 Reagents

Synthesized electrophile-attached Z33 reagents are listed in Table S1. Each modified peptide was synthesized as follows:

*Synthesis of $N_3$-**1-I***

**1** (15 mg) was weighed into a 1.5 mL microcentrifuge tube, and DMF (200 μL) was added. Additionally, **BB-14** (7.1 mg) was weighed into a 0.6 mL microcentrifuge tube, and DMF (0.18 mL) was added to prepare a separate brown solution using sonication. The **BB-14** solution was added to the solution of **1**, and the mixture was vortexed briefly and allowed to stand at room temperature for 30 min. Then the reaction mixture was transferred to a 50 mL centrifuge tube and diluted to 5 mL with water. Subsequently, the reaction solution was directly loaded onto a Biotage® Sfär C18 D column (12 g) and purified using the following conditions: 5 CV 5% B, 1 CV 5-10% B, 20 CV 10-35% B. 1 μL of a 20-fold diluted solution of the obtained fractions was analyzed by LC-MS, and the fractions containing only the target product were collected and lyophilized to obtain $N_3$-**1-I** (6.7 mg, 45%) as a white solid.

*Synthesis of $N_3$-**2-I***

**2** (15.0 mg) and **BB-14** (7.1 mg) were used to synthesize $N_3$-**2-I** following the same procedure as for *$N_3$-**1-I***. $N_3$-**2-I** (6.7 mg, 42%) was obtained as a white solid.

*Synthesis of $N_3$-**3-I***

**3** (15.7 mg) and **BB-14** (6.7 mg) were used to synthesize $N_3$-**3-I** following the same procedure as for $N_3$-**1-I**. $N_3$-**3-I** (9.5 mg, 58%) was obtained as a white solid.

*Synthesis of $N_3$-**4-I***

**4** (5.0 mg) and **BB-14** (2.3 mg) were used to synthesize $N_3$-**4-I** following the same procedure as for $N_3$-**1-I**. $N_3$-**4-I** was obtained as a white solid.

*Synthesis of $N_3$-**5-I***

**5** (5.0 mg) and **BB-14** (2.3 mg) were used to synthesize $N_3$-**5-I** following the same procedure as for $N_3$-**1-I**. $N_3$-**5-I** was obtained as a white solid.

*Synthesis of $N_3$-**6-I***

**6** (5.0 mg) and **BB-14** (2.3 mg) were used to synthesize $N_3$-**6-I** following the same procedure as for $N_3$-**1-I**. $N_3$-**6-I** was obtained as a white solid.

*Synthesis of **BB-21***

**2** (5.0 mg) and **BB-15** (2.5 mg) were used to synthesize **BB-21** following the same procedure as for **$N_3$-1-I**. **BB-21** (2.5 mg, 47%) was obtained as a white solid.

*Synthesis of **BB-22***

**2** (5.0 mg) and **BB-16** (2.2 mg) were used to synthesize **BB-22** following the same procedure as for **$N_3$-1-I**. **BB-22** (3.4 mg, 65%) was obtained as a white solid.

*Synthesis of **BB-23***

**2** (5.0 mg) and **BB-17** (2.4 mg) were used to synthesize **BB-23** following the same procedure as for **$N_3$-1-I**. **BB-23** (1.2 mg, 23%) was obtained as a white solid.

*Synthesis of **$N_3$-12-VIII***

**2** (5.0 mg) and **BB-20** (3.3 mg) were used to synthesize ***$N_3$-12-VIII*** following the same procedure as for **$N_3$-1-I**. **$N_3$-12-VIII** (0.32 mg, 6%) was obtained as a white solid.

*Synthesis of **$N_3$-7-VI***

**BB-2** (5.0 mg) and **BB-18** (2.6 mg) were used to synthesize ***$N_3$-7-VI*** following the same procedure as for **$N_3$-1-I**. **$N_3$-7-VI** (3.2 mg, 59%) was obtained as a white solid.

*Synthesis of **$N_3$-8-VI***

**BB-3** (5.0 mg) and **BB-18** (2.6 mg) were used to synthesize ***$N_3$-8-VI*** following the same procedure as for **$N_3$-1-I**. **$N_3$-8-VI** (3.1 mg, 57%) was obtained as a white solid.

*Synthesis of **$N_3$-7-VII***

**BB-2** (5.0 mg) and **BB-19** (2.6 mg) were used to synthesize ***$N_3$-7-VII*** following the same procedure as for **$N_3$-1-I**. **$N_3$-7-VII** (3.2 mg, 59%) was obtained as a white solid.

*Synthesis of* ***$N_3$-8-VII***

**BB-3** (5.0 mg) and **BB-19** (2.6 mg) were used to synthesize ***$N_3$-8-VII*** following the same procedure as for **$N_3$-1-I**. **$N_3$-8-VII** (4.5 mg, 83%) was obtained as a white solid.

*Synthesis of* ***$N_3$-9-VII***

**BB-4** (5.0 mg) and **BB-19** (2.6 mg) were used to synthesize ***$N_3$-9-VII*** following the same procedure as for **$N_3$-1-I**. **$N_3$-9-VII** (3.5 mg, 64%) was obtained as a white solid.

*Synthesis of* ***$N_3$-10-VII***

**BB-5** (5.0 mg) and **BB-19** (2.6 mg) were used to synthesize ***$N_3$-10-VII*** following the same procedure as for **$N_3$-1-I**. **$N_3$-10-VII** (0.5 mg, 9%) was obtained as a white solid.

*Synthesis of* ***$N_3$-11-VII***

**BB-6** (5.0 mg) and **BB-19** (2.6 mg) were used to synthesize ***$N_3$-11-VII*** following the same procedure as for **$N_3$-1-I**. **$N_3$-11-VII** (0.48 mg, 9%) was obtained as a white solid.

### 4.2. Click handle transfer to IgGs using electrophile-Z33 variants.

#### 4.2.1. Bioconjugation to Trastuzumab (IgG1)

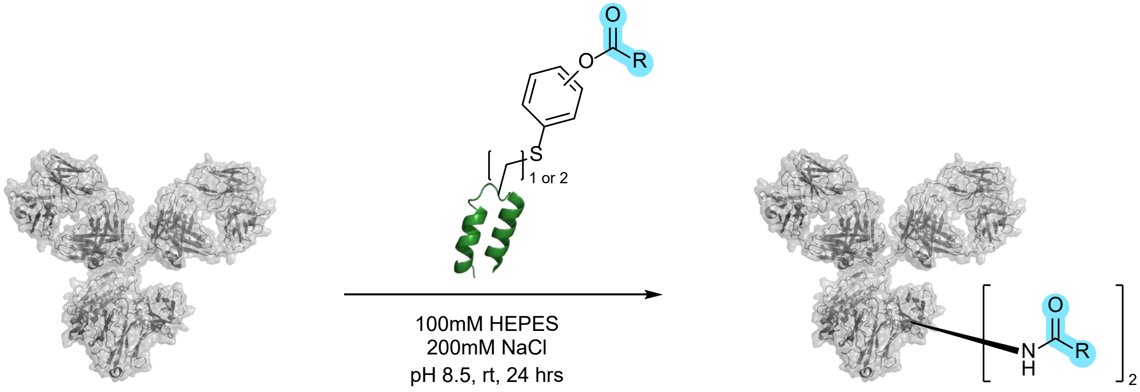

*<u>General Reaction Procedure</u>*

Reactions were performed on a 20 µL scale using 0.2 mL PCR tubes. First, the electrophile-attached Z33 peptide reagent was dissolved in water to prepare a 2 mg/mL solution. Trastuzumab (MedChemExpress LLC. or Bio X Cell), electrophile-attached Z33 peptide reagent (2 mg/mL), HEPES (pH 8.5), and NaCl were added to achieve final concentrations of 10 µM, 100 µM, 100 mM, and 200 mM respectively. The contents were mixed using a 10 µL pipette, followed by incubation at room temperature in the dark for 24 hours. The resulting reaction solution was quenched with a 100 mM glycine buffer or 100 mM Tris buffer and analyzed by LC-MS.

*General Procedure of LC-MS Analysis*

A reaction solution corresponding to 9.0 μg of IgG was taken into a PCR tube and diluted with Tris buffer (100 mM, pH 8.1) to a final volume of 6μL. Separately, a mixed solution of PNGase F (New England BioLabs, 63 units) and Glycoprotein Denaturing Buffer (New England Biolabs, 1X, 4μL) was prepared and added to the reaction solution, followed by incubation at 37 °C for 4 to 12 hours. DTT (200 mM in water, 2 μL) was added, and the mixture was further incubated at 37 °C for 1 hour. Finally, the solution was diluted to a volume of 40 μL with a mixture of water and acetonitrile (95:5), and 8 μL of the diluted solution was injected into LC-MS.

*Procedure using reagent **BB-23** or $N_3$**-12-VIII***

Reactions were performed on a 20 μL scale using 0.2 mL PCR tubes. First, the electrophile-attached Z33 peptide reagent was dissolved in water to prepare a 2 mg/mL solution. Trastuzumab (Bio X Cell), electrophile-attached Z33 peptide reagent (2 mg/mL), HEPES (pH 8.5), and NaCl were added to achieve final concentrations of 10 μM, 100 μM, 100 mM, and 200 mM respectively. The contents were mixed using a 10 μL pipette, followed by incubation at 37 °C in the dark for 24 hours. 4 hours after the start of incubation, an additional 10 equivalent of electrophile-attached Z33 peptide was added (200 μM in total). The resulting reaction solution was quenched with a 100 mM glycine buffer or 100 mM Tris buffer and analyzed by LC-MS.

#### 4.2.2. Bioconjugation to other IgGs (human IgG1,2,4 & mouse IgG1)

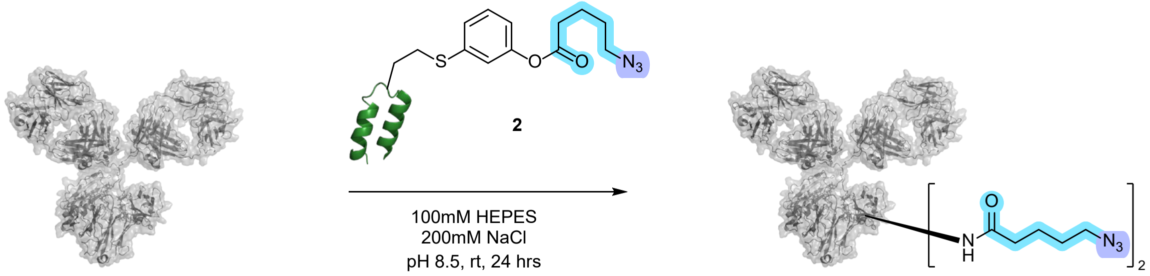

Reactions were performed on a 10 μL scale using 0.2 mL PCR tubes. First, the electrophile-attached Z33 peptide reagent was dissolved in water to prepare a 2 mg/mL solution. Denosumab (Invitrogen), dupilumab (Invitrogen), or mouse IgG1 (Invitrogen, subtype controlled)), electrophile-attached Z33 peptide reagent (2 mg/mL), HEPES buffer (pH 8.5), and NaCl solution were added to achieve final concentrations of 5 μM, 100 μM, 100 mM, and 200 mM. The contents were mixed using a 5 μL pipette and incubated at room temperature in the dark for 24 hours. The resulting reaction solution was quenched with a 100 mM Tris buffer and analyzed by LC-MS. Sample preparation for LC-MS analysis was performed in the same manner as described in 4.2.1.

#### 4.3. Determination of the Modification Sites

The modified trastuzumab (30 μg) from entry 6 of 5.2.1 was diluted to 36 μL using Tris buffer (50 mM, pH 8.0). PNGase F (New England BioLabs, 200 units) was added to the solution and incubated at 37 °C for 4 hours. The reaction solution was then brought to room temperature, and urea solution (6 M in 50 mM Tris pH 8.0, 14 μL) and DTT solution (200 mM, 1.5 μL) were added and incubated at 37°C for 1 hour. After that, iodoacetamide (800 mM, 1 µL) was added and incubated at room temperature for 30 min. The resulting solution was diluted 2-fold using Tris buffer (50 mM, pH 8.0), and then Trypsin/Lys-C mix (Promega, 0.2 µg/µL in solution, 6 µL) was added and incubated at 37 °C for 18 hours. The reaction solution was then purified by pipetting with Ziptip, lyophilized, and analyzed by nLC-MS/MS (using nLC-MS/MS method B). Obtained raw data were analyzed using Thermo Scientific FreeStyleTM 1.6. De novo sequencing was performed with PEAKS 8.5 with the following search parameters: Parent Mass Error Tolerance = 15.0 ppm; Fragment Mass Error Tolerance = 0.02 Da, Enzyme = None; modification setting are listed in the table below; Max Variable PTM Per Peptide = 10; Report # Peptides = 10. Sequencing results were exported as .csv reports for all de novo candidates. Finally, fragments corresponding to trastuzumab sequences were manually searched from the .csv file, and the main modification sites were estimated using each fragment's peak area.

| PTM type | Amino acids | Molecular weight | Setting Reason |
|---|---|---|---|
| Fixed | C | 57.0214 | Capped by iodoacetamide |
| Variable | K | 125.0589 | Modified by the electrophile attached peptide reagents |
| Variable | M | 15.9949 | Unintentional natural oxidation |

* Since the mass of the modified lysine is the same as the mass of Pro-Arg (271.1644), both possibilities were considered in the fragment search.

### 4.4. Expanding the Chemistry with Other Z33 Variants

#### 4.4.1. Using the Electrophile-attached Z33M3C/Hcy Reagents

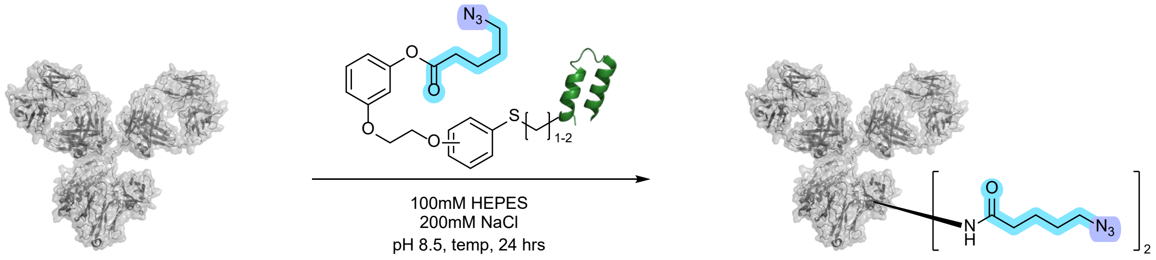

Reactions were performed on a 20 µL scale using 0.2 mL PCR tubes. First, the electrophile-attached Z33 peptide reagent was dissolved in water to prepare a 2 mg/mL solution. Trastuzumab (Bio X Cell), electrophile-attached Z33 peptide reagent (2 mg/mL), HEPES buffer (pH 8.5), and NaCl solution were added to achieve final concentrations of 10 µM, 200 µM, 100 mM, and 200 mM. The contents were mixed using a 10 µL pipette, followed by incubation at room temperature or 37 °C in the dark for 24 hours. The resulting reaction solution was quenched with a 100 mM Tris buffer and analyzed by LC-MS. Sample preparation for LC-MS analysis was performed in the same manner as described in 4.2.1.

#### 4.4.2. Using the Electrophile-attached Z33R31C/Hcy and D32C/Hcy Reagents

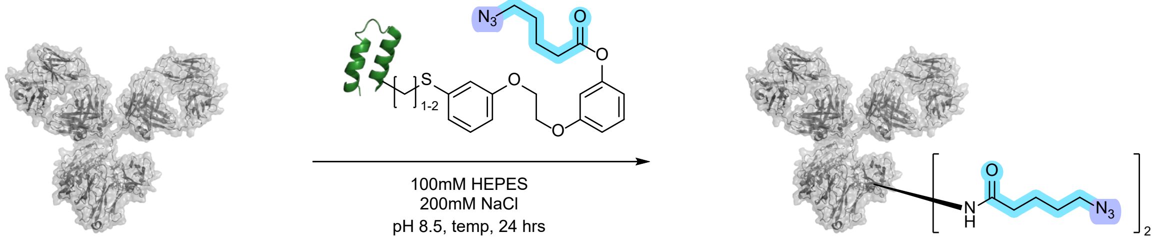

Reactions were performed on a 20 µL scale using 0.2 mL PCR tubes. First, the electrophile-attached Z33 peptide reagent was dissolved in water to prepare a 2 mg/mL solution. Trastuzumab (Bio X Cell), electrophile-attached Z33 peptide reagent (2 mg/mL), HEPES buffer (pH 8.5), and NaCl solution were added to achieve final concentrations of 10 µM, 200 uM, 100 mM, and 200 mM. The contents were mixed using a 10 µL pipette, followed by incubation at room temperature or 37 °C in the dark for 24 hours. The resulting reaction solution was quenched with a 100 mM Tris buffer and analyzed by LC-MS. Sample preparation for LC-MS analysis was performed in the same manner as described in 4.2.1.

#### 4.5. Dual Modification of the Trastuzumab Heavy Chain by Stepwise Reactions

First Tmab modification reaction was performed on a 200 µL scale using 1.5 mL microcentrifuge tube. $N_3$-**2-I** was dissolved in water to prepare a 2 mg/mL solution. Trastuzumab (Bio X Cell), $N_3$-**2-I** (2 mg/mL), HEPES buffer (pH 8.5), NaCl solution were added to achieve final concentrations of 10 µM, 100 µM, 100 mM, and 200 mM. The contents were mixed using a 200 µL pipette and incubated at room temperature for 24 hours. The resulting reaction solution was transferred to an Amicon 30K MWCO filter, and 200 µL of a 100 mM citric acid (pH 2.7) was added. The solvent was removed by centrifugation, and 200 µL of a 50 mM Tris buffer (pH 8.0) was added. Another round of centrifugation, 30 µL of a reaction mixture was obtained. Sample preparation for LC-MS analysis was performed in the same manner as described in 4.2.1. The concentration of the modified Tmab was determined to be 2.2 mg/mL by measuring absorbance at 280 nm.

Second Tmab modification was performed on a 20 µL scale using 0.2 mL PCR tubes. $N_3$-8-VI was dissolved in water to prepare a 2 mg/mL solution. Modified Tmab above, $N_3$-8-VI (2 mg/mL), HEPES buffer (pH 8.5), NaCl solution were added to achieve final concentrations of 10 µM, 100 µM, 100 mM, and 200 mM. The contents were mixed using a 20 µL pipette and incubated at 37 °C for 24 hours. The resulting reaction solution was quenched with a 100 mM Tris buffer and analyzed by LC-MS. Sample preparation for LC-MS analysis was performed in the same manner as described in 4.2.1.

#### 4.6. Trastuzumab Modification in the Presence of RNase A

Reactions were performed on a 20 µL scale using 0.2 mL PCR tubes. First, $N_3$-**2-I** was dissolved in water to prepare a 2 mg/mL solution. Tmab (Bio X Cell), RNAse A (VWR), $N_3$-**2-I** (2 mg/mL), and PBS were added to achieve final concentrations of 10 µM, 10 µM, 200 µM, and 1X. The contents were mixed using a 5 µL pipette and incubated at 37 °C for 24 hours. The resulting reaction solution was transferred to an Amicon 10K MWCO filter, and 200 µL of a

100 mM citric acid (pH 2.7) was added. The solvent was removed by centrifugation, and 200 µL of a 100 mM Tris buffer (pH 8.1) was added. Another round of centrifugation, 30 µL of a reaction mixture was obtained. Sample preparation for LC-MS analysis was performed in the same manner as described in 4.2.1.

**4.7. Antibody-drug conjugate (ADC) preparation with MMAE transfer reagent**
Reactions were performed on a 20 µL scale using 0.2 mL PCR tubes. First, $N_3$-**2-I** and **DBCO-PEG4-Val-Cit-PAB-MMAE** (BROADPHARM) were dissolved to prepare 2 mg/mL (in water) and 2 mM (in DMSO) solutions, respectively. A mixture of HEPES buffer and NaCl solution or PBS was added to the tubes, followed by $N_3$-**2-I** (2 mg/mL, 4.29 µL) and **DBCO-PEG4-Val-Cit-PAB-MMAE** (2 mM, 1.1 µL) were added, mixed using a 10 µL pipette, and incubated at 4°C for 30 min. The final concentrations of HEPES buffer (pH 8.5), NaCl solution, and PBS buffer were 100 mM, 200 mM and 1X, respectively. Tmab (Bio X Cell, 9.2 mg/mL, 1.61 µL) was then added to the reaction solution and incubated at room temperature or 37 °C for 24 hours. The resulting reaction solution was quenched with 100 mM Tris buffer and analyzed by SDS-PAGE.

*Sample Preparation for SDS-PAGE Analysis*

The reaction mixture corresponding to 3.5 µg of IgG was taken to a PCR tube, and diluted with Tris buffer (100 mM, pH 8.1) to a final volume of 3 µL. Then, Laemmli sample buffer (BIO-RAD, 2X +5% 2-mercaptoethanol, 3 µL) was added, and the mixture was heated at 70 °C for 5 min. After returning the reaction solution to room temparature, 2 µL was loaded onto an SDS-Gel. Subsequently, electrophoresis, washing, staining, and analysis were performedusing the methods described in General Information Section.

**5. Animal Experiments**
All animal experiments were performed under an MIT institute approved IACUC protocol following federal, state, and local guidelines for the care and use of animals (protocol number #0821-058-24). The studies were performed either on 35-45 g (12-16 weeks-old) wild-type female Swiss mice, 25-30 g (14-20 weeks-old) male and female C57BL/6J mice, or 55-70 g (16 weeks-old) male and female obese *Lep*$^{ob/ob}$ mice, all purchased from The Jackson Laboratory (MA, USA). Mice were housed with free access to normal food diet and water *ad libitum* unless stated otherwise for the need of the experiment. For subcutaneous (SC) injections, the mice were shortly anesthetized beforehand with 2-3% isoflurane along with $O_2$, then shaved on the right flank. Intravenous injections (IV) were performed through IV catheterization; the mice were anesthetized with 2-3% isoflurane along with $O_2$, then a catheter was inserted in the lateral tail vein to ensure the injection was properly done. Intraperitoneal injections (IP) were performed on non-anesthetized mice, in the lower right abdominal quadrant. Blood collection for the ELISA assays and biodistributions was performed as a terminal procedure by heart puncture immediately after the sacrifice. For Ip-GTT and Ip-ITT mice were restrained and gently pressed at the tail following tail pricks. The animals were sacrificed by $CO_2$ inhalation followed by cervical dislocation. All compounds injected in mice were either USP grade, sterile, or filtered using 0.22 µm.

### 5.1. Mouse IgG Painting with Azido Moieties

#### 5.1.1. Enzyme-linked Immunosorbent Assay (ELISA)

Two sandwich ELISA assays have been developed in-house, one for the detection of the azido moiety conjugated mouse IgG (mIgG, E1), the other one for the detection of the whole mIgGs (modified and unmodified) in the sample (E2). Both E1 and E2 were run simultaneously on the same plate, using the same initial stock sample, and were processed in parallel at the same time for every step. The same azido-free capture goat anti-mouse polyclonal antibody (Creative Diagnostics) was used for E1-E2 assays at 1 µg/mL (100 µL per well) for incubation overnight at 4 °C. Washing steps were performed thoroughly between each incubation by using 0.02% Tween 20-PBS. Saturation step (1 h, RT) and sample dilutions were all carried in 5% milk-PBS. For E1, the detection was performed using 1:150 dilution of DBCO-PEG4-Biotin (Jena Bioscience) (in 5% milk-PBS) while for E2 the detection was made using 1:2000 azido-free biotin-conjugated donkey anti-mouse polyclonal antibody (Creative Diagnostics) in 5% milk-PBS, both for 150 min at RT, in the dark. A dilution of 1:500 HRP-conjugated streptavidin (Thermo Fisher Scientific Inc.) in 5% milk-PBS was then added for 1 h, RT, in the dark. TMB substrate (Thermo Fisher Scientific Inc.) was added (100 µL per well, RT) simultaneously in E1 and E2 wells, and the reaction was stopped with 2 N $H_2SO_4$ stop solution (Thermo Fisher Scientific Inc.) after 5 to 10 min. Absorbances were measured at 450 nm and 540 nm ($A_0$ = $A_{450nm}$-$A_{540nm}$) using a Spark microplate reader (Tecan, USA). The background noise ($A_{blank}$) from the serum was subtracted to get the final absorbance of the sample ($A_{sample}$ = $A_0$-$A_{blank}$). The ratio $A_{sample}$ (E1)/ $A_{sample}$ (E2) was calculated to estimate the percentage of azido transfer on the total amount of IgG in the sample.

#### 5.1.2. Qualitative Assessment of Azido-transfer to Mouse IgG in Serum

Reactions were performed on a 40 µL scale using 0.2 mL PCR tubes. First, the azido electrophile-attached Z33 peptide reagent **1**, **2** or **3** was dissolved in water to prepare a 2 mg/mL solution. Mouse serum (Invitrogen, S/N 24-5544-94, 31.2 µL) and electrophile-attached Z33 peptide reagent (2 mg/mL, 8.8 µL) were added to the PCR tubes to a final concentration of 100 mM of electrophile-attached Z33 reagent. The contents were mixed using a 10 µL pipette, followed by incubation at 37 °C in the dark for 2, 6, or 24 hours. To remove the excess amount of the remaining electrophile-attached Z33 reagent, the reaction mixture was filtered using an Amicon® filter. The reaction mixture was diluted to 200 µL with citrate buffer (100 mM, pH 2.7) and filtered through an Amicon® filter (0.5 mL, 30KMWCO) at 14000 rcf for 7 min. After filtration, 400 µL of PBS buffer was added and centrifuged again. Finally, the remaining solution (roughly 20 µL) on the Amicon® filter was transferred to a microcentrifuge tube. Obtained samples were then diluted 1:100 to 1:2000 in 5% milk-PBS for dosage using E1/E2 assays.

#### 5.1.3. Qualitative Assessment of Azido-transfer to Mouse IgG in Vivo

Female Swiss mice (n=3 per group) were either IP or SC injected with 10 mg/kg (~90 nmol) to 30 mg/kg (~280 nmol) of $N_3$-**1**, $N_3$-**2**, or $N_3$-**3** (diluted in 1X PBS). Blood was recovered by intracardiac puncture (terminal procedure) at 24 hours post injection, then was centrifuged 10 min at 15,000 rcf to get serum. Obtained samples were then diluted 1:2 in 5% milk-PBS for dosage using E1/E2 assays.

### 5.2. Mouse IgG Painting with Radionuclides

#### 5.2.1. Preparation of Deferoxamine B-attached Z33 Reagents and Radiolabeling with Zirconium-89

First, **1**, **2, 3** were dissolved in HEPES buffer (100 mM in ultra-trace elemental water; Fisher Scientific, pH 6.7) to prepare a 2 mg/mL solution of each. Deferoxamine-DBCO (obtained from Macrocycles, 1 mM in DMSO, 11.1 µL) and **1**, **2, 3** (360 µL) solutions were then added to the tube, stirred gently by hand, and the tube was placed on ice for 1 hour.

The obtained samples were incubated with 37 MBq of [$^{89}$Zr]Zr-oxalate (obtained from Mallinckrodt Institute of Radiology, Nuclear Pharmacy Cyclotron; dilution in 0.5 M HEPES, pH 6.7) for 1 hour in the wet ice under mild agitation. Purification was then performed using a Zeba spin column (0.5 mL, 7K MWCO) to remove the non-complexed [$^{89}$Zr]Zr-oxalate, for 5 min at 1,200 rcf. Instant-thin layer chromatography (iTLC) was performed to verify the success of the radiolabeling with elution in 0.1 M citrate buffer.

Representative instant-thin layer chromatography iTLC before and after purification:

Free [$^{89}$Zr]Zr-oxalate:

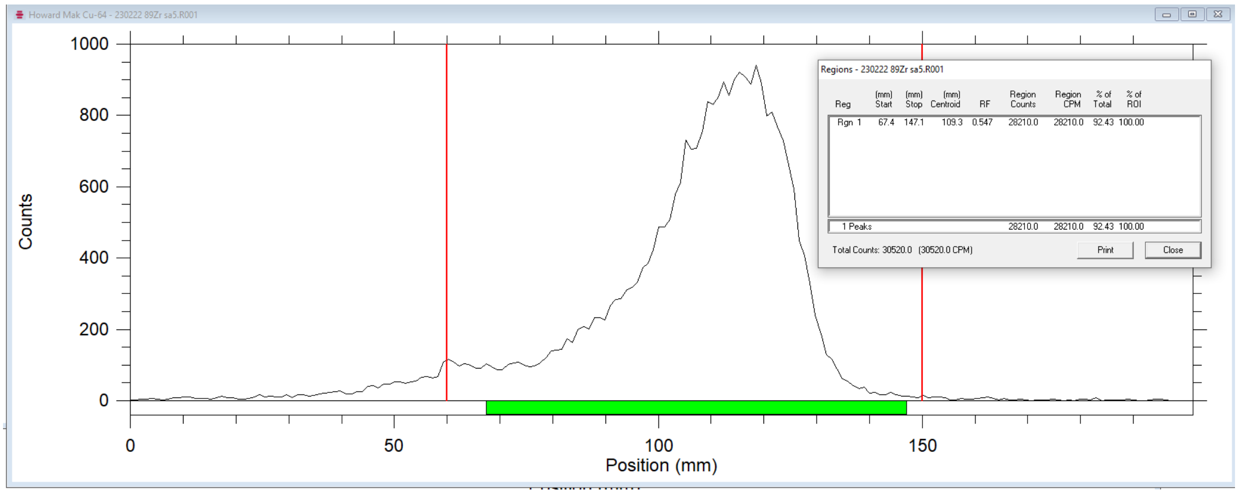

Peptide: [$^{89}$Zr]Zr-**1**: (pre-purification using ZebaSpin columns)

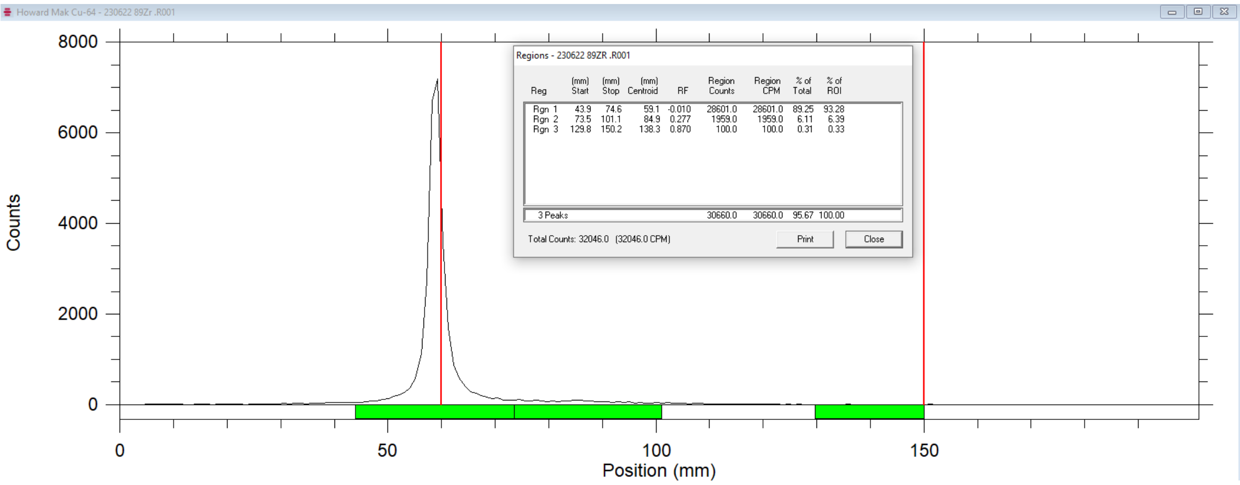

Peptide: [$^{89}$Zr]Zr-**1**: (post-purification using ZebaSpin columns)

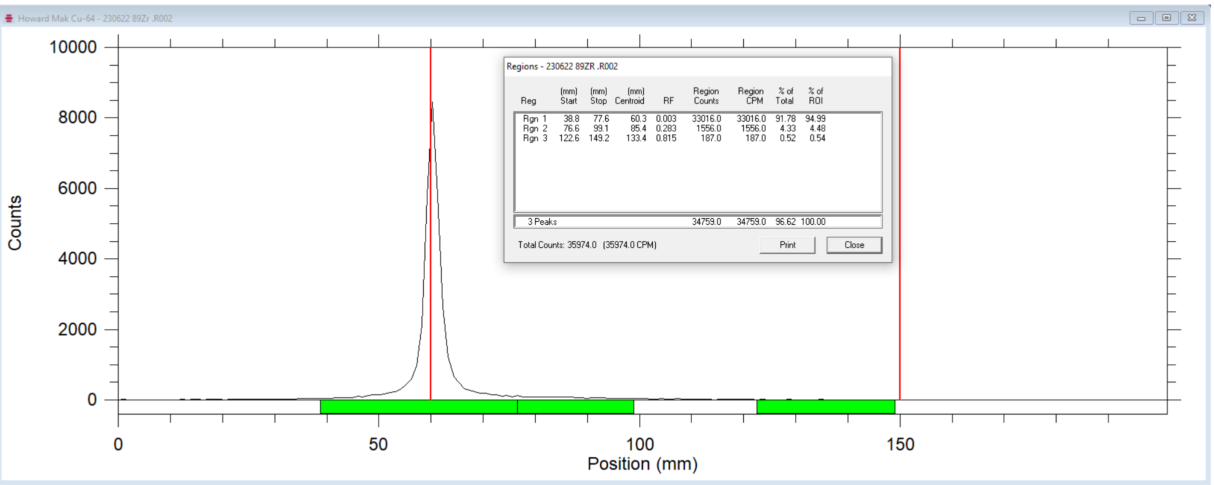

Peptide: [$^{89}$Zr]Zr-**2**: (pre-purification using ZebaSpin columns)

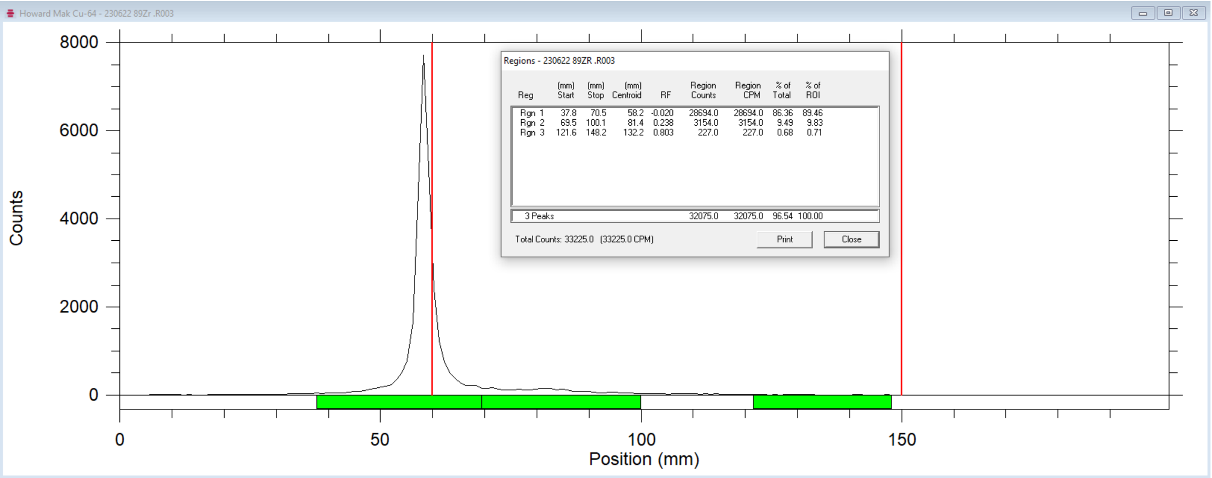

Peptide: [$^{89}$Zr]Zr-**2**: (post-purification using ZebaSpin columns)

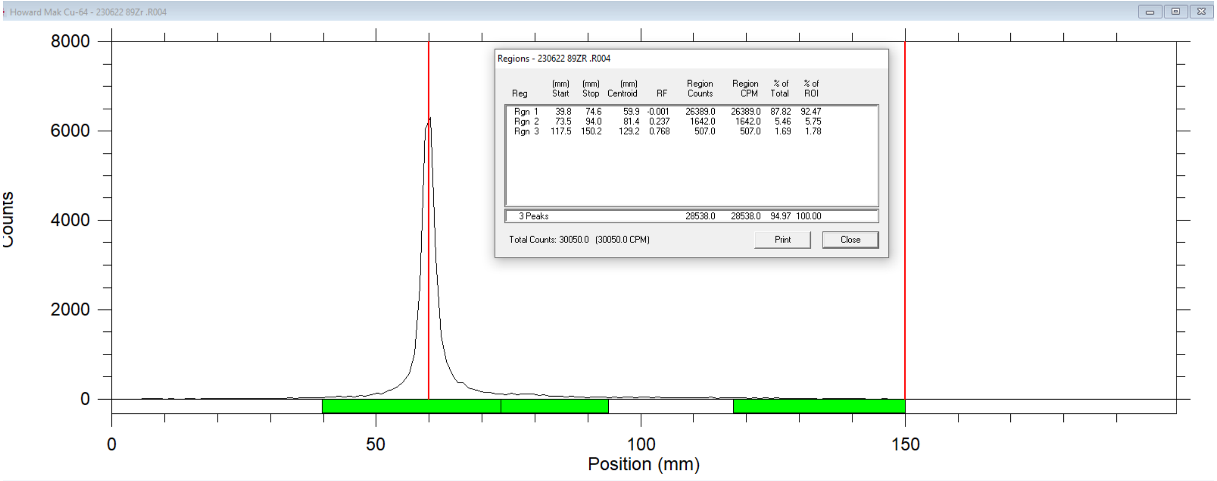

#### 5.2.2. Pharmacokinetics of Radionuclide Transfer to mIgGs and Whole-body Biodistribution

For injection in animals, samples were incubated with 55.5 MBq of [$^{89}$Zr]Zr-oxalate (obtained from Mallinckrodt Institute of Radiology, Nuclear Pharmacy Cyclotron; dilution in 0.5M HEPES, pH 6.7) for 1.5 hours in wet ice under mild agitation. No purification step was performed as our previous iTLC analysis showed no significant purity difference between the pre- and post-purification of samples.

*Intravenous injections.* Female WT Swiss mice (n=4 per group) were anesthetized using 2.5% isoflurane mixed with $O_2$, then a catheter (29 gauge) was inserted in the tail vein. About 1.2-1.5 MBq of [$^{89}$Zr]Zr, [$^{89}$Zr]Zr-**1**, [$^{89}$Zr]Zr-**2**, or [$^{89}$Zr]Zr-**3** was then injected. The injected radioactive material was eventually flushed with 100 µL of sterile saline solution. Longitudinal PET-CT was performed then mice were sacrificed at the last time point (144 h) for harvesting organs and fluid for quantification.

*Intraperitoneal injections*. Female WT Swiss mice (n=16 per group, 4 per time point) were manually restrained and injected in the lower right part of the abdomen with 1.2 to 1.5 MBq of [$^{89}$Zr]Zr, [$^{89}$Zr]Zr-**1** or [$^{89}$Zr]Zr-**2** then sacrificed after 2 h, 24 h, 72 h, or 144 h for harvesting tissues for quantification.

*Subcutaneous injections*. Female WT Swiss mice (n=20 per group, 4 per time point) were anesthetized using 2.5% isoflurane mixed with $O_2$, shaved on the right flank, then subcutaneously injected using a 29-gauge insulin syringe. Longitudinal PET-CT was performed then mice were sacrificed at the last time point (240 h) for harvesting tissues for quantification.

*PET-CT imaging*. Static PET-CT acquisitions were made using a Gator8 PET imager (Perkin Elmer LLC). PET acquisitions were performed for 15 min, followed by 5 min CT scan. Images were analyzed using VivoQuant software (Perkin Elmer LLC).

*Biodistribution*. After sacrifice, the following tissues were collected: blood, urine, feces, heart, lungs, liver, spleen, stomach, small intestines, colon, kidneys, bladder, pancreas, caecum, uterus + ovaries, gastrocnemius muscle, bone, skin, brain, and eyes. For subcutaneous group we also collected the inguinal and axillary lymph nodes, and the thymus. Each sample was weighted using a milligram precision scale then counted using a gamma counter (Wizard$^2$ 3", Perkin Elmer LLC) to get the counts per min (cpm) for determining the percentage of injected activity per gram (%IA/g) in each organ. Cpm were decay-corrected by using the following formula: $A = A_0\ e^{((-(\ln 2)\Delta t)/T)}$ with A = decay-corrected cpm, $A_0$ = raw cpm, Δt = time difference between the injection and the cpm measurement, and T = half-life of the radionuclide (*i.e.* 78.41 h for Zr-89).

#### 5.3. Mouse IgG Painting with Bioactive GLP-1 peptide analogs

##### 5.3.1. Preparation of the GLP-1 analog-attached Z33 transfer reagents.

*Synthesis of GLP-1 peptide analog* ***GLP-1a**, **1b**, and **1c***

**GLP-1a** or **GLP-1b** were synthesized according to the method in section 2, except for the following conditions: ChemMatrix® H-Rink Amide resin (0.19 mmol/g, 150 mg) was used, and the noncanonical amino acid Fmoc-Aib-OH was prepared as a 0.40 M DMF solution and incorporated into the sequence by AFPS. For the introduction of Aib and N-terminal H, the condensation reagent was changed to PyAOP, and pump stroke number was changed to 35. A mixture of TFA, water, thioanisole, phenol, and TIPS in a ratio of 33:2:2:1 was used as the cleavage solution from the resin. Additionally, if $CO_2$ adducts were observed in the crude product, the crude product was dissolved in a 95:5 water-acetonitrile solution (5 mL), and 200 µL of acetic acid was added before purification. The mixture was incubated at 37°C for 1 hour and then loaded into a Biotage® for purification. 22.0 mg of **GLP-1a** and 20.0 mg of **GLP-1b** were obtained.

**GLP-1c** was synthesized using the method described above from Ala-21 to Ser-39. The resin in a plastic fritted syringe was transferred from the AFPS system to a manifold. Fmoc-protected azidolysine (10 equivalents for the peptide on the resin) in HATU/DMF (0.38 M, 9.5 equivalents) solution and DIEA (15 equivalents) were added to the resin and allowed to react for 50 min at room temperature. The reaction solution was stirred with a spatula every 10 min. After the reaction, the reaction solution was removed, and the resin in the syringe was washed three times with 5 mL of DMF. Then, 3 mL of 40% piperidine/DMF solution (+2% formic acid) was added and allowed to react for 10 min. The reaction solution was removed, and 3 mL of 40% piperidine/DMF solution (+2% formic acid) was added again and allowed to react for 10 min. The reaction solution was removed, and the resin was washed three times with 4 mL of DMF. Subsequent synthesis of the sequence using natural amino acids and Aib from Tyr-1 to Gln-19 was again performed according to the procedure described above. A mixture of TFA, water, thioanisole, phenol, and TIPS in a ratio of 33:2:2:1 was used as the cleavage solution from the resin. Additionally, if $CO_2$ adducts were observed in the crude product, the crude product was dissolved in a 95:5 water-acetonitrile solution (5 mL), and 200 µL of acetic acid was added before purification. The mixture was incubated at 37°C for 1 hr. Agilent automated flash chromatography system Agilent 1290 Infinity II Preparative LC system (column; Zorbax 300SB-C3, 9.4 x 250 mm, 5 µm) was used for the purification of the half amount of the crude product, and 6.5 mg of the **GLP-1c** was obtained.

| Peptide | Sequence | Calculated Mass (Da) | Observed Mass (Da) |
|---|---|---|---|
| GLP-1**a** | HAibEGTFTSDVSSYLEGQAA**K**EFIAW LVRGRG | 3396.8 | 3397.3 |
| GLP-1**b** | HAEGTFTSDVSSYLEGQAA**K**EFIAWLVRGRG | 3382.7 | 3383.3 |
| GLP-1**c** | YAibEGTFTSDYSIAibLDKIAQ**Lys{N$_3$}**AFVQW LIAGGPSSGAPPPS | 4095.6 | 4095.7 |

*Synthesis of **GLP-1a** analog and compound **2a**:*

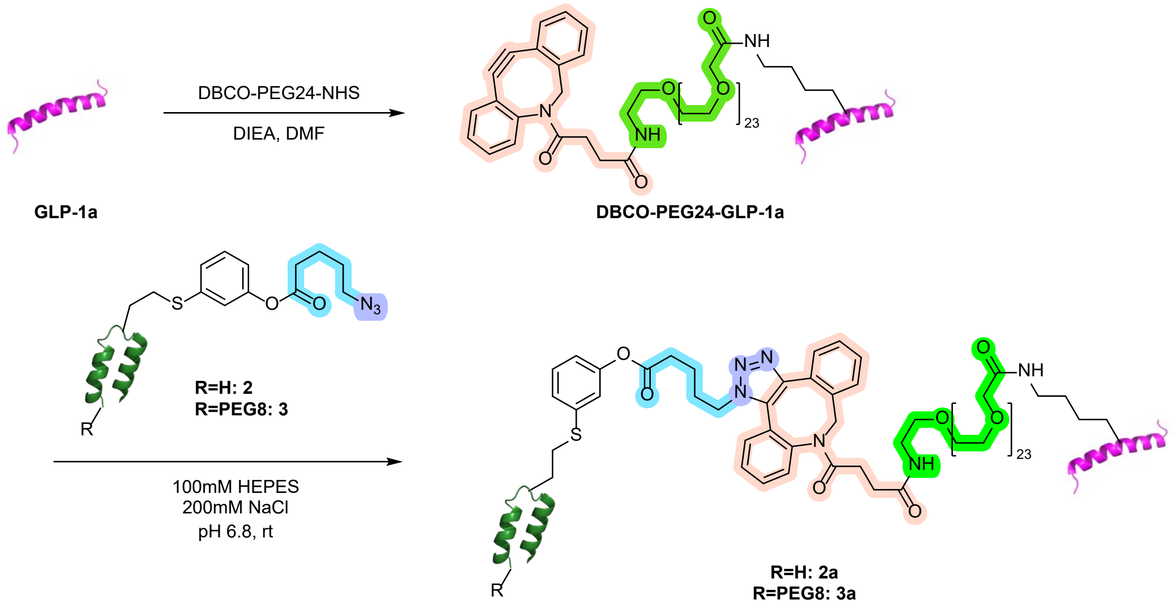

The reaction was performed using a 1.5 mL microcentrifuge tube. **GLP-1a** (18.3 mg) was dissolved in 600 µL of DMF. After adding 9 µL of DIEA, a separately prepared solution of DBCO-PEG24-NHS (BROADPHARM, 30 mg/mL in DMF, 300 µL) was added. The mixture was incubated at room temperature for 30 min, and the reaction was quenched with 1-naphthylmethylamine (9 µL). Then the reaction mixture was transferred to a 50 mL centrifuge tube and diluted to 5 mL with water. Subsequently, the reaction solution was directly loaded onto a Biotage® Sfär C18 D column (12 g) and purified using the following conditions: 5 CV 5% B, 1 CV 5-35% B , 6 CV 35% B, and 32 CV 35-50% B. 1 µL of a 5-fold diluted solution of the obtained fractions was analyzed by LC-MS, and the fractions containing only the target product were collected and lyophilized to obtain **DBCO-PEG$_{24}$-GLP-1a** (9.6 mg, 37%) as a white solid.

The modification site of **DBCO-PEG$_{24}$-GLP-1a** was determined using nLC-MS/MS. **DBCO-PEG$_{24}$-GLP-1a** (100 nmol, 1 µL) was injected into the nLC-MS/MS system, and nLC-MS/MS method B was used for this experiment.

Finally, **2** (4.0 mg) and **DBCO-PEG$_{24}$-GLP-1a** (4.9 mg) were dissolved in HEPES buffer (100 mM, pH 6.7) to prepare a 2 mg/mL solution of each. Both solutions were then added to a 15 mL centrifuge tube, stirred gently by hand, and placed on ice for 30 min. Then, the reaction solution was directly loaded onto a Biotage® Sfär C18 D column (12 g) and purified using the following conditions: 5 CV 5% B, 1CV 5-20% B, 25CV 20-60% B. 1 µL of a 5-fold diluted solution of the obtained fractions was analyzed by LC-MS, and the fractions containing only the target product were collected and lyophilized to obtain **2a** (6.2 mg, 73%) as a white solid. For **3a**, **3** (7.0 mg) and **DBCO-PEG$_{24}$-GLP-1a** (7.8 mg) were used, and 10.2 mg (59%) of **3a** was obtained as a white solid.

*<u>Synthesis of **GLP-1b** analog and compound **2b**:</u>*

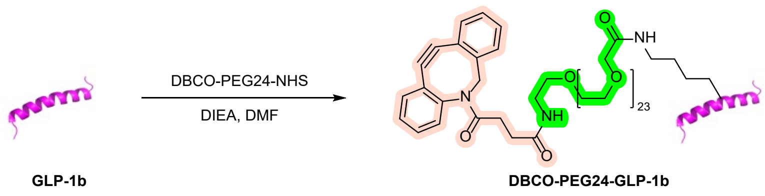

Synthesis of **Z33-PEG24-GLP-1b** was performed according to the same procedure as **DBCO-PEG24-GLP-1a**. 5.0 mg of GLP-1b was used, and 2.4 mg (34%) of DBCO-PEG24-GLP-1b was obtained as a white solid.

*<u>Synthesis of **GLP-1c** analog and compound **2c**:</u>*

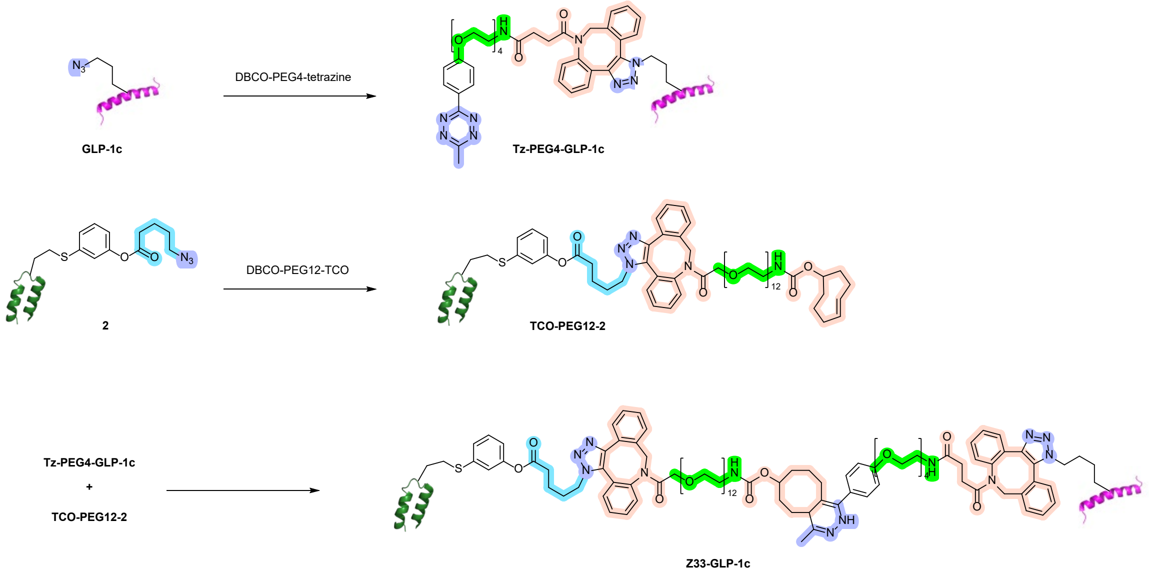

The reaction was performed using a 2.0 mL microcentrifuge tube. The **GLP-1c** (1.5 mg) was dissolved in 1.5 mL of water. After adding 500 µL of DMSO, a separately prepared solution of DBCO-PEG4-methyltetrazine (BROADPHARM, 10 mM in DMSO, 73.2 µL) was added. The mixture was incubated at 4 °C for 80 min. The reaction was used for the next step without further purification.

The reaction was performed using a 2.0 mL microcentrifuge tube. The **2** (1.5 mg) was dissolved in 0.75 mL of water. After adding HEPES buffer (465 mM, 215 µL), a separately prepared solution of DBCO-PEG12-TCO (BROADPHARM, 10 mM in DMSO, 34.8 µL) was added. The mixture was incubated at 4 °C for 80 min. The reaction was used for the next step without further purification.

The reaction was performed using a 5.0 mL microcentrifuge tube. The prepared **Tz-PEG4-GLP-1c** and **TCO-PEG12-2** solution were mixed and incubated at 4 °C for 1 hour. Then the reaction mixture was directly loaded onto a Biotage® Sfär C18 D column (12 g) and purified using the following conditions: 5 CV 5% B, 1 CV 5-25% B, and 70 CV 25-60% B. 1 µL of a 5-fold diluted solution of the obtained fractions were analyzed by LC-MS, and the fractions containing only the target product were collected and lyophilized to obtain **Z33-GLP-1c** (1.0 mg, 24%) as a white solid.

#### 5.3.2. Assessment of GLP-1 Transfer using human IgG$_1$ Trastuzumab

*With **GLP-1a***

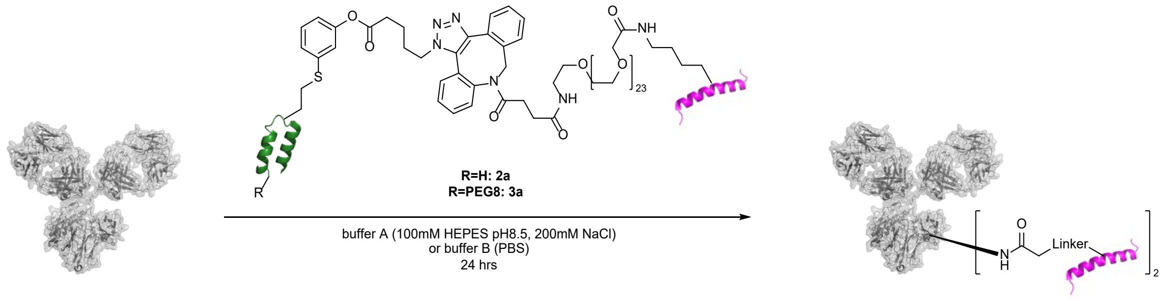

Reactions were performed on a 20 µL scale using 1.5 mL microcentrifuge tubes. First, **2a** or **3a** was dissolved in water to prepare a 2 mg/mL solution. A mixture of HEPES buffer and NaCl solution or PBS was added to the tubes, followed by trastuzumab (Bio X Cell, 9.2 mg/mL, 1.61 µL), **2a** (2 mg/mL, 9.12 µL) or **3a** (2 mg/mL, 9.55 µL) were added, mixed using a 10 µL pipette, and incubated at room temperature or 37 °C for 24 hours. The final concentrations of HEPES buffer, NaCl solution, and PBS buffer were 100 mM, 200 mM, and 1X, respectively. The reaction mixture was then diluted to 200 µL with citrate buffer (100 mM, pH 2.7) and filtered through an Amicon® filter (0.5 mL, 50K MWCO) at 14000 rcf for 7 min. After filtration, 400 µL of Tris buffer (100 mM, pH 8.1) was added and centrifuged again. Finally, the remaining solution (roughly 20 µL) on the Amicon® filter was transferred to a microcentrifuge tube, and the Amicon® filter was washed twice with 10 µL of Tris buffer (100 mM, pH 8.1). Sample preparation for LC-MS or SDS-PAGE analysis was performed in the same manner as described in 4.2.1. or 4.7.

*With **GLP-1b***

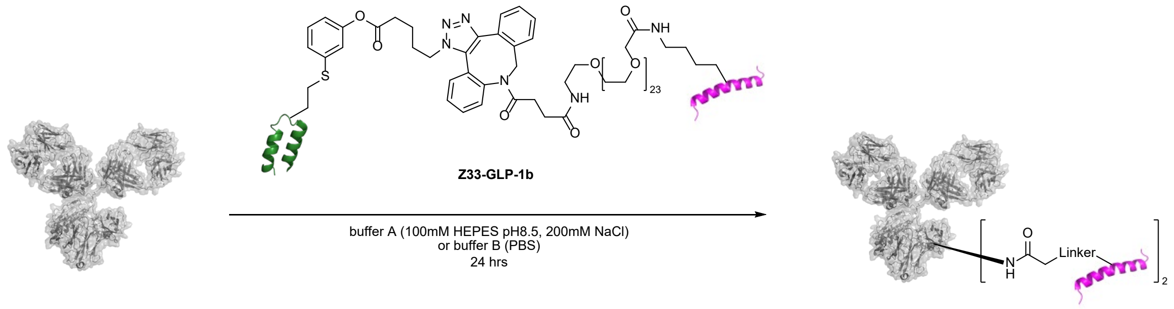

Reactions were performed on a 20-µL scale using 1.5-mL microcentrifuge tubes. First, **2** and **DBCO-PEG24-GLP-1b** were dissolved in water to prepare 2 mg/mL and 1 mg/mL solutions, respectively. A mixture of HEPES buffer and NaCl solution or PBS was added to the tubes, followed by 2 (2 mg/mL, 4.29 µL) and **DBCO-PEG24-GLP-1b** (1 mg /mL, 9.80 µL) were added, mixed using a 10 µL pipette, and incubated at 4 °C for 30 min. The final concentrations of HEPES buffer, NaCl solution, and PBS buffer were 100 mM, 200 mM and 1X, respectively. Trastuzumab (Bio X Cell, 9.2 mg/mL, 1.61 µL) was then added to the reaction solution and incubated at room temperature or 37 °C for 24 hours. The resulting reaction solution was quenched with 100 mM Tris buffer and analyzed by SDS-PAGE. Sample preparation for SDS-PAGE analysis was performed in the same manner as described in 4.7.

*With **GLP-1c***

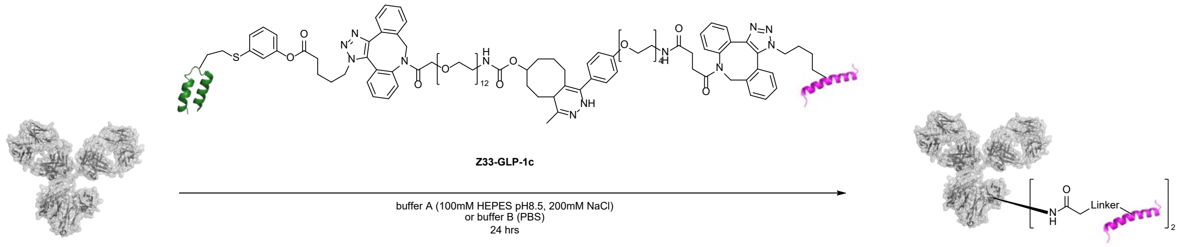

Reactions were performed on a 20 µL scale using 1.5 mL microcentrifuge tubes. First, **Z33-GLP-1c** was dissolved in water to prepare a 2 mg/mL solution. A mixture of HEPES buffer and NaCl solution or PBS was added to the tubes, followed by trastuzumab (Bio X Cell, 9.2 mg/mL, 1.61 µL), **2a** (2 mg/mL, 9.02 µL) were added, mixed using a 10 µL pipette, and incubated at room temperature or 37 °C for 24 hours. The final concentrations of HEPES buffer, NaCl solution, and PBS buffer were 100 mM, 200 mM, and 1X, respectively. The reaction mixture was then diluted to 200 µL with citrate buffer (100 mM, pH 2.7) and filtered through an Amicon® filter (0.5 mL, 50K MWCO) at 14000 rcf for 7 min. After filtration, 400 µL of Tris buffer (100 mM, pH 8.1) was added and centrifuged again. Finally, the remaining solution (roughly 20 µL) on the Amicon® filter was transferred to a microcentrifuge tube, and the Amicon® filter was washed twice with 10 µL of Tris buffer (100 mM, pH 8.1). Sample preparation for SDS-PAGE analysis was performed in the same manner as described in 4.7.

#### 5.3.3. Toxicity and dose response of semaglutide in wild-type Swiss mice.

The toxicity and maximum tolerated dose (MTD) of semaglutide (purchased from Adipogen Inc.) were assessed in wild-type female WT Swiss mice after one single SC injection of either 0.5 mg/kg (n=5), 3 mg/kg (n=5), or 10 mg/kg (n=15). The fasting blood glucose was measured after 24 h to determine the hypoglycemic status of the animal. The aspect and behavior of the animal, such as its hair condition, a change in the animal activity or socialization, aggressiveness, and body weight loss (≥ 20%) were also assessed to determine the presence of significant side-adverse toxicity. Ip-GTT was also performed at 24 h post injection to evaluate the effect on blood glucose and validate the wild-type Swiss mice model for studying the IgG painting strategy with GLP-1 analogs.

#### 5.3.4. Pharmacodynamic studies, toxicity, and dose response of in vivo IgG Painting with GLP-1 analogs in wild-type Swiss Mice.

Female WT Swiss mice were SC injected with 10 mg/kg of semaglutide (Adipogen Inc.) (n=15), **2a** (n=15), or **3a** (n=15). Body weight was monitored over 21 days for the entire cohort. At 24 h, 72 h, or 144 hours post injection, an IP-glucose tolerance test (Ip-GTT) was performed.

*Ip-GTT.* Six hours before the Ip-GTT, the mice were fasted by removing the food, the enrichment, and by placing them in a new cage to avoid coprophagy. Access to water was, however kept free and *ad libitum* for the entire experiment. Five min before the Ip-GTT, the fasting blood glucose was measured by taking a small drop of blood (< 5 µL) using tail pricks. Then, 2 g/kg of a 20% solution of dextrose (Sigma Aldrich Inc.) was IP injected followed by blood glucose measurements over 120 min using a glucometer (Contour®). *NB: At the 24-hour time-point, the test was stopped after 30-45 min as the glycemia of the mice was severely dropping and started showing moderate to severe hypoglycemia.*

The dose response to **2a** and **3a** was assessed following two different dose regimens, either one single SC injection of 30 mg/kg, or 3 SC injections of 10 mg/kg at a rate of 1 injection per week for 3 weeks. The toxicity of **2a** and **3a** was assessed following the same protocol as for semaglutide.

#### 5.3.5. Efficacy of in vivo IgG painting with GLP-1 analogs in obese *Lep*$^{ob/ob}$ mice.

Male and female *Lep*$^{ob/ob}$ obese mice (n=6 per compound; 3 males, 3 females) received one single SC injection of 10 mg/kg of semaglutide, **2a**, or **3a**. Body weight was measured over 21 days. Blood glucose was measured after 6 hours of fasting. For fasting, the food and enrichment were removed, and the mice placed in a new cage to avoid coprophagy. Access to water was kept free and *ad libitum* for the entire experiment. Normal C57BL/6J mice (n= 6; 3 males, 3 females) were also included as a second naïve control to obtain baseline values for non-diabetic non-obese animals. The entire study was duplicated to assess the efficacy after one single IP injection of 10 mg/kg of semaglutide, **2a**, or **3a**.

*Ip-GTT assays*. Ip-GTT was performed at 24 h and 144 h after the drug injection, through the IP injection of 2 g/kg of a 20% solution of dextrose (Sigma Aldrich Inc.), followed by blood glucose measurements over 120 min using a glucometer (Contour®).

*Ip-ITT assays*. Intraperitoneal insulin tolerance test (Ip-ITT) was performed at 240 h after drug injection, through the IP injection of 2.5 UI/kg of insulin (Humulin R®, Fisher Scientific), followed by blood glucose measurements over 90 min using a glucometer (Contour®). The insulin sensitivity of the animals was determined using the plasma glucose disappearance rate (K-ITT) calculated using the Ip-ITT data. *NB: The normal C57BL/6J mice were excluded from the IP-ITT assay due to the impossibility to inject the same amount of insulin (0.8 UI/kg maximum) as the insulin resistant $Lep^{ob/ob}$ mice (2 UI/kg minimum).*

### Supplementary LC-MS traces

In all cases below, the characterization data includes figures depicting from top to bottom: (a) the total ion count chromatogram (TICC) showing the extracted spectral region, (b) the extracted mass-to-charge (m/z) mass spectrum from the marked region of the TICC (a), (c) the deconvoluted mass spectrum corresponding to the m/z spectrum in (b), and (d) the UHPLC chromatogram (in select cases as shown).

Peptide: **1**

Structure:

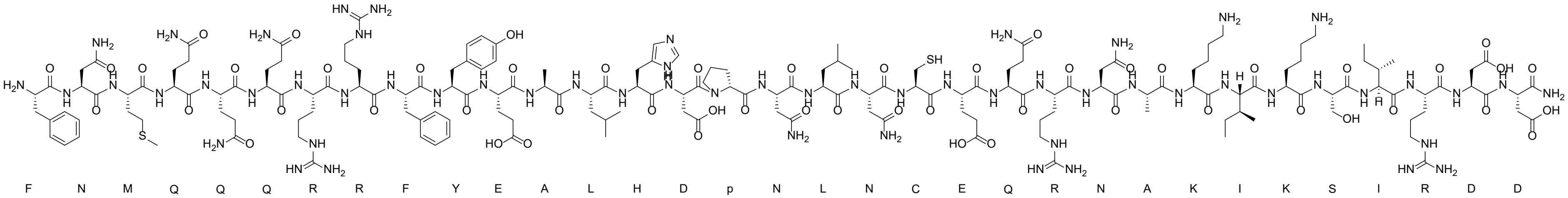

LC-MS analysis: Method A

Calculated Mass: 4078.6 Da

Observed Mass: 4078.6 Da

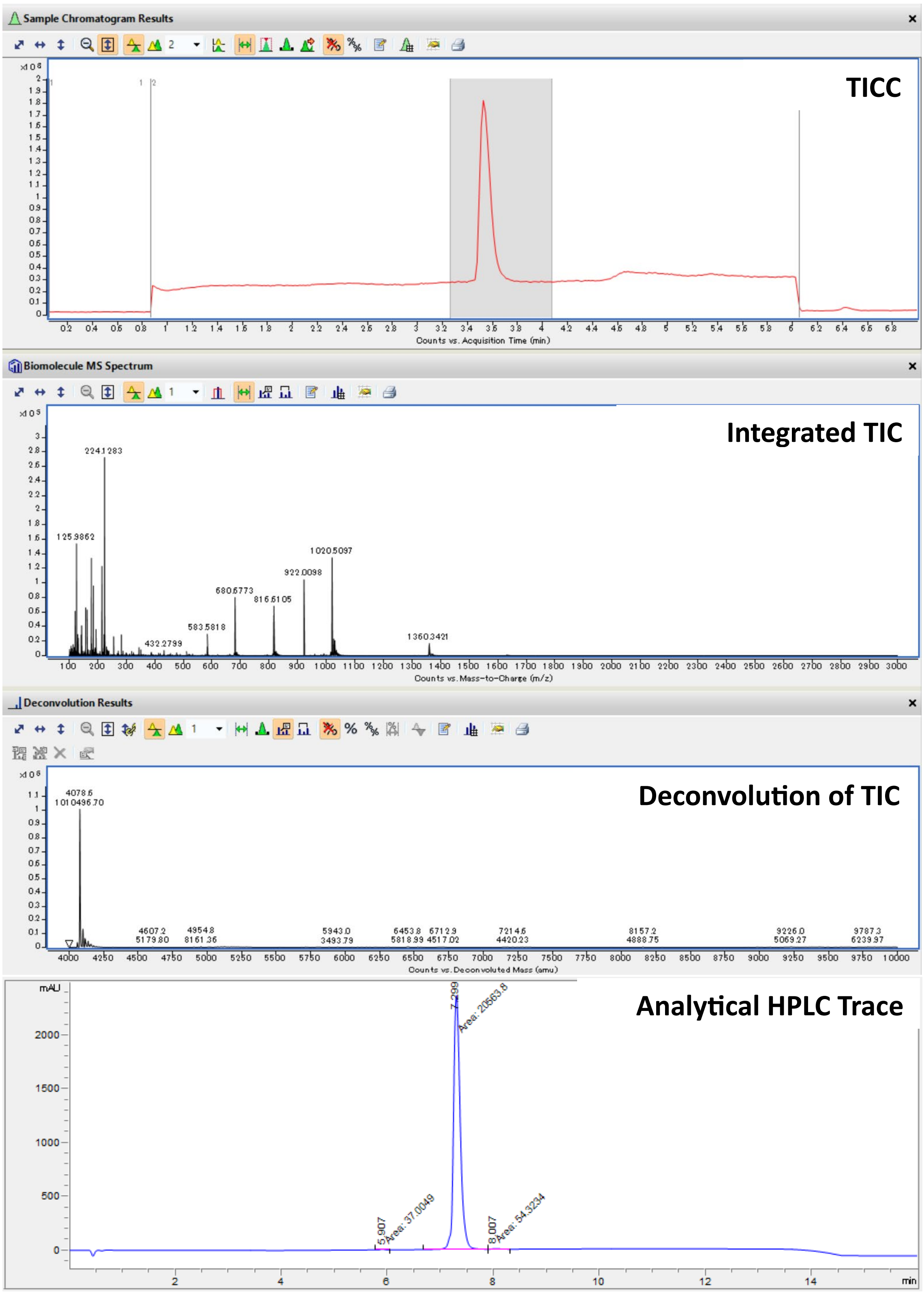

TICC: Total Ion Current Chromatogram; TIC: Total Ion Current.

Peptide: **2**

Structure:

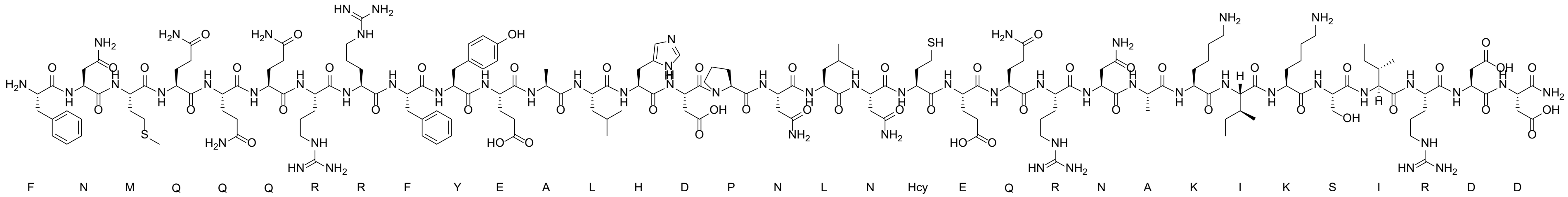

LC-MS analysis: Method D

Calculated Mass: 4092.6 Da

Observed Mass: 4093.1 Da

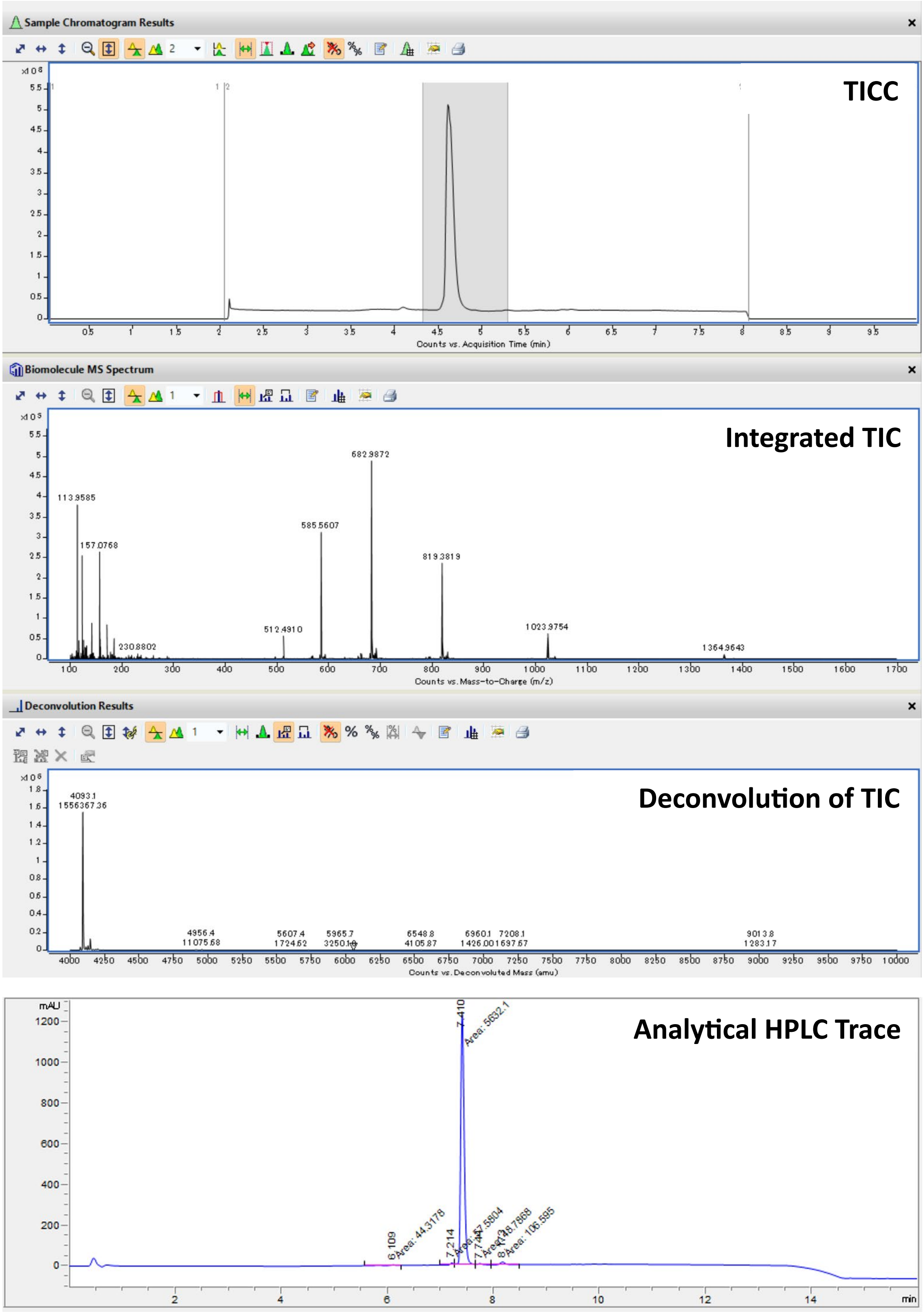

TICC: Total Ion Current Chromatogram; TIC: Total Ion Current.

Peptide: **3**

Structure:

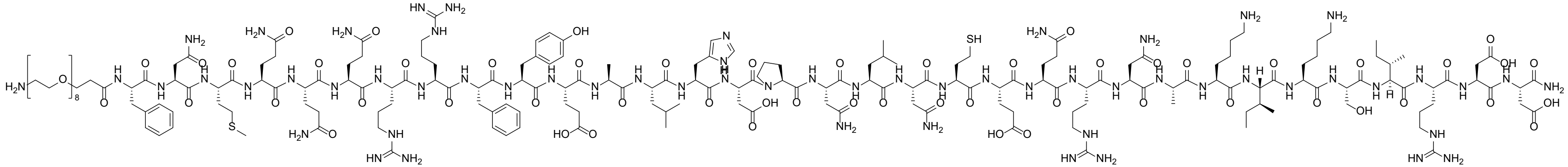

LC-MS analysis: Method C

Calculated Mass: 4516.1 Da

Observed Mass: 4516.2 Da

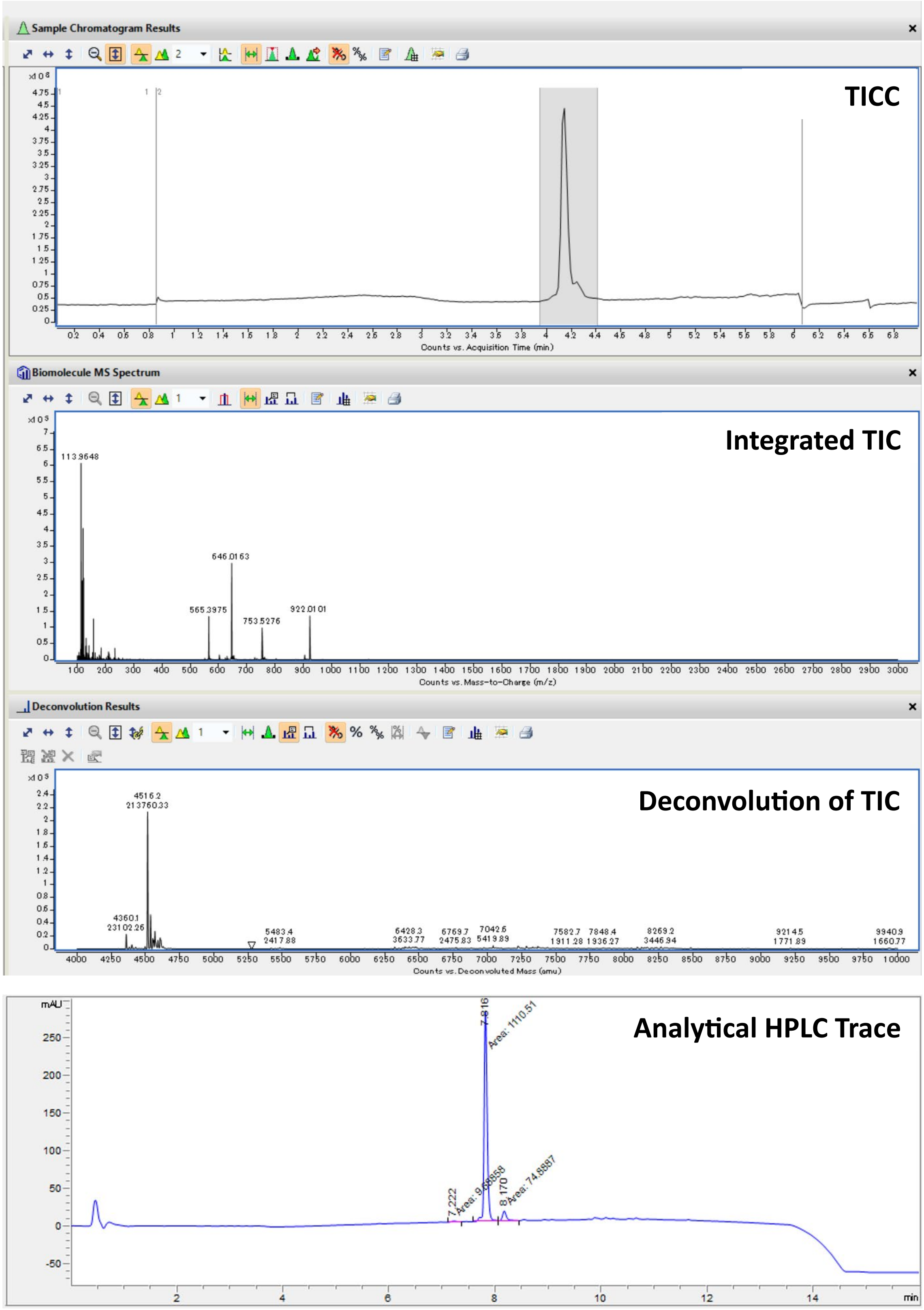

TICC: Total Ion Current Chromatogram; TIC: Total Ion Current.

Peptide: **4**

Structure:

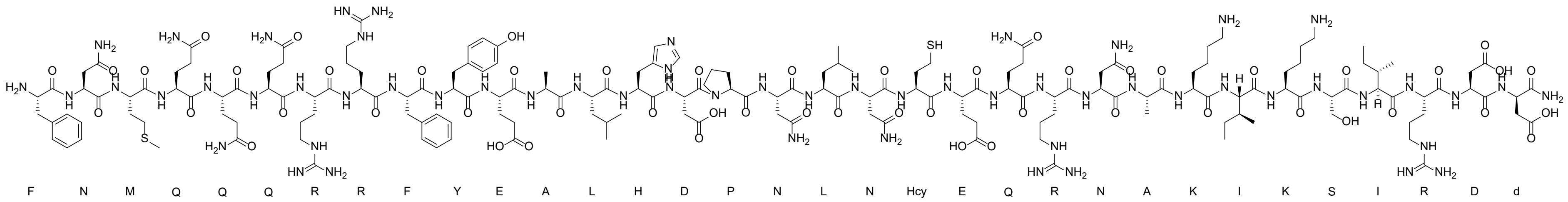

LC-MS analysis: Method E

Calculated Mass: 4092.6 Da

Observed Mass: 4092.5 Da

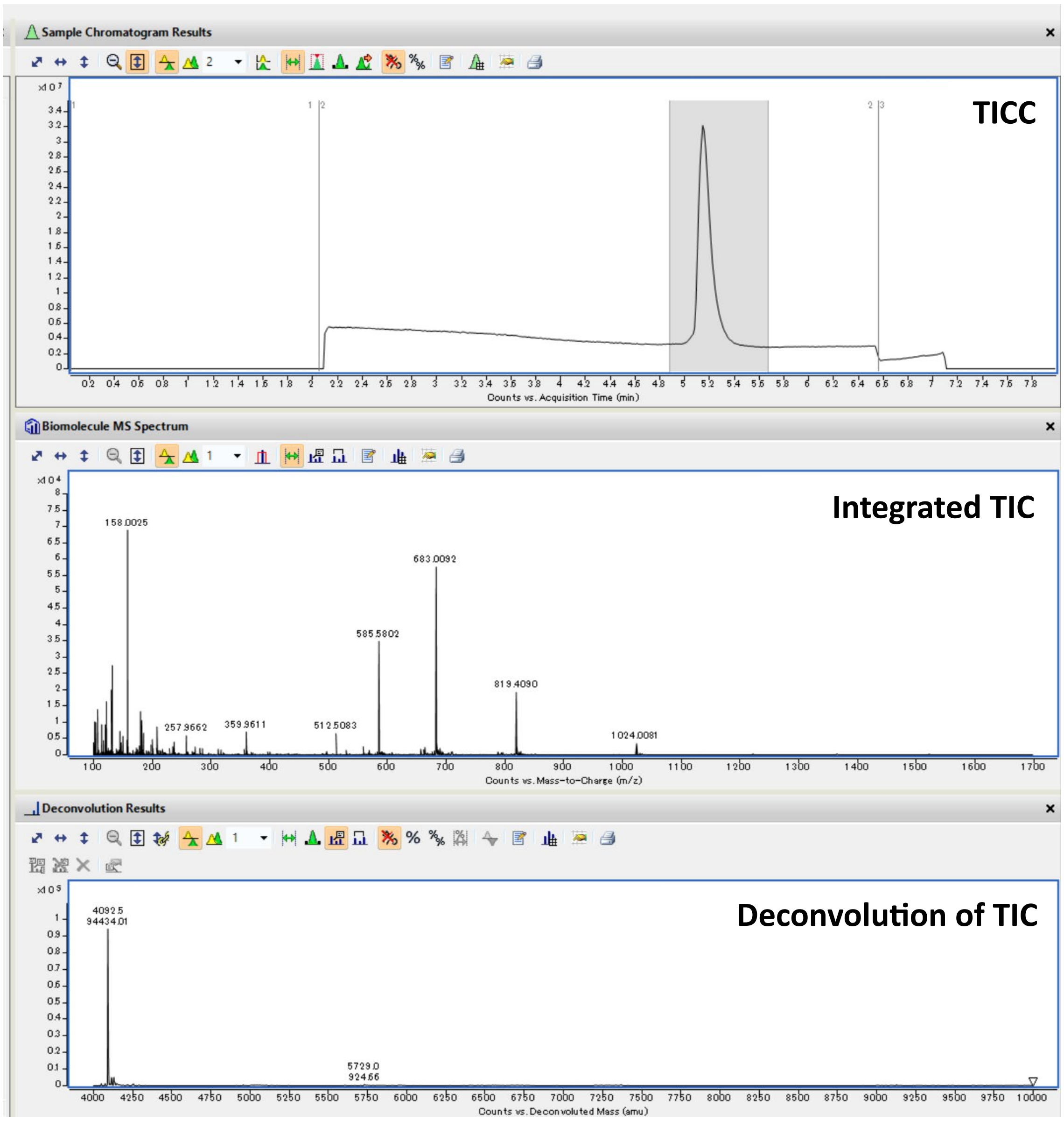

TICC: Total Ion Current Chromatogram; TIC: Total Ion Current.

Peptide: **5**

Structure:

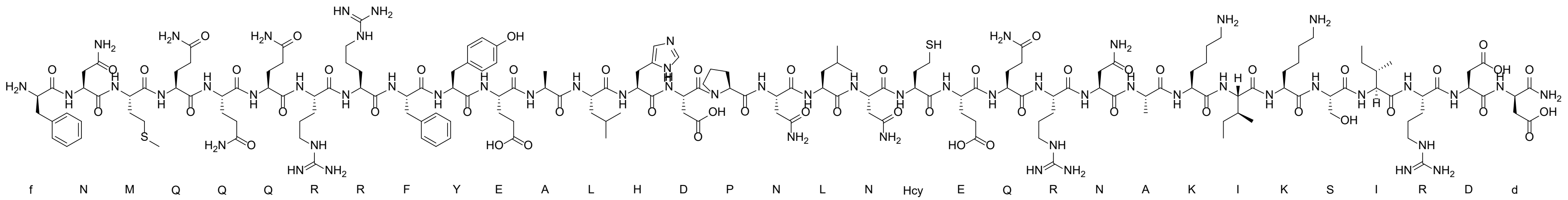

LC-MS analysis: Method E

Calculated Mass: 4092.6 Da

Observed Mass: 4092.5 Da

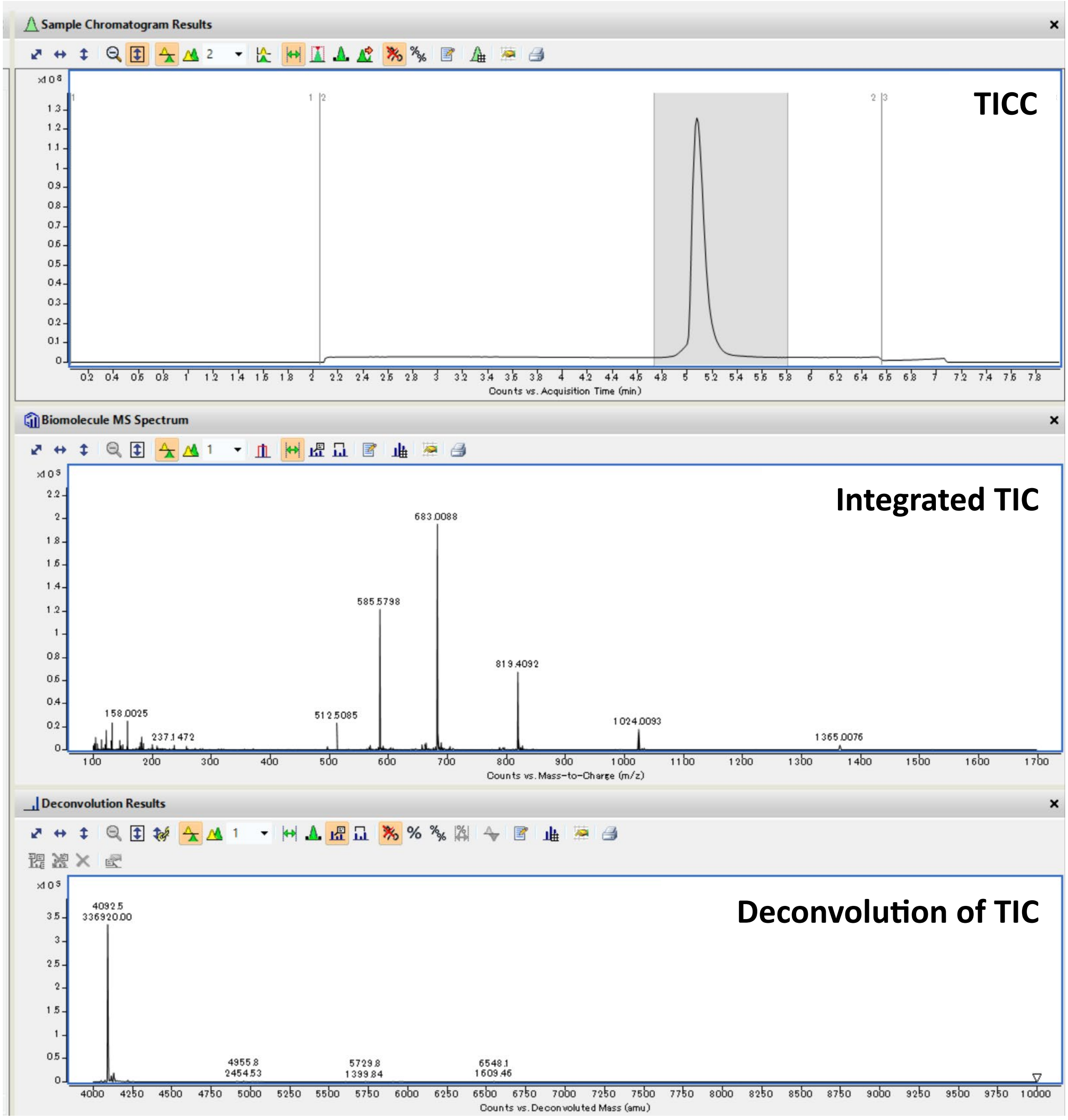

TICC: Total Ion Current Chromatogram; TIC: Total Ion Current.

Peptide: **6**

Structure:

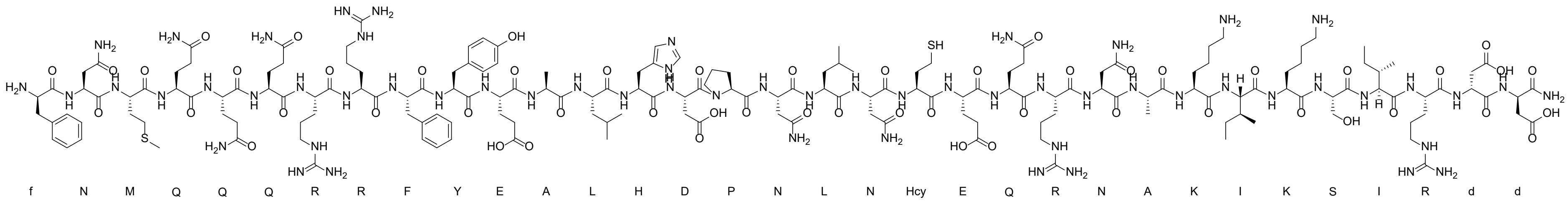

LC-MS analysis: Method E

Calculated Mass: 4092.6 Da

Observed Mass: 4092.6 Da

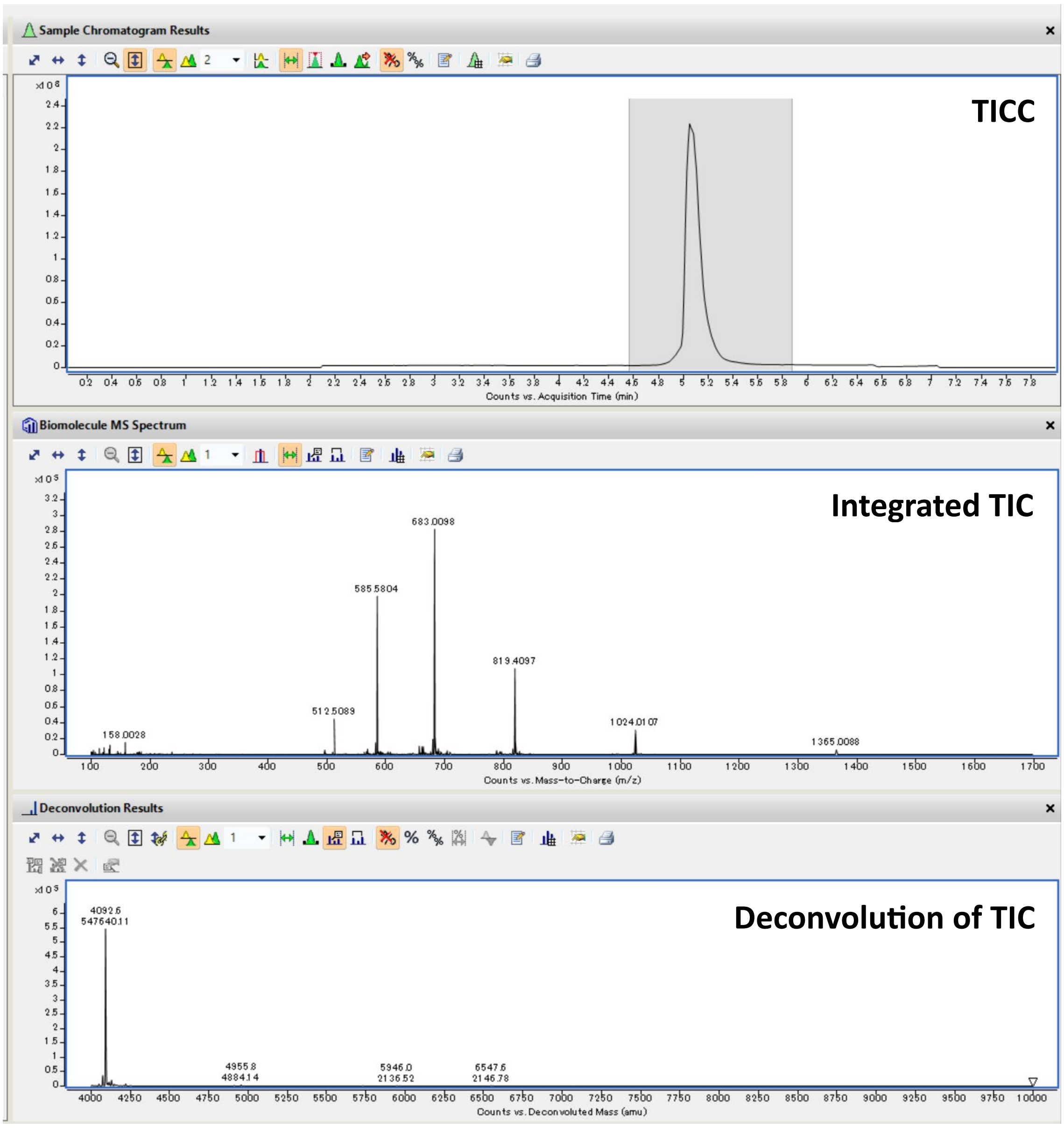

TICC: Total Ion Current Chromatogram; TIC: Total Ion Current.

Peptide: BB-1

Structure:

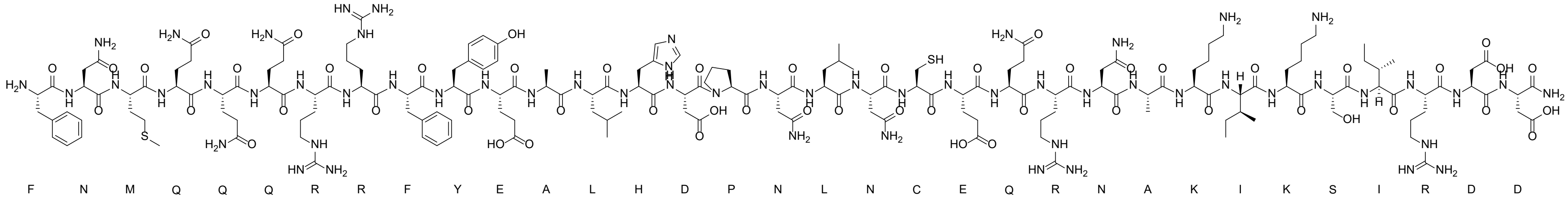

LC-MS analysis: Method D

Calculated Mass: 4078.6 Da

Observed Mass: 4078.5 Da

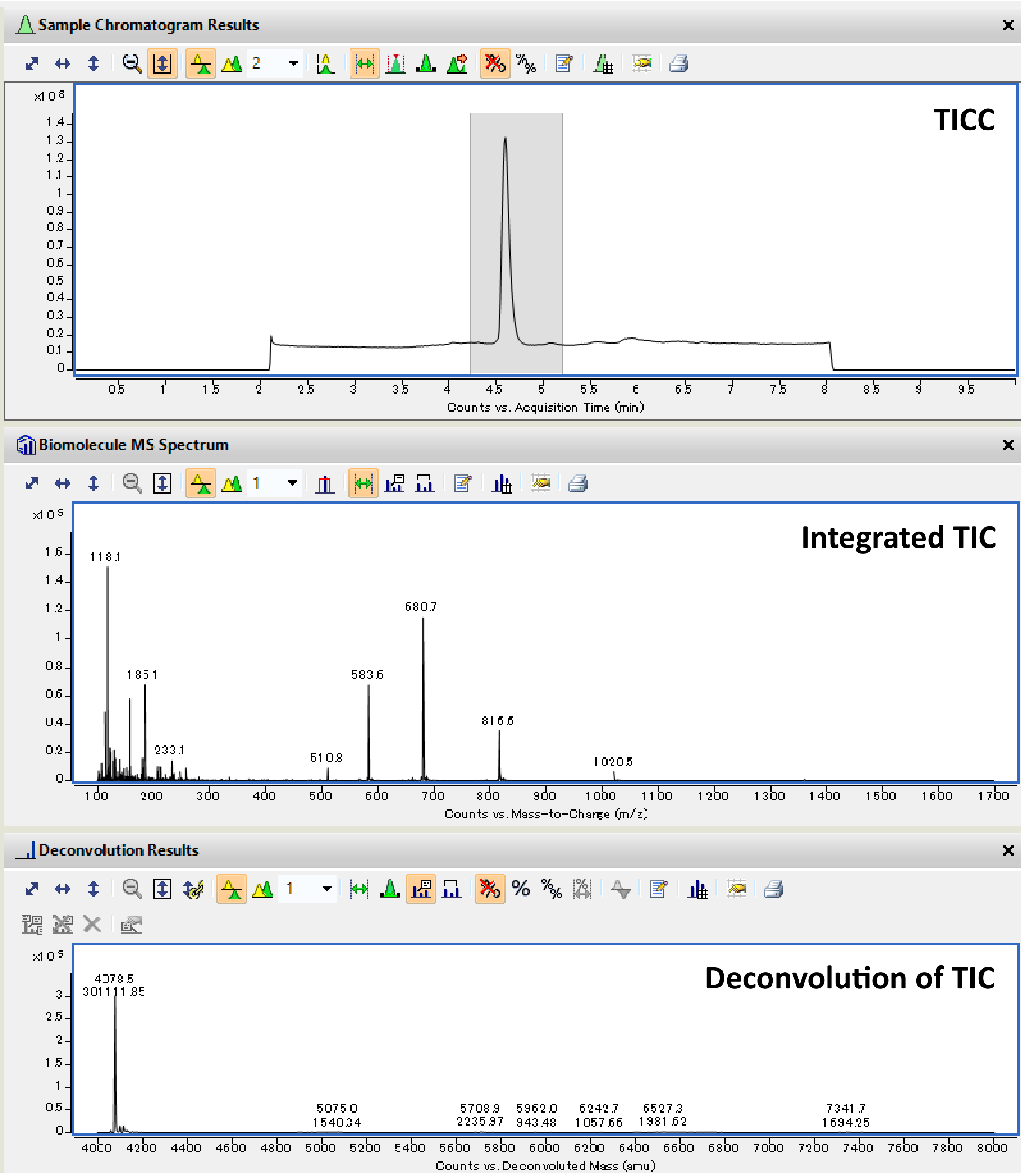

TICC: Total Ion Current Chromatogram; TIC: Total Ion Current.

Peptide: **BB-3**

Structure:

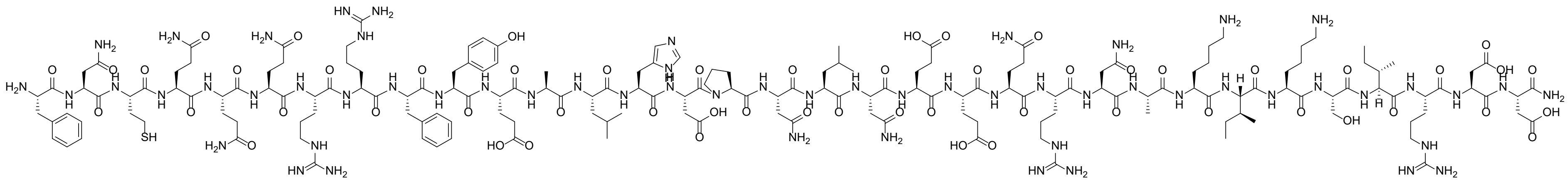

LC-MS analysis: Method A

Calculated Mass: 4090.5 Da

Observed Mass: 4090.7 Da

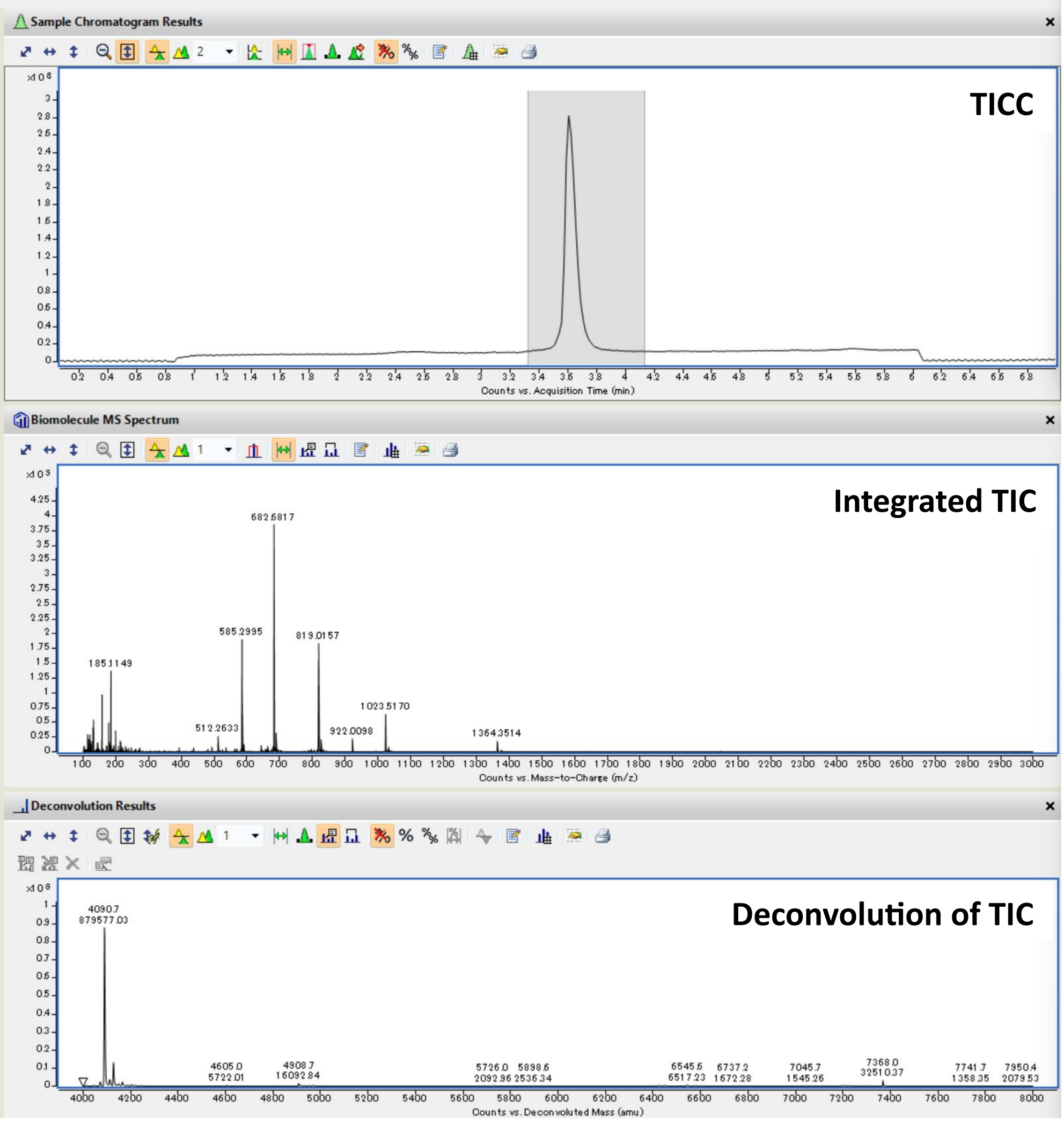

TICC: Total Ion Current Chromatogram; TIC: Total Ion Current.

Peptide: **BB-4**

Structure:

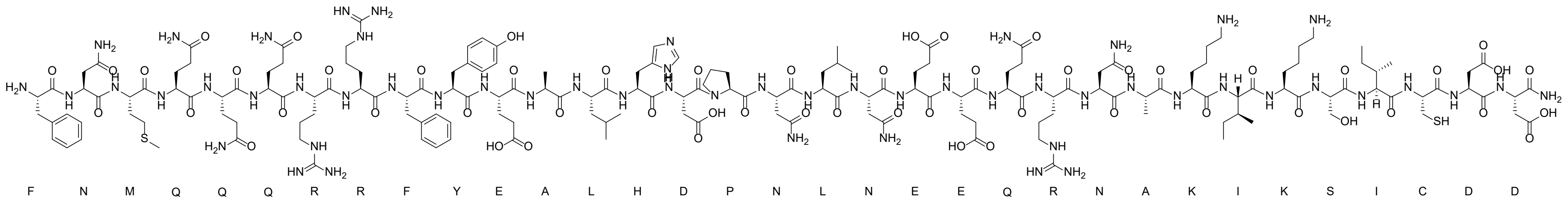

LC-MS analysis: Method E

Calculated Mass: 4051.5 Da

Observed Mass: 4052.3 Da

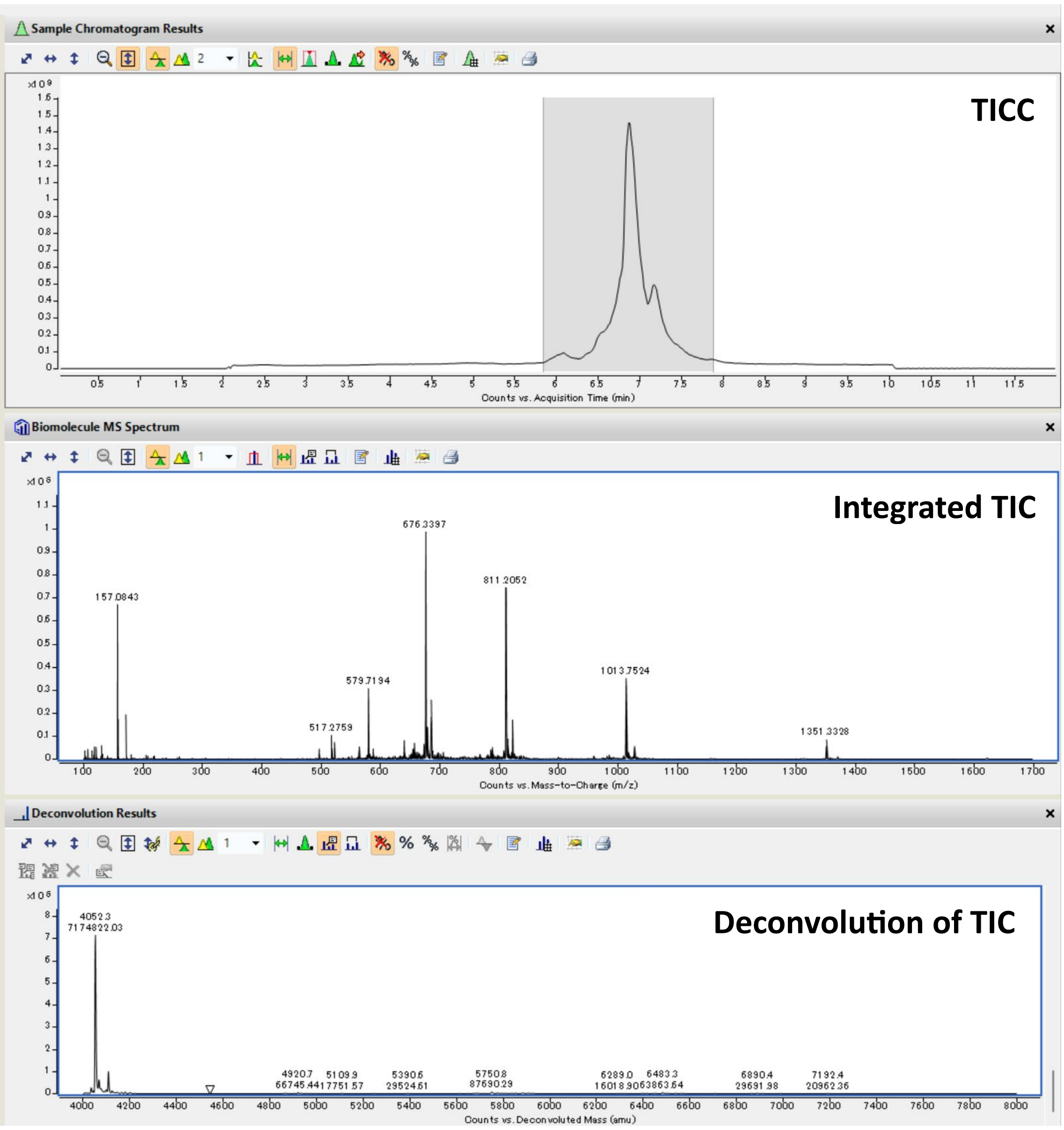

TICC: Total Ion Current Chromatogram; TIC: Total Ion Current.

Peptide: **BB-5**

Structure:

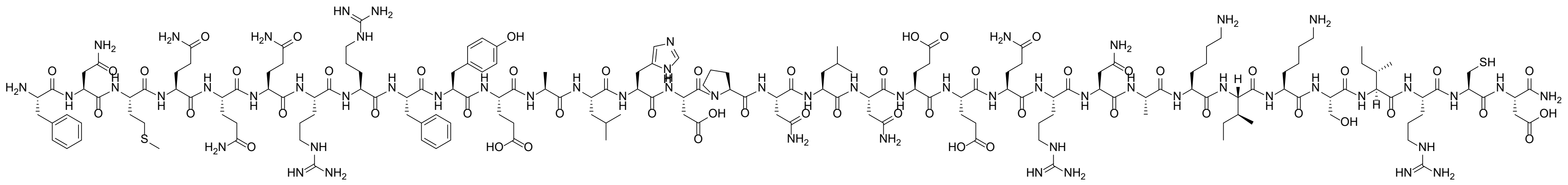

LC-MS analysis: method A

Calculated Mass: 4092.6 Da

Observed Mass: 4092.7 Da

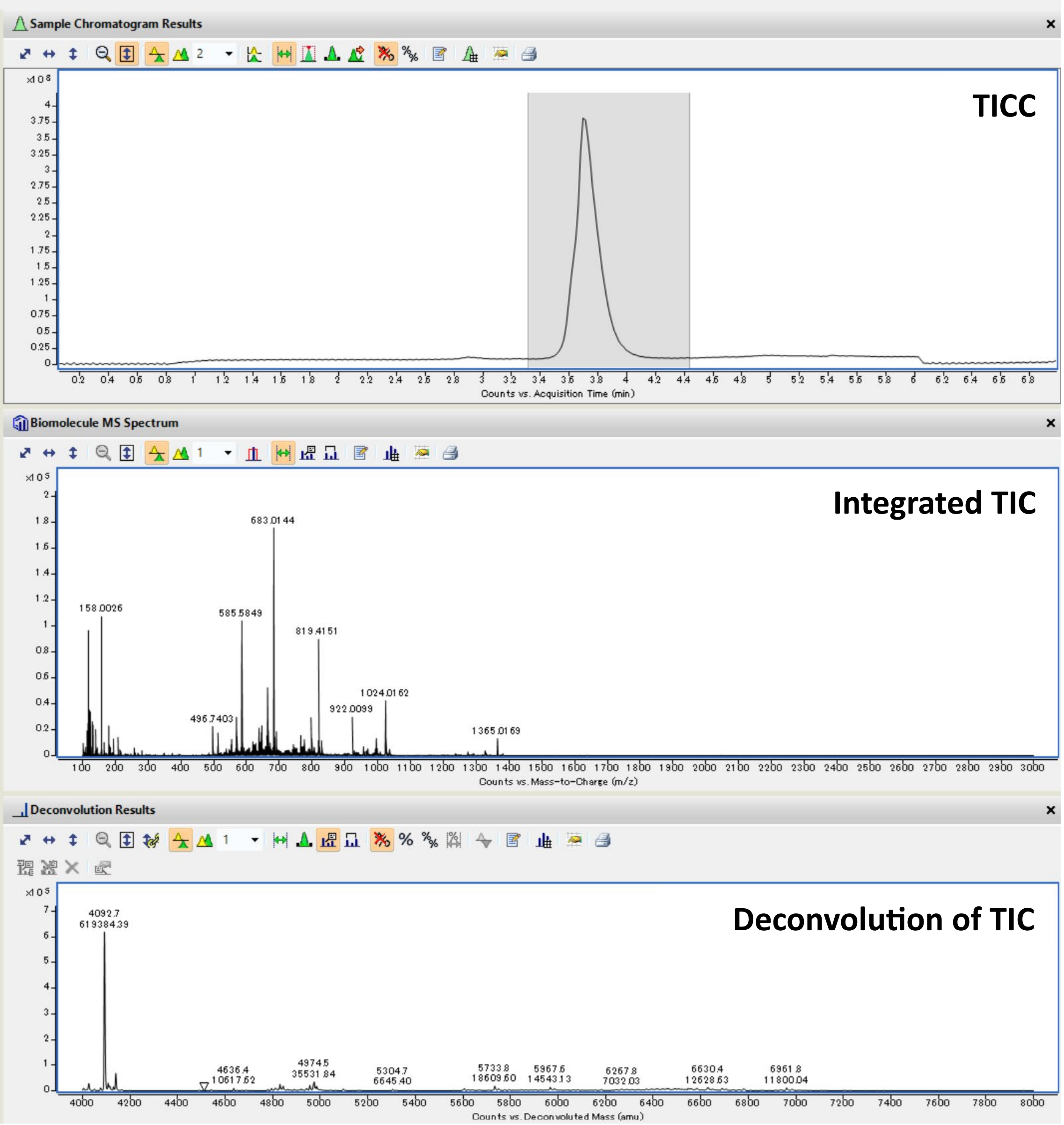

TICC: Total Ion Current Chromatogram; TIC: Total Ion Current.

Peptide: **BB-6**

Structure:

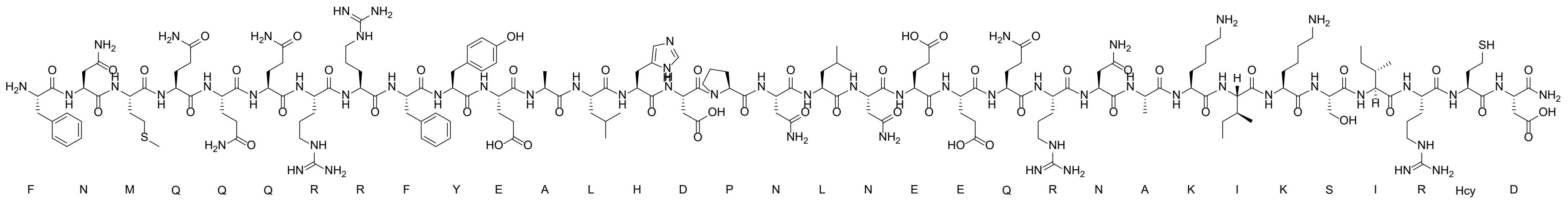

LC-MS analysis: Method E

Calculated Mass: 4106.6 Da

Observed Mass: 4106.5 Da

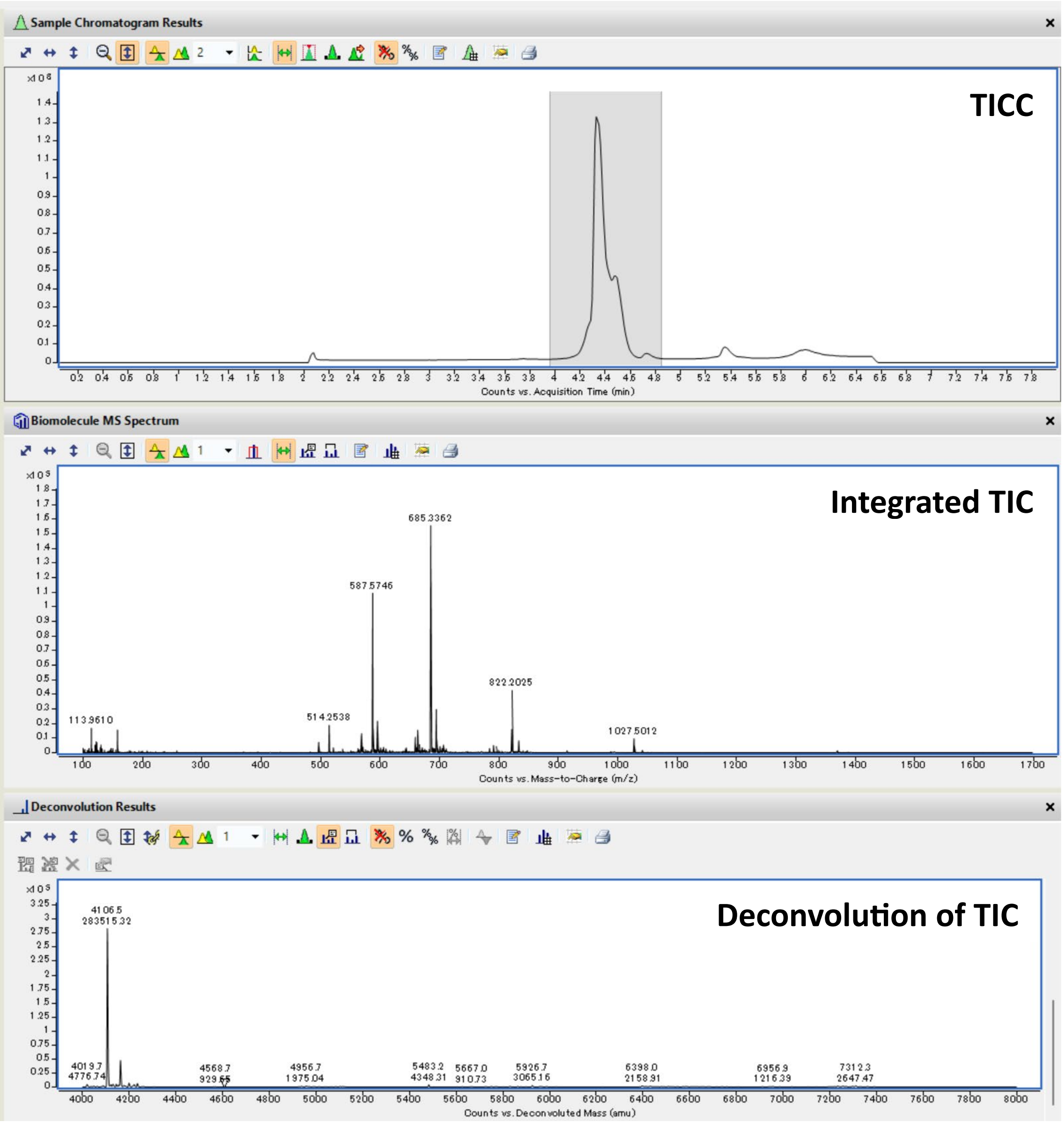

TICC: Total Ion Current Chromatogram; TIC: Total Ion Current.

Peptide: **N$_3$-1-I**

Structure:

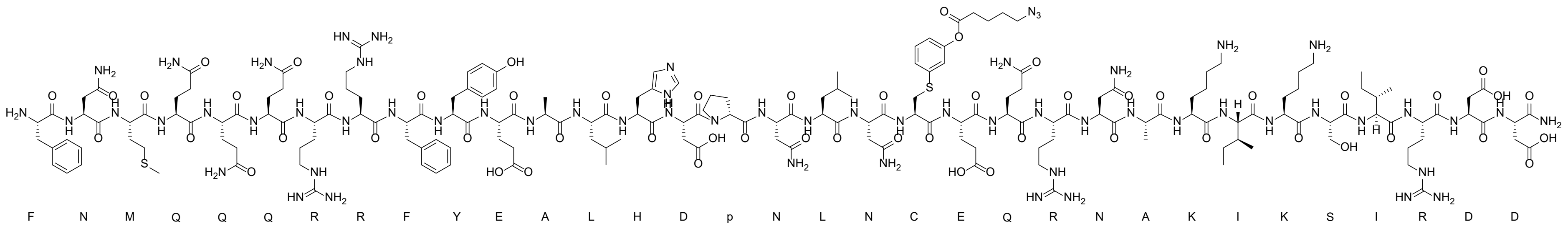

LC-MS analysis: Method A

Calculated Mass: 4295.8 Da

Observed Mass: 4296.1 Da

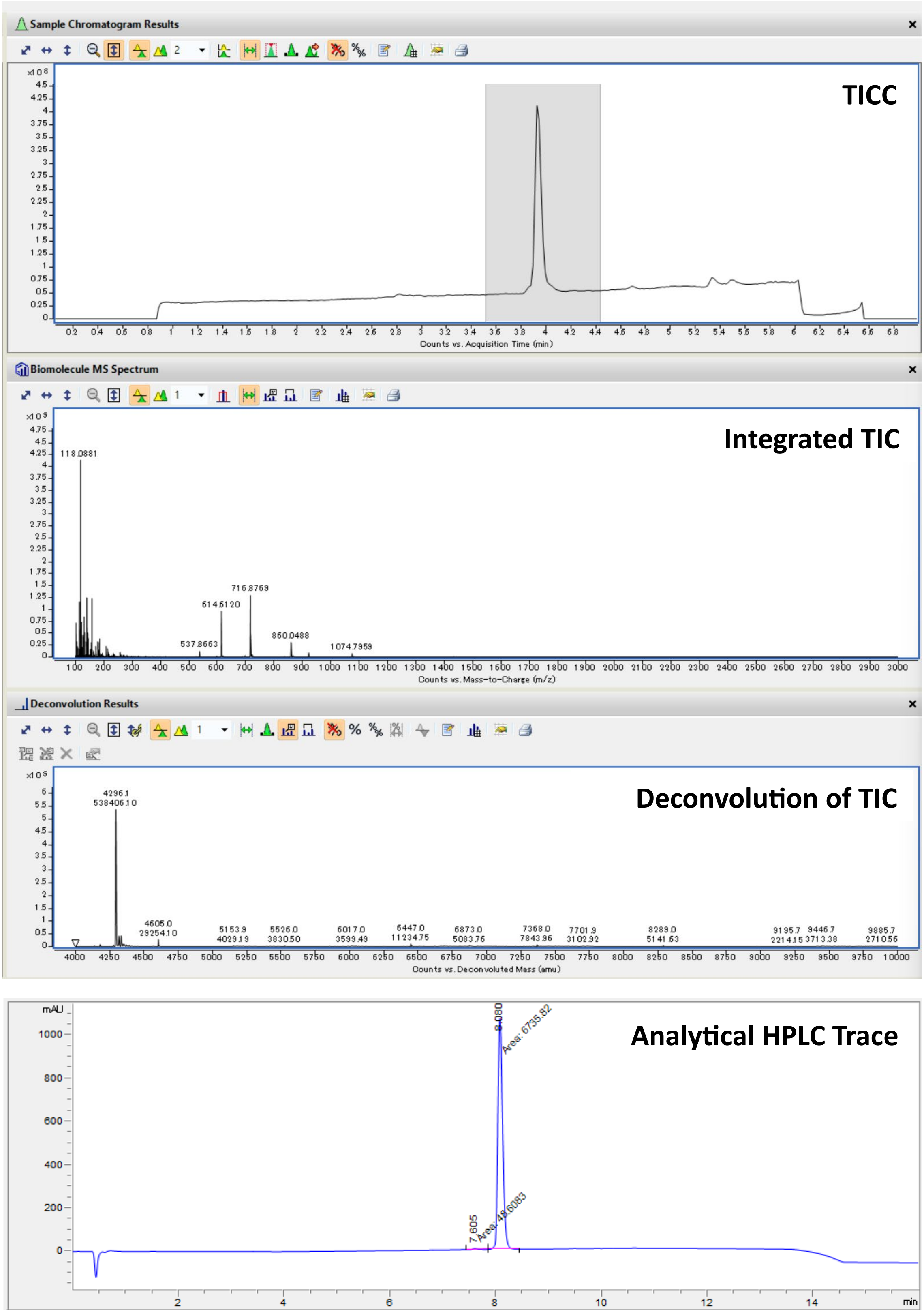

TICC: Total Ion Current Chromatogram; TIC: Total Ion Current.

Peptide: **$N_3$-2-I**

Structure:

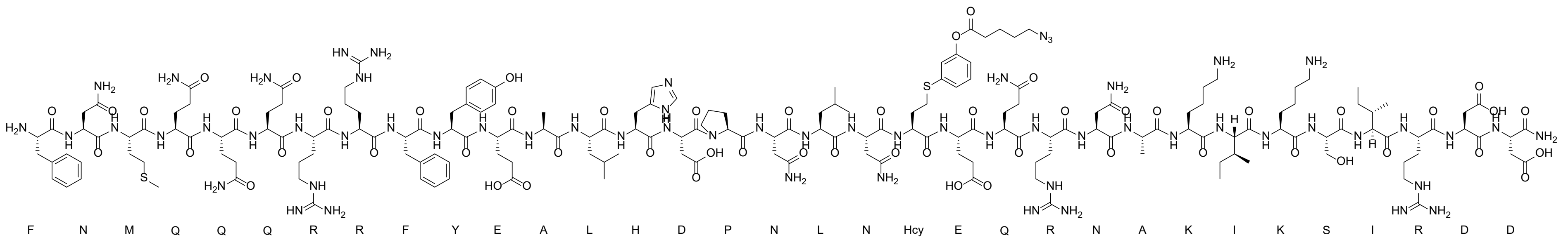

LC-MS analysis: Method A

Calculated Mass: 4309.8 Da

Observed Mass: 4310.0 Da

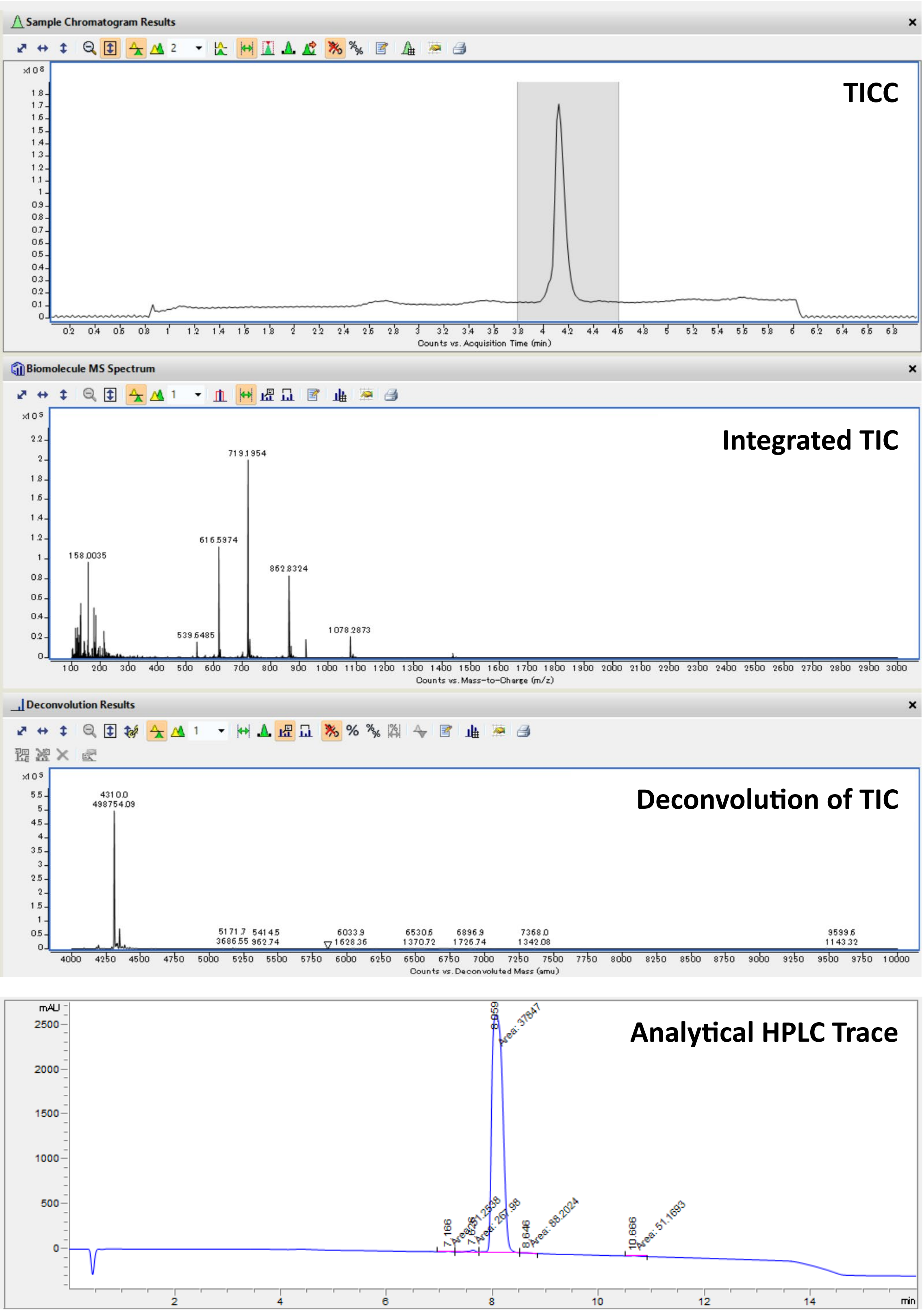

TICC: Total Ion Current Chromatogram; TIC: Total Ion Current.

Peptide: **$N_3$-3-I**

Structure:

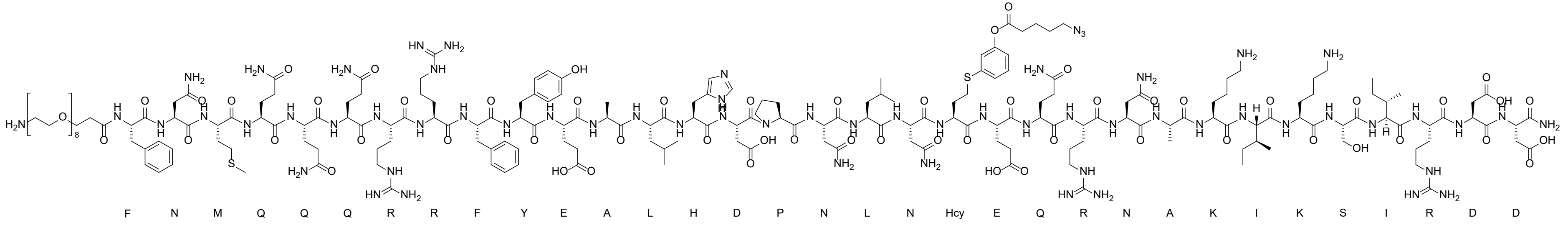

LC-MS analysis: Method D

Calculated Mass: 4733.3 Da

Observed Mass: 4733.3 Da

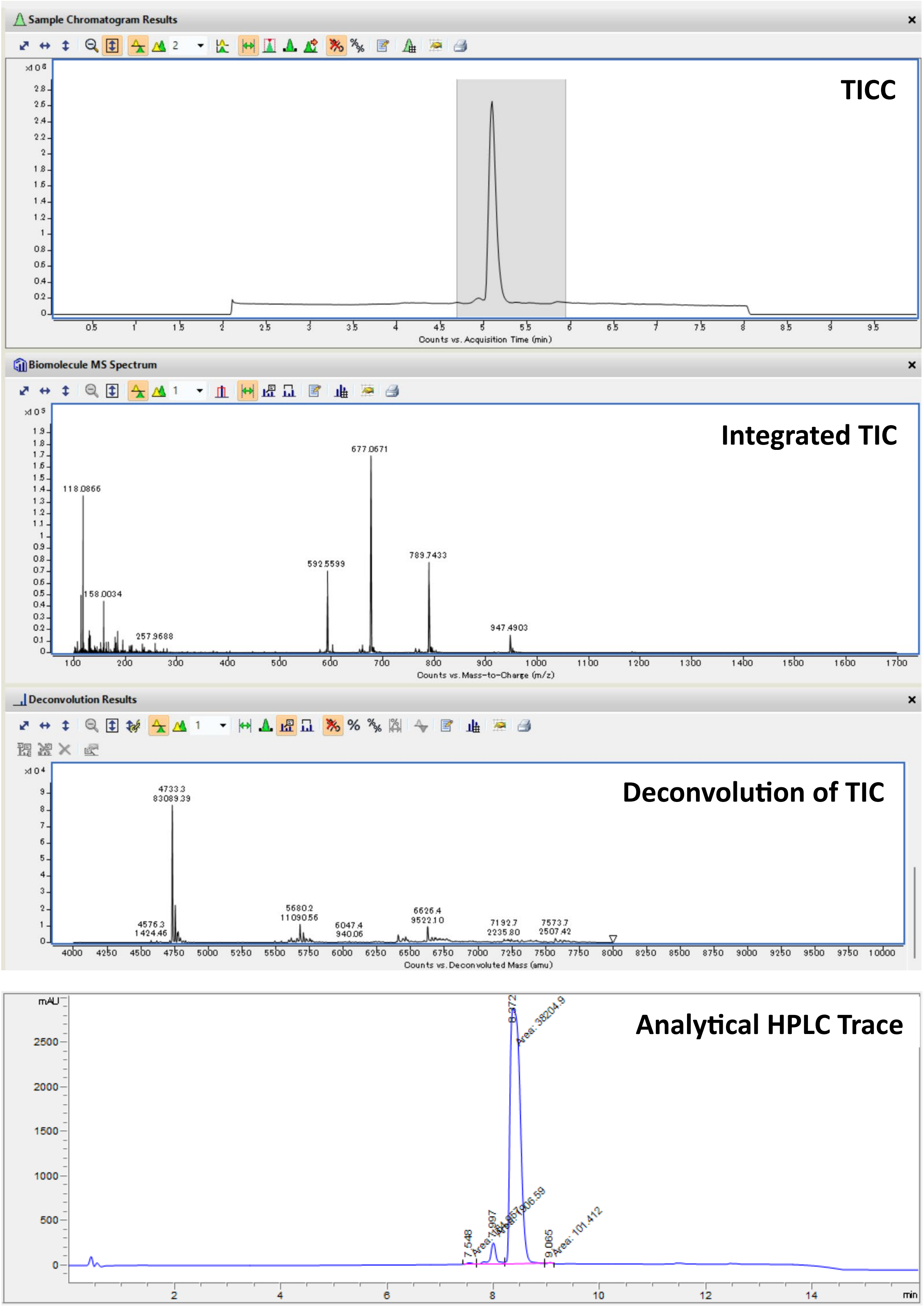

TICC: Total Ion Current Chromatogram; TIC: Total Ion Current.

Peptide: **$N_3$-4-I**

Structure:

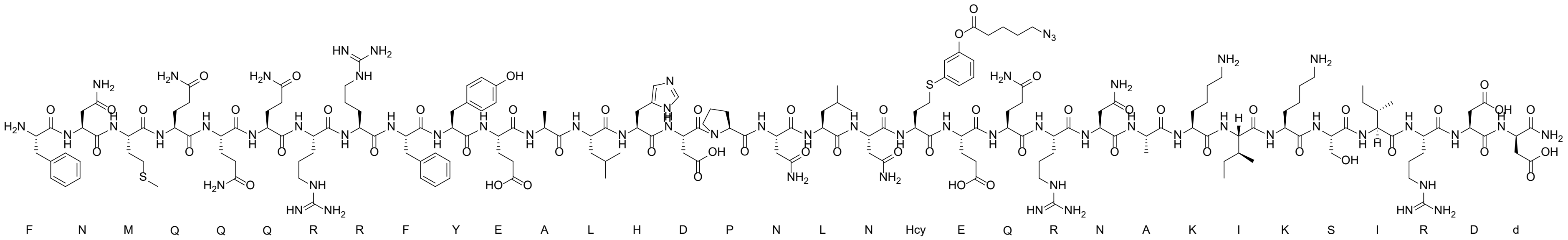

LC-MS analysis: Method E

Calculated Mass: 4309.8 Da

Observed Mass: 4309.7 Da

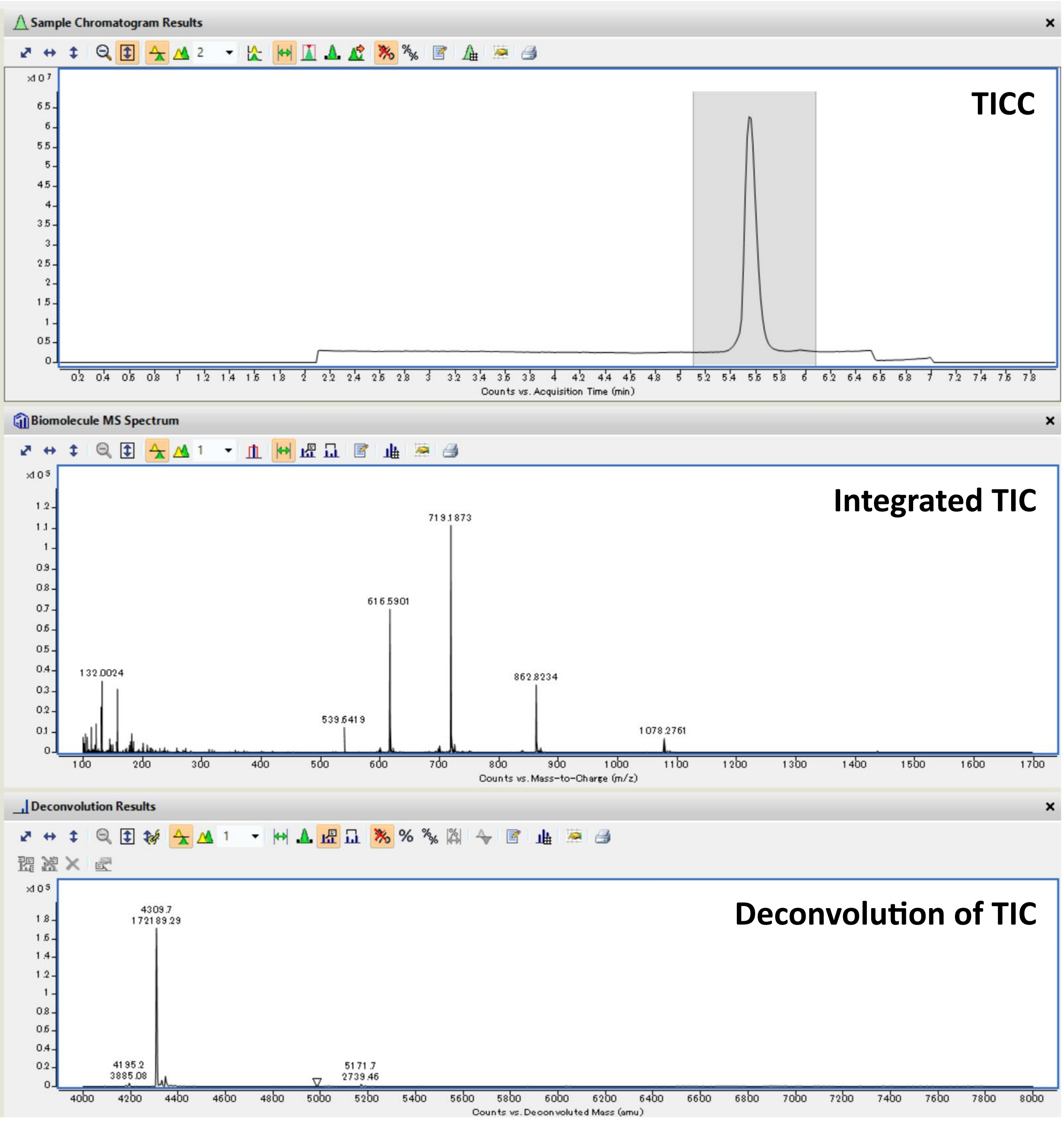

TICC: Total Ion Current Chromatogram; TIC: Total Ion Current.

Peptide: **$N_3$-5-I**

Structure:

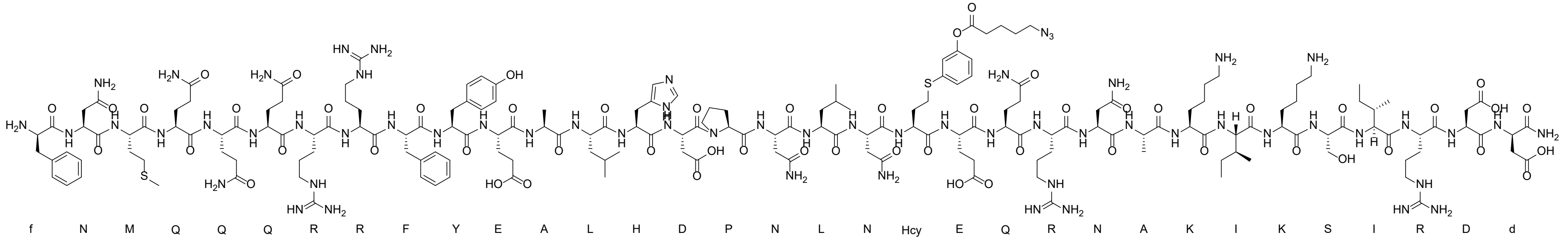

LC-MS analysis: Method E

Calculated Mass: 4309.8 Da

Observed Mass: 4309.7 Da

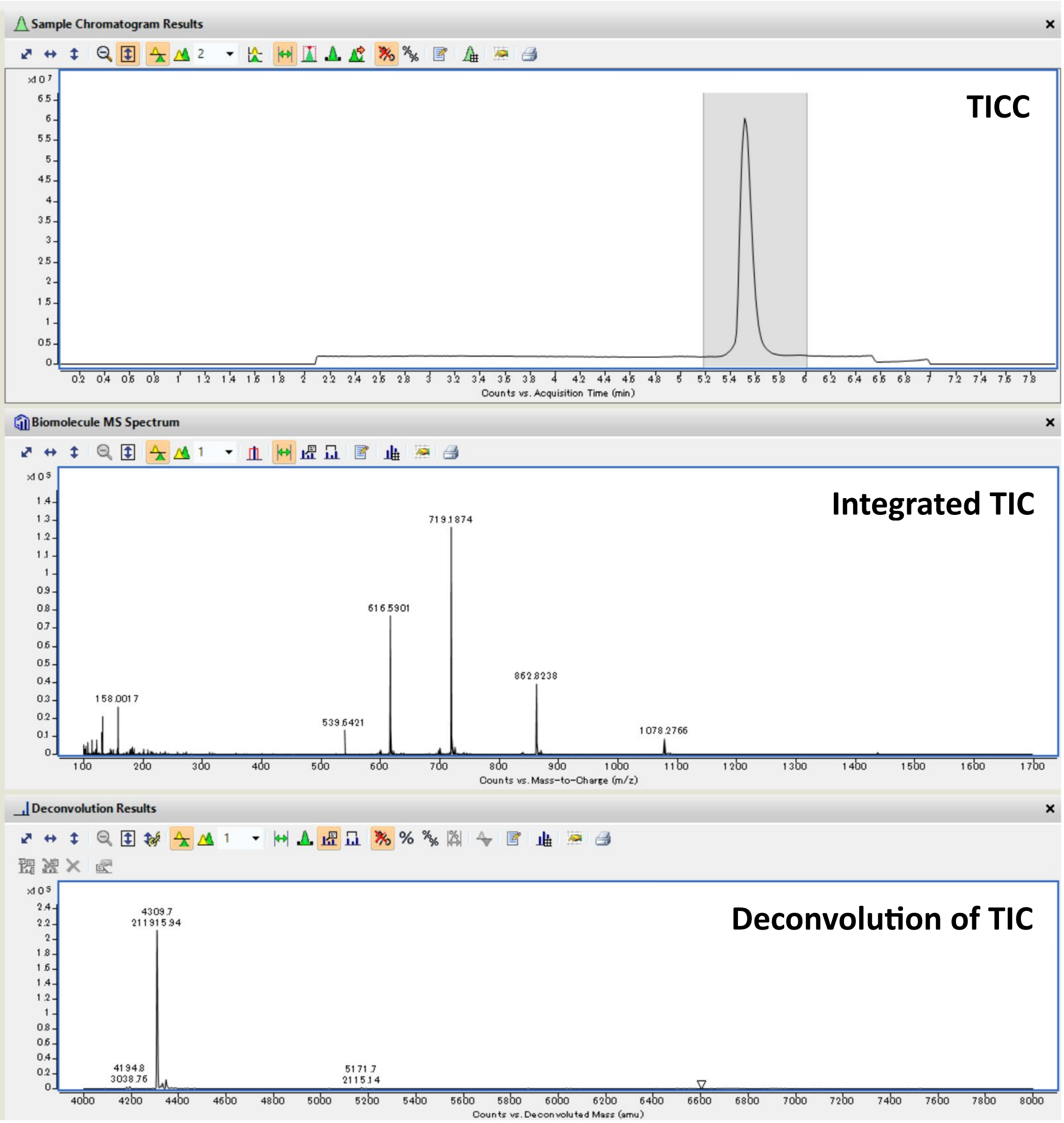

TICC: Total Ion Current Chromatogram; TIC: Total Ion Current.

Peptide: **$N_3$-6-I**

Structure:

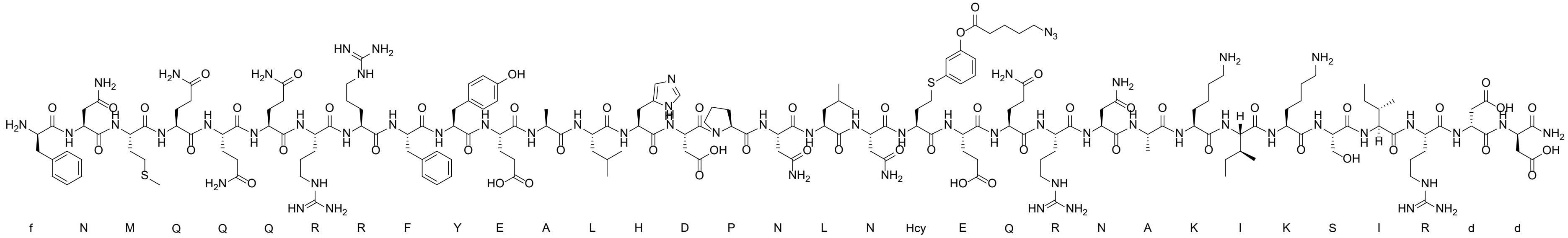

LC-MS analysis: Method E

Calculated Mass: 4309.8 Da

Observed Mass: 4309.7 Da

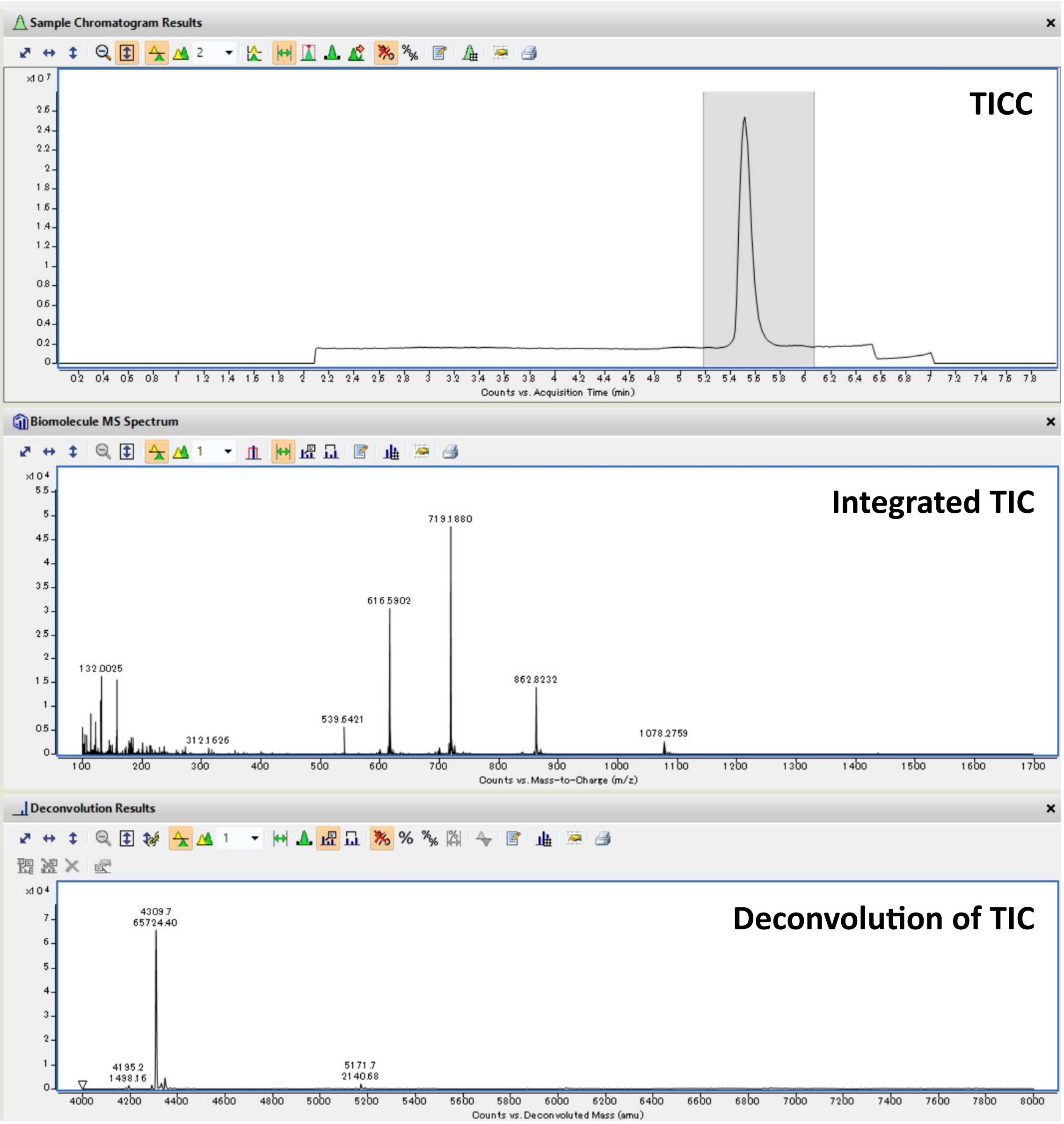

TICC: Total Ion Current Chromatogram; TIC: Total Ion Current.

Peptide: **BB-21**

Structure:

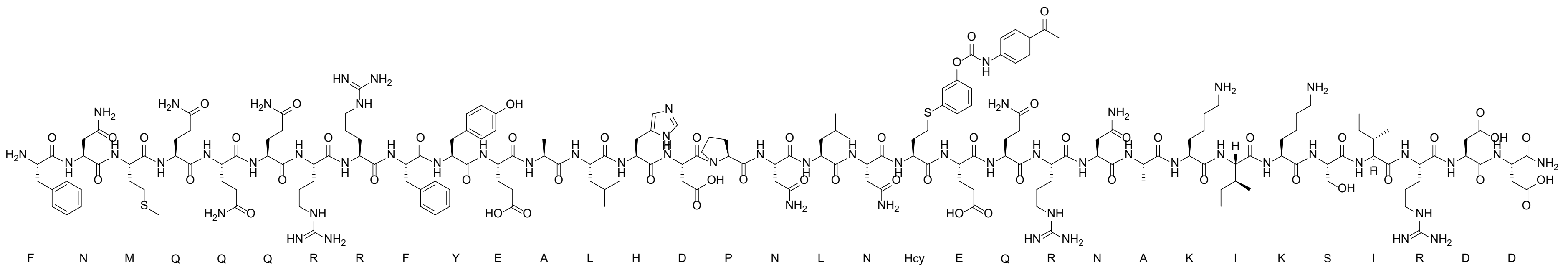

LC-MS analysis: Method E

Calculated Mass: 4345.9 Da

Observed Mass: 4345.7 Da

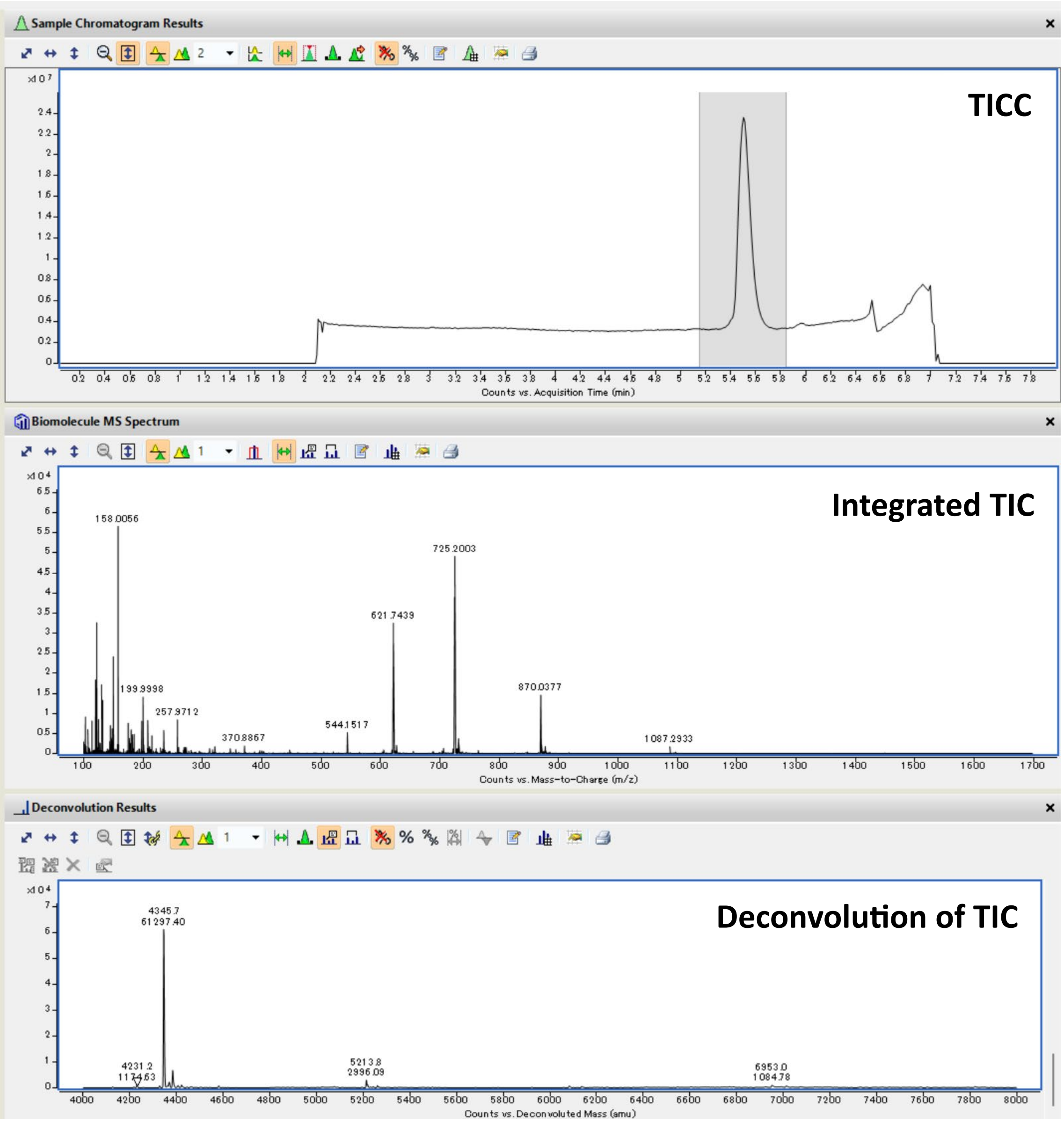

TICC: Total Ion Current Chromatogram; TIC: Total Ion Current.

Peptide: **BB-22**

Structure:

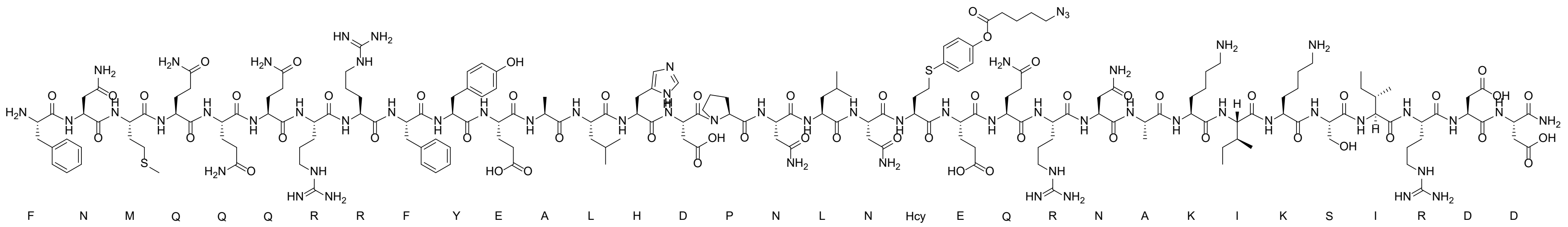

LC-MS analysis: Method E

Calculated Mass: 4309.8 Da

Observed Mass: 4309.7 Da

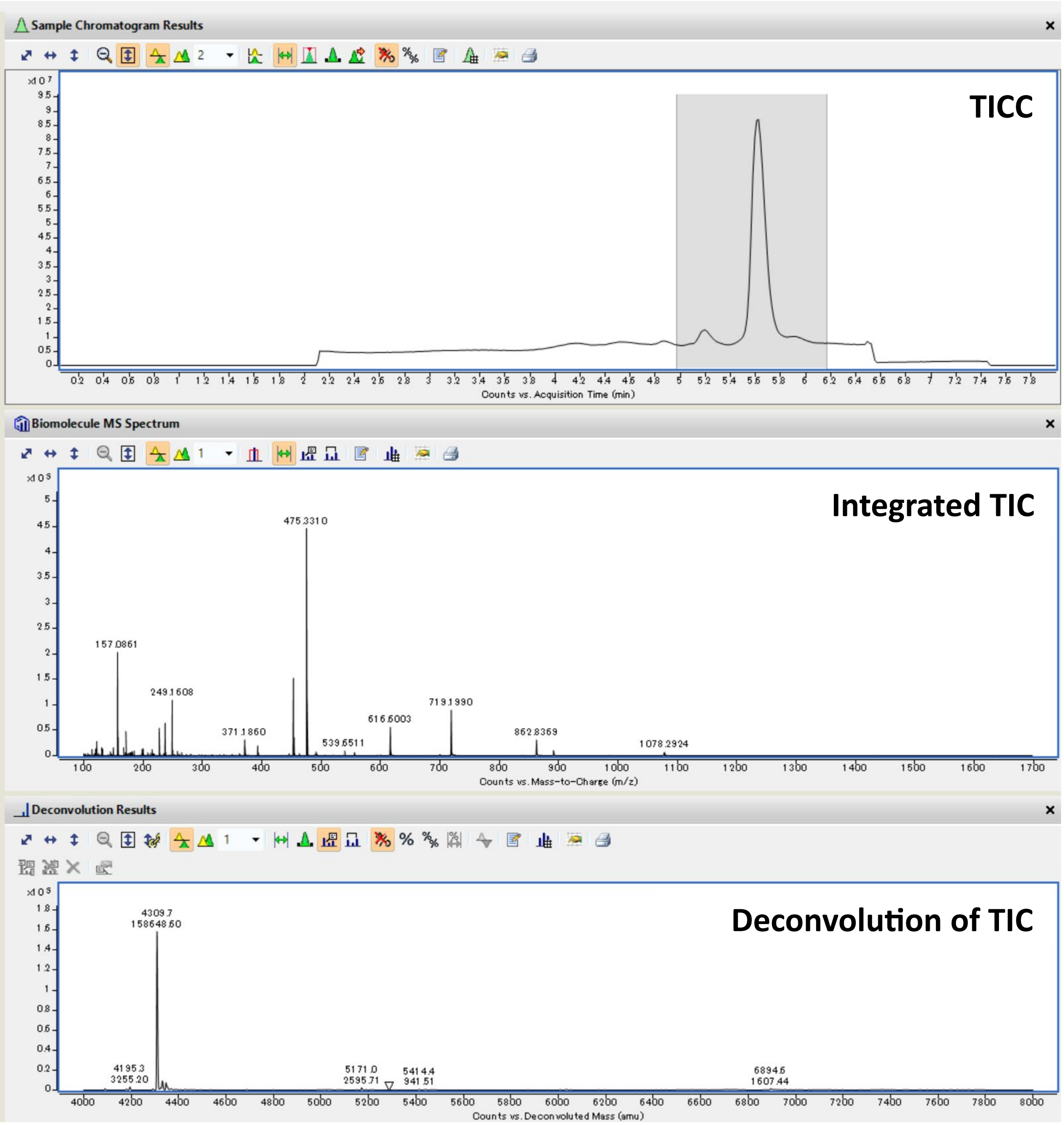

TICC: Total Ion Current Chromatogram; TIC: Total Ion Current.

Peptide: **BB-23**

Structure:

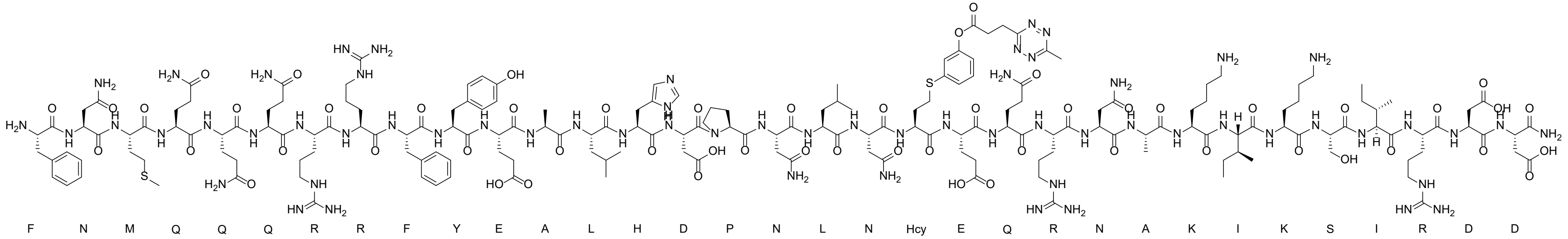

LC-MS analysis: Method E

Calculated Mass: 4334.8 Da

Observed Mass: 4334.7 Da

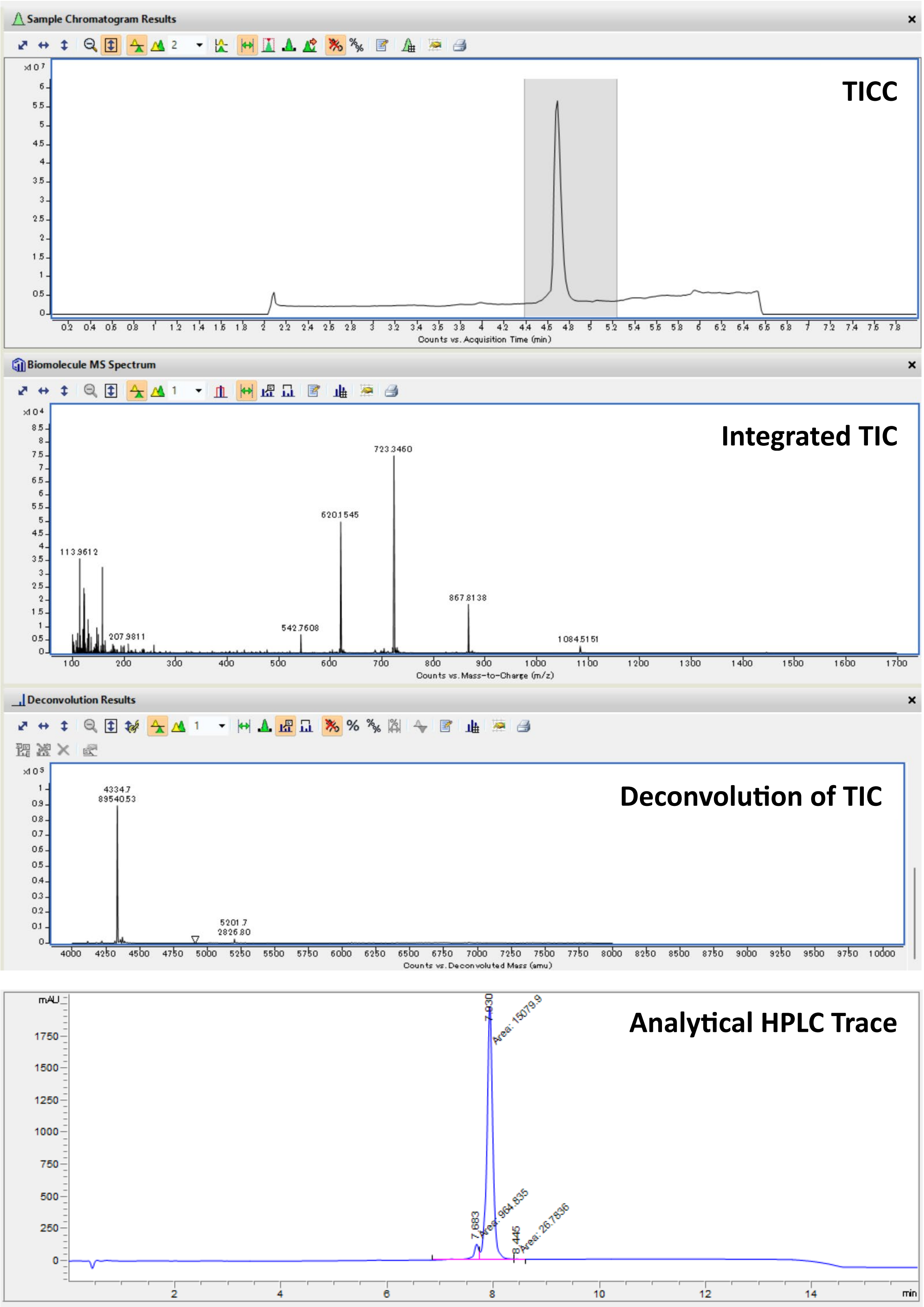

TICC: Total Ion Current Chromatogram; TIC: Total Ion Current.

Peptide: **$N_3$-12-VIII**

Structure:

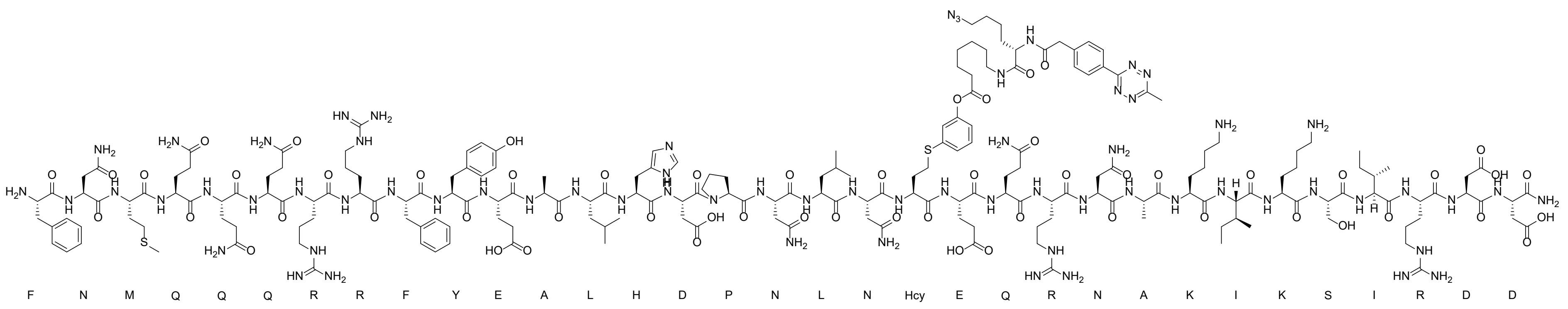

LC-MS analysis: Method A

Calculated Mass: 4678.3 Da

Observed Mass: 4678.4 Da

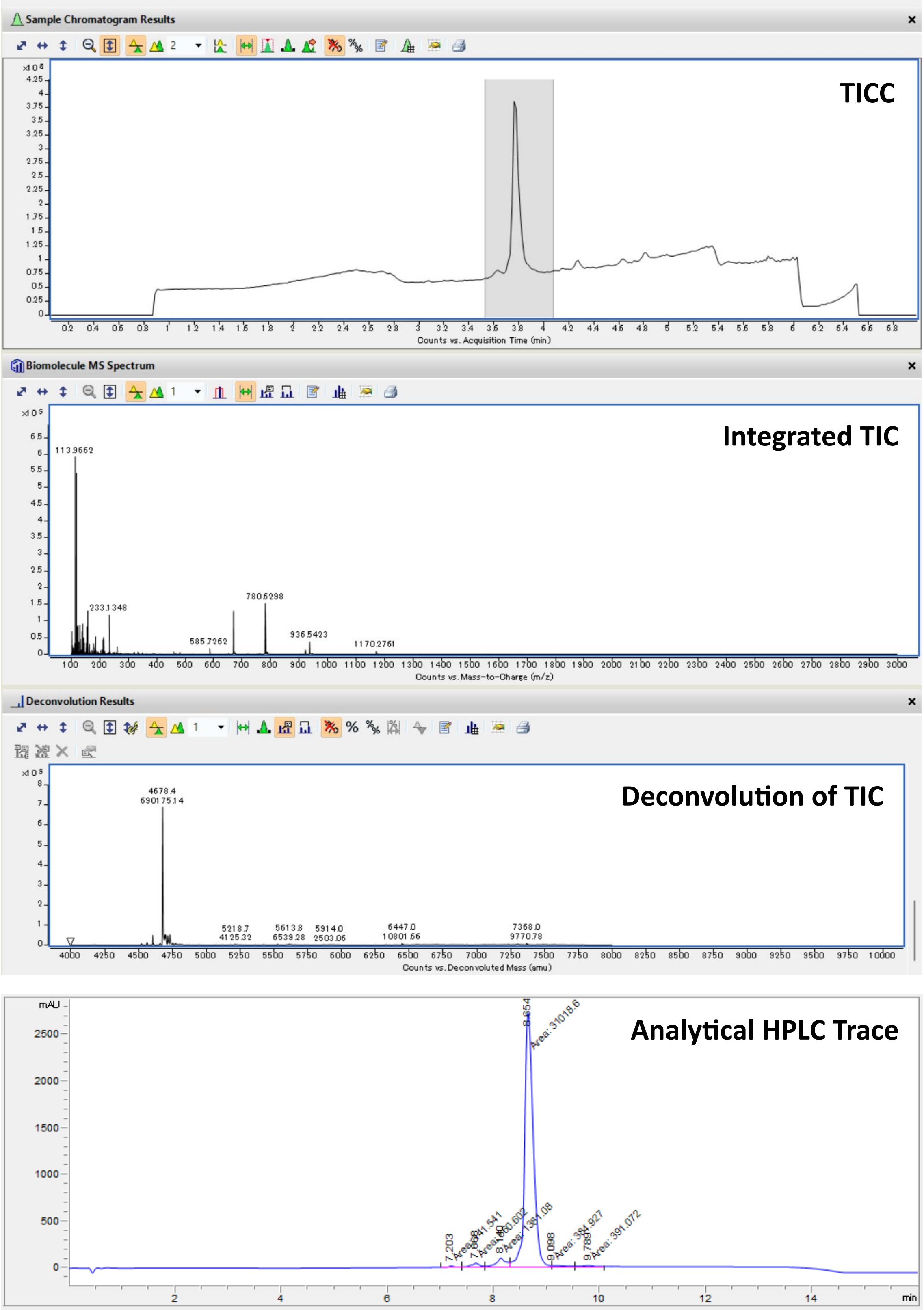

TICC: Total Ion Current Chromatogram; TIC: Total Ion Current.

Peptide: **$N_3$-7-VI**

Structure:

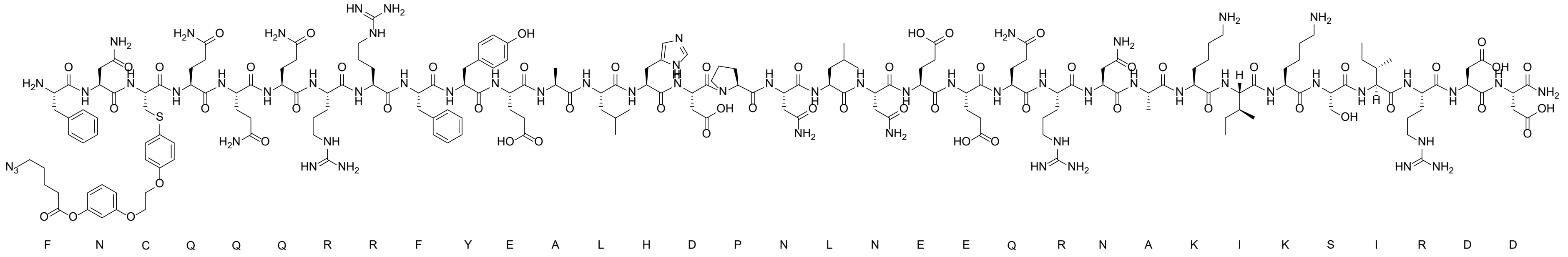

LC-MS analysis: Method A

Calculated Mass: 4429.9 Da

Observed Mass: 4430.1 Da

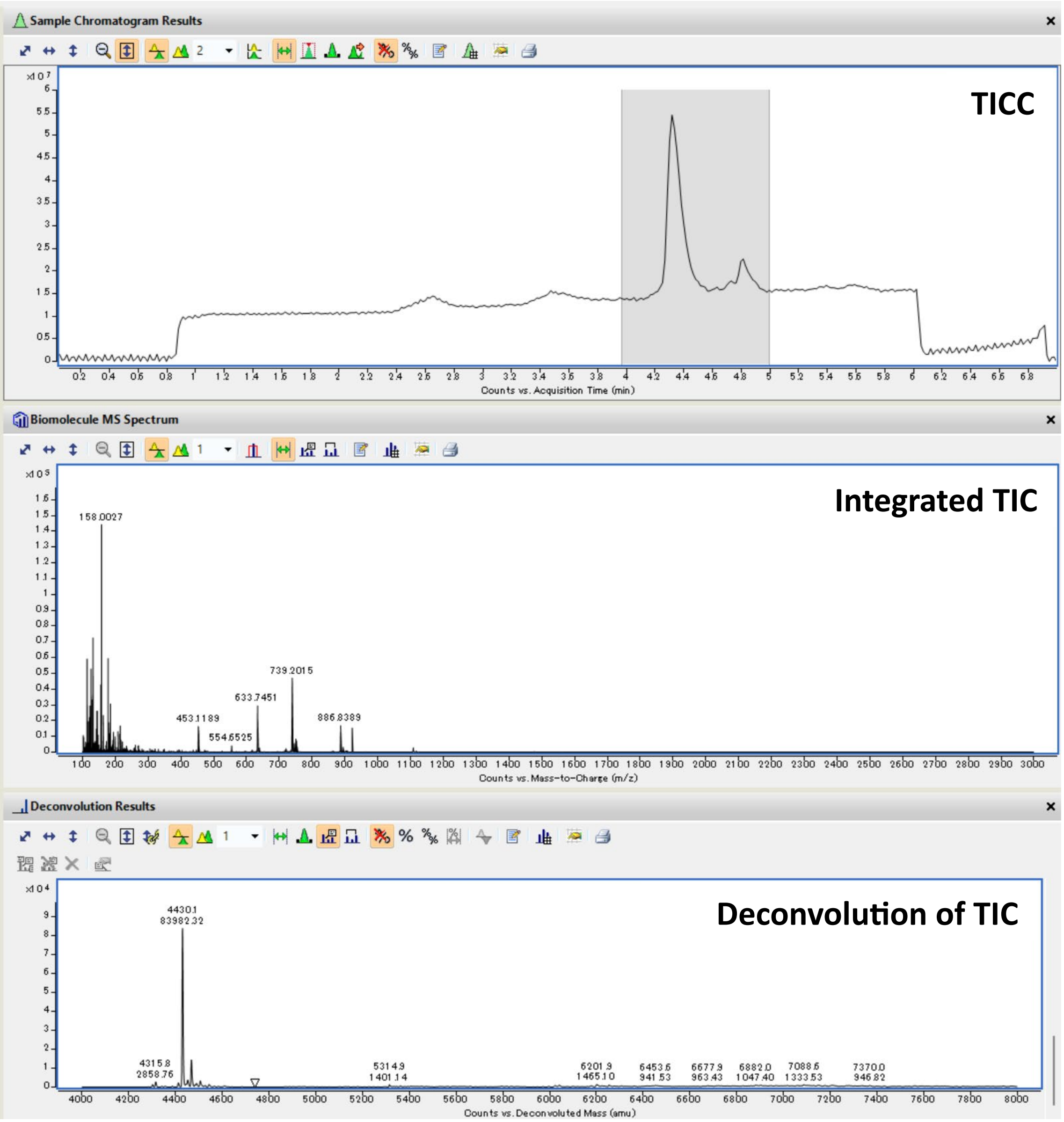

TICC: Total Ion Current Chromatogram; TIC: Total Ion Current.

Peptide: **$N_3$-8-VI**

Structure:

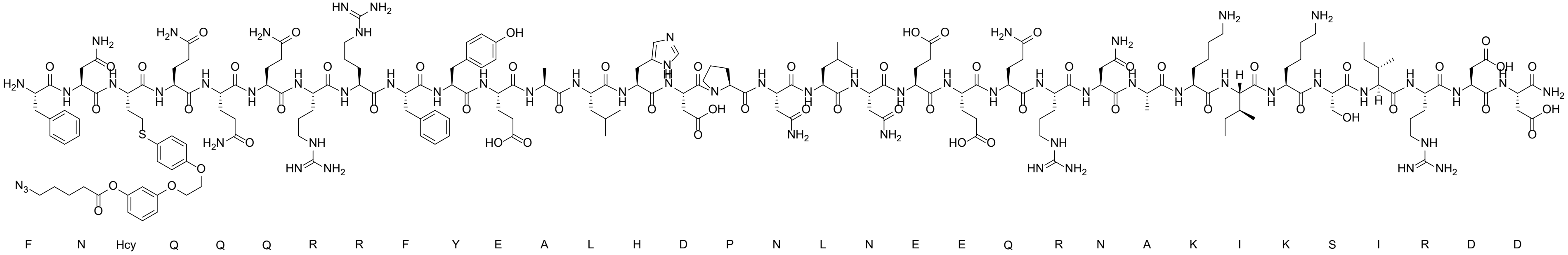

LC-MS analysis: Method E

Calculated Mass: 4443.9 Da

Observed Mass: 4443.8 Da

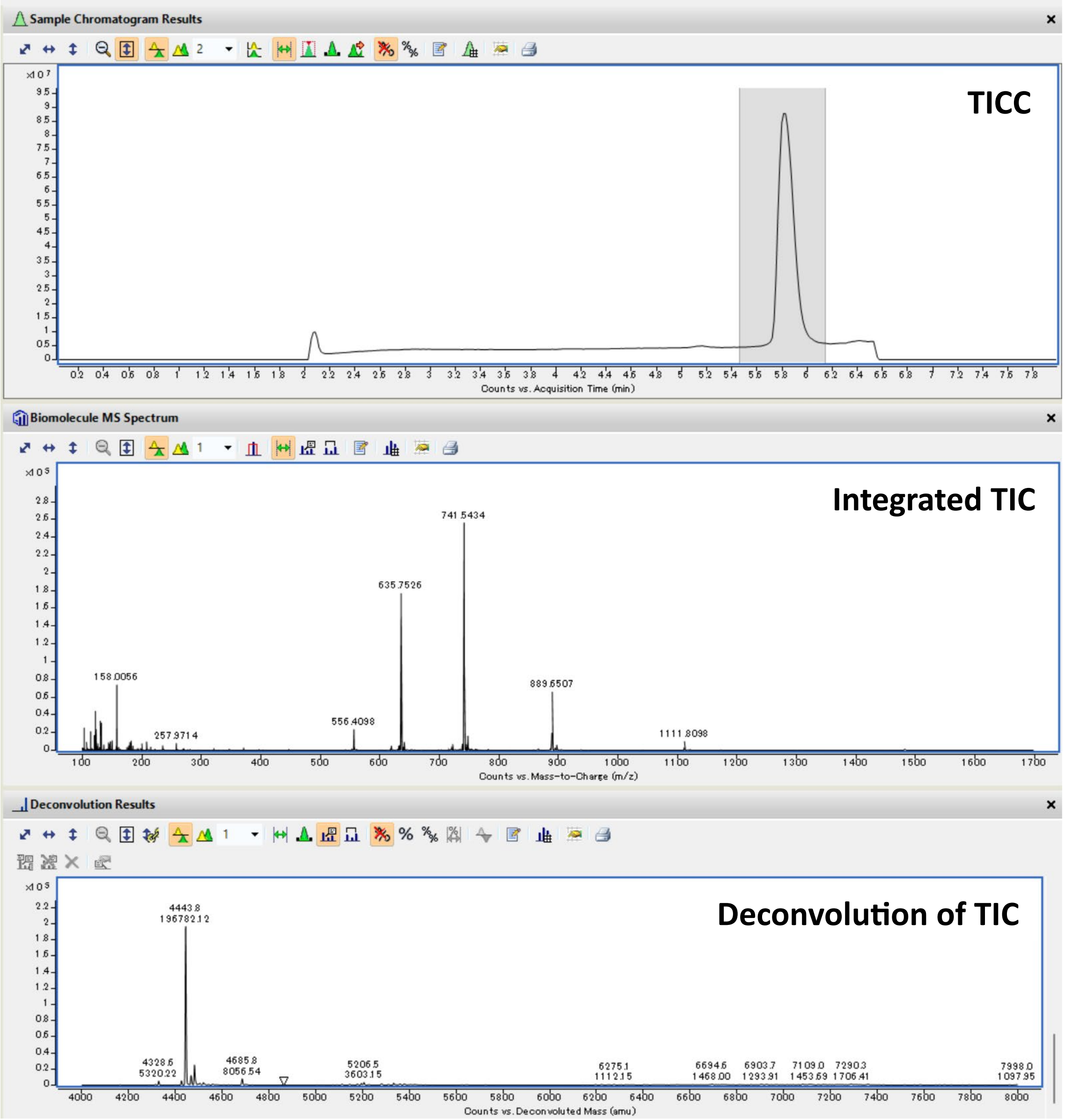

TICC: Total Ion Current Chromatogram; TIC: Total Ion Current.

Peptide: **$N_3$-7-VII**

Structure:

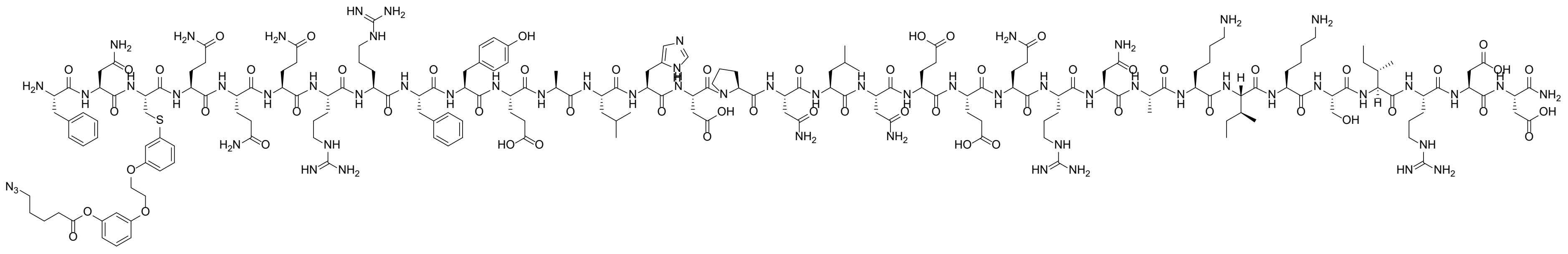

LC-MS analysis: Method E

Calculated Mass: 4429.9 Da

Observed Mass: 4430.6 Da

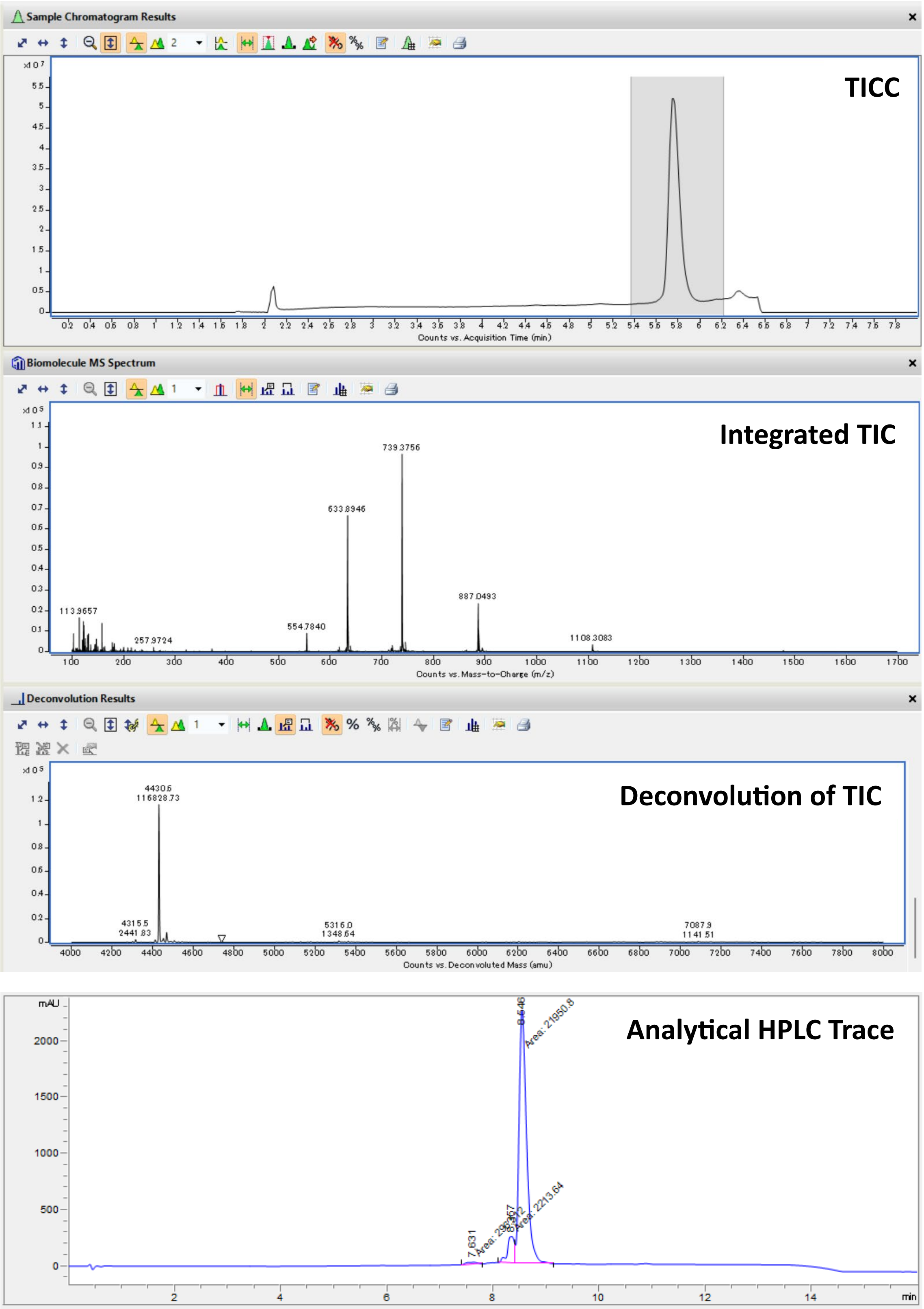

TICC: Total Ion Current Chromatogram; TIC: Total Ion Current.

Peptide: **$N_3$-8-VII**

Structure:

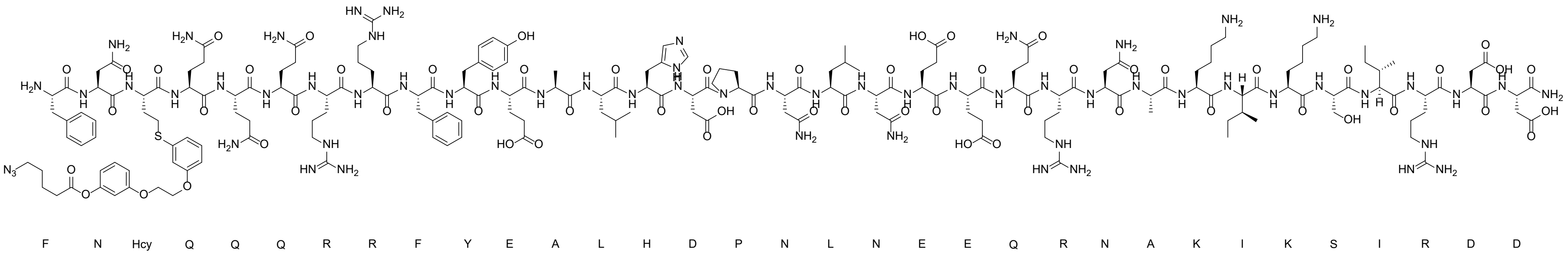

LC-MS analysis: Method E

Calculated Mass:4443.9 Da

Observed Mass: 4444.4 Da

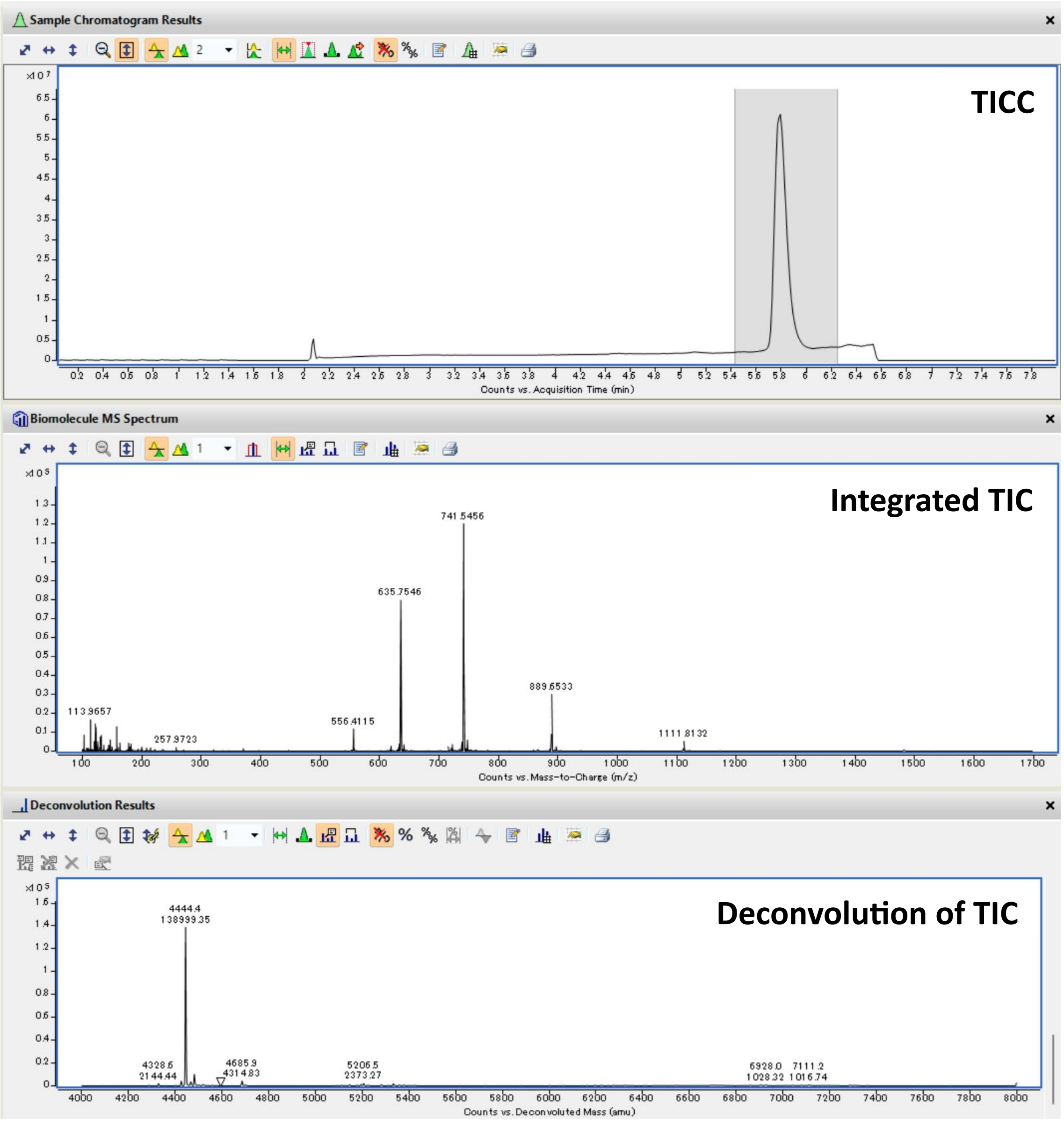

TICC: Total Ion Current Chromatogram; TIC: Total Ion Current.

Peptide: **$N_3$-9-VII**

Structure:

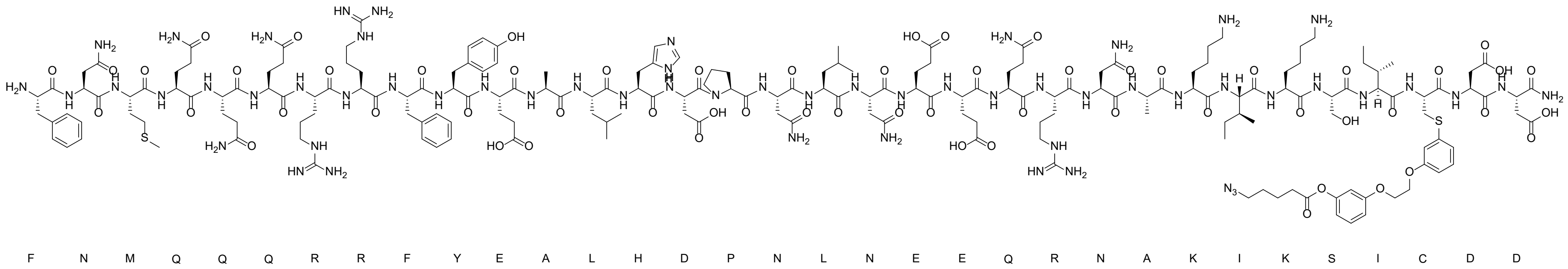

LC-MS analysis: Method B

Calculated Mass: 4404.9 Da

Observed Mass: 4405.8 Da

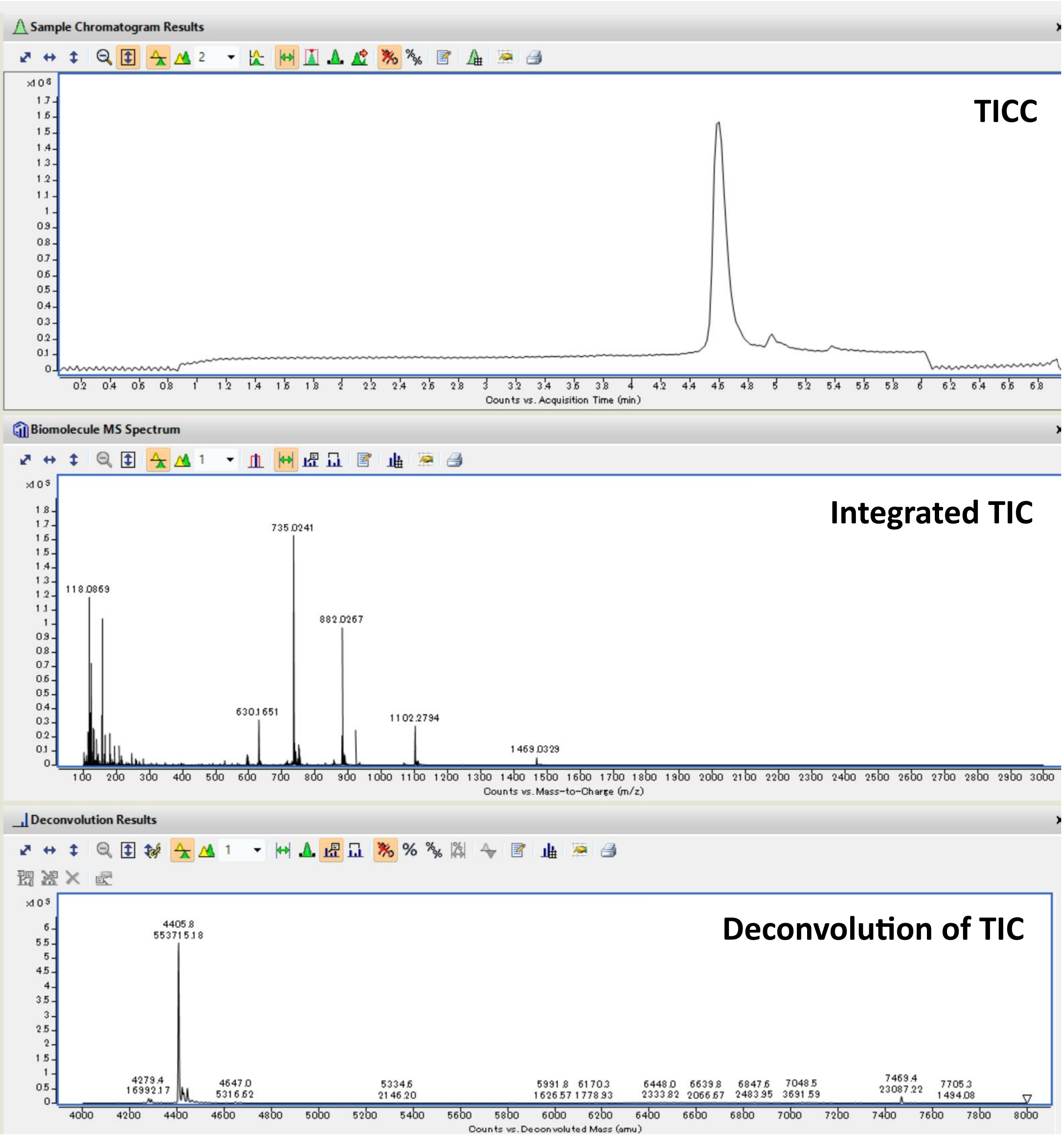

TICC: Total Ion Current Chromatogram; TIC: Total Ion Current.

Peptide: **$N_3$-10-VII**

Structure:

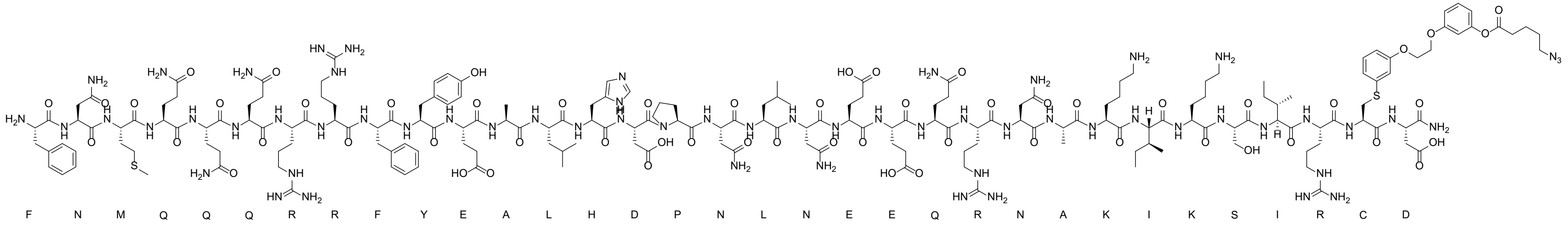

LC-MS analysis: Method B

Calculated Mass: 4446.0 Da

Observed Mass: 4445.7 Da

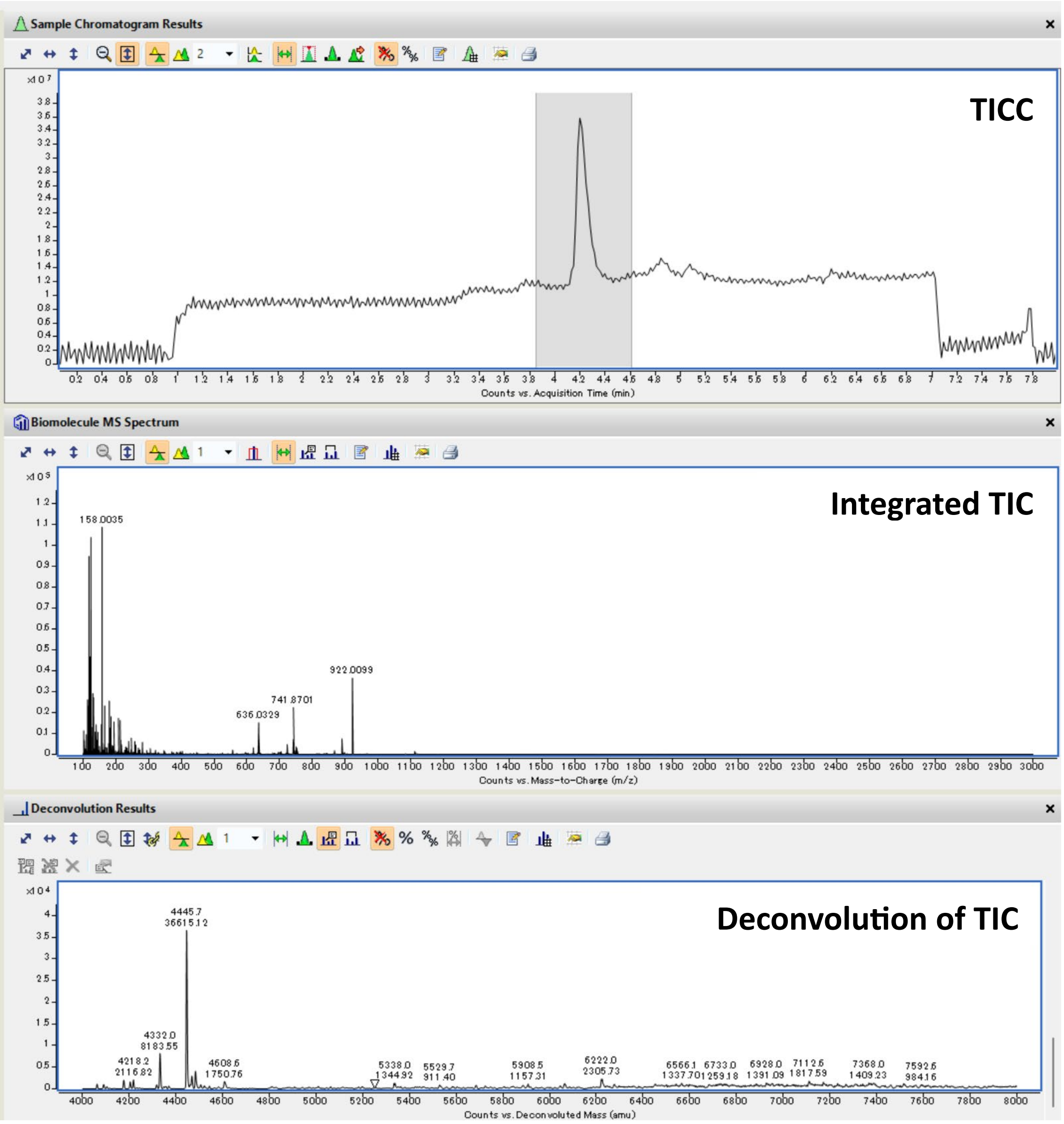

TICC: Total Ion Current Chromatogram; TIC: Total Ion Current.

Peptide: **$N_3$-11-VII**

Structure:

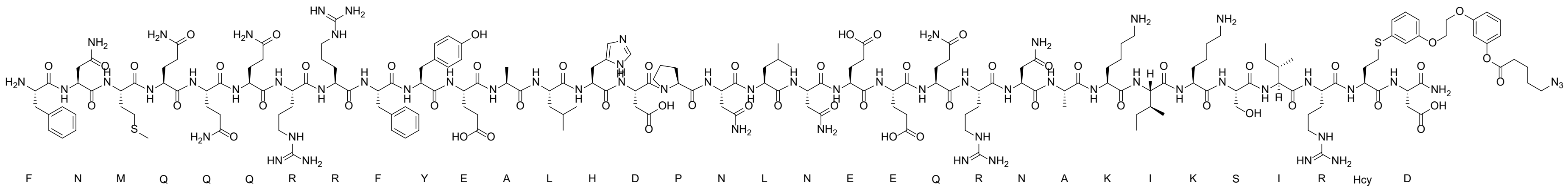

LC-MS analysis: Method E

Calculated Mass: 4460.0 Da

Observed Mass: 4460.2 Da

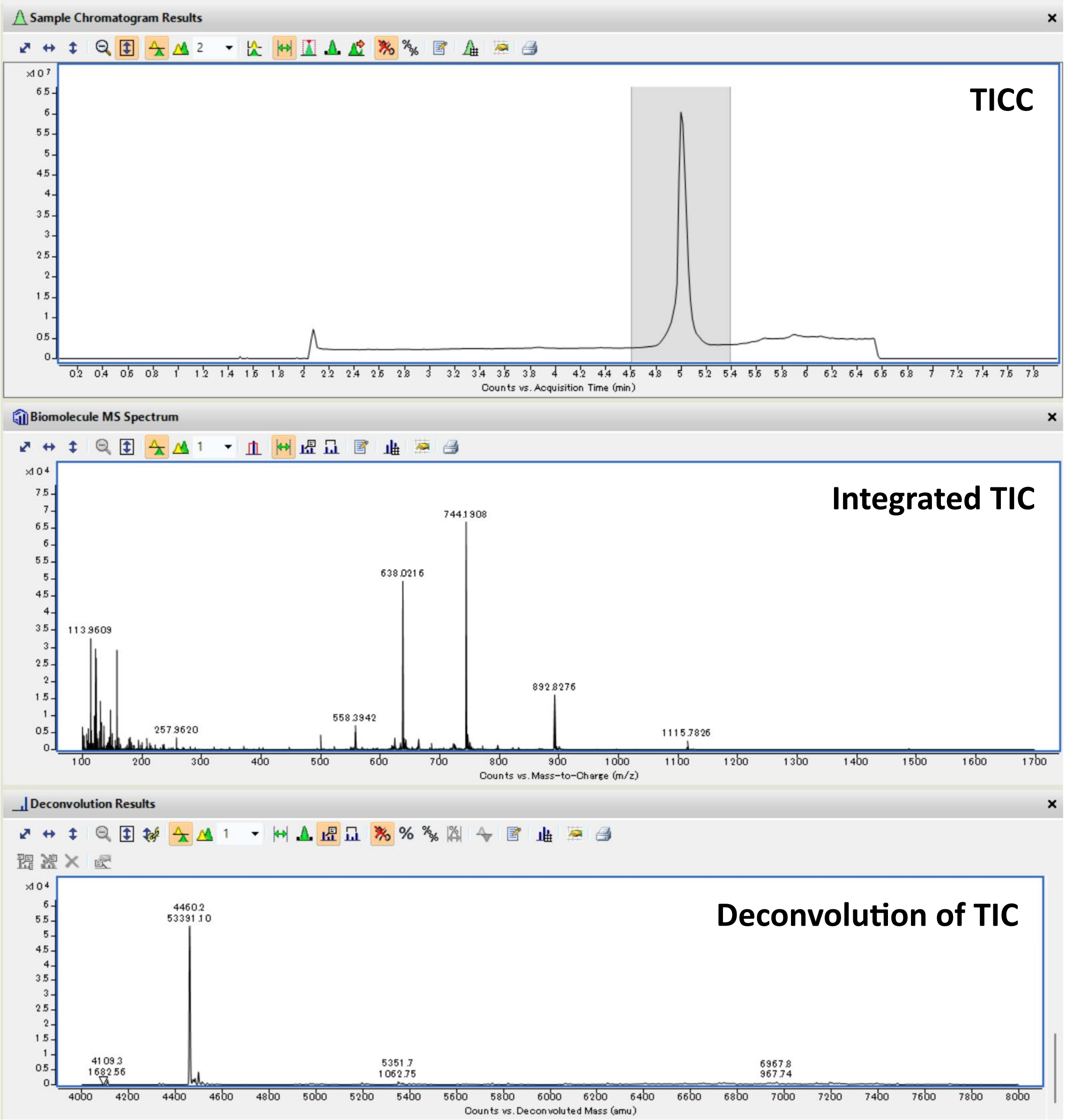

TICC: Total Ion Current Chromatogram; TIC: Total Ion Current.

Peptide: **$N_3$-13-VIII**

Structure:

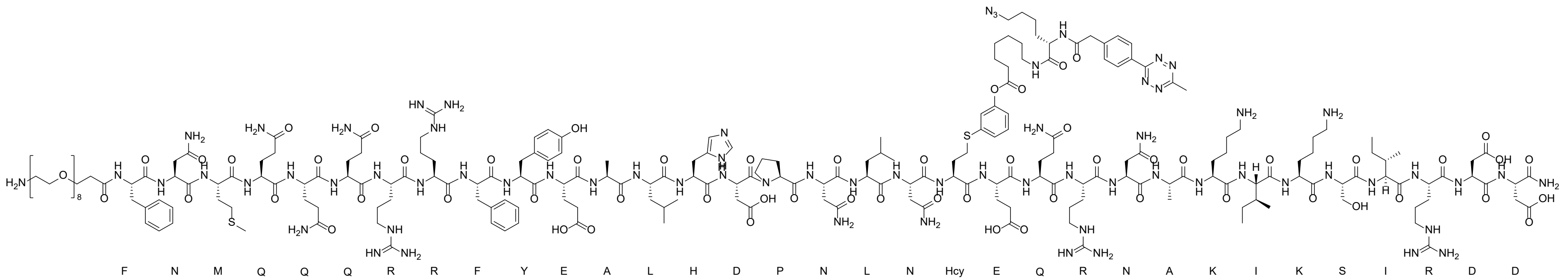

LC-MS analysis: Method A

Calculated Mass: 5101.8 Da

Observed Mass: 5102.2 Da

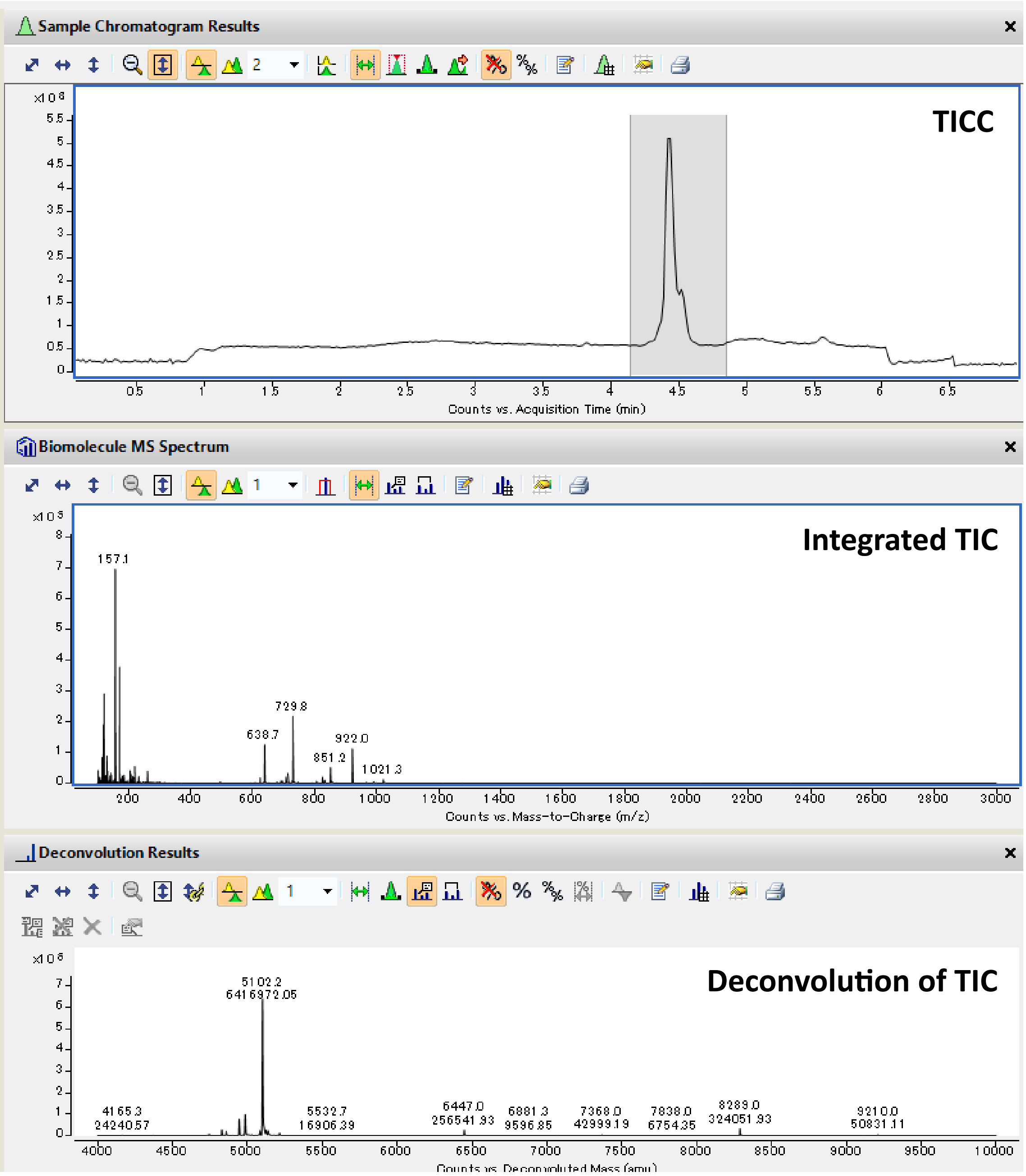

TICC: Total Ion Current Chromatogram; TIC: Total Ion Current.

Peptide: **12a**

Structure:

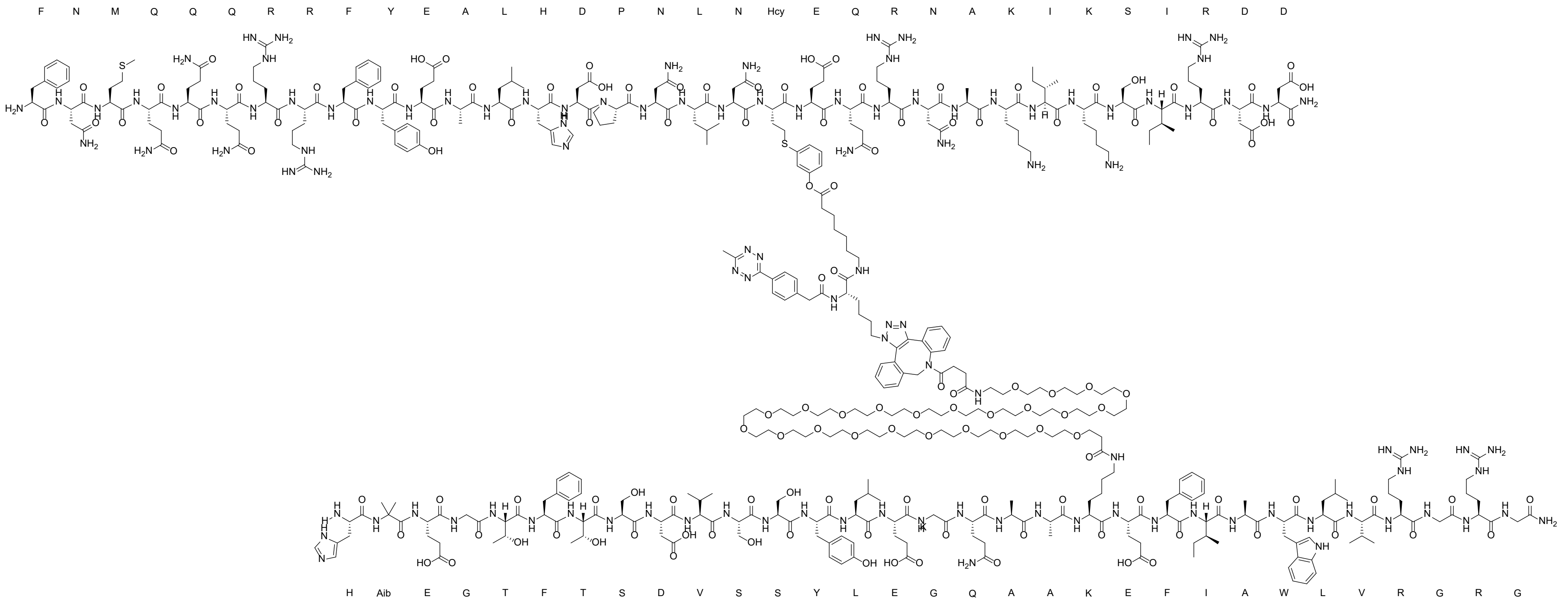

LC-MS analysis*: Method B (* This data was obtained using crude reaction mixture.)

Calculated Mass: 9490.7 Da

Observed Mass: 9491.0 Da

UHPLC analysis** (** This data was obtained using product after purification.)

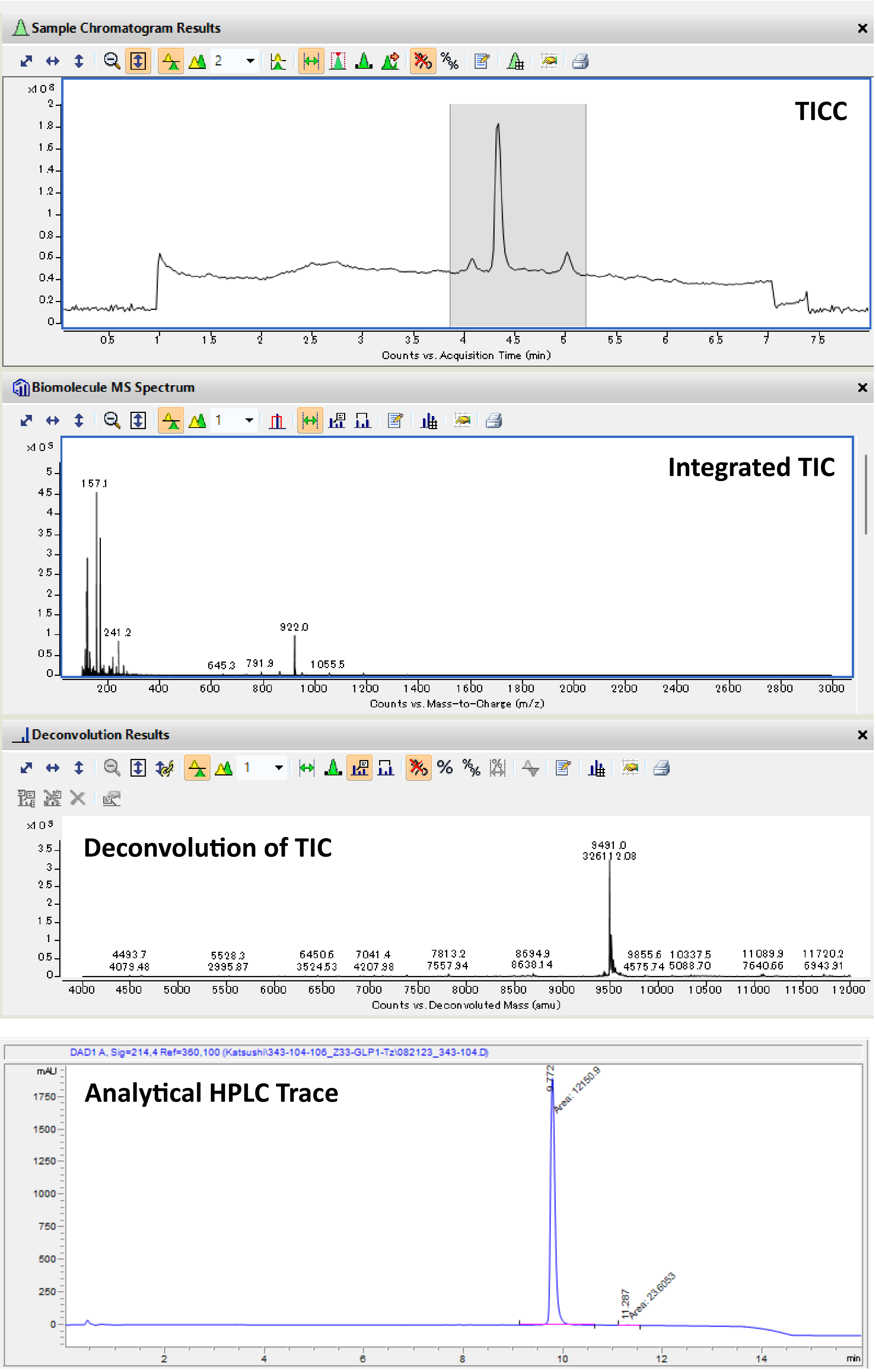

TICC: Total Ion Current Chromatogram; TIC: Total Ion Current.

Peptide: **13a**

Structure:

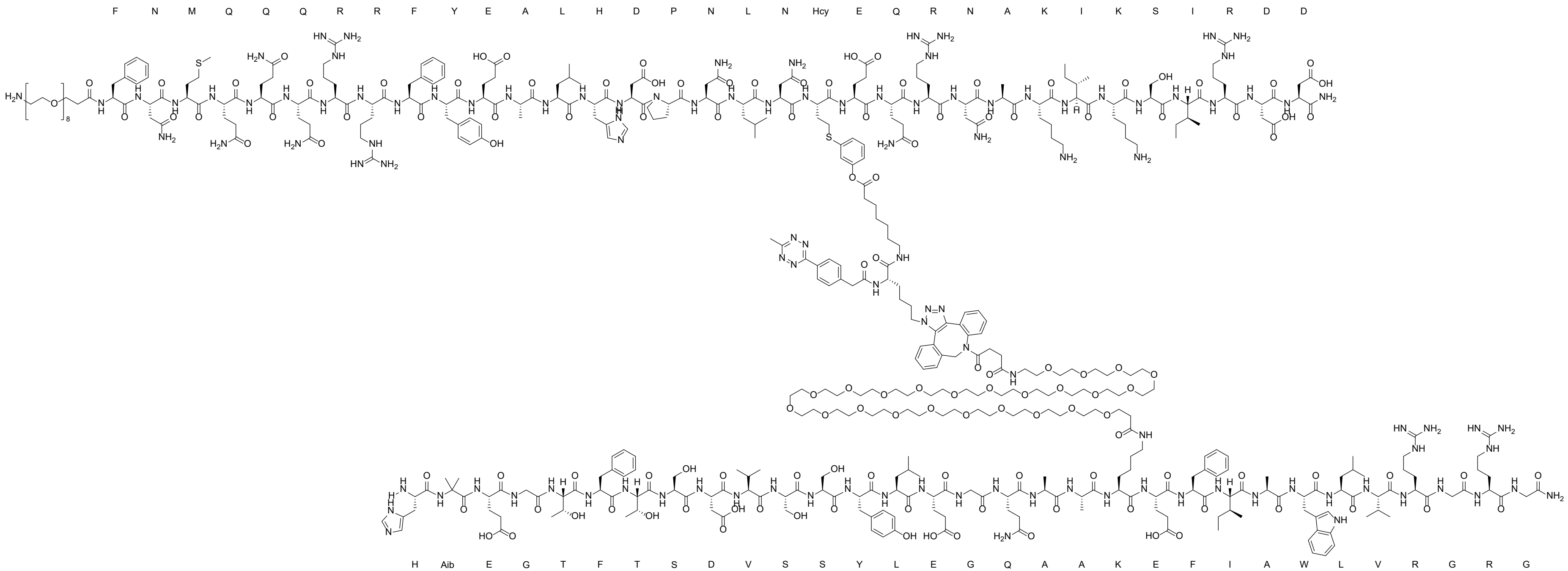

LC-MS analysis: Method B

Calculated Mass: 9914.2 Da

Observed Mass*: 9914.7 Da (*The peak observed around 5 min elution time was also detected in the blank measurement, suggesting its origin as an unidentified compound retained in the column.)

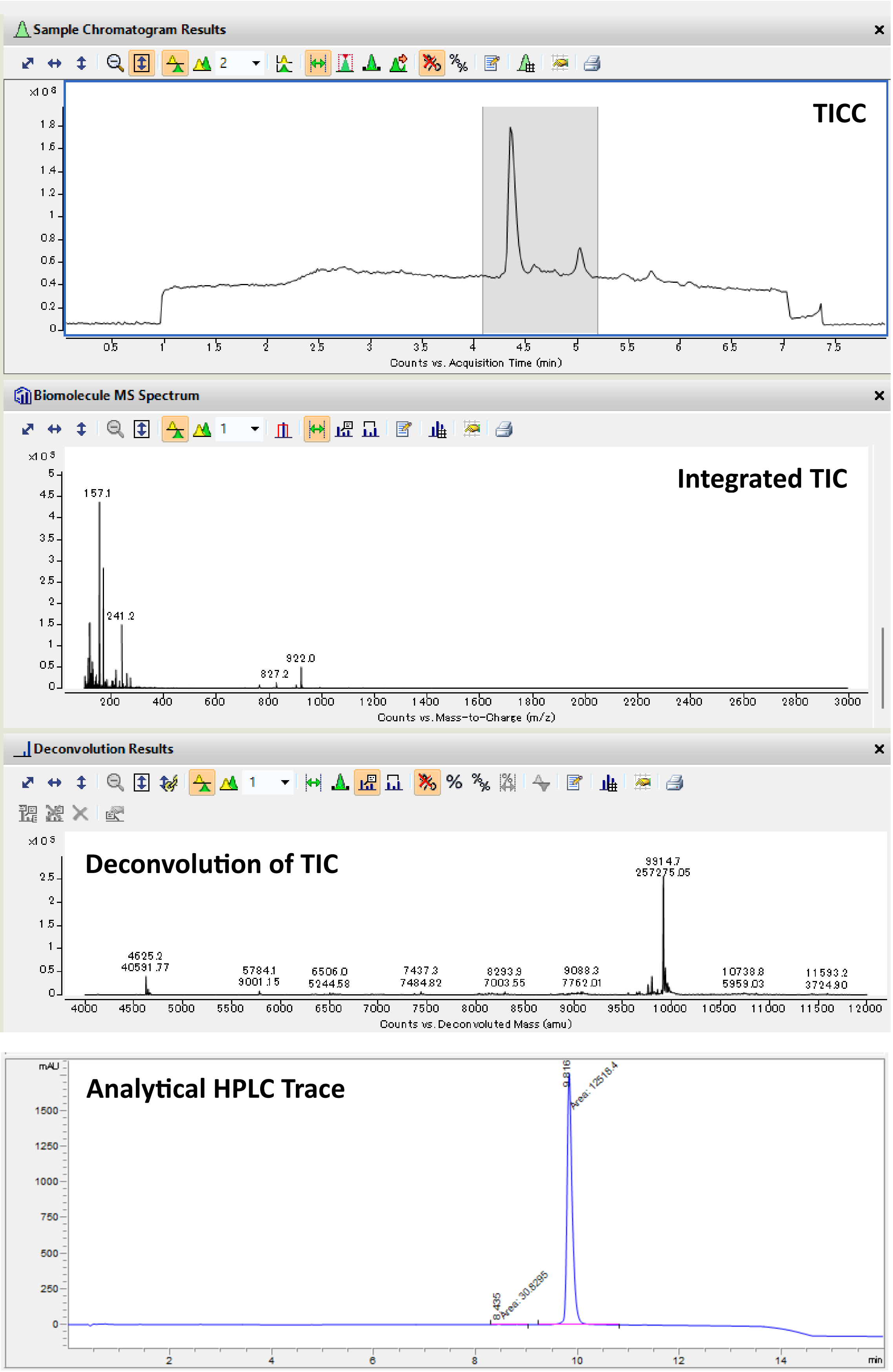

TICC: Total Ion Current Chromatogram; TIC: Total Ion Current.

Peptide: **GLP-1a**

Structure:

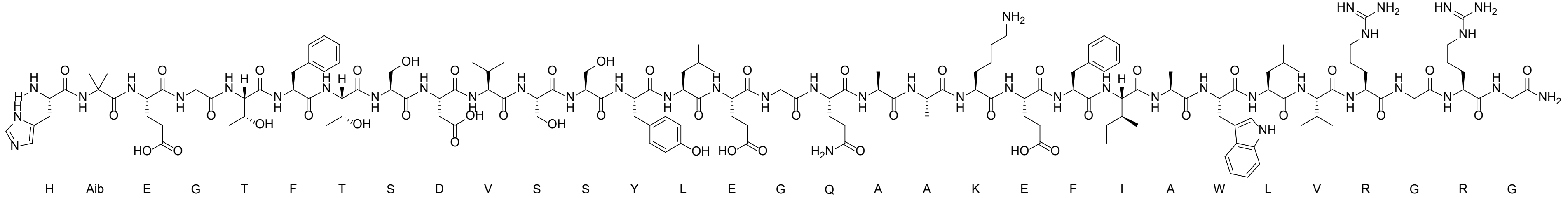

LC-MS analysis: Method D

Calculated Exact Mass: 3396.8 Da

Observed Mass: 3397.3 Da

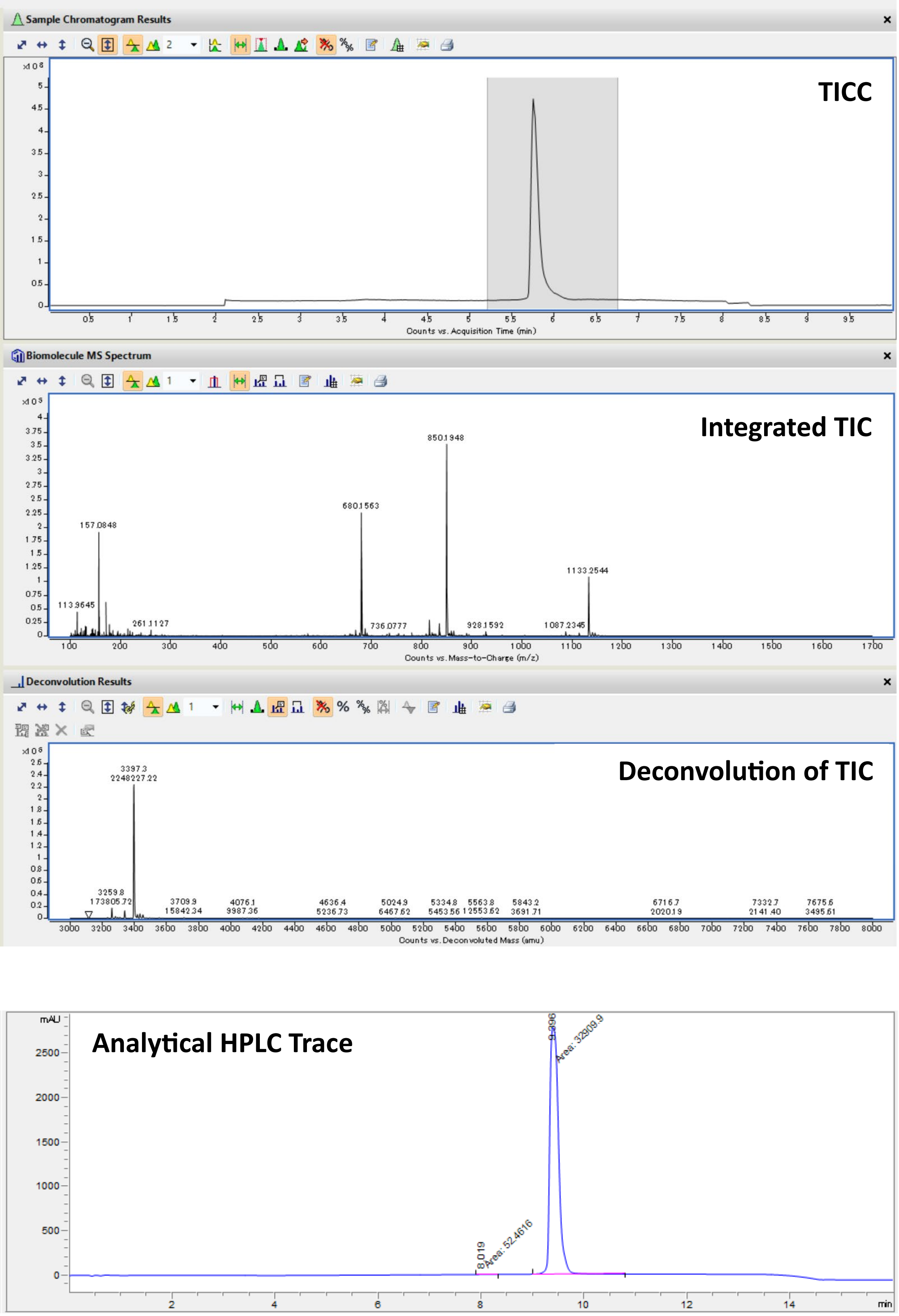

TICC: Total Ion Current Chromatogram; TIC: Total Ion Current.

Peptide: **2a**

Structure:

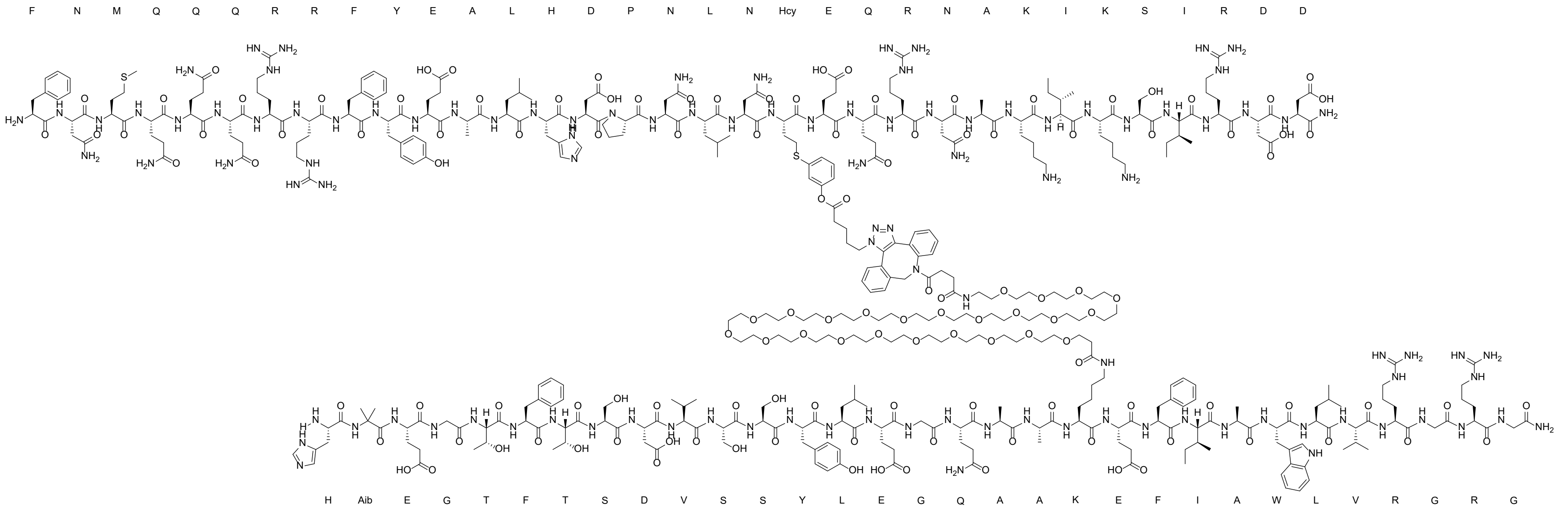

LC-MS analysis: Method B

Calculated Mass: 9122.3 Da

Observed Mass: 9122.7 Da

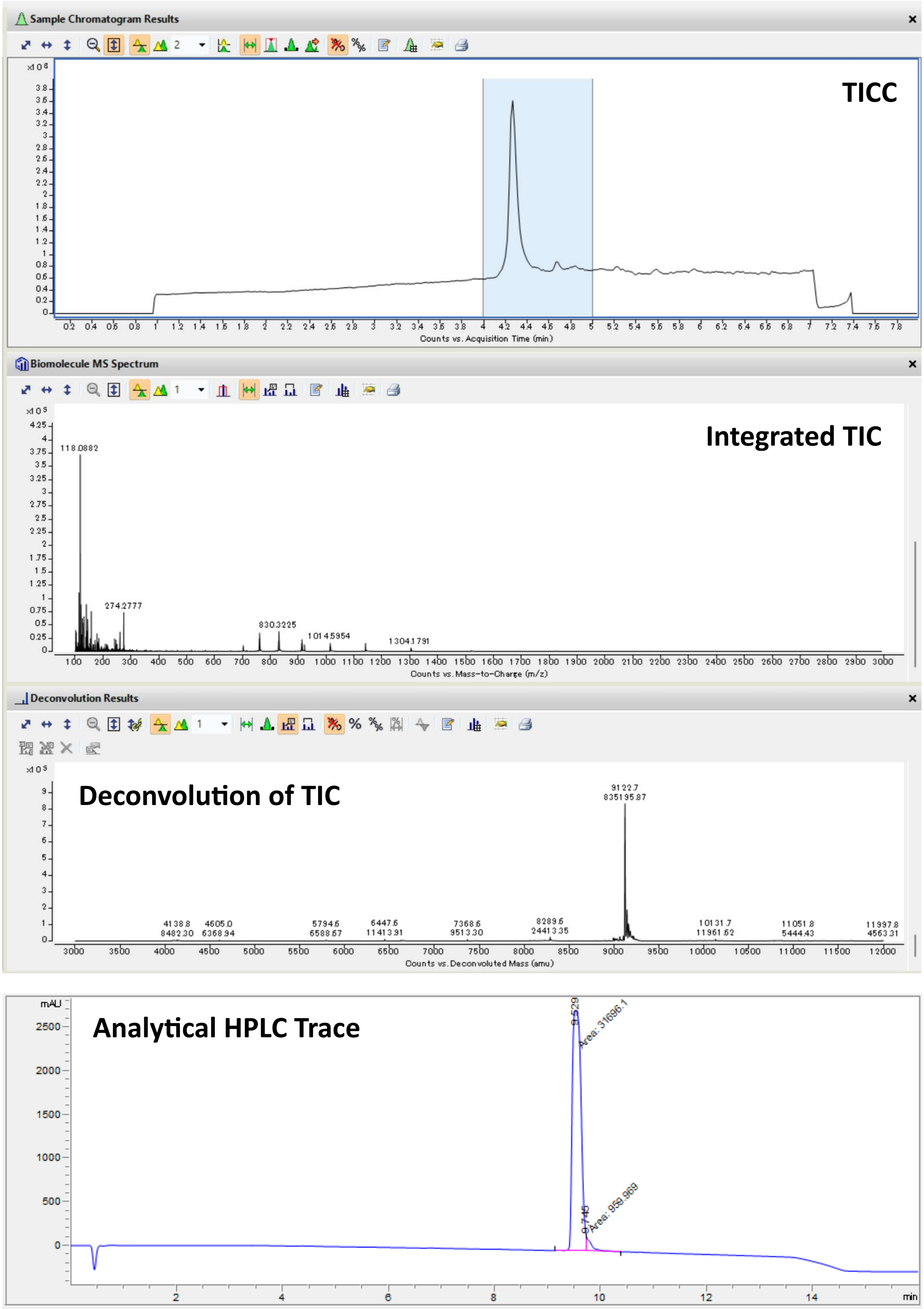

TICC: Total Ion Current Chromatogram; TIC: Total Ion Current.

Peptide: **3a**

Structure:

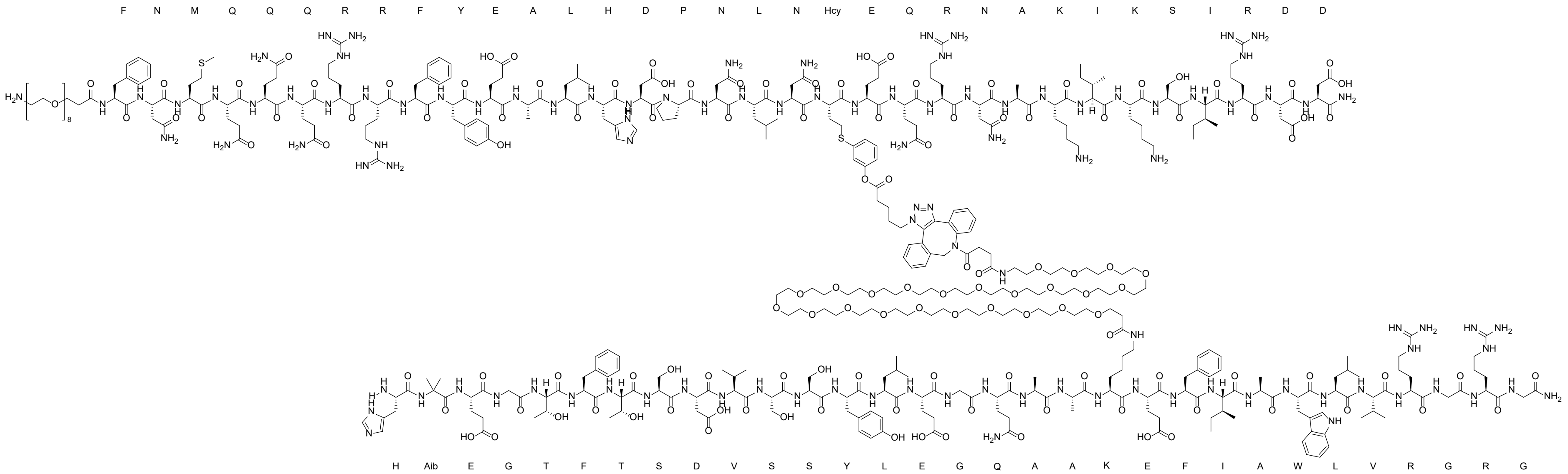

LC-MS analysis: Method D

Calculated Mass: 9545.8 Da

Observed Mass: 9545.8 Da

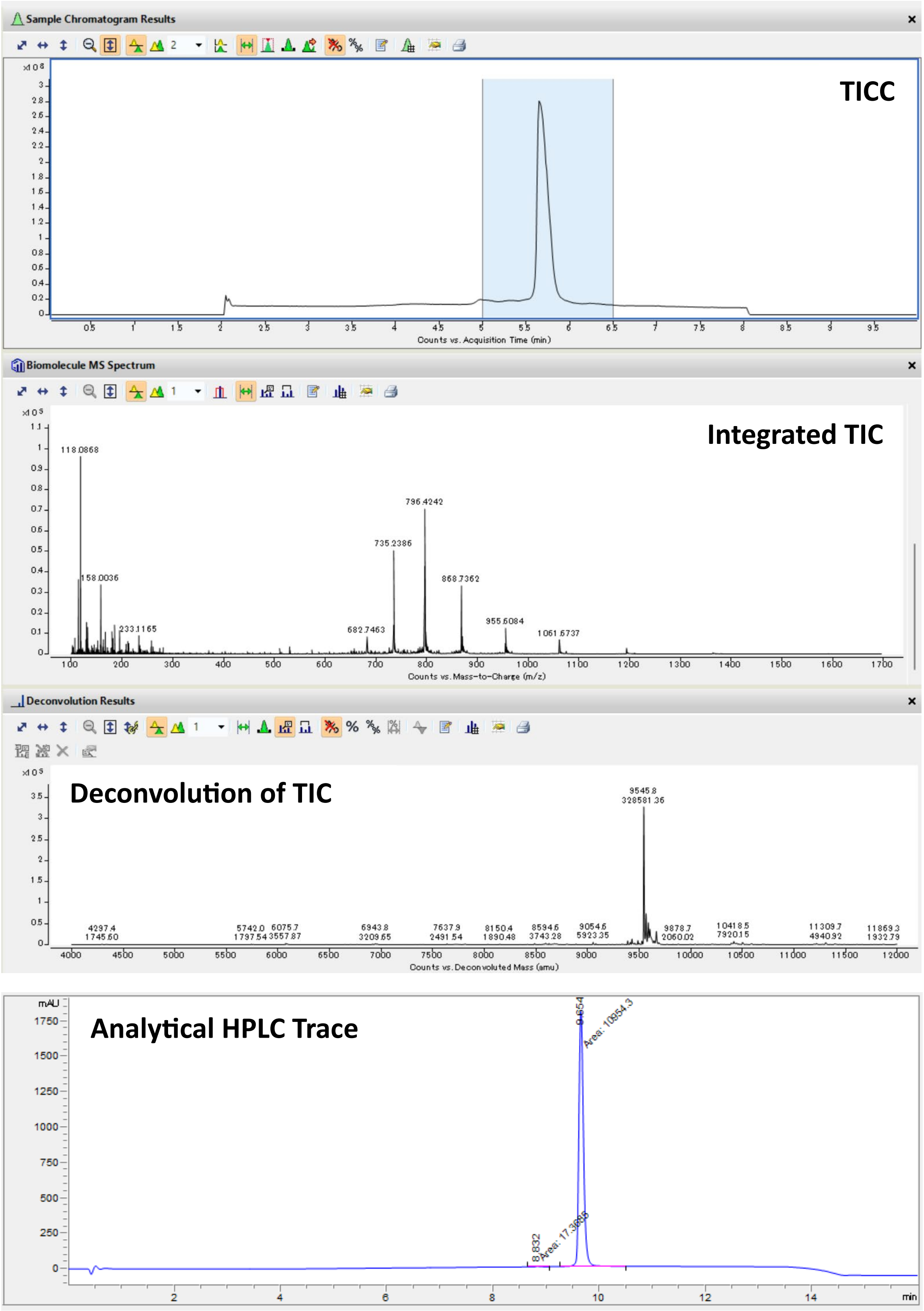

TICC: Total Ion Current Chromatogram; TIC: Total Ion Current.

Peptide: **$N_3$-GLP-1a**

Structure:

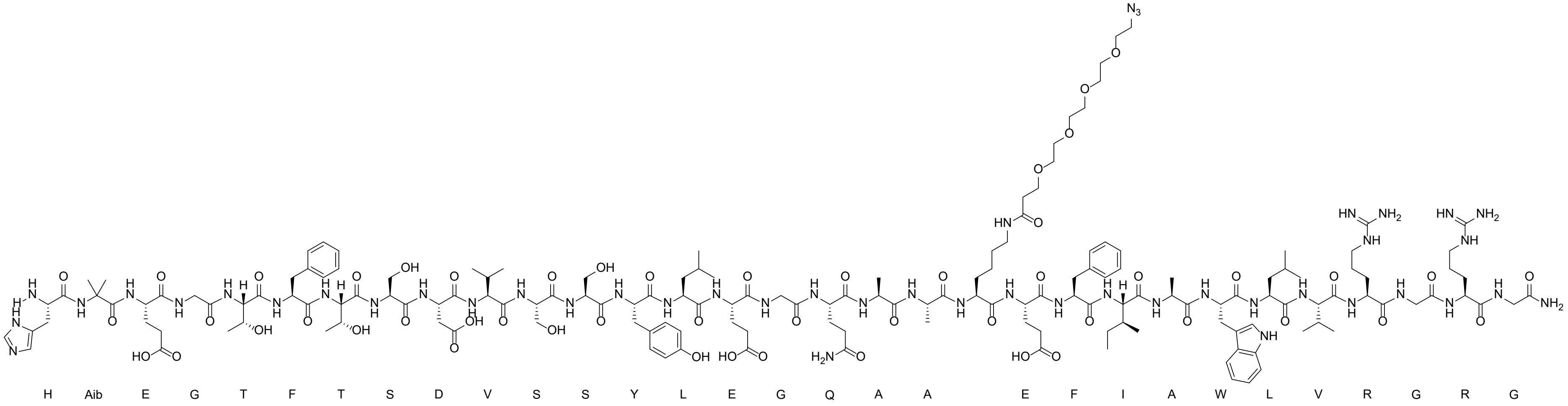

LC-MS analysis: Method B

Calculated Exact Mass: 3670.1 Da

Observed Mass*: 3670.4 Da (*The peak observed around 5 min elution time was also detected in the blank measurement, suggestiong its origin as an unidentified compound retained in the column.)

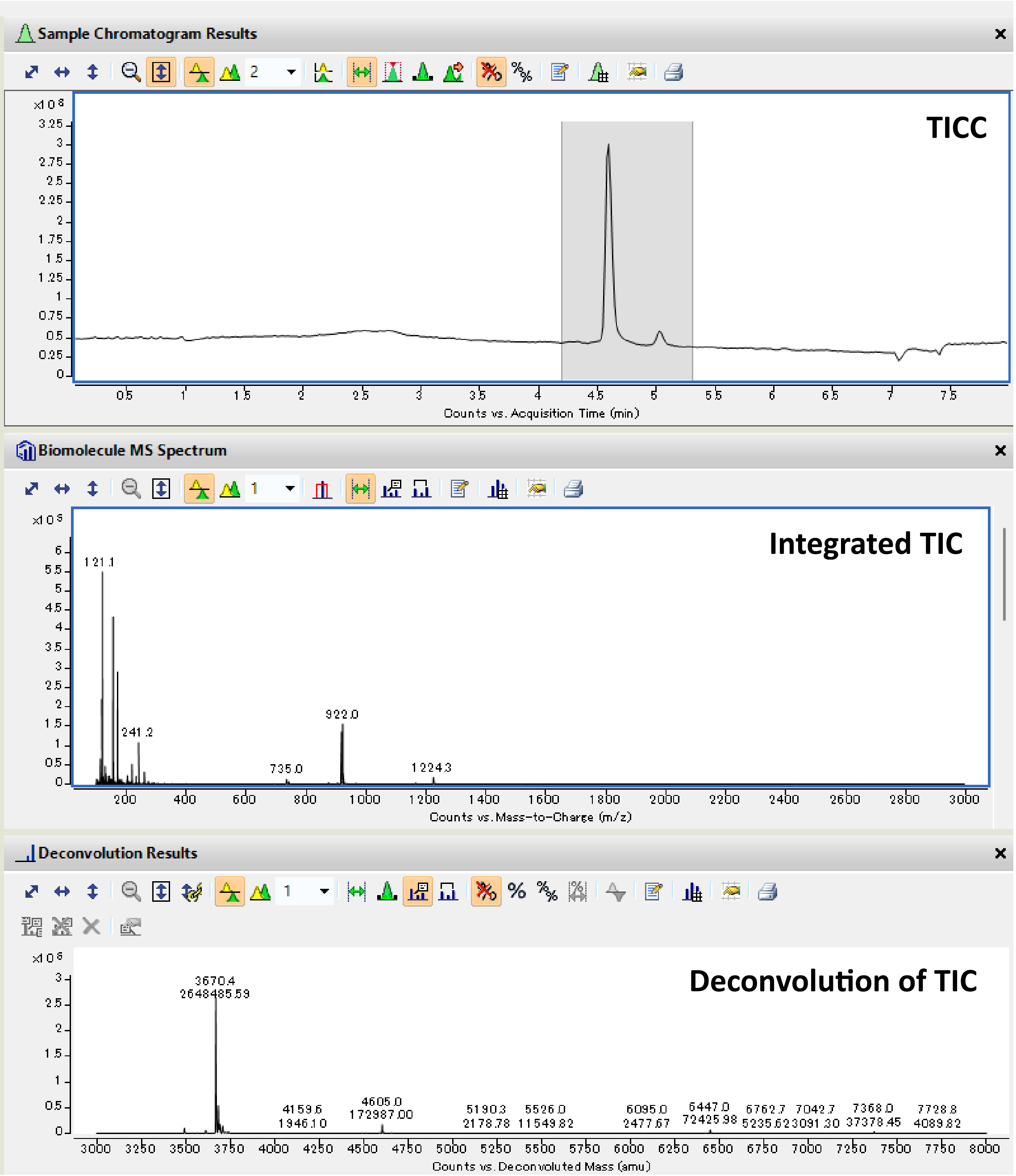

TICC: Total Ion Current Chromatogram; TIC: Total Ion Current.

Peptide: **DBCO-PEG24-GLP-1a**

Structure:

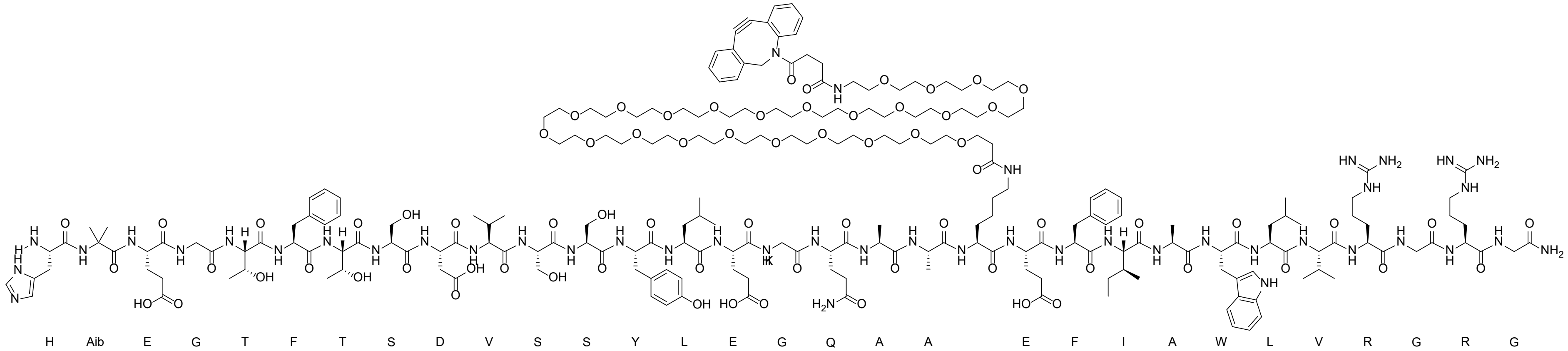

LC-MS analysis: Method B

Calculated Mass: 4812.4 Da

Observed Mass: 4812.5 Da

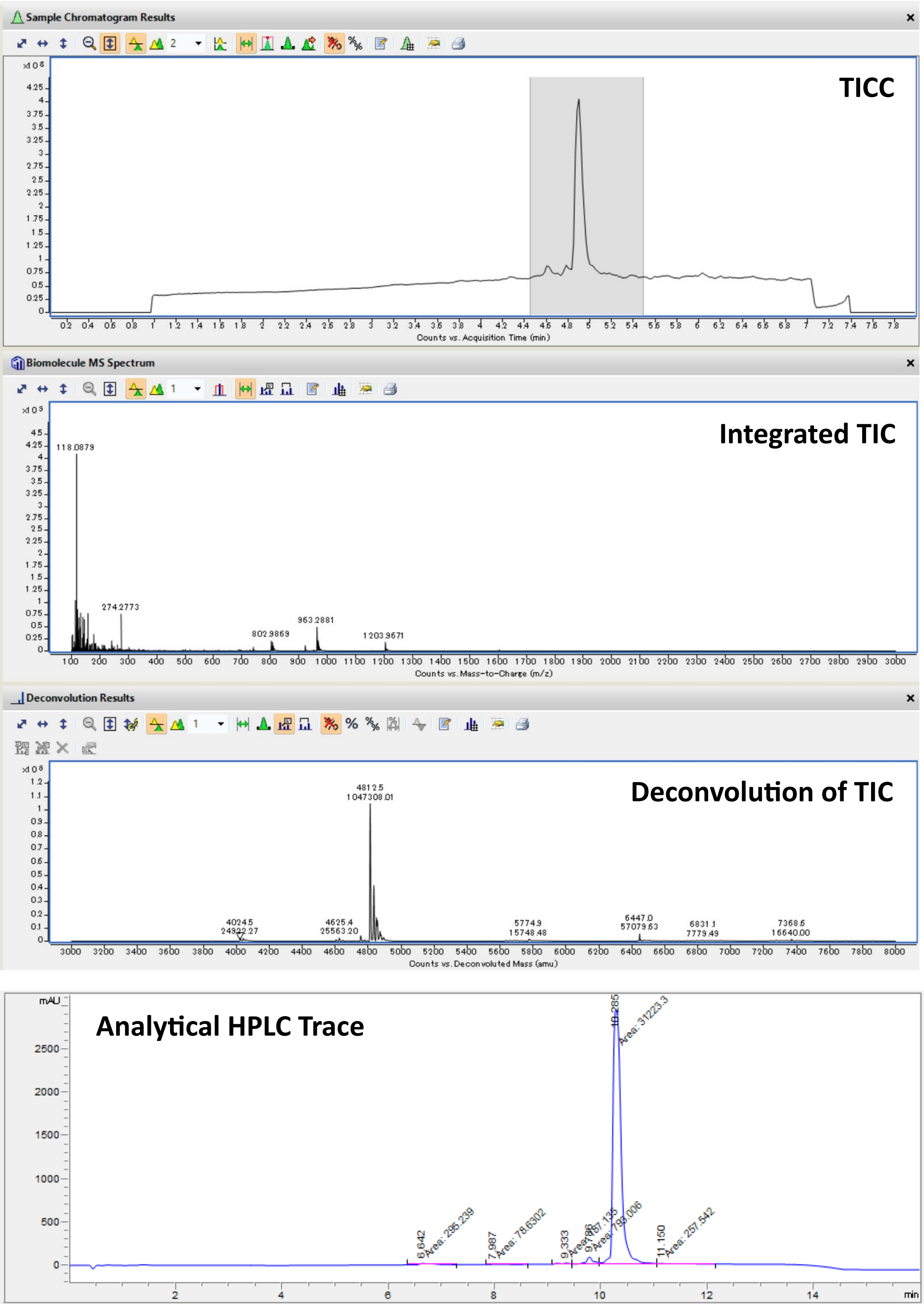

TICC: Total Ion Current Chromatogram; TIC: Total Ion Current.

Peptide: **GLP-1b**

Structure:

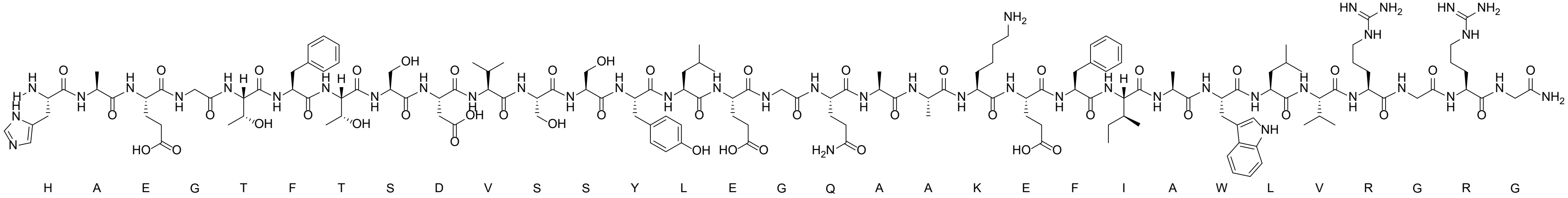

LC-MS analysis: Method D

Calculated Mass: 3382.7 Da

Observed Mass: 3383.3 Da

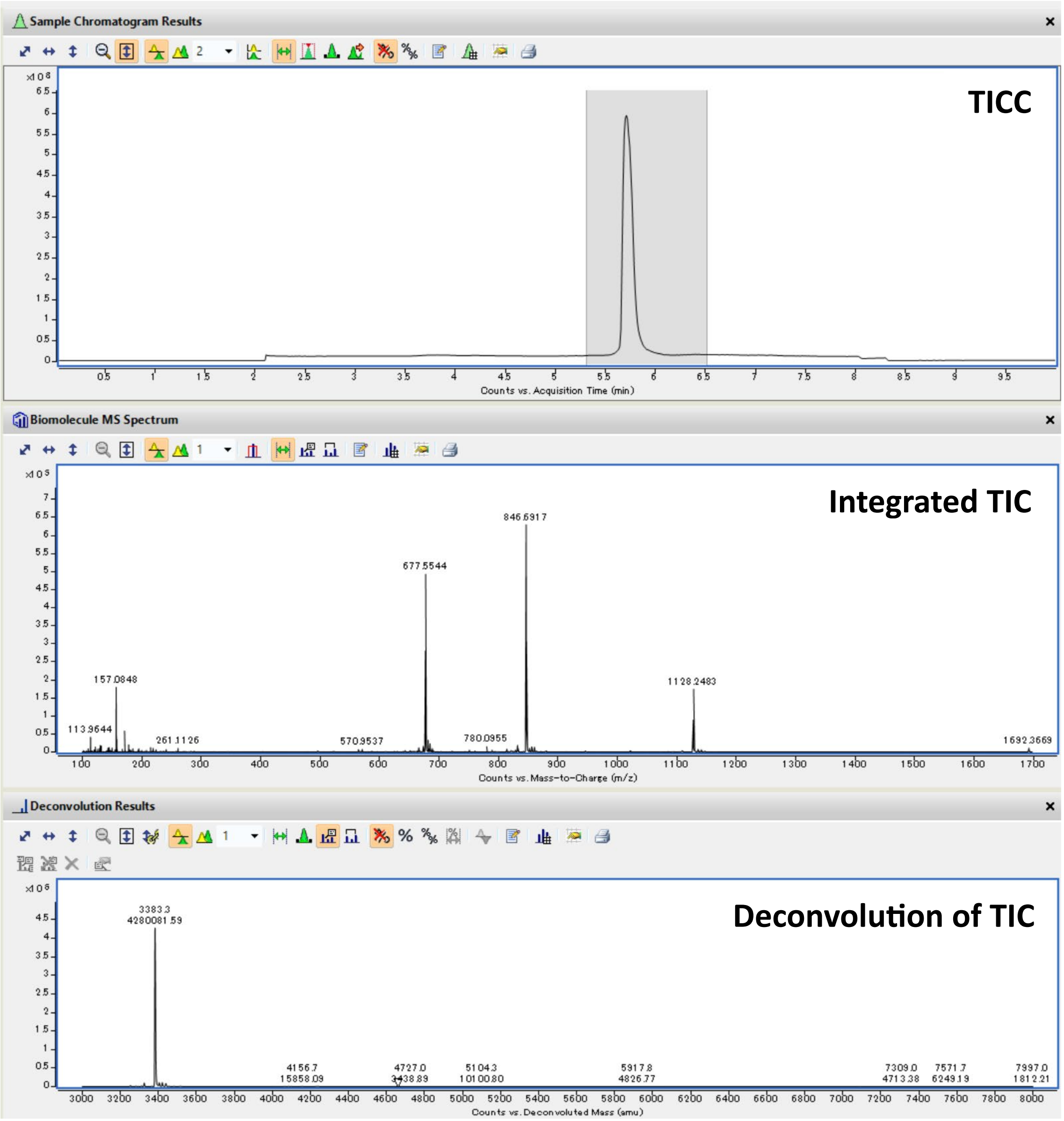

TICC: Total Ion Current Chromatogram; TIC: Total Ion Current.

Peptide: **GLP-1c**

Structure:

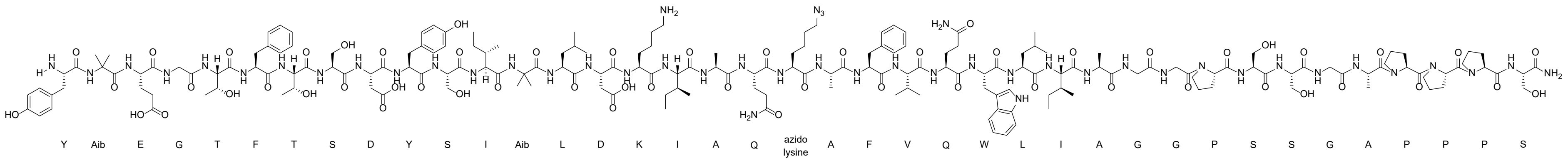

LC-MS analysis: Method B

Calculated Mass: 4095.6 Da

Observed Mass: 4095.7 Da

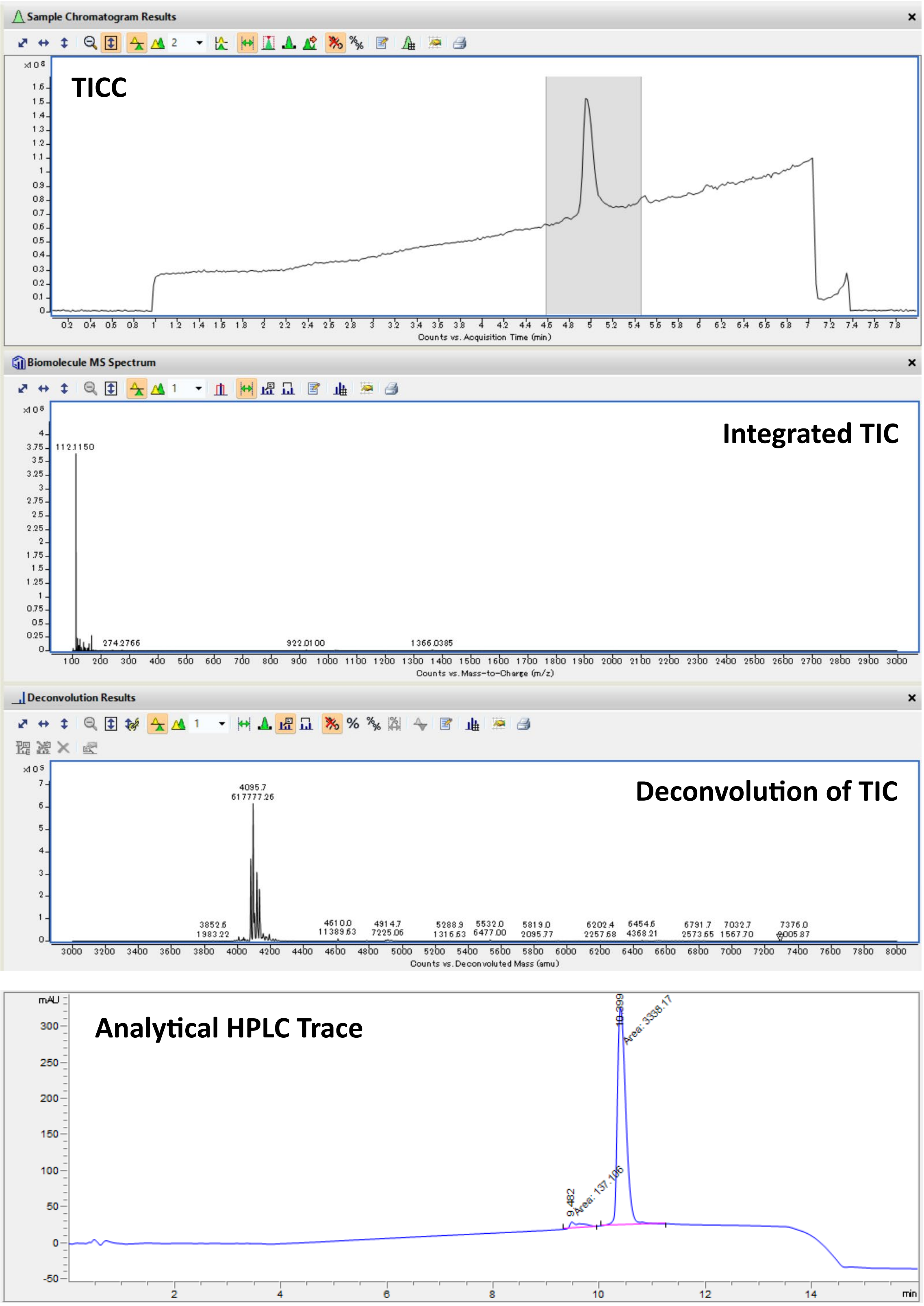

TICC: Total Ion Current Chromatogram; TIC: Total Ion Current.

Peptide: **Z33-GLP-1c**

Structure:

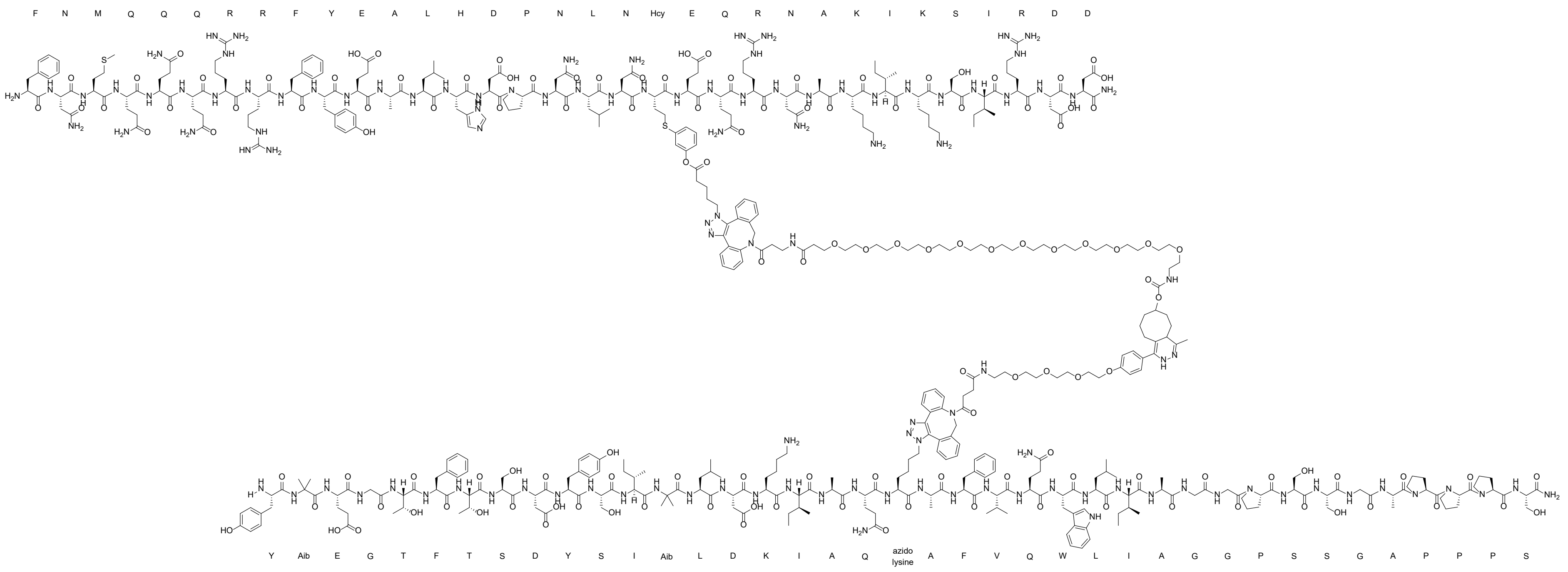

LC-MS analysis: Method B

Calculated Mass: 10056 Da

Observed Mass: 10057 Da

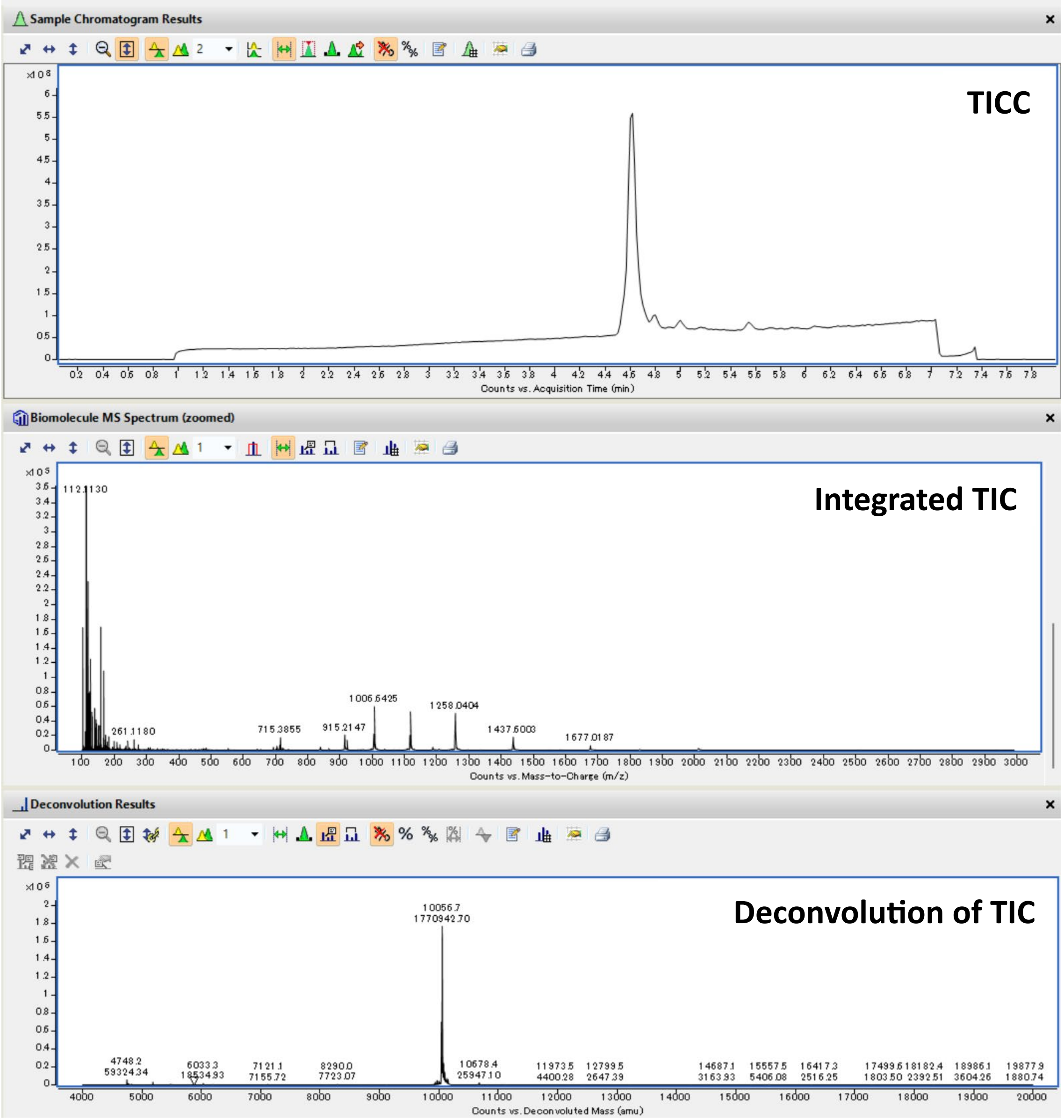

TICC: Total Ion Current Chromatogram; TIC: Total Ion Current.

**Supplementary NMR Spectral Data**

Compound: **BB-7**

$^{1}H$:

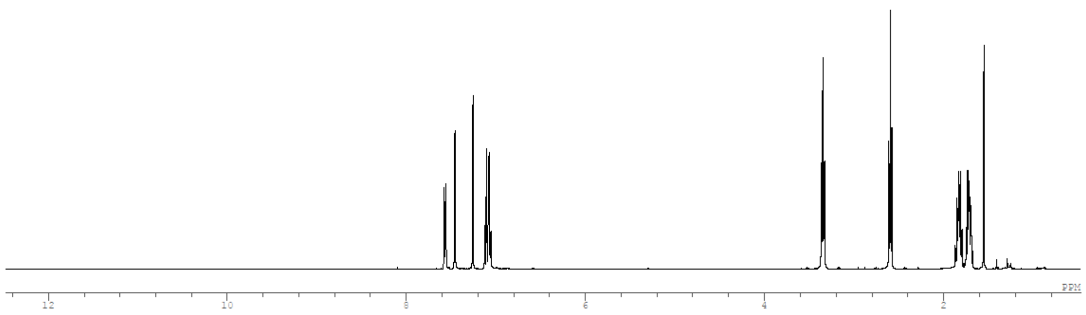

$^{13}C$:

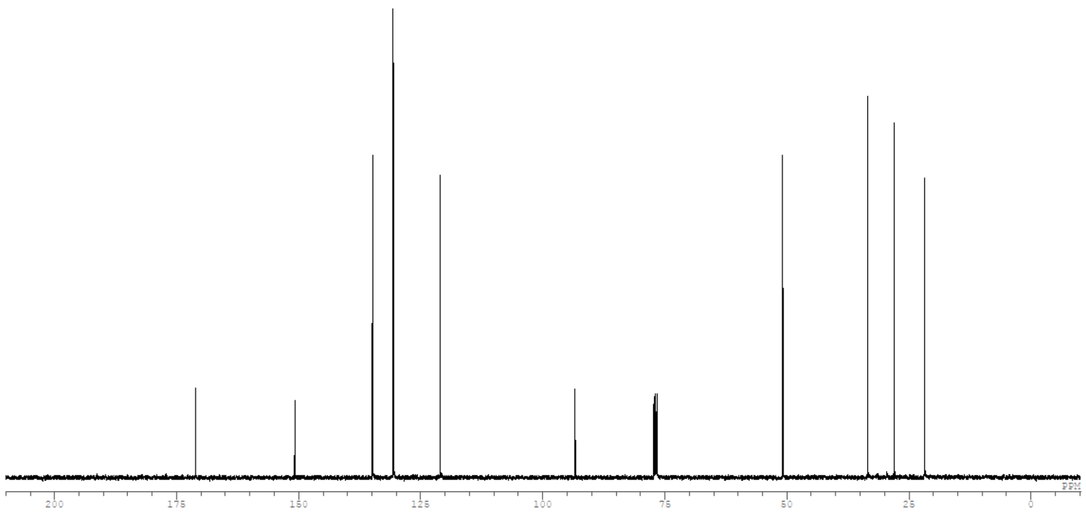

Compound: **BB-8**

$^{1}H$:

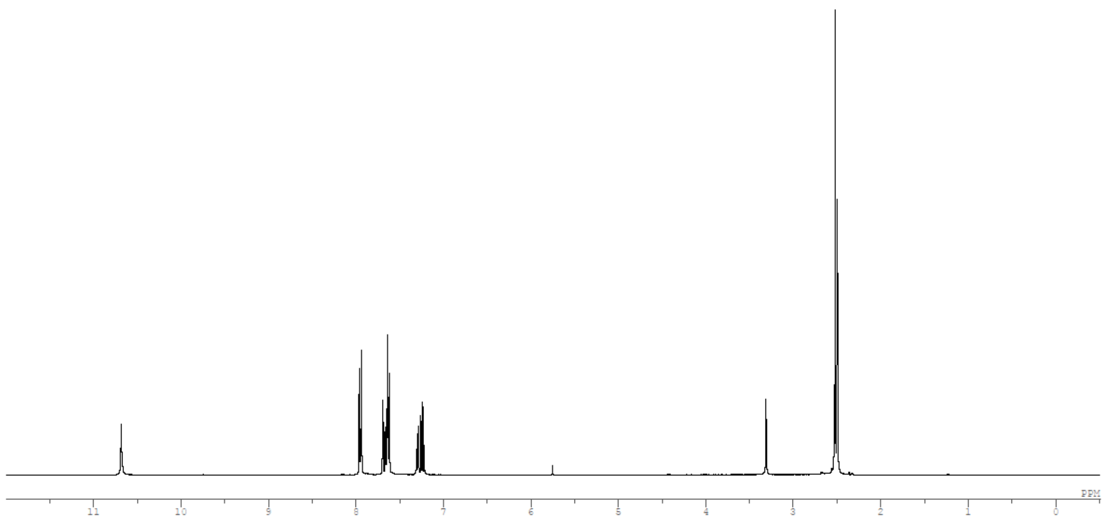

Compound: **BB-9**

$^{1}H$:

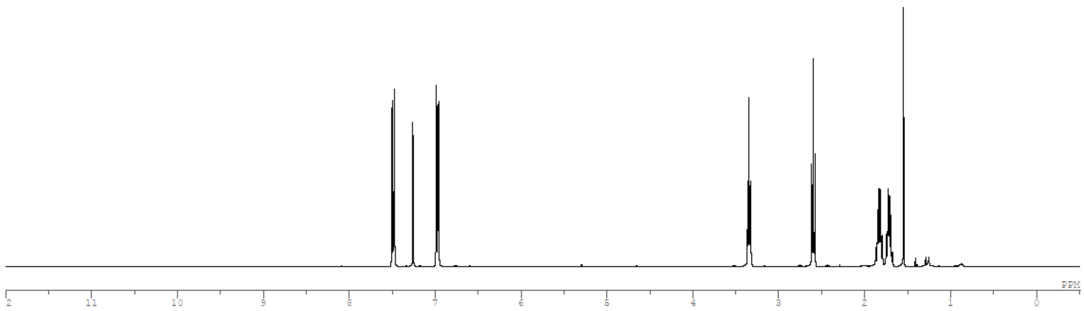

$^{13}C$:

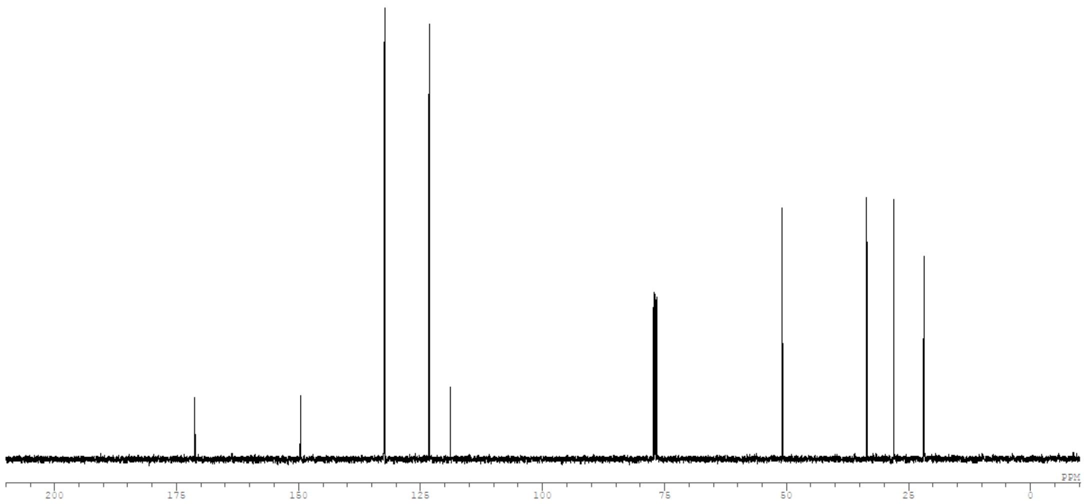

Compound: **BB-10**

$^{1}H$:

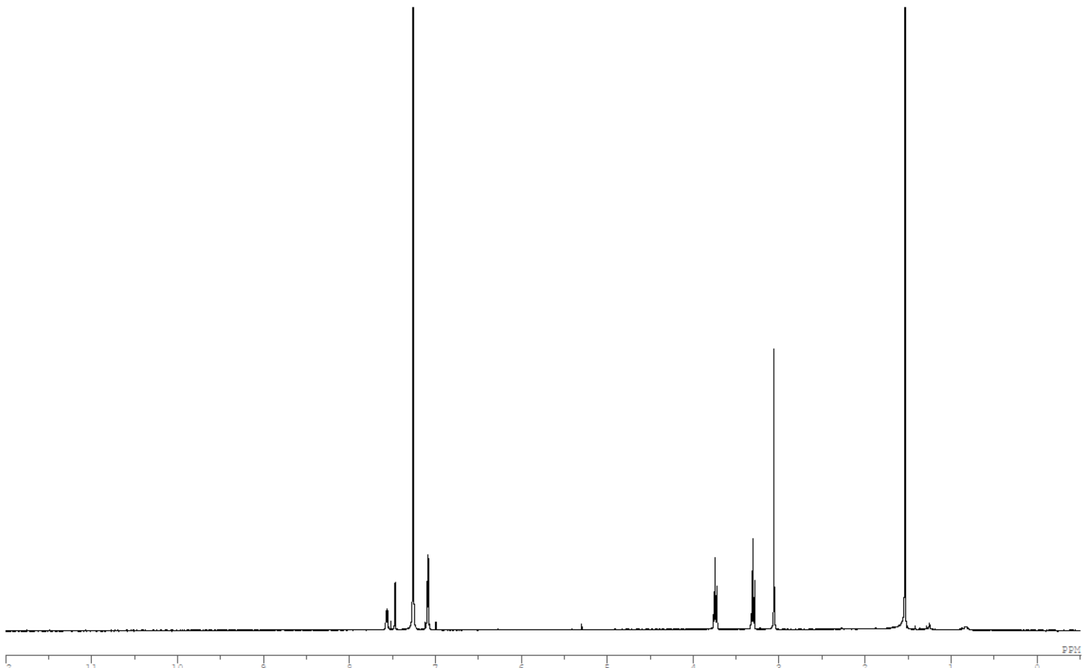

Compound: **BB-11 intermediate**

$^{1}H$:

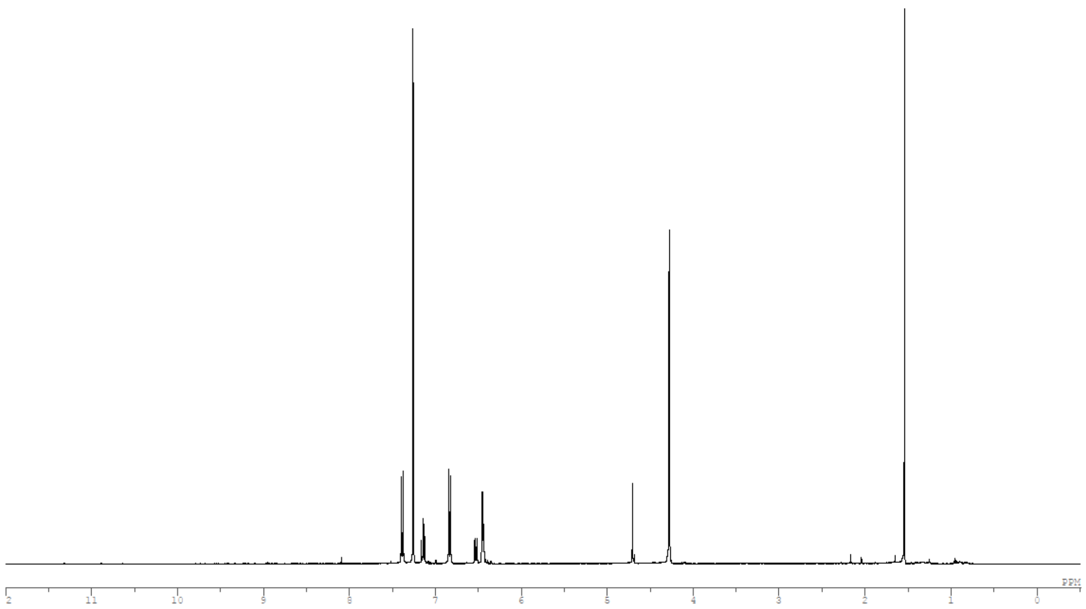

$^{13}C$:

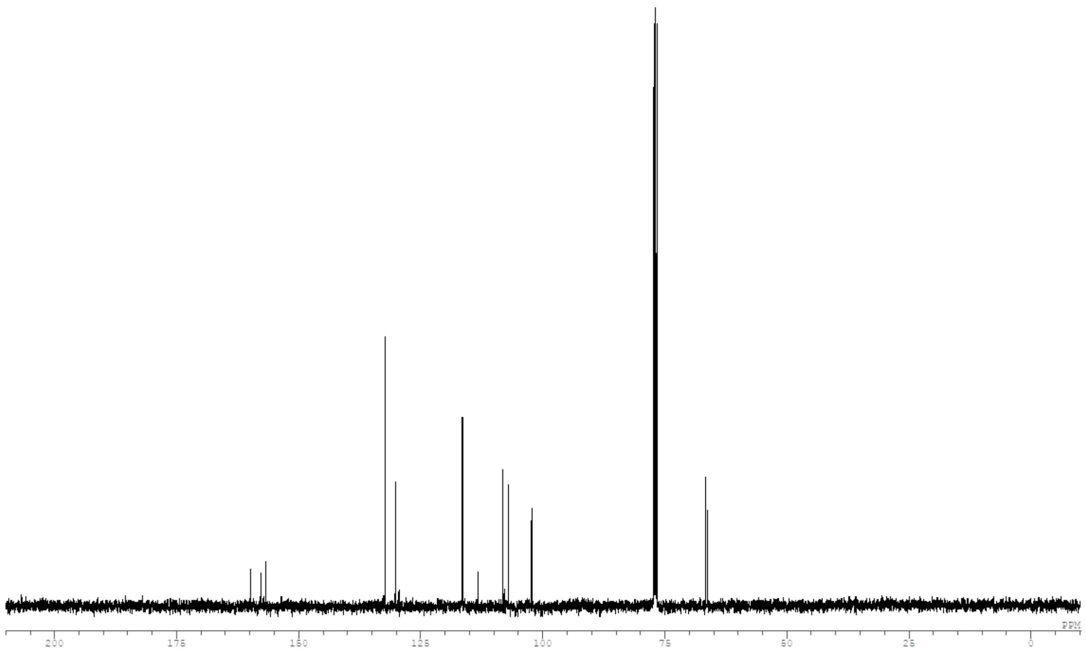

Compound: **BB-11**

$^{1}H$:

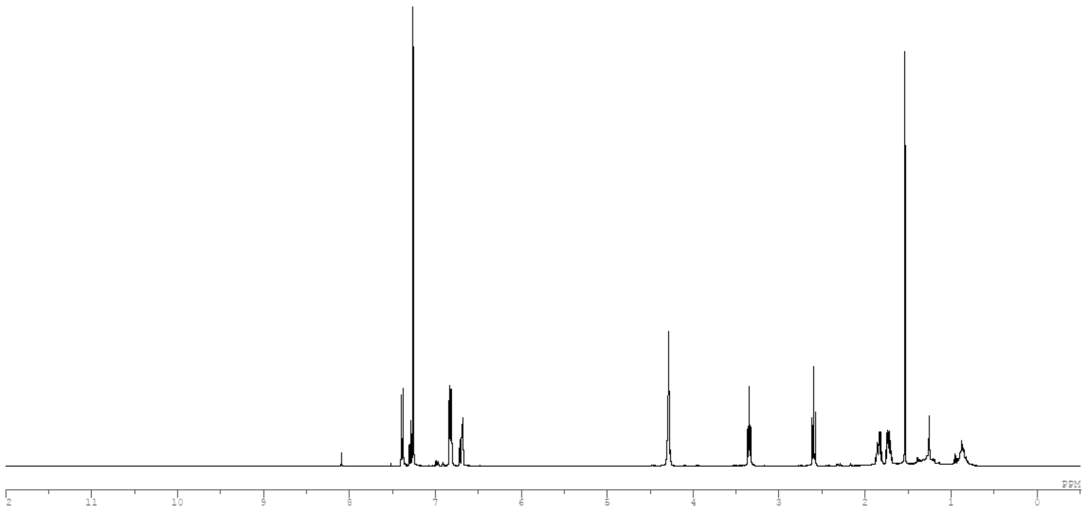

$^{13}C$:

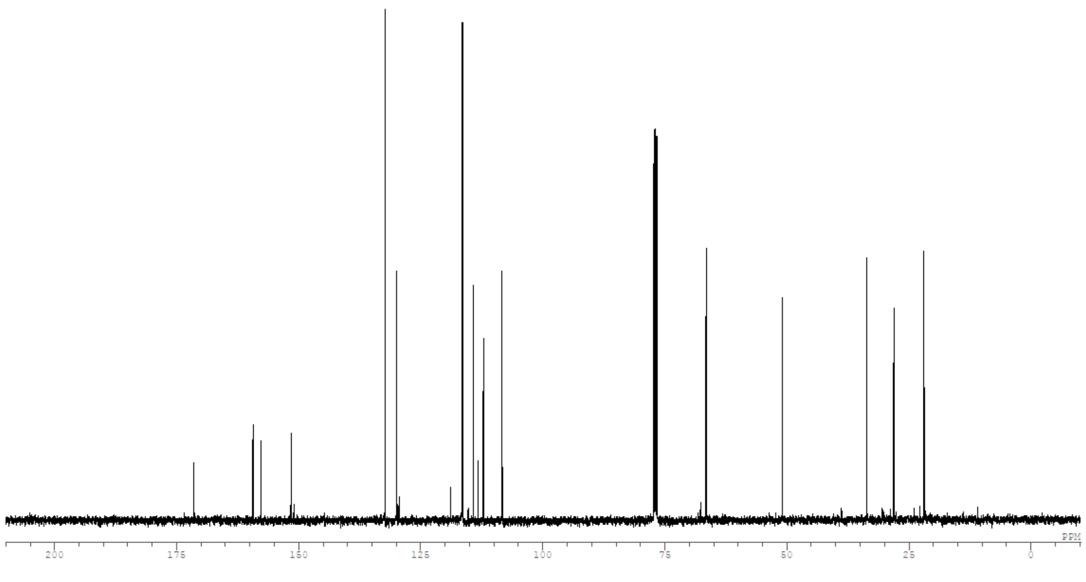

Compound: **BB-12 intermediate**

$^{1}H$:

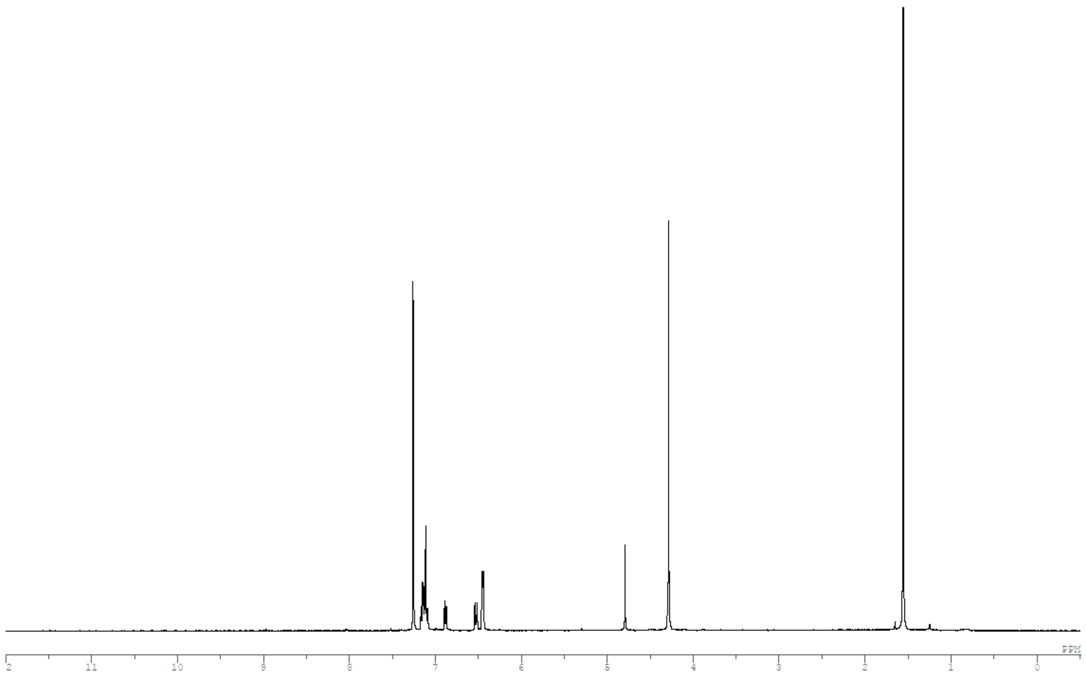

$^{13}C$:

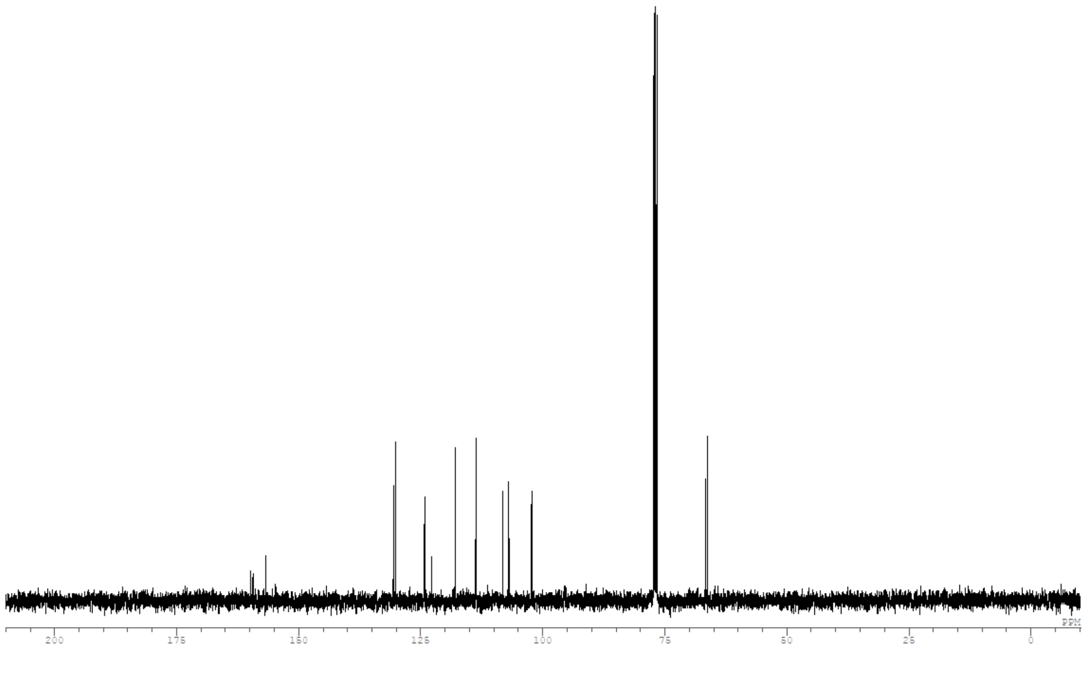

Compound: **BB-12**

$^{1}$H:

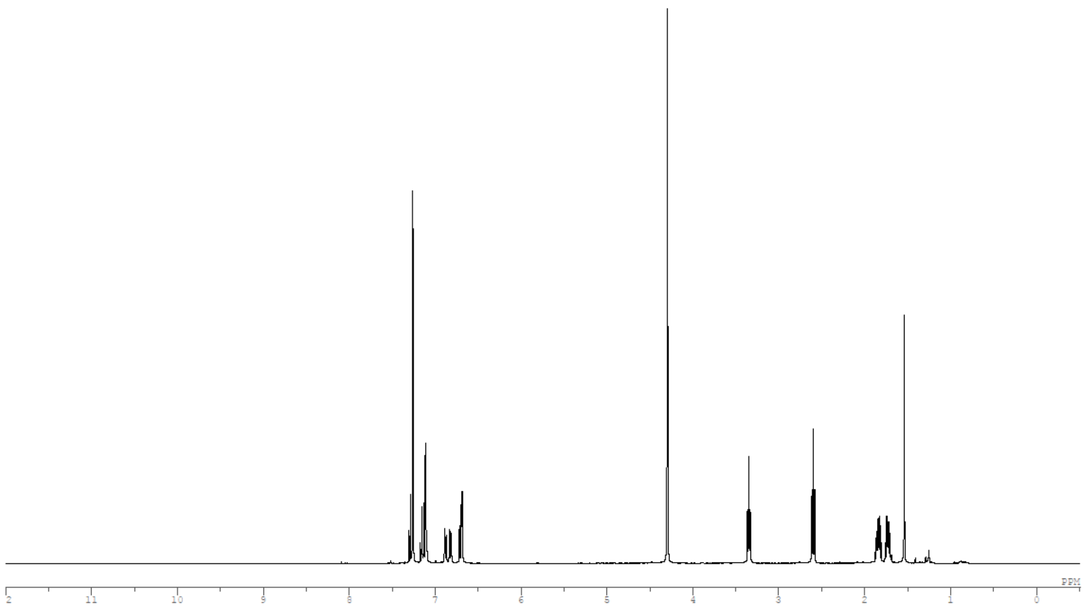

$^{13}$C:

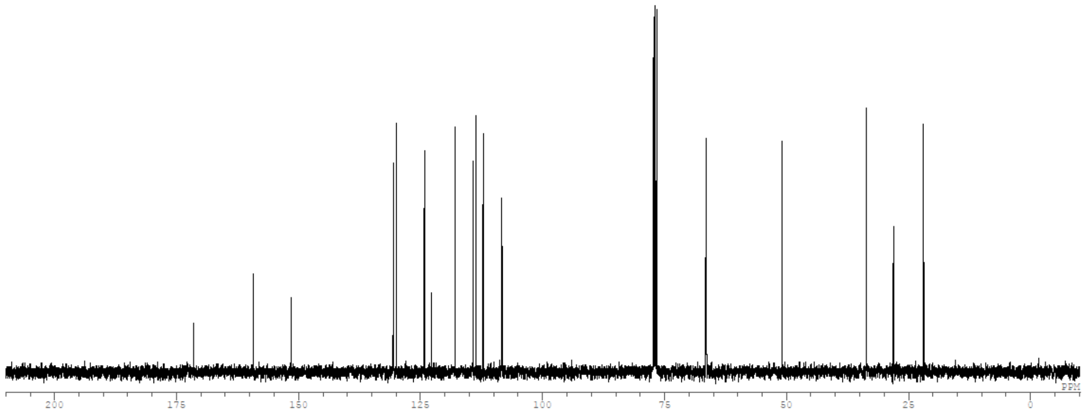

Compound: **BB-13 intermediate 1**

[1]H:

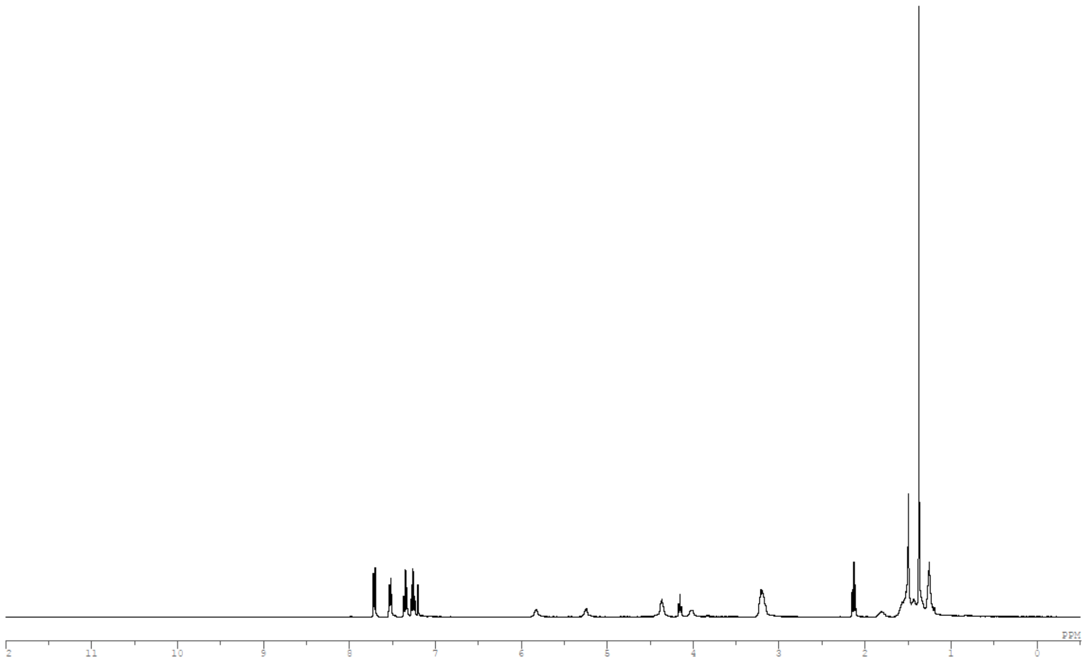

[13]C:

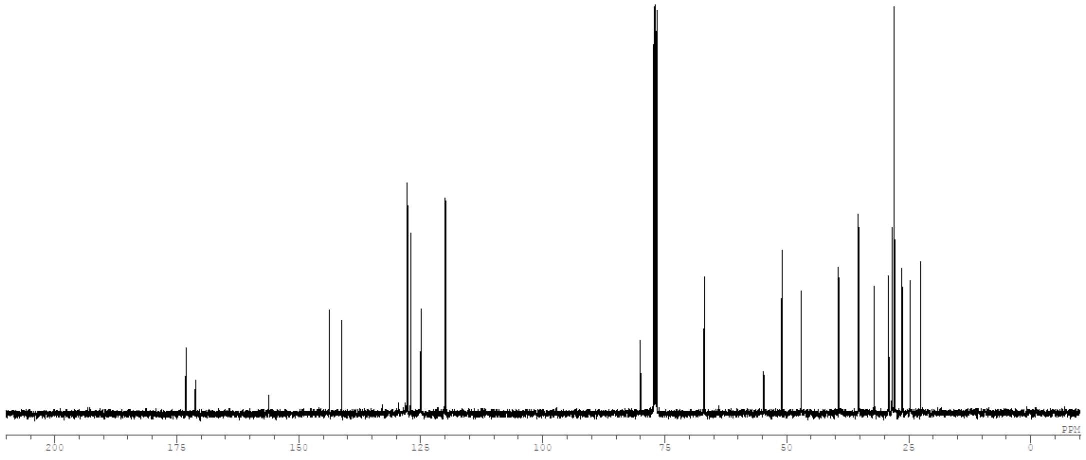

Compound: **BB-13 intermediate 2**

$^{1}H$:

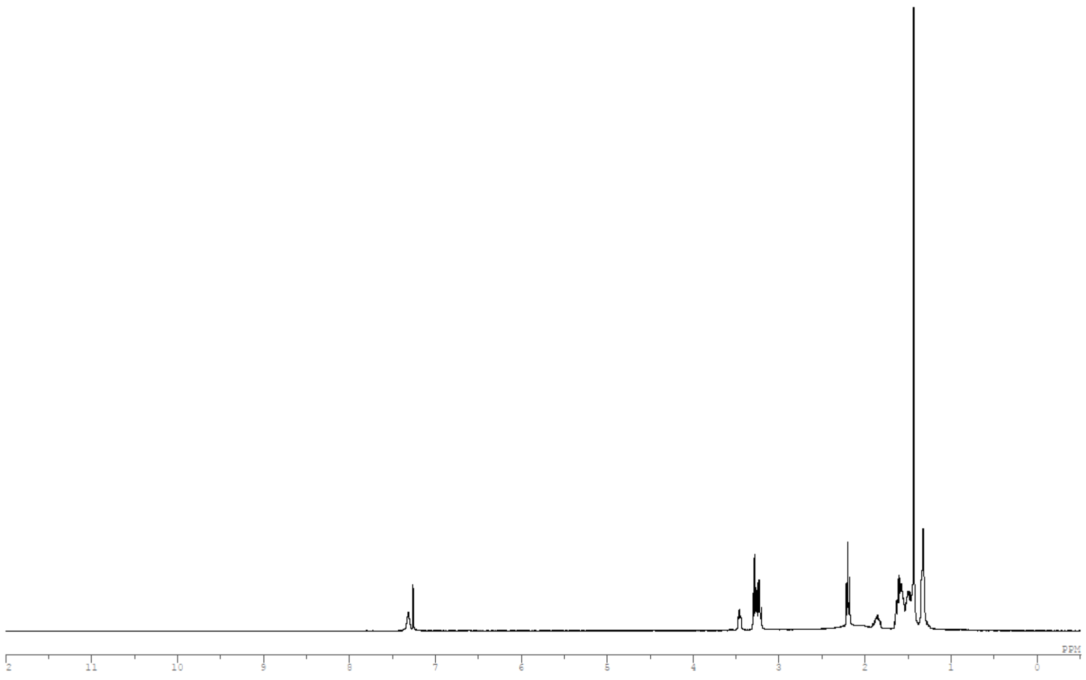

$^{13}C$:

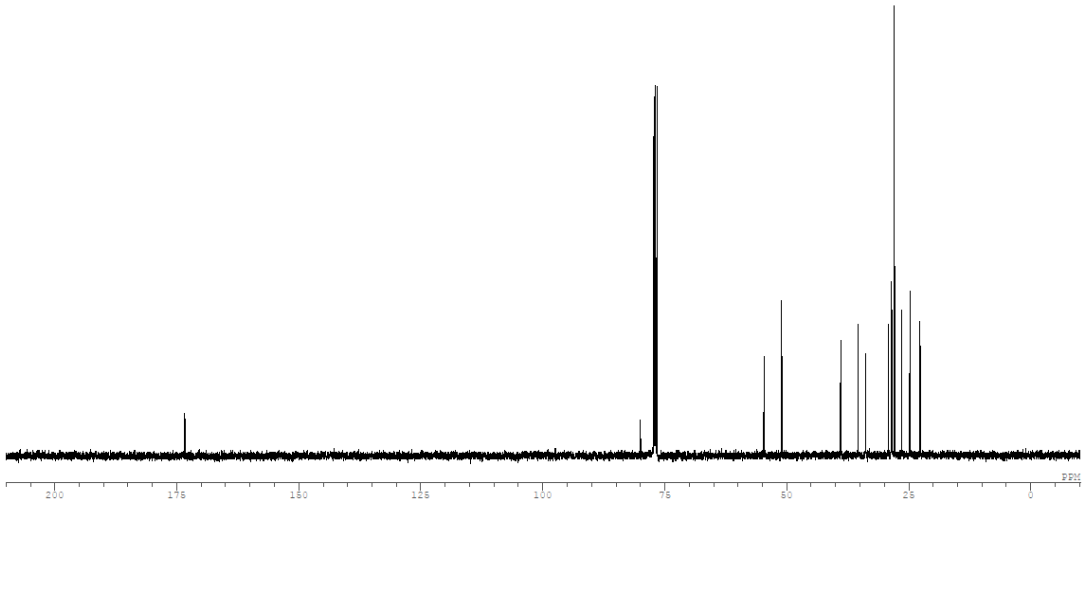

Compound: **BB-13 intermediate 3**

$^{1}H$:

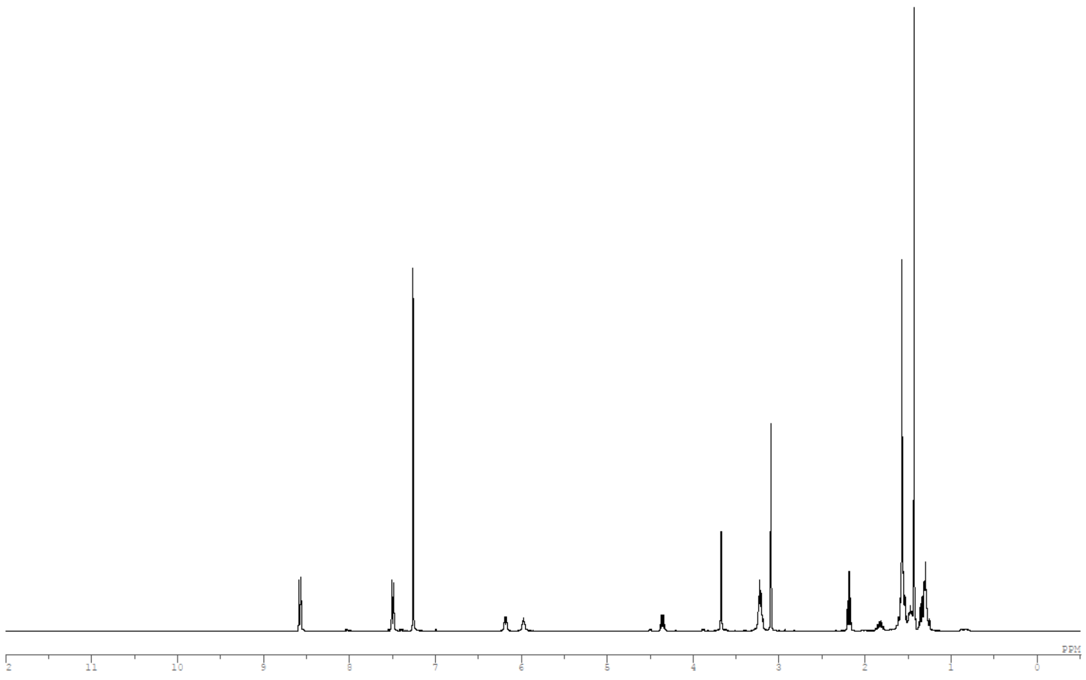

$^{13}C$:

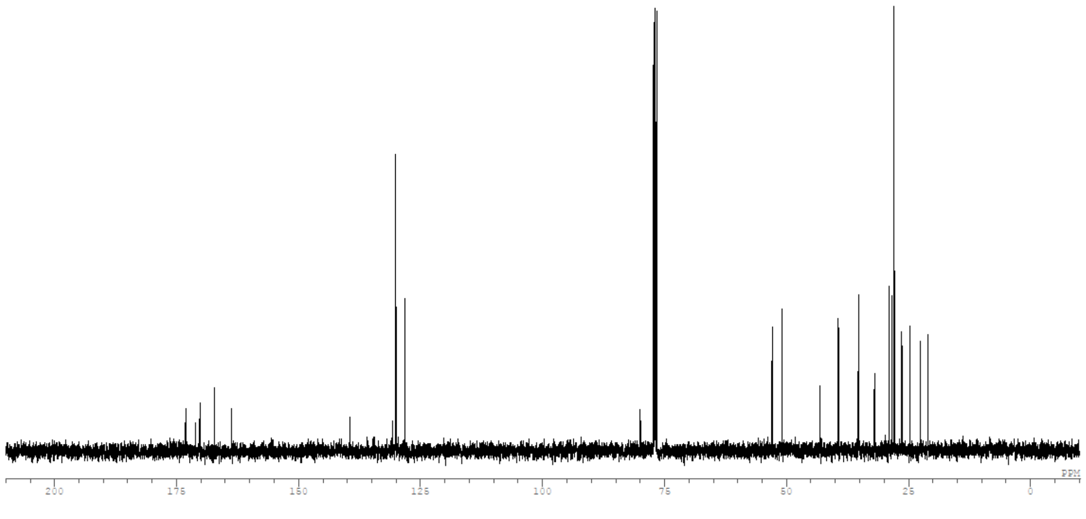

Compound: **BB-13**

$^{1}$H:

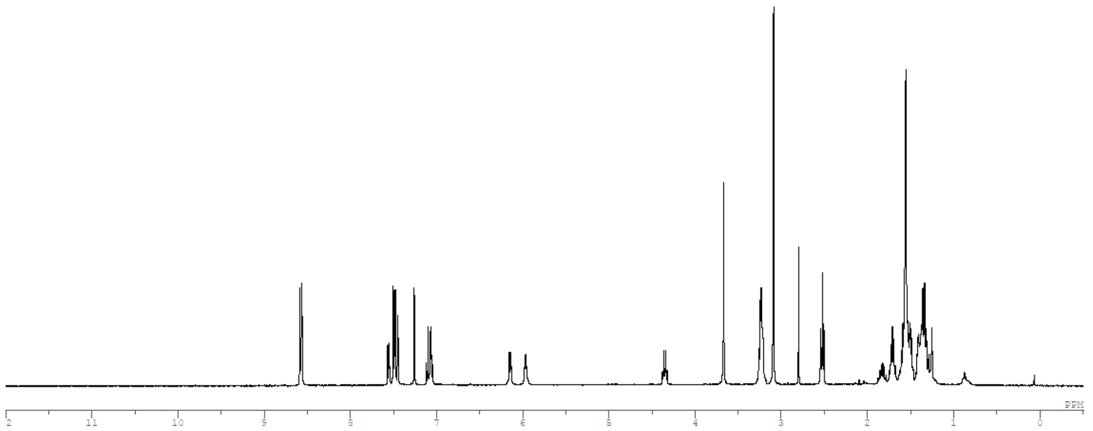

$^{13}$C:

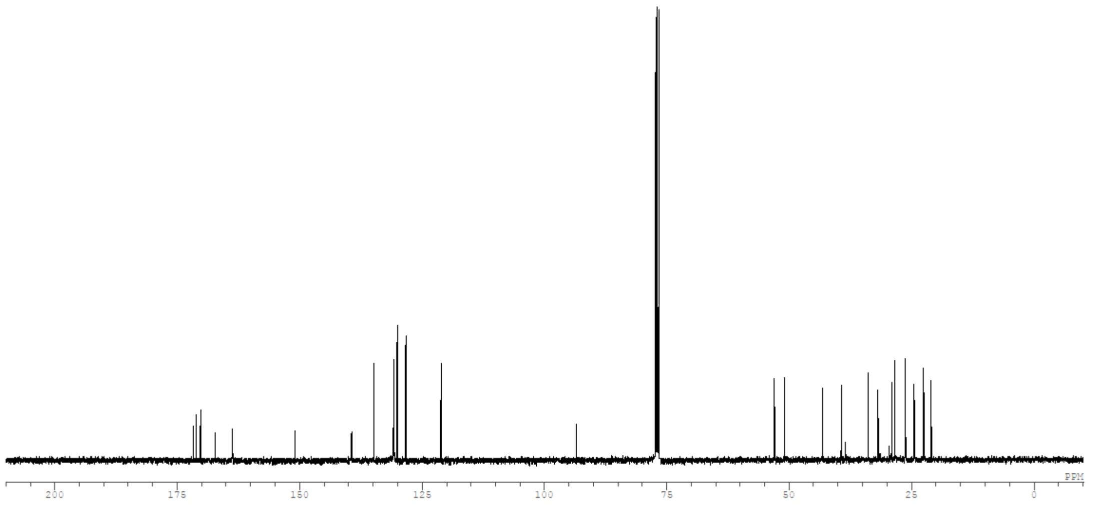

### Supplementary Tables:

**Table S1**. Sequences and mass of the synthesized Z33 variants.

| Code | Sequence[a] | Electrophile | Calculated Mass (Da) | Observed Mass (Da)[c] |
|---|---|---|---|---|
| $N_3$-**1-I** | FNMQQQRRFYEALHD<br>pNLN**C**EQRNAKIKSI<br>RDD | 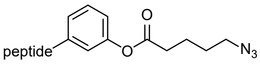 | 4295.8 | 4296.1 |
| $N_3$-**2-I** | FNMQQQRRFYEALHD<br>PNLN**Hcy**EQRNAKIKSI<br>RDD | 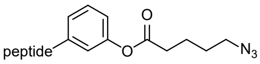 | 4309.8 | 4310.0 |
| $N_3$-**3-I** | $H_2N$-PEG8-<br>FNMQQQRRFYEALHD<br>PNLN**Hcy**EQRNAKIKSI<br>RDD | 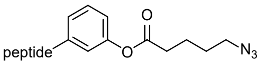 | 4733.3 | 4733.3 |
| $N_3$-**4-I** | FNMQQQRRFYEALHD<br>PNLN**Hcy**EQRNAKIKSI<br>RDd | 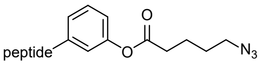 | 4309.8 | 4309.7 |
| $N_3$-**5-I** | fNMQQQRRFYEALHD<br>PNLN**Hcy**EQRNAKIKSI<br>RDd | 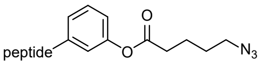 | 4309.8 | 4309.7 |
| $N_3$-**6-I** | fNMQQQRRFYEALHD<br>PNLN**Hcy**EQRNAKIKSI<br>Rdd | 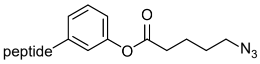 | 4309.8 | 4309.7 |
| $N_3$-**7-VI** | FN**C**QQQRRFYEALHD<br>PNLNEEQRNAKIKSI<br>RDD | 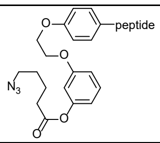 | 4429.9 | 4430.1 |
| $N_3$-8-**VI** | FN**Hcy**QQQRRFYEALHD<br>PNLNEEQRNAKIKSI<br>RDD | 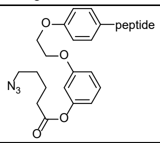 | 4443.9 | 4443.8 |
| $N_3$-**7-VII** | FN**C**QQQRRFYEALHD<br>PNLNEEQRNAKIKSI<br>RDD | 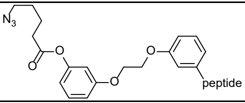 | 4429.9 | 4430.6 |
| $N_3$-**8-VII** | FN**Hcy**QQQRRFYEALHD<br>PNLNEEQRNAKIKSI<br>RDD | 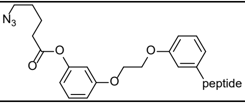 | 4443.9 | 4444.4 |
| $N_3$-**9-VII** | FNMQQQRRFYEALHD<br>PNLNEEQRNAKIKSI<br>**C**DD | 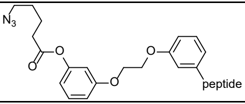 | 4404.9 | 4405.8 |
| $N_3$-**10-VII** | FNMQQQRRFYEALHD<br>PNLNEEQRNAKIKSI<br>R**C**D | 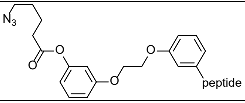 | 4446.0 | 4445.7 |
| $N_3$-**11-VII** | FNMQQQRRFYEALHD<br>PNLNEEQRNAKIKSI<br>R**Hcy**D | 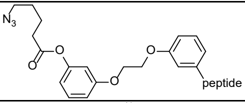 | 4460.0 | 4460.2 |
| $N_3$-**12-VIII** | FNMQQQRRFYEALHD<br>PNLN**Hcy**EQRNAKIKSI<br>RDD | 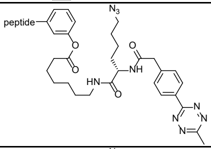 | 4678.1 | 4678.5 |
| $N_3$-**13-VIII** | $H_2N$-PEG8-<br>FNMQQQRRFYEALHD<br>PNLN**Hcy**EQRNAKIKSI<br>RDD | 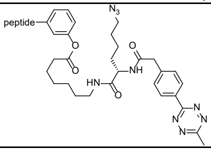 | 5101.8 | 5102.2 |

**Table S2**. Quantification of azido transfer on pure human $IgG_1$ Trastuzumab.

| Peptide | Elect rophi le | R | Equivalent of reagent | Reaction buffer | Reaction Temp. | Result analyzed by Q-ToF LC-MS | | |
|---|---|---|---|---|---|---|---|---|
| | | | | | | Light chain total conversion | Heavy chain total conversion | Heavy chain total conversion ratio (+1/+2) |
| $N_3$-1 | I | 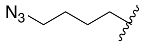 | 10 | 100mM HEPES pH 8.5 | rt | <1% | 17% | >99% / <1% |
| $N_3$-1 | I | 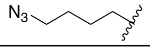 | 10 | PBS | rt | <1% | 12% | >99% / <1% |
| $N_3$-2 | I | 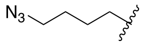 | 10 | 100mM HEPES pH 8.5 | rt | 5% | 99% | >99% / <1% |
| $N_3$-2 | I | 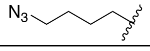 | 20 | PBS | rt | 12% | 68% | 88/12 |
| $N_3$-3 | I | 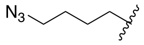 | 10 | 100mM HEPES pH 8.5 | rt | 5% | 95% | 95/5 |
| $N_3$-3 | I | 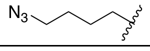 | 10 | PBS | 37 | 4% | 86% | 95/5 |
| $N_3$-12 | I | 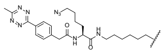 | 10+10[a] | 100mM HEPES pH 8.7 | 37 | <1% | 84% | >99% / <1% |

[a] 10 equivalents of the reagent were added at the beginning and after 4 hours of reaction, respectively.

**Table S3**. Quantification of azido transfer on different pure human and mouse IgG subtypes.

| IgG | Peptide | Electrophile | Equivalent of reagent | Reaction temp. | Result analyzed by Q-ToF LC-MS | | |
|---|---|---|---|---|---|---|---|
| | | | | | Light chain total conversion | Light chain total conversion | Heavy chain total conversion ratio (+1/+2) |
| Denosumab (hIgG2) | $N_3$-2 | I | 20 | rt | 1% | >99% | >99% / <1% |
| Dupilumab (hIgG4) | $N_3$-2 | I | 20 | rt | 13% | 96% | >99% / <1% |
| Mouse IgG1 | $N_3$-2 | I | 20 | rt | 2% | 69% | >99% / <1% |

**Table S4**. Quantification of GLP-1a transfer on pure human $IgG_1$ Trastuzumab.

| Peptide | Equivalent of reagent | Buffer | Reaction Temp. | Result analyzed by Q-ToF LC-MS | | |
|---|---|---|---|---|---|---|
| | | | | Light chain total conversion | Heavy chain total conversion | Heavy chain total conversion ratio (+1/+2) |
| 2a | 20 | HEPES | rt | <1% | 27% | >99% / <1% |
| 2a | 20 | HEPES | 37 °C | <1% | 33% | 94/6 |
| 3a | 20 | HEPES | rt | <1% | 26% | >99% / <1% |
| 3a | 20 | HEPES | 37 °C | <1% | 28% | >99% / <1% |
| 3a | 20 | PBS | 37 °C | <1% | 16% | >99% / <1% |

**Table S5.** Safety profile of GLP-1a conjugates in female WT Swiss mice. All GLP-1a drugs were injected subcutaneously. Every cohort contains 15 mice total, except for semaglutide at 0.5 mg/kg and 3 mg/kg, and **2a** stacking dose (3 × 10 mg/kg) which contain 5 mice each. Measured mean fasting blood glucose in naïve female WT Swiss mice: 134 ± 10 mg/dL.

| Compound | Dose (mg/kg) | Mean fasting blood glucose (mg/dL)* | Number of mice with Hypoglycemia* | | | Number of mice with Piloerection | | | Number of mice showing significant aggressiveness | MTD |
|---|---|---|---|---|---|---|---|---|---|---|
| | | | Mild (40-60 mg/dL) | Moderate (20-40 mg/dL) | Extreme (≤ 20 mg/dL) | Mild | Moderate | Severe | | |
| Semaglutide | 0.5 | 60.8 ± 9.4 | 2 | - | - | - | - | - | - | 10 mg/kg |
| | 3 | 61.4 ± 13.5 | 1 | 1 | - | - | 2[¤] | 3[¤] | 5[¤] | |
| | 10 | 40.4 ± 12.1 | 1 | 2 | 1 | - | - | 15[£] | 15[£] | |
| GLP-1**a** | 10 | 40.8 ± 11.8 | - | 3 | 1 | - | 15[¤] | - | 15[¤] | 10 mg/kg |
| **2a** | 10 | 81.8 ± 9.4 | - | - | - | - | - | - | - | ≥ 30 mg/kg |
| | 3 × 10, over 3 weeks | 93.2 ± 11.4 | - | - | - | - | - | - | - | |
| | 30 | 94.0 ± 4.7 | - | - | - | 2* | - | - | - | |
| **3a** | 10 | 82.4 ± 9.7 | - | - | - | - | - | - | - | > 10 mg/kg |

* At 24 h post drug injection
¤ Over 48 h post injection
£ Over 72 h post injection
MTD: Maximum tolerated dose among the assessed range.

**Table S6**. Safety profile after 10 mg/kg of GLP-1a conjugates in male (M) and female (F) Lep$^{ob/ob}$ mice. IP: intraperitoneal injection. SC: subcutaneous injection. Cohorts of 3 mice per row. Measured mean fasting blood glucose in female C57BL/6J mice: 174 ± 15 mg/dL; male C57BL/6J mice:184 ± 10 mg/dL; female Lep$^{ob/ob}$ mice: 374 ± 29 mg/dL; male Lep$^{ob/ob}$ mice: 407 ± 44 mg/dL.

| Compound | Sexe and route of injection | Mean fasting blood glucose (mg/dL)* | Number of mice with Hypoglycemia* | | | Number of mice with Piloerection | | | Number of mice showing significant aggressiveness |
|---|---|---|---|---|---|---|---|---|---|
| | | | Mild (40-60 mg/dL) | Moderate (20-40 mg/dL) | Extreme (≤ 20 mg/dL) | Mild | Moderate | Severe | |
| Semaglutide | F, IP | 104±6 | - | - | - | - | - | 3 | 3 |
| | F, SC | 97±7 | - | - | - | - | 3 | - | 3 |
| | M, IP | 111±14 | - | - | - | - | - | 3 | 3 |
| | M, SC | 75±12 | 2 | - | - | - | 3 | - | 3 |
| **2a** | F, IP | 137±8 | - | - | - | 2 | 1 | - | - |
| | F, SC | 104±5 | - | - | - | - | - | - | - |
| | M, IP | 149±32 | - | - | - | - | 3 | - | 1 |
| | M, SC | 108±4 | - | - | - | 1 | - | - | - |
| **3a** | F, IP | 147±35 | - | - | - | 1 | - | - | - |
| | F, SC | 130±12 | - | - | - | - | - | - | - |
| | M, IP | 167±22 | - | - | - | 2 | - | - | - |
| | M, SC | 102±9 | - | - | - | - | - | - | - |

**Table S7**. Biodistribution of the affinity peptides [$^{89}$Zr]Zr-**1-3** in the main organs at 144 h after intravenous injection in female WT Swiss mice. Quantification was performed using Wizard$^2$ 3" (Perkin Elmer) gamma counter. Data are represented as mean ± SEM (n=3-4 mice per group).

| **Mean ±SEM %IA/g** | **[$^{89}$Zr]Zr-1** | **[$^{89}$Zr]Zr-2** | **[$^{89}$Zr]Zr-3** |
|---|---|---|---|
| Blood | 0.09 ± 0.01 | 0.09 ± 0.00 | 0.07 ± 0.04 |
| Urine | 2.79 ± 3.11 | 1.67 ± 0.61 | 1.38 ± 0.27 |
| Heart | 0.21 ± 0.01 | 0.18 ± 0.01 | 0.15 ± 0.01 |
| Lungs | 18.0 ± 3.25 | 0.56 ± 0.25 | 0.34 ± 0.03 |
| Liver | 25.7 ± 2.79 | 34.7 ± 3.18 | 23.8 ± 2.03 |
| Stomach | 0.70 ± 0.45 | 0.95 ± 1.05 | 0.20 ± 0.14 |
| Pancreas | 0.28 ± 0.10 | 0.18 ± 0.03 | 0.16 ± 0.02 |
| Spleen | 11.5 ± 1.34 | 42.8 ± 1.26 | 25.5 ± 2.29 |
| Caecum | 0.18 ± 0.04 | 0.14 ± 0.03 | 0.13 ± 0.03 |
| Small intestines | 0.14 ± 0.04 | 0.17 ± 0.08 | 0.11 ± 0.02 |
| Colon | 0.18 ± 0.02 | 0.16 ± 0.02 | 0.13 ± 0.02 |
| Feces | 0.27 ± 0.02 | 0.30 ± 0.06 | 0.26 ± 0.06 |
| Kidneys | 44.2 ± 6.24 | 34.7 ± 10.2 | 51.6 ± 10.3 |
| Bladder | 0.54 ± 0.08 | 0.50 ± 0.14 | 0.55 ± 0.05 |
| Uterus | 0.19 ± 0.07 | 0.20 ± 0.07 | 0.10 ± 0.03 |
| Gastroc. Muscle | 0.10 ± 0.01 | 0.10 ± 0.01 | 0.07 ± 0.01 |
| Bone | 1.09 ± 0.28 | 0.74 ± 0.18 | 0.58 ± 0.15 |
| Skin | 0.20 ± 0.03 | 0.21 ± 0.02 | 0.12 ± 0.02 |
| Brain | 0.03 ± 0.00 | 0.03 ± 0.01 | 0.03 ± 0.00 |
| Eyes | 0.20 ± 0.01 | 0.29 ± 0.03 | 0.23 ± 0.03 |

%IA/g: percent of injected activity per gram; Gastroc. muscle: gastrocnemius muscle.

**Table S8**. Biodistribution of the affinity peptides [$^{89}$Zr]Zr-**2** in the main organs at 2 h, 24 h, 72 h, 144 h, and 240 h after subcutaneous injection in female WT Swiss mice. Quantification was performed using Wizard$^2$ 3'' (Perkin Elmer) gamma counter. Data are represented as mean ± SEM (n= 3-4 mice per group for each time point).

| **Mean ±SEM %IA/g** | **[$^{89}$Zr]Zr-2** | | | | |
|---|---|---|---|---|---|
| | **2 h** | **24 h** | **72 h** | **144 h** | **240 h** |
| Blood | 0.05 ± 0.07 | 0.07 ±0.04 | 0.04 ± 0.01 | 0.03 ± 0.01 | 0.02 ± 0.00 |
| Urine | 67.0 ± 35.6 | 0.66 ± 0.29 | 0.54 ± 0.29 | 0.45 ± 0.41 | 0.08 ± 0.02 |
| Heart | 0.35 ± 0.07 | 0.06 ± 0.00 | 0.06 ± 0.02 | 0.04 ± 0.01 | 0.04 ± 0.00 |
| Lungs | 0.80 ± 0.36 | 0.12 ± 0.04 | 0.11 ± 0.05 | 0.09 ± 0.04 | 0.05 ± 0.01 |
| Liver | 2.45 ± 0.42 | 1.22 ± 0.65 | 1.34 ± 0.74 | 1.08 ± 0.73 | 0.49 ± 0.05 |
| Stomach | 0.19 ± 0.09 | 0.14 ± 0.10 | 0.05 ± 0.03 | 0.07 ± 0.06 | 0.01 ± 0.00 |
| Pancreas | 0.33 ± 0.13 | 0.08 ± 0.06 | 0.10 ± 0.06 | 0.07 ± 0.06 | 0.03 ± 0.01 |
| Spleen | 0.39 ± 0.15 | 0.12 ± 0.03 | 0.13 ± 0.04 | 0.12 ± 0.05 | 0.07 ± 0.01 |
| Caecum | 10.7 ± 10.5 | 0.39 ± 0.13 | 0.09 ± 0.04 | 0.06 ± 0.04 | 0.02 ± 0.00 |
| Small intestines | 9.21 ± 5.60 | 0.07 ± 0.02 | 0.04 ± 0.01 | 0.03 ± 0.03 | 0.01 ± 0.00 |
| Colon | 2.44 ± 2.09 | 0.21 ± 0.06 | 0.08 ± 0.04 | 0.07 ± 0.05 | 0.02 ± 0.00 |
| Feces | 0.57 ± 0.45 | 1.58 ± 0.80 | 0.19 ± 0.07 | 0.19 ± 0.14 | 0.05 ± 0.01 |
| Kidneys | 32.1 ± 13.3 | 17.3 ± 9.68 | 16.1 ± 11.5 | 15.2 ± 10.7 | 2.41 ± 0.95 |
| Bladder | 3.29 ± 1.97 | 0.22 ± 0.03 | 0.20 ± 0.04 | 0.18 ± 0.06 | 0.12 ± 0.02 |
| Uterus | 0.27 ± 0.06 | 0.04 ± 0.01 | 0.06 ± 0.04 | 0.04 ± 0.02 | 0.04 ± 0.01 |
| Gastroc. Muscle | 0.16 ± 0.03 | 0.03 ± 0.00 | 0.03 ± 0.01 | 0.03 ± 0.01 | 0.02 ± 0.00 |
| Bone | 0.20 ± 0.03 | 0.06 ± 0.02 | 0.17 ± 0.10 | 0.10 ± 0.05 | 0.06 ± 0.00 |
| Skin | 0.51 ± 0.12 | 0.07 ± 0.02 | 0.10 ± 0.02 | 0.06 ± 0.03 | 0.05 ± 0.00 |
| Brain | 0.05± 0.01 | 0.01 ± 0.00 | 0.01 ± 0.00 | 0.01 ± 0.00 | 0.01 ± 0.00 |
| Eyes | 0.26 ± 0.04 | 0.05 ± 0.01 | 0.05 ±0.00 | 0.05 ± 0.00 | 0.08 ± 0.02 |
| Inguinal lymph nodes | 1.49 ± 0.49 | 0.49 ± 0.55 | 0..36 ± 0.23 | 0.23 ± 0.16 | 0.15 ± 0.06 |
| Axillary lymph nodes | 0.53 ± 0.06 | 0.23 ± 0.05 | 0.20 ± 0.02 | 0.13 ± 0.02 | 0.14 ± 0.05 |
| Thymus | 0.42 ± 0.13 | 0.08 ± 0.01 | 0.09 ± 0.04 | 0.07 ± 0.02 | 0.08 ± 0.03 |

%IA/g: percent of injected activity per gram; Gastroc. muscle: gastrocnemius muscle.

**Table S9**. Biodistribution of the affinity peptides [$^{89}$Zr]Zr-**3** in the main organs at 2 h, 24 h, 72 h, 144 h, and 240 h after subcutaneous injection in female WT Swiss mice. Quantification was performed using Wizard$^2$ 3'' (Perkin Elmer) gamma counter. Data are represented as mean ± SEM (n= 3-4 mice per group for each time point).

| **Mean ±SEM %IA/g** | **[$^{89}$Zr]Zr-3** | | | | |
|---|---|---|---|---|---|
| | **2 h** | **24 h** | **72 h** | **144 h** | **240 h** |
| Blood | 0.49 ± 0.01 | 0.11 ±0.03 | 0.04 ± 0.01 | 0.03 ± 0.01 | 0.01 ± 0.00 |
| Urine | 62.5 ± 21.3 | 0.65 ± 0.22 | 0.78 ± 0.66 | 0.71 ± 0.85 | 0.08 ± 0.02 |
| Heart | 0.48 ± 0.29 | 0.06 ± 0.01 | 0.04 ± 0.00 | 0.04 ± 0.00 | 0.03 ± 0.01 |
| Lungs | 0.47 ± 0.03 | 0.10 ± 0.01 | 0.06 ± 0.01 | 0.05 ± 0.00 | 0.04 ± 0.01 |
| Liver | 5.30 ± 0.22 | 1.87 ± 0.50 | 1.53 ± 0.11 | 1.50 ± 0.35 | 1.23 ± 0.34 |
| Stomach | 0.42 ± 0.20 | 0.13 ± 0.09 | 0.03 ± 0.01 | 0.03 ± 0.01 | 0.01 ± 0.01 |
| Pancreas | 0.33 ± 0.11 | 0.05 ± 0.01 | 0.04 ± 0.01 | 0.04 ± 0.02 | 0.03 ± 0.00 |
| Spleen | 0.46 ± 0.06 | 0.18 ± 0.05 | 0.16 ± 0.01 | 0.14 ± 0.01 | 0.10 ± 0.01 |
| Caecum | 14.4 ± 6.85 | 0.62 ± 0.09 | 0.05 ± 0.01 | 0.08 ± 0.05 | 0.01 ± 0.00 |
| Small intestines | 14.8 ± 1.69 | 0.13 ± 0.03 | 0.03 ± 0.00 | 0.03 ± 0.00 | 0.01 ± 0.00 |
| Colon | 4.10 ± 1.78 | 0.38 ± 0.09 | 0.05 ± 0.02 | 0.05 ± 0.04 | 0.02 ± 0.00 |
| Feces | 1.12 ± 1.10 | 2.05 ± 0.90 | 0.17 ± 0.05 | 0.20 ± 0.16 | 0.03 ± 0.01 |
| Kidneys | 13.5 ± 1.31 | 5.58 ± 0.83 | 4.69 ± 0.17 | 4.53 ± 1.32 | 1.86 ± 1.21 |
| Bladder | 5.89 ± 5.40 | 0.17 ± 0.02 | 0.11 ± 0.01 | 0.10 ± 0.02 | 0.11 ± 0.03 |
| Uterus | 0.26 ± 0.07 | 0.05 ± 0.01 | 0.03 ± 0.01 | 0.04 ± 0.01 | 0.03 ± 0.01 |
| Gastroc. Muscle | 0.20 ± 0.07 | 0.03 ± 0.00 | 0.02 ± 0.00 | 0.02 ± 0.00 | 0.02 ± 0.00 |
| Bone | 0.26 ± 0.08 | 0.07 ± 0.01 | 0.09 ± 0.01 | 0.07 ± 0.01 | 0.08 ± 0.01 |
| Skin | 0.36 ± 0.04 | 0.07 ± 0.01 | 0.06 ± 0.00 | 0.05 ± 0.01 | 0.05 ± 0.01 |
| Brain | 0.07± 0.03 | 0.01 ± 0.00 | 0.01 ± 0.00 | 0.01 ± 0.00 | 0.01 ± 0.00 |
| Eyes | 0.25 ± 0.14 | 0.05 ± 0.01 | 0.03 ±0.00 | 0.05 ± 0.01 | 0.05 ± 0.01 |
| Inguinal lymph nodes | 2.70 ± 1.83 | 0.12 ± 0.02 | 0.16 ± 0.04 | 0.22 ± 0.19 | 0.14 ± 0.06 |
| Axillary lymph nodes | 0.69 ± 0.27 | 0.18 ± 0.08 | 0.18 ± 0.03 | 0.13 ± 0.03 | 0.12 ± 0.02 |
| Thymus | 0.28 ± 0.03 | 0.07 ± 0.01 | 0.05 ± 0.01 | 0.05 ± 0.01 | 0.05 ± 0.01 |

%IA/g: percent of injected activity per gram; Gastroc. muscle: gastrocnemius muscle.

**Table S20**. Biodistribution of the affinity peptides [$^{89}$Zr]Zr-**1,2** in the main organs at 2 h, 24 h, 72 h, and 144 h after intraperitoneal injection in female WT Swiss mice. Quantification was performed using Wizard$^2$ 3" (Perkin Elmer) gamma counter. Data are represented as mean ± SEM (n= 3-4 mice per group for each time point).

| **Mean ±SEM %IA/g** | **[$^{89}$Zr]Zr-1** | | | | **[$^{89}$Zr]Zr-2** | | | |
|---|---|---|---|---|---|---|---|---|
| | **2 h** | **24 h** | **72 h** | **144 h** | **2 h** | **24 h** | **72 h** | **144 h** |
| Blood | 0.52 ±0.09 | 0.10 ± 0.01 | 0.03 ± 0.00 | 0.04 ± 0.01 | 0.34 ± 0.19 | 0.28 ± 0.06 | 0.16 ± 0.01 | 0.10 ± 0.01 |
| Urine | 93.9 ± 23.3 | 0.38 ± 0.18 | 0.14 ± 0.04 | 0.40 ± 0.24 | 68.2 ± 9.98 | 1.84 ± 0.39 | 0.71 ± 0.50 | 0.72 ± 0.11 |
| Heart | 0.36 ± 0.07 | 0.09 ± 0.04 | 0.08 ± 0.04 | 0.08 ± 0.01 | 0.39 ± 0.06 | 0.18 ± 0.05 | 0.20 ± 0.04 | 0.22 ± 0.05 |
| Lungs | 0.59 ± 0.09 | 0.12 ± 0.03 | 0.32± 0.25 | 0.27 ± 0.17 | 1.46 ± 1.07 | 0.21 ± 0.09 | 0.55 ± 0.29 | 0.37 ± 0.11 |
| Liver | 5.21 ± 0.49 | 0.82 ± 0.32 | 0.56 ± 0.04 | 0.79 ± 0.07 | 3.12 ± 0.42 | 2.25 ± 0.27 | 2.04 ± 0.70 | 2.89 ± 0.92 |
| Stomach | 1.31 ± 0.58 | 0.30 ± 0.08 | 0.15 ± 0.02 | 0.29 ± 0.09 | 2.06 ± 0.82 | 3.71 ± 0.99 | 3.03 ± 0.97 | 2.53 ± 0.76 |
| Pancreas | 1.01 ± 0.23 | 0.68 ± 0.20 | 0.53 ± 0.16 | 0.48 ± 0.05 | 4.81 ± 1.76 | 3.00 ± 0.71 | 4.51 ± 0.51 | 9.13 ± 0.97 |
| Spleen | 0.95 ± 0.32 | 0.61 ±0.11 | 0.64 ± 0.05 | 0.86 ± 0.06 | 3.15 ± 0.39 | 4.87 ± 1.16 | 5.85 ± 1.55 | 11.7 ± 0.49 |
| Caecum | 16.51 ± 5.42 | 0.21 ± 0.04 | 0.17 ± 0.13 | 0.11 ± 0.03 | 11.7 ± 1.28 | 0.71 ± 0.22 | 0.41 ± 0.14 | 0.52 ± 0.16 |
| Small intestines | 16.37 ± 3.24 | 0.17 ± 0.04 | 0.10 ± 0.03 | 0.18 ± 0.03 | 8.04 ± 1.15 | 1.11 ± 0.42 | 1.25 ± 1.03 | 0.90 ± 0.53 |
| Colon | 1.66 ± 0.69 | 0.19 ± 0.06 | 0.10 ± 0.02 | 0.13 ± 0.02 | 2.83 ± 0.35 | 0.71 ± 0.09 | 0.70 ± 0.48 | 1.39 ± 1.06 |
| Feces | 2.56 ± 3.03 | 0.40 ± 0.13 | 0.13 ± 0.01 | 0.15 ± 0.03 | 0.59 ± 0.24 | 2.01 ± 0.93 | 0.73 ± 0.45 | 0.55 ± 0.32 |
| Kidneys | 10.8 ± 3.02 | 3.66 ± 0.58 | 3.27 ± 0.32 | 2.90 ± 0.34 | 12.3 ± 0.72 | 6.71 ± 1.08 | 7.00 ± 1.31 | 7.39 ± 1.63 |
| Bladder | 3.83 ± 1.54 | 0.29 ± 0.03 | 0.24 ± 0.08 | 0.34 ± 0.03 | 3.41 ± 0.90 | 0.70 ± 0.21 | 0.65 ± 0.22 | 1.00 ± 0.27 |
| Uterus | 0.62 ± 0.18 | 0.16 ± 0.06 | 0.15 ± 0.02 | 0.18 ± 0.03 | 1.07 ± 0.45 | 0.64 ± 0.33 | 1.14 ± 0.54 | 1.09 ± 0.69 |
| Gastroc. Muscle | 0.26 ± 0.06 | 0.05 ± 0.01 | 0.04 ± 0.02 | 0.05 ± 0.01 | 0.45 ± 0.27 | 0.21 ± 0.14 | 0.19 ± 0.09 | 0.13 ± 0.05 |
| Bone | 0.22 ± 0.06 | 0.06 ± 0.01 | 0.07 ± 0.00 | 0.09 ± 0.00 | 0.34 ± 0.20 | 0.17 ± 0.05 | 0.51 ± 0.16 | 0.54 ± 0.16 |
| Skin | 0.76 ± 0.34 | 0.11 ± 0.01 | 0.08 ± 0.02 | 0.25 ± 0.04 | 1.03 ± 0.80 | 0.25 ± 0.22 | 0.45 ± 0.19 | 0.50 ± 0.29 |
| Brain | 0.08 ± 0.01 | 0.01 ± 0.00 | 0.01 ± 0.00 | 0.01 ± 0.00 | 0.21 ± 0.20 | 0.03 ± 0.01 | 0.03 ± 0.01 | 0.04 ± 0.02 |
| Eyes | 0.26 ± 0.09 | 0.05 ± 0.01 | 0.05 ± 0.01 | 0.07 ± 0.03 | 0.24 ± 0.03 | 0.06 ± 0.02 | 0.10 ± 0.03 | 0.09 ± 0.01 |

%IA/g: percent of injected activity per gram; Gastroc. muscle: gastrocnemius muscle.

**Supplementary Figures**

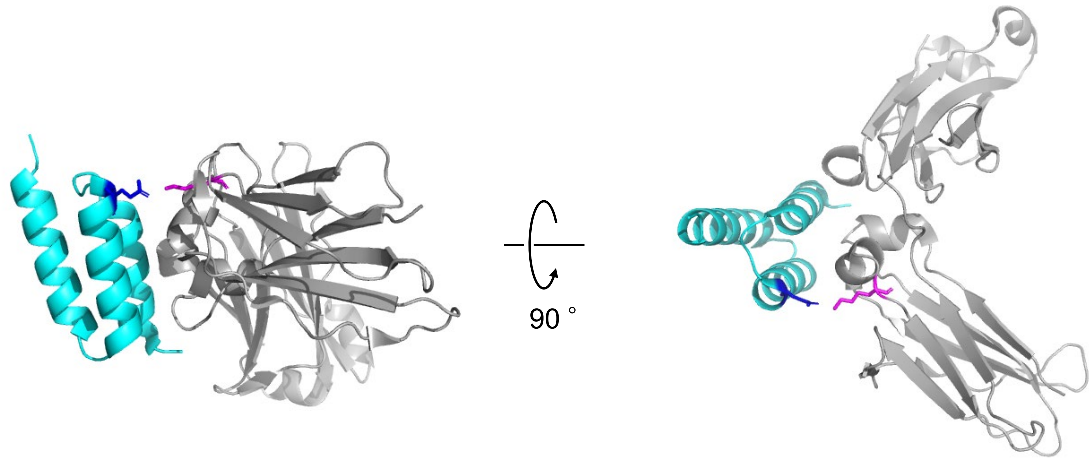

**Figure S1.** The co-crystal structure of the human IgG Fc fragment and the B domain of Protein A (PDB: 1FC2). Gray: Human Fc fragment, Cyan: B domain of protein A, Magenta: K317 of the human IgG Fc fragment, Blue: E20 of the B domain of protein A.

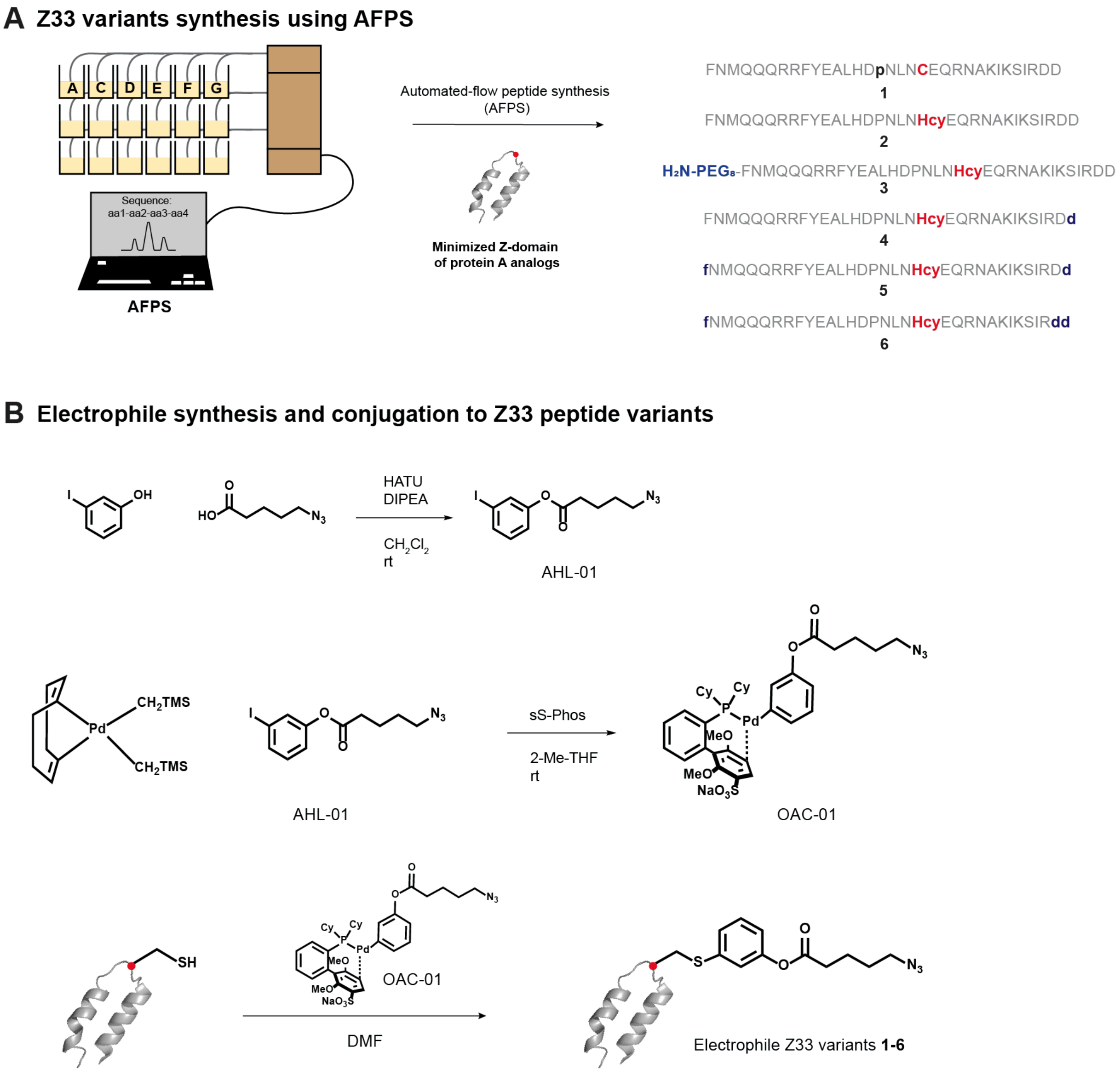

**Figure S2**. Synthesis of the electrophile affinity peptides **1-6**. (A) Variants of the minimized Z-domain of protein A (Z33 peptide) were synthesized using automated fast-flow peptide synthesis (AFPS) as previously described (2). (B) Conjugation of the electrophile to the Z33 peptide variants **1-6** to obtain $N_3$-**1-6** via an arylation reaction using palladium reagent as previously described (3).

**A**

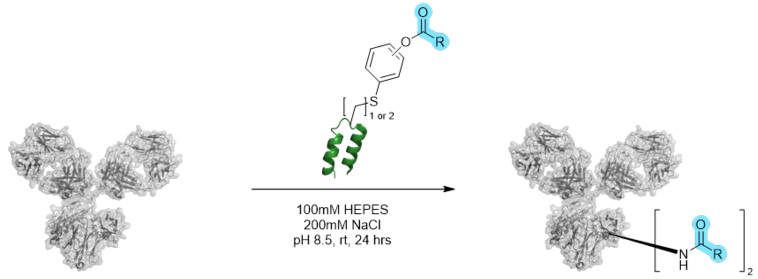

| Entry | Sequence of hIgG-fc Binding Peptide | Structure Attached on the C or Hcy | Light Chain Reactivity | Heavy Chain Reactivity | Heavy Chain reaction selectivity |
|---|---|---|---|---|---|
| 1 | FNMQQQRRFYEALHDPNLN**C**EQRNAKIKSIRDD | I | 6% | 61% | 90% |
| 2 | | I | 6% | **99%** | **>99%** |
| 3 | FNMQQQRRFYEALHDPNLN**Hcy**EQRNAKIKSIRDD | II | 10% | 47% | >99% |
| 4 | | III | 20% | 47% | 77% |
| 5 | FNMQQQRRFYEALHD**p**NLN**C**EQRNAKIKSIRDD | I | <1% | 17% | >99% |

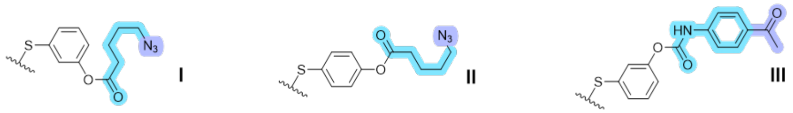

**B**

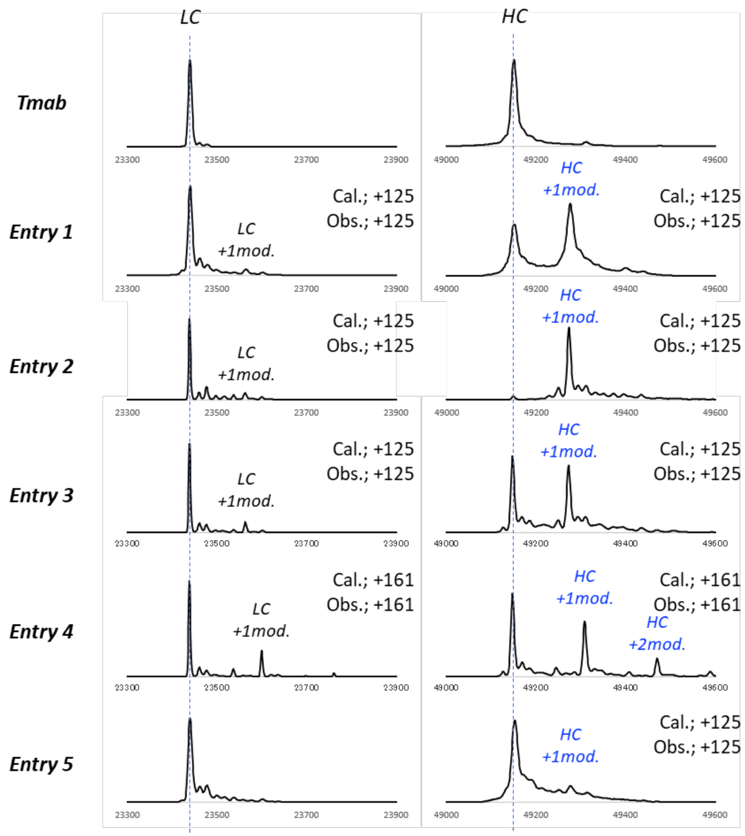

**Figure S3**. In vitro reactivity of selected electrophile affinity peptides for human IgG1 Trastuzumab (Tmab). (A) Percentage of modification on heavy and light chains of Tmab depending on the amino acid sequence of the affinity peptide and structure of the electrophiles. The conjugation site on the affinity peptide is highlighted in blue. “Heavy chain reaction selectivity” was calculated based on the percentage of lysine residues on the heavy chain with at least one modification. (B) Mass spectra (MS) post reaction. Tmab was deglycosylated and reduced beforehand. Left: MS for the light chain; Right: MS for the heavy chain.

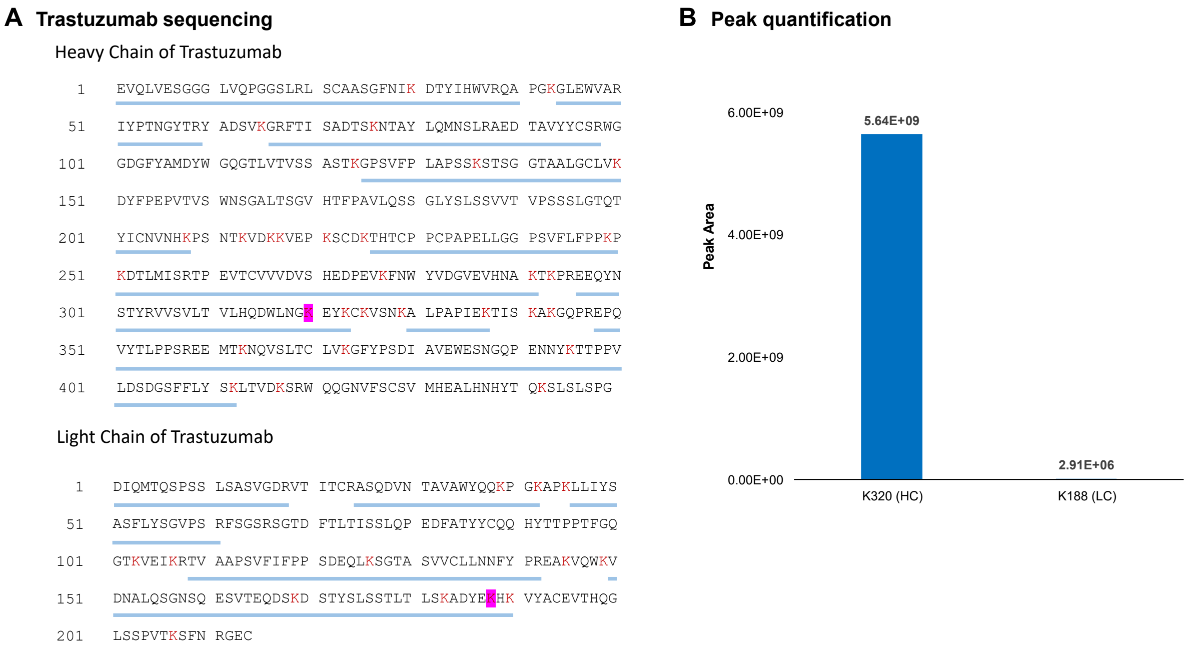

**Figure S4**. Identification of the sites of modification on IgG1 Trastuzumab (Tmab) after in vitro reaction with compound **2**. (A) Sequencing of Tmab after digestion with trypsin and analysis by LC-MS/MS. Lysines are in red. Pale blue bars below the sequence indicate the observed fragments. The lysines exhibiting modifications are highlighted in magenta. (B) Peak areas corresponding to the lysines modified by the affinity peptides.

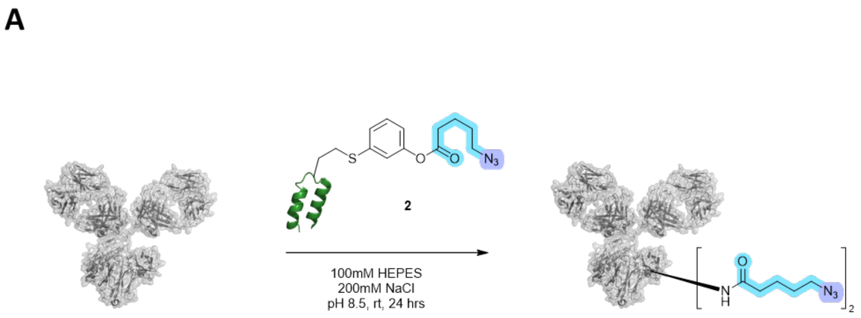

| Entry | Starting Material | Light Chain Reactivity | Heavy Chain Reactivity | Heavy Chain Reaction Selectivity |
|---|---|---|---|---|
| 1 | denosumab (human IgG2) | 1% | >99% | >99% |
| 2 | dupilumab (human IgG4) | 13% | 96% | >99% |
| 3 | mouse IgG1 | 2% | 69% | >99% |

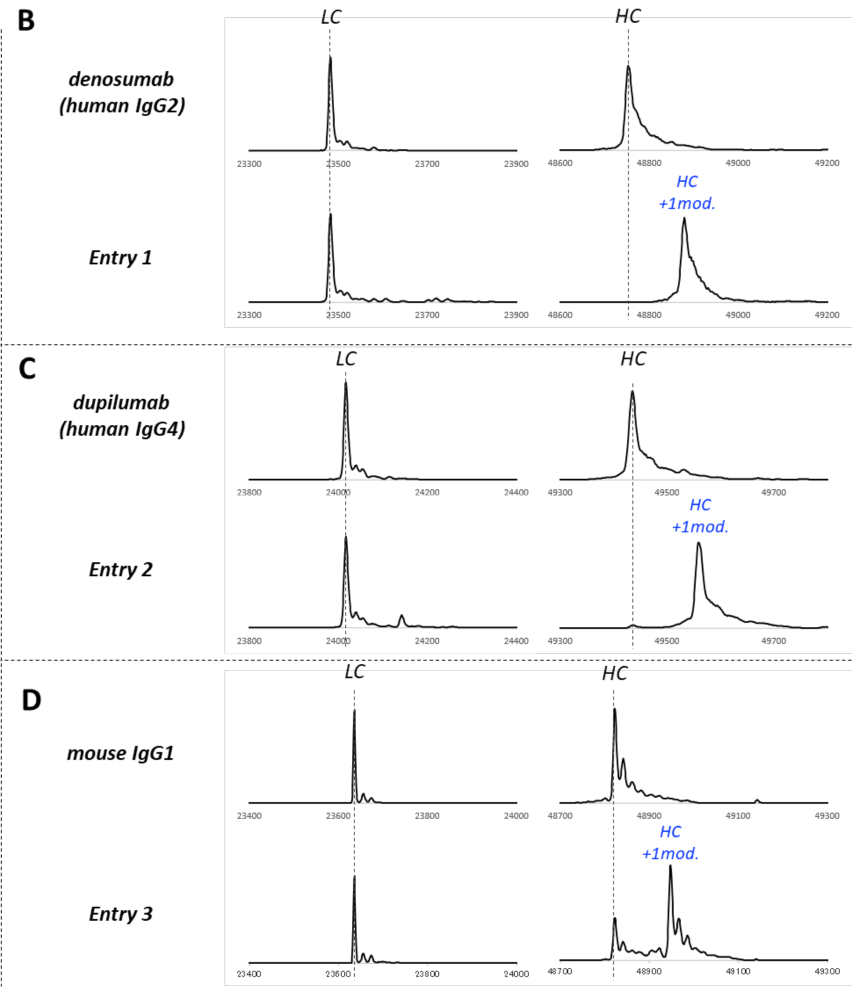

**Figure S5**. In vitro reactivity of compound **2** for other IgG subclasses and species. (A) "Heavy chain reaction selectivity" was calculated based on the percentage of lysine residues with at least one modification. (B-D) Mass spectra (MS) post reaction. IgGs were deglycosylated and reduced beforehand. Left: MS for the light chain; Right: MS for the heavy chain.

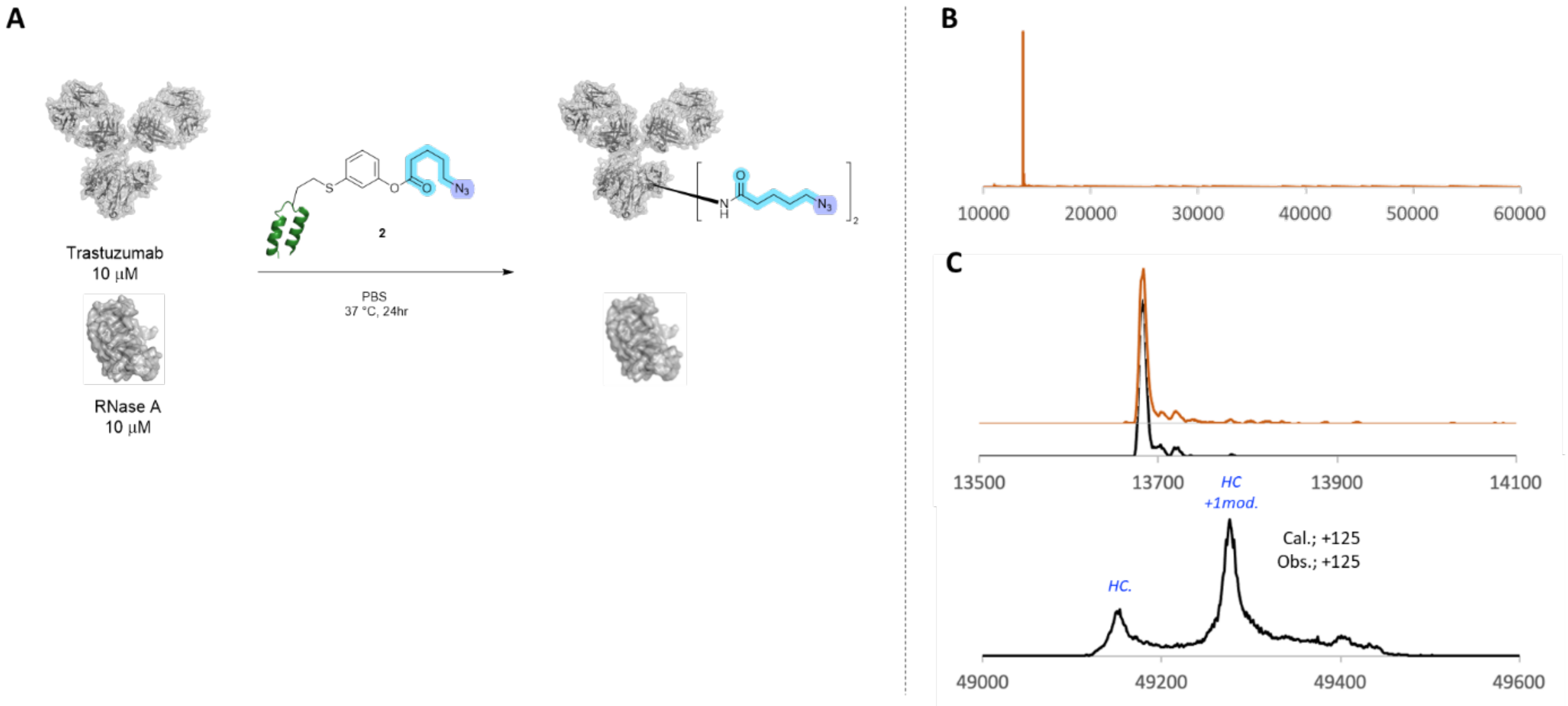

**Figure S6**. In vitro selectivity of compound **2** for IgG1 (A) Compound **2** incubation with Tmab and RNAse A (B) Mass spectrometry of native RNase A (C) Mass spectrometry after reaction with compound **2**. Data were obtained after glycan removal and disulfide reduction of Tmab. The top panel shows the mass of RNase A, and the bottom panel shows the mass of Tmab heavy chains. Orange trace indicates native RNase A, and black trace shows the mass obtained after reaction with compound **2**.

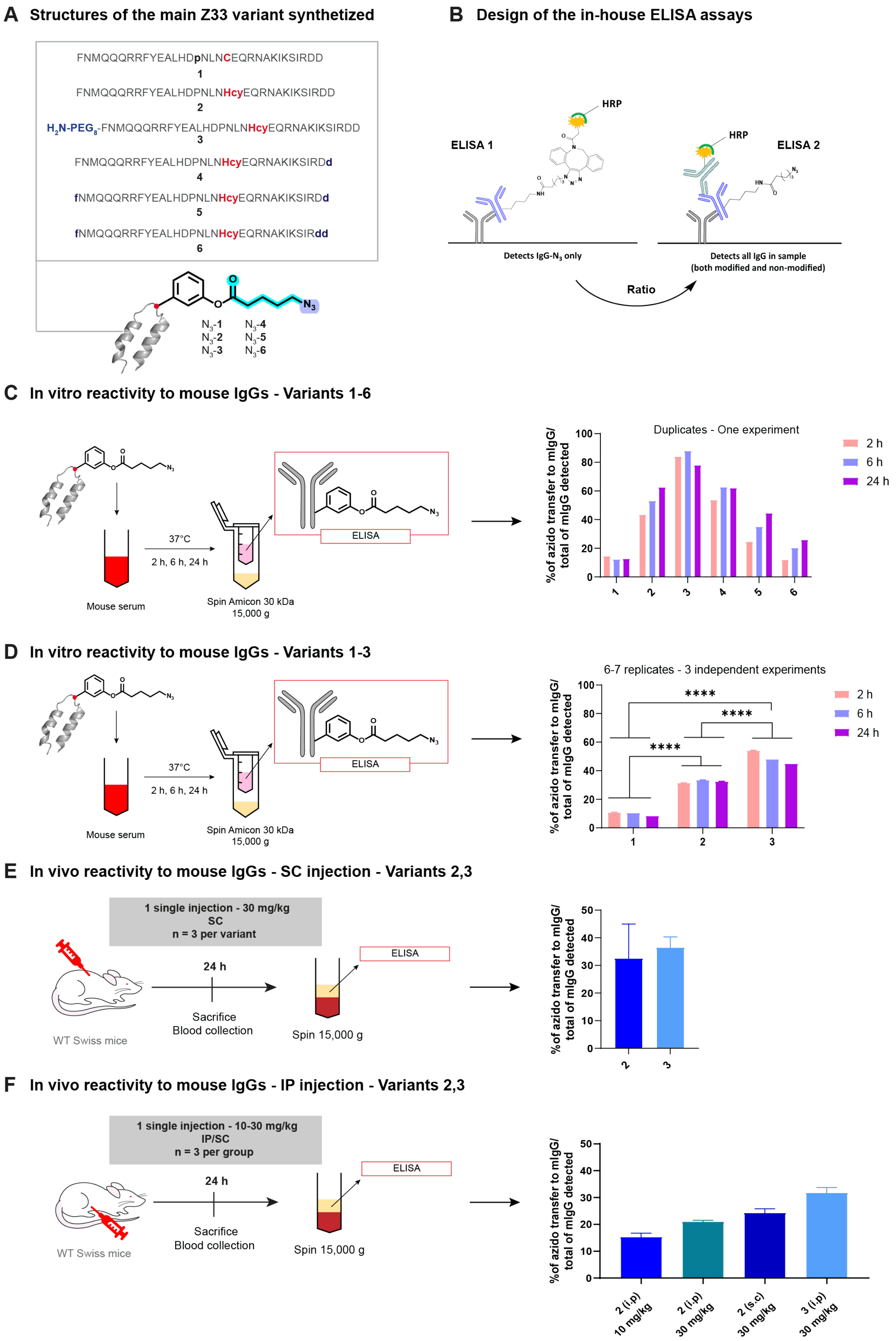
A Structures of the main Z33 variant synthetized
FNMQQQRRFYEALHDpNLNCEQRNAKIKSIRDD
1
FNMQQQRRFYEALHDPNLNHcyEQRNAKIKSIRDD
2
H2N-PEG8-FNMQQQRRFYEALHDPNLNHcyEQRNAKIKSIRDD
3
FNMQQQRRFYEALHDPNLNHcyEQRNAKIKSIRDd
4
fNMQQQRRFYEALHDPNLNHcyEQRNAKIKSIRDd
5
fNMQQQRRFYEALHDPNLNHcyEQRNAKIKSIRdd
6
N3-1 N3-2 N3-3 N3-4 N3-5 N3-6
B Design of the in-house ELISA assays
ELISA 1
HRP
ELISA 2
Detects IgG-N3 only
Detects all IgG in sample (both modified and non-modified)
Ratio
C In vitro reactivity to mouse IgGs - Variants 1-6
Mouse serum
37°C
2 h, 6 h, 24 h
Spin Amicon 30 kDa 15,000 g
ELISA
Duplicates - One experiment
%of azido transfer to mIgG/ total of mIgG detected
2 h
6 h
24 h
D In vitro reactivity to mouse IgGs - Variants 1-3
6-7 replicates - 3 independent experiments
****
E In vivo reactivity to mouse IgGs - SC injection - Variants 2,3
1 single injection - 30 mg/kg
SC
n = 3 per variant
24 h
Sacrifice
Blood collection
Spin 15,000 g
WT Swiss mice
F In vivo reactivity to mouse IgGs - IP injection - Variants 2,3
1 single injection - 10-30 mg/kg
IP/SC
n = 3 per group
2 (i.p) 10 mg/kg
2 (i.p) 30 mg/kg
2 (s.c) 30 mg/kg
3 (i.p) 30 mg/kg

**Figure S7**. In vitro and in vivo azido transfer to mouse IgG (mIgG) using electrophile affinity peptides $N_3$-**1-6**. (A) Sequences and structures of electrophile affinity peptide $N_3$-**1-6**. (B) ELISA assays E1 and E2 developed for the quantification of azido transfer to native mIgG. (C) In vitro azido transfer to mIgG using electrophile affinity peptides $N_3$-**1-6** quantified by E1/E2 after 2, 6, or 24 h of incubation (~4 nmol of $N_3$-**1-6**) in mouse sera. (D) In vitro azido transfer to mIgG using peptides electrophile affinity peptides $N_3$-**1-3** quantified by E1/E2 after 2, 6, or 24 h of incubation in mouse sera (~4 nmol of $N_3$-**1-6**). Data show mean ± SD from 6-7 replicates over 3 independent experiments. Statistical analysis was performed using non-parametric T-tests. *$P<0.05$, **$P<0.01$, ***$P<0.001$, ****$P<0.0001$. (E) In vivo azido transfer to mIgG in mice sera quantified by E1/E2 at 24 h after subcutaneous (SC) injection of 30 mg/kg (~280 nmol) of electrophile affinity peptides $N_3$-**2,3** in female WT Swiss mice. (F) In vivo azido transfer to mIgG in mice sera quantified by E1/E2 at 24 h after either SC or intraperitoneal (IP) injection of 10 mg/kg (~90 nmol) or 30 mg/kg (~280 nmol) of electrophile affinity peptides $N_3$-**2,3** in female WT Swiss mice.

**A Synthesis of DBCO-Semaglutide, DBCO-Liraglutide, and Tz-Tirzepatide**

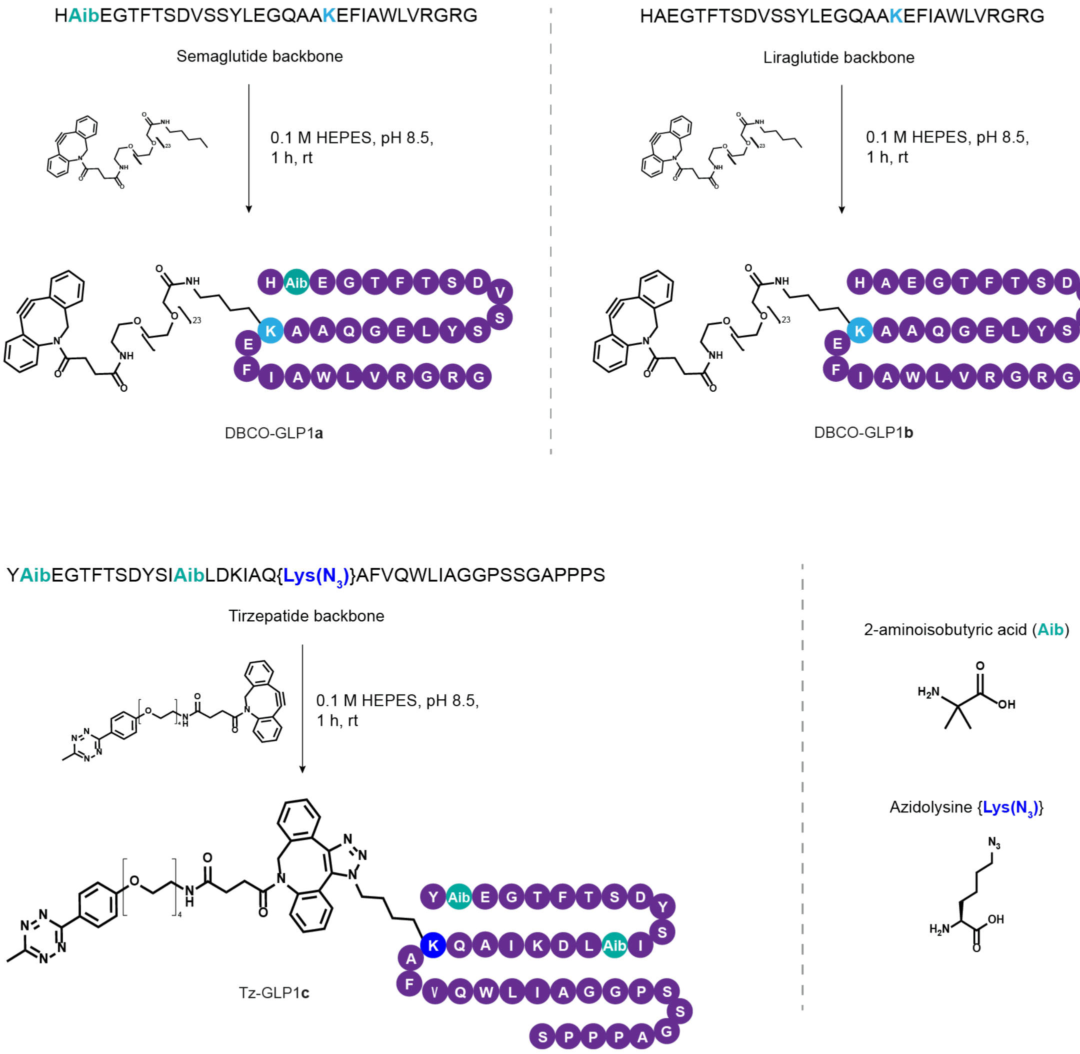

**B Multi-step synthesis of affinity peptide 2c**

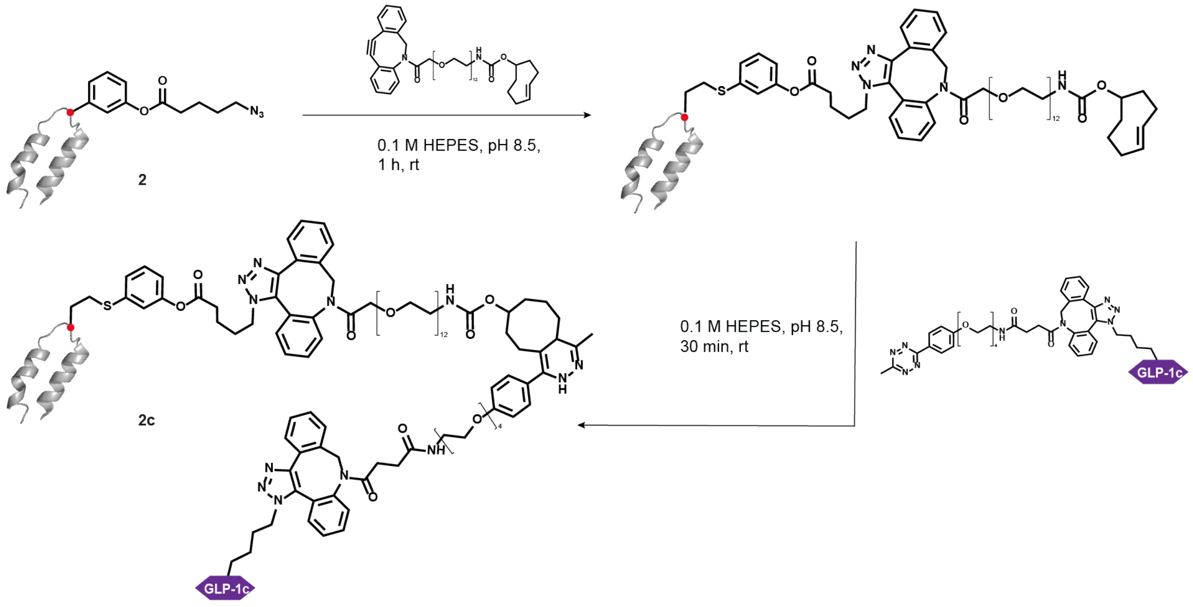

**Figure S8**. Synthesis of GLP-1 conjugates. (A) Semaglutide, Liraglutide, and Tirzepatide peptide backbones were synthesized using automated fast-flow peptide synthesis (AFPS) (2). Semaglutide and Liraglutide were then conjugated to dibenzyl cyclooctyne (DBCO)-$PEG_{23}$ moiety to obtain DBCO-GLP-1-**a,b** reagents. Tirzepatide was conjugated to bifunctional DBCO-$PEG_4$-tetrazine (Tz) to obtain reagent Tz-GLP-1**c**. (B) Synthesis of electrophile affinity peptide **2c**. Affinity peptide **2** was first conjugated to DBCO-$PEG_{12}$-transcyclooctene, then conjugated to Tz-$PEG_4$-GLP1**c** through the inverse-electron demand Diels-Alder click cycloaddition to obtain reagent **2c**.

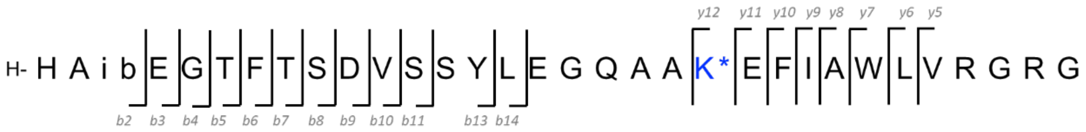

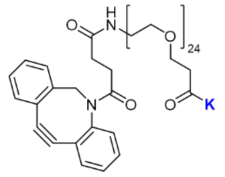

* Structure attached to the K

| Fragment | Theoretical m/z | Observed m/z | Error Value (ppm) |
|---|---|---|---|
| *b2* | 223.1190 | 223.1191 | 0.4 |
| *b3* | 352.1615 | 352.1621 | 1.7 |
| *b4* | 409.1830 | 409.1840 | 2.4 |
| *b5* | 510.2307 | 510.2318 | 2.2 |
| *b6* | 657.2991 | 657.3004 | 2.0 |
| *b7* | 758.3468 | 758.3481 | 1.7 |
| *b8* | 845.3788 | 845.3800 | 1.4 |
| *b9* | 960.4058 | 960.4073 | 1.6 |
| *b10* | 1059.4742 | 1059.4757 | 1.4 |
| *b11* | 1146.5062 | 1146.5089 | 2.4 |
| *b12* | 1233.5382 | 1233.5397 | 1.2 |
| *b13* | 1396.6016 | 1396.6027 | 0.8 |
| *b14* | 1509.6856 | 1509.6885 | 1.9 |

| Fragment | Theoretical m/z | Observed m/z | Error Value (ppm) |
|---|---|---|---|
| *y5-NH$_3$* | 526.3208 | 526.3226 | 3.4 |
| *y6-NH$_3$* | 639.4049 | 639.4048 | -0.2 |
| *y7-NH$_3$* | 825.4842 | 825.4872 | 3.6 |
| *y8-NH$_3$* | 896.5213 | 896.5217 | 0.4 |
| *y9* | 1026.6319<br>513.8196 | 513.8218 | 4.3 |
| *y10* | 1173.7004<br>587.3538 | 587.3549 | 1.9 |
| *y11* | 1302.7430<br>651.8751 | 651.8768 | 2.6 |
| *y12* | 2845.5988<br>1423.3030 | 1423.3193 | 11.5 |

**Figure S9**. Analyzed sequences of GLP-1a and conjugated click dibenzylcyclooctyne moiety. The indicated fragments were analyzed using LC/MS-MS to confirm the site of modification.

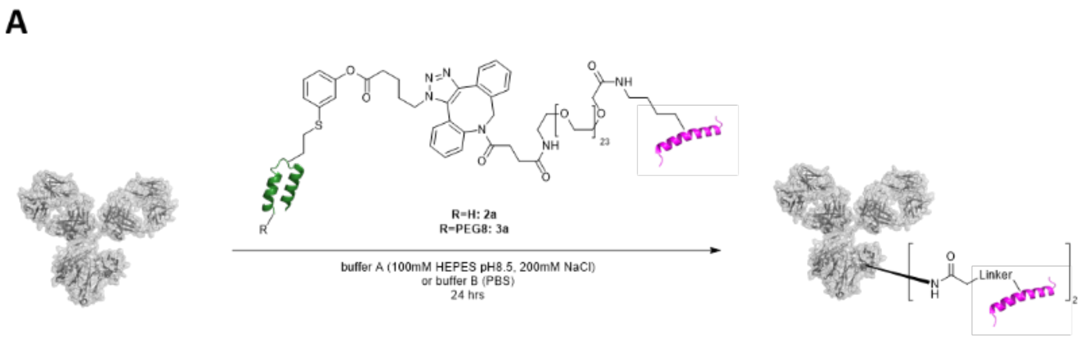

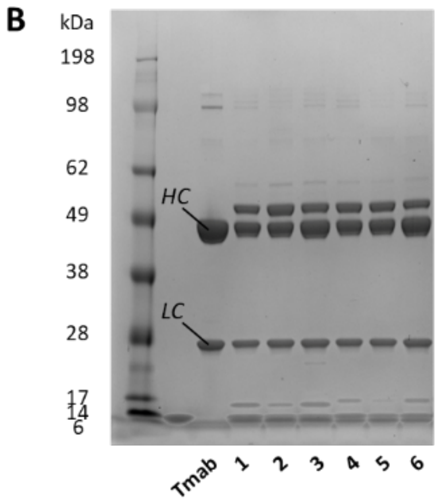

| Entry | Reagent | Buffer | Temp. | Light Chain Reactivity | Heavy Chain Reactivity |
|---|---|---|---|---|---|
| 1 | 2a | A | rt | <1% | 34% |
| 2 | 2a | A | 37°C | <1% | 41% |
| 3 | 2a | B | 37°C | <1% | 29% |
| 4 | 3a | A | rt | <1% | 36% |
| 5 | 3a | A | 37°C | <1% | 35% |
| 6 | 3a | B | 37°C | <1% | 26% |

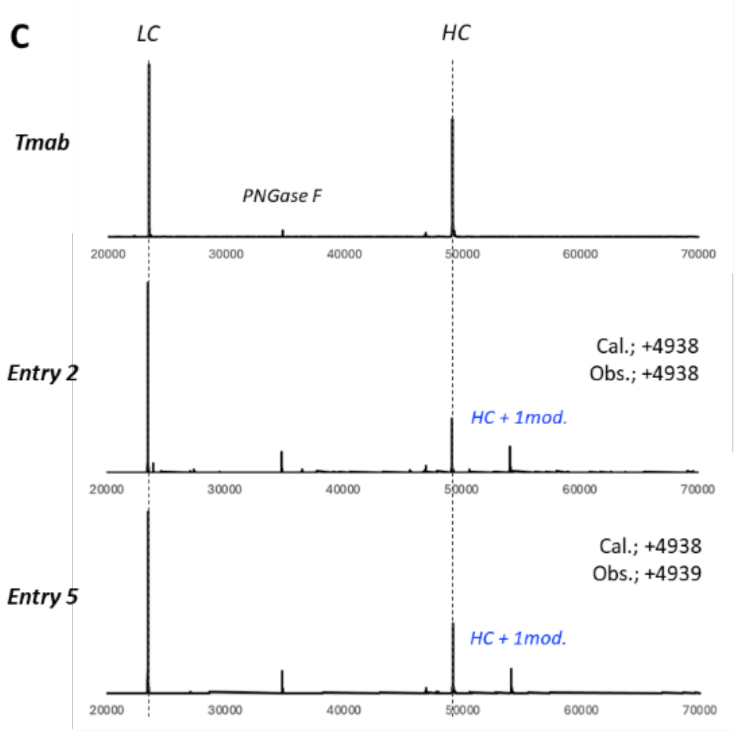

| Entry | Reagent | Buffer | Temp. | Light Chain Reactivity | Heavy Chain Reactivity |
|---|---|---|---|---|---|
| 2 | 2a | A | 37°C | <1% | 33% |
| 5 | 3a | A | 37°C | <1% | 28% |

**Figure S10**. In vitro reactivity of compounds **2a** and **3a** for human IgG1 Trastuzumab (Tmab). (A) Scheme of the reaction (B) SDS-PAGE result post reaction with corresponding quantification. Tmab was reduced beforehand. (C) Mass spectra (MS) post reaction. Tmab was deglycosylated and reduced beforehand.

**A**

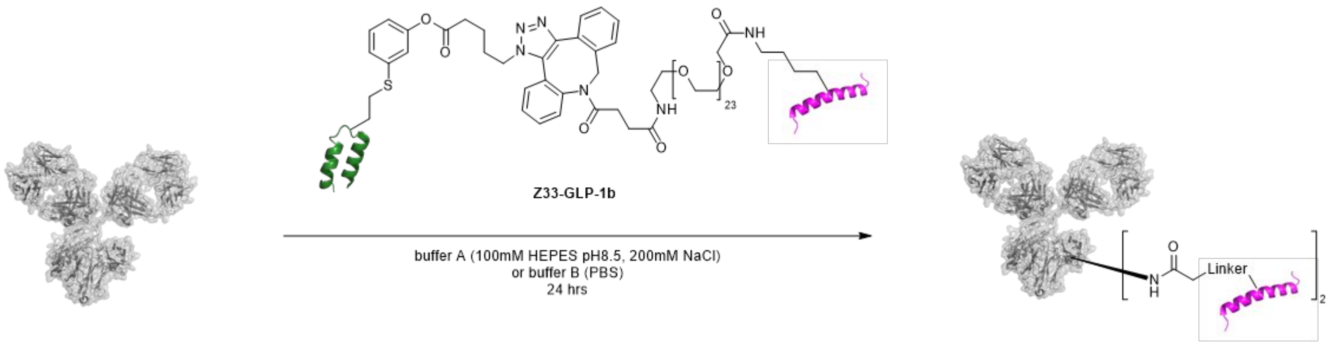

**B**

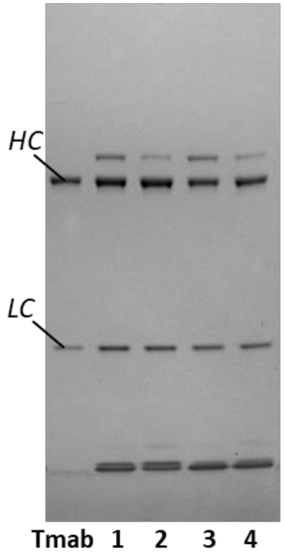

| Entry | Buffer | Temp. | Light Chain Reactivity | Heavy Chain Reactivity |
|---|---|---|---|---|
| **1** | A | rt | <1% | 23% |
| **2** | B | rt | <1% | 9% |
| **3** | A | 37°C | <1% | 26% |
| **4** | B | 37°C | <1% | 11% |

**Figure S11**. In vitro reactivity of compound **2b** for human IgG1 Trastuzumab (Tmab). (A) Scheme of the reaction (B) SDS-PAGE result post reaction and corresponding quantification. Tmab was reduced beforehand.

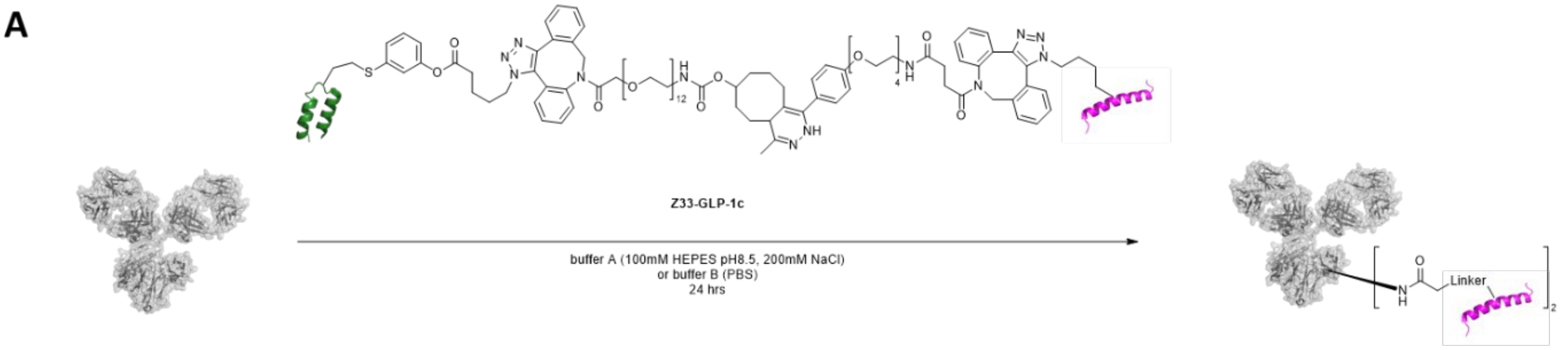

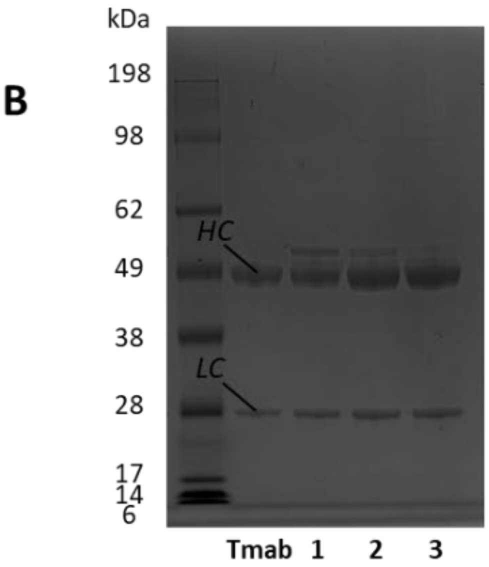

| Entry | Buffer | Temp. | Light Chain Reactivity | Heavy Chain Reactivity |
|---|---|---|---|---|
| **1** | A | rt | <1% | 20% |
| **2** | A | 37°C | <1% | 10% |
| **3** | B | 37°C | <1% | 4% |

**Figure S12.** Reactivity of compound **2c** for human IgG1 Trastuzumab (Tmab). (A) Scheme of the reaction. (B) SDS-PAGE result post reaction and corresponding quantification. Tmab was reduced beforehand.

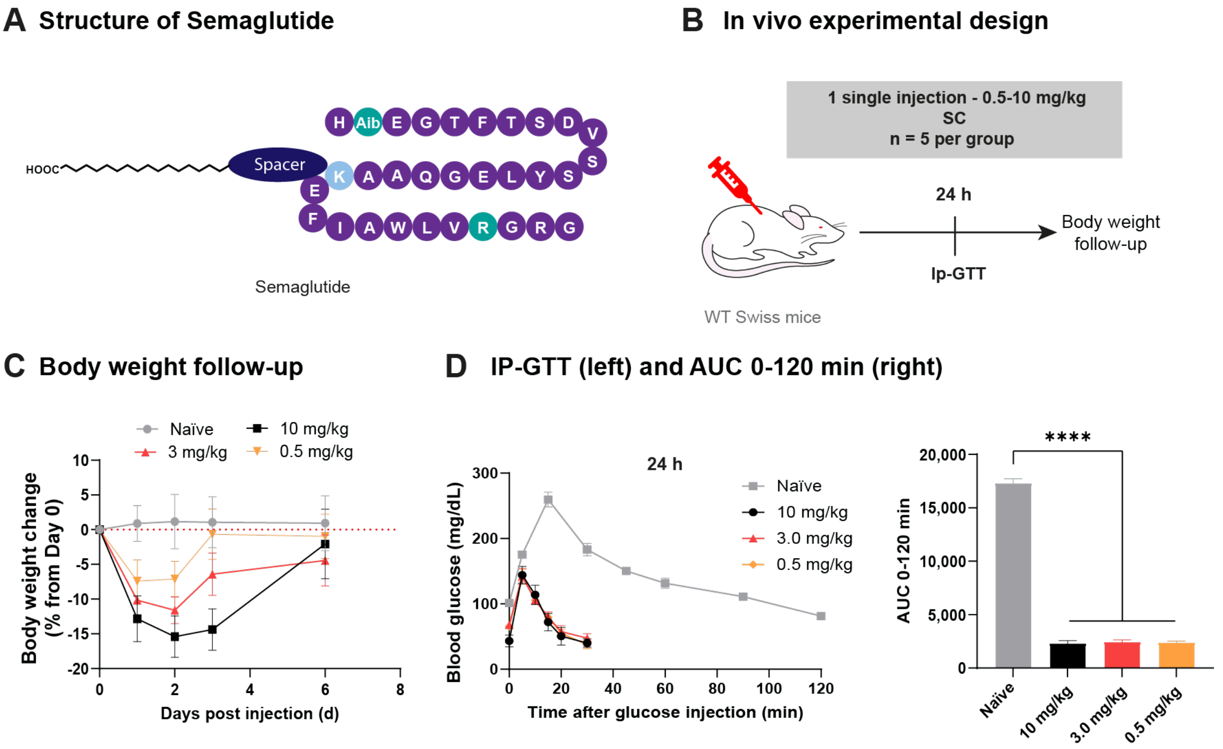

**Figure S13.** Dose response of semaglutide in wild-type Swiss mice. (A) Structure of commercial semaglutide. (B) Experimental design: female WT Swiss mice were subcutaneously (SC) injected with either 0.5 mg/kg (~5 nmol), 3 mg/kg (~30 nmol), or 10 mg/kg (~100 nmol) of semaglutide. Intraperitoneal glucose tolerance test (Ip-GTT) was performed after 24 h. (C) Body weight change from Day 0. (D) Ip-GTT curve (left) and area under the curve (AUC) from 0 to 120 min (right) performed at 24 h post Semaglutide injection. Statistical analysis was performed using non-parametric T-tests: *P<0.05, **P<0.01, ***P<0.001, ****P<0.0001.

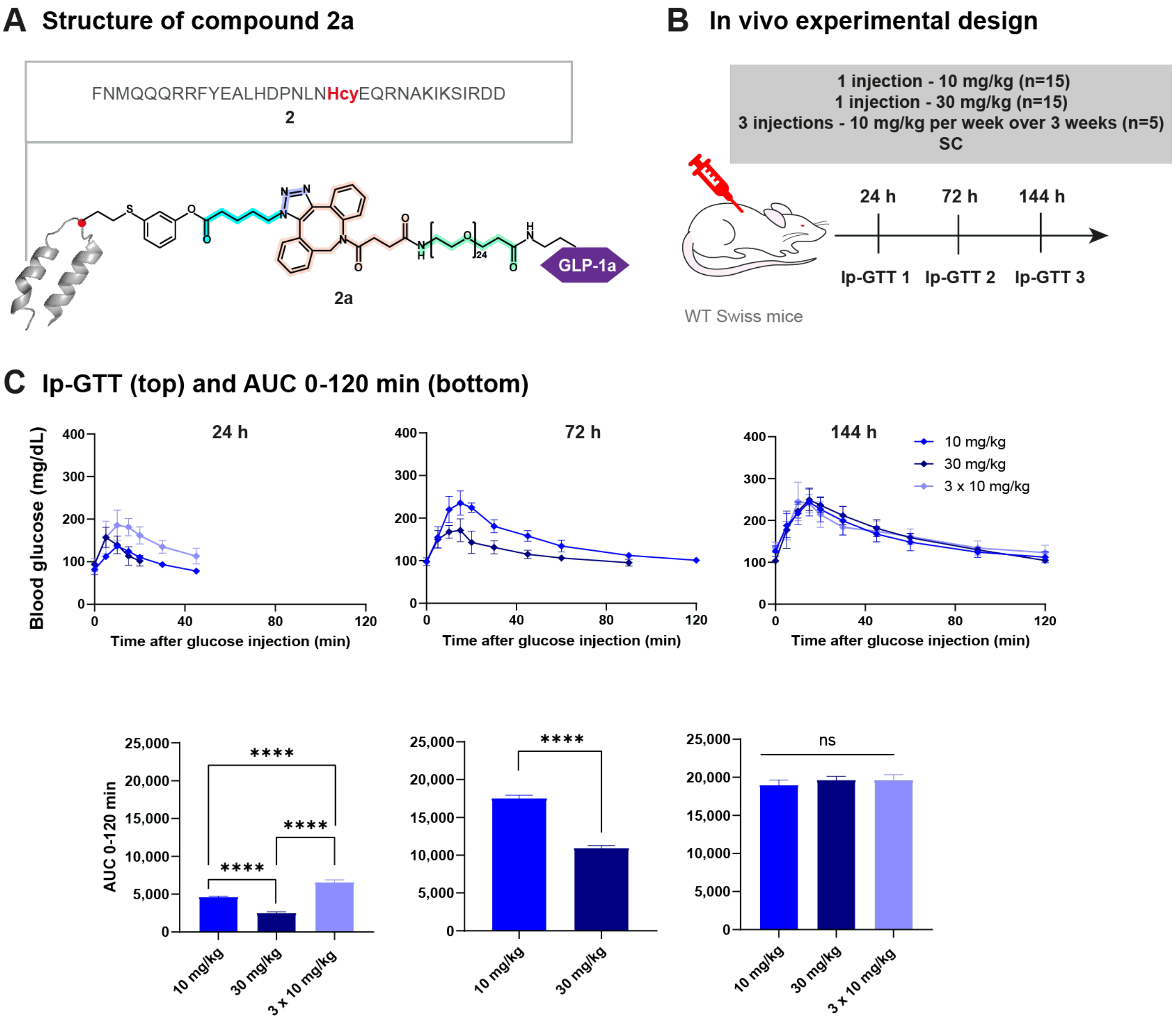

**Figure S14.** Dose response of **2a**. (A) Structure of the affinity peptide **2a**. (B) Experimental design: female WT Swiss mice were subcutaneously (SC) injected either with one single injection of 10 mg/kg (25 nmol) or 30 mg/kg (75 nmol) of **2a**, or with a stacking dose of 3 injections of 10 mg/kg (75 nmol total) through 1 injection per week for 3 weeks. (C) Intraperitoneal glucose tolerance tests (Ip-GTT) performed at 24 h, 72 h, and 144 h post **2a** injection. Top: Ip-GTT curves; Bottom: area under the curves (AUC) from 0 to 120 min. For the stacking dose (3 x 10 mg/kg), Ip-GTT were performed after the 3rd injection of **2a**. Statistical analysis was performed using non-parametric T-tests: *P<0.05, **P< 0.01, ***P< 0.001, ****P<0.0001.

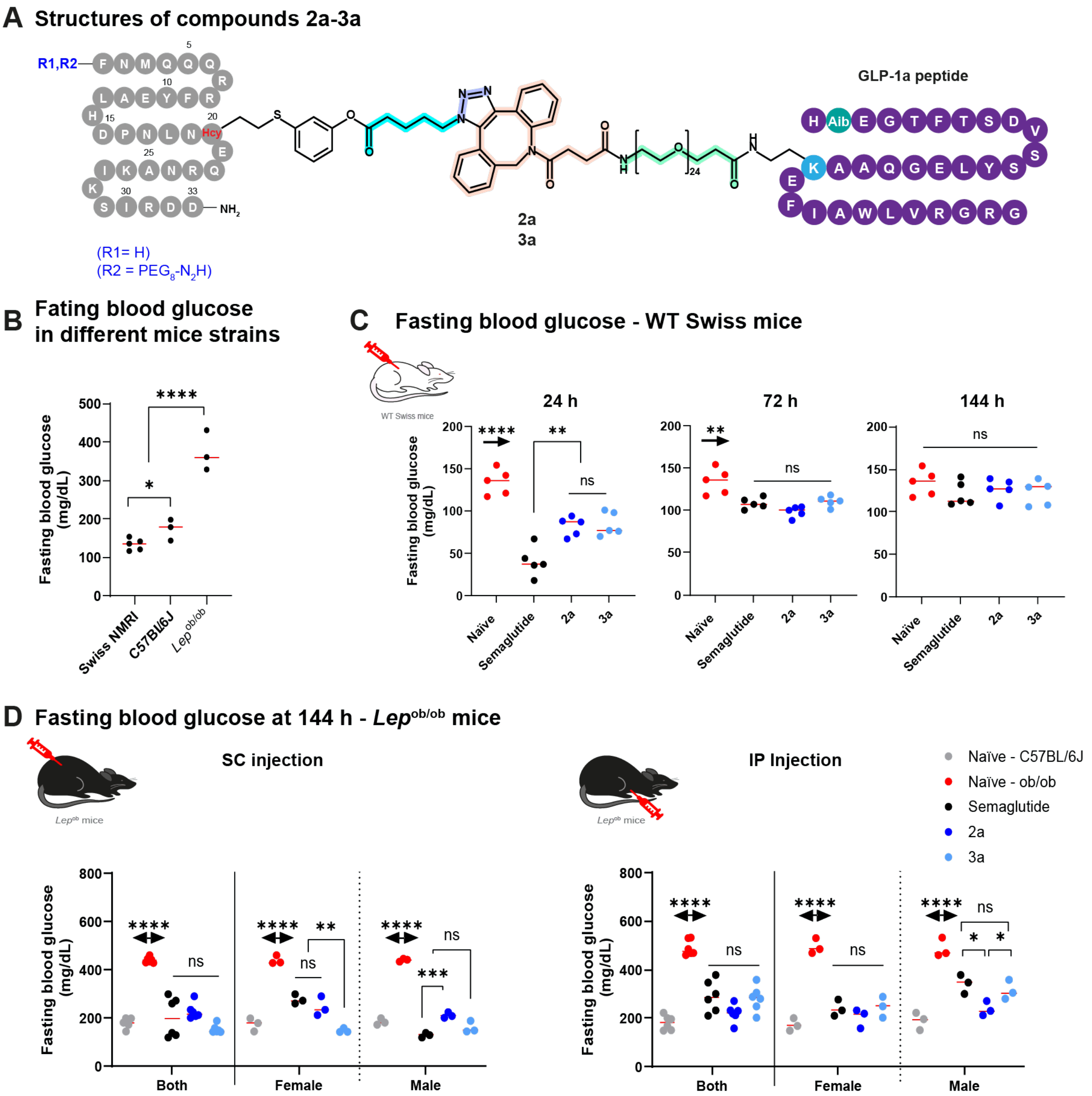

**Figure S15.** Complementary data set for fasting blood glucose in WT Swiss mice and Lep$^{ob/ob}$ mice. Complete main manuscript Figure 2-3 and SI Fig.S11-13. (A) Structure of compounds **2a,3a**. (B) Comparison of fasting blood glucose, after 6 hours of fasting, in different female mice strains. (C) Fasting blood glucose in female WT Swiss mice at 24 h, 72 h, and 144 h post SC injection of 10 mg/kg (~100 nmol) of Semaglutide (Ozempic®), or 10 mg/kg (~25 nmol) of compound **2a,3a**. (D) Fasting blood glucose in Lep$^{ob/ob}$ mice at 144 h post injection of 10 mg/kg (~170 nmol) of Semaglutide (Ozempic®), or 10 mg/kg (~45 nmol) of compound **2a,3a** following SC route (left) or IP (right).

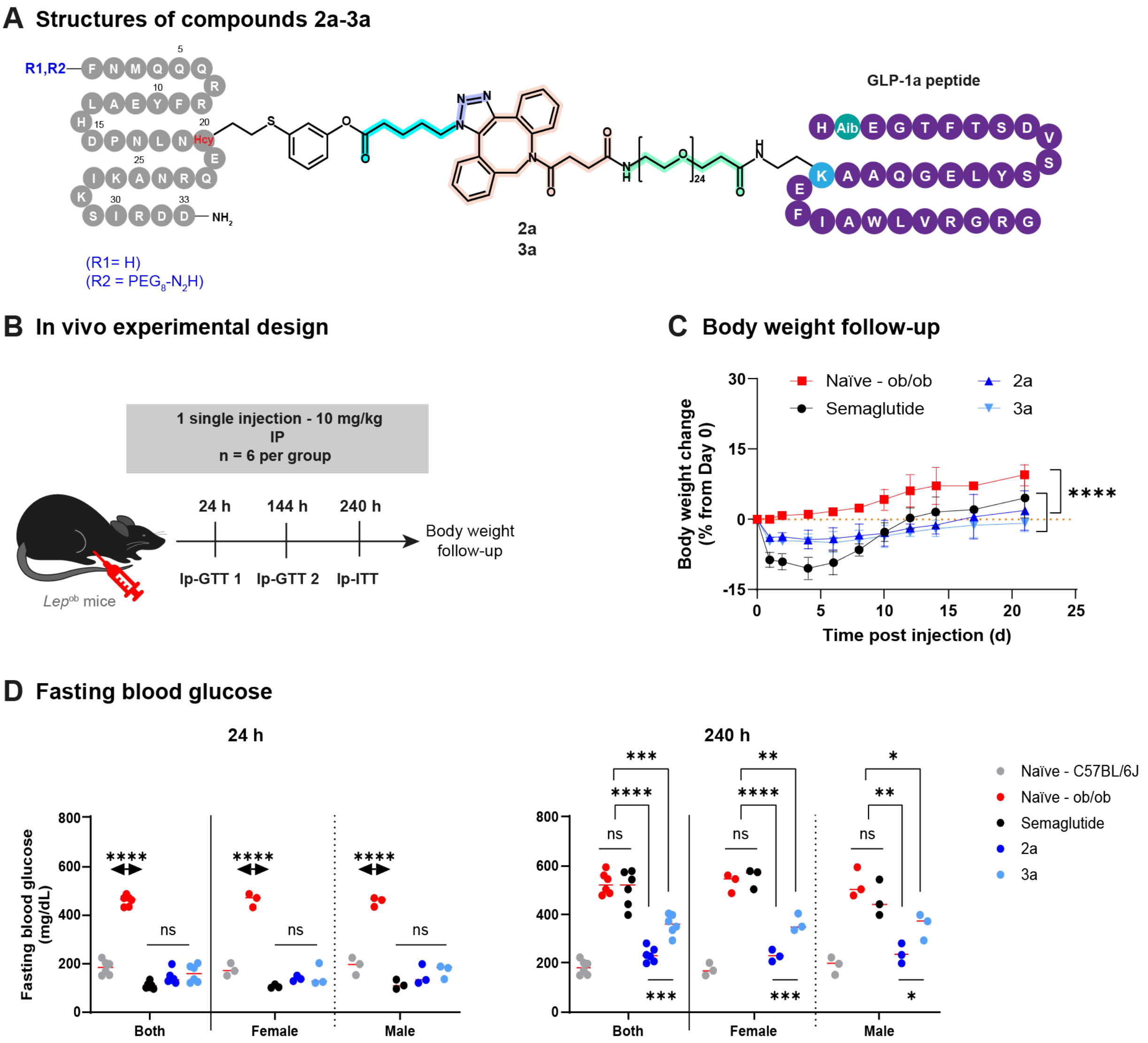

A Structures of compounds 2a-3a
R1,R2
(R1= H)
(R2 = PEG8-N2H)
Hcy
NH2
24
2a
3a
GLP-1a peptide
Aib
B In vivo experimental design
1 single injection - 10 mg/kg
IP
n = 6 per group
24 h
144 h
240 h
Ip-GTT 1
Ip-GTT 2
Ip-ITT
Body weight
follow-up
Lep^ob mice
C Body weight follow-up
Naïve - ob/ob
Semaglutide
2a
3a
Body weight change
(% from Day 0)
Time post injection (d)
****
D Fasting blood glucose
24 h
240 h
Fasting blood glucose
(mg/dL)
Both
Female
Male
ns
Naïve - C57BL/6J
Naïve - ob/ob
Semaglutide
2a
3a

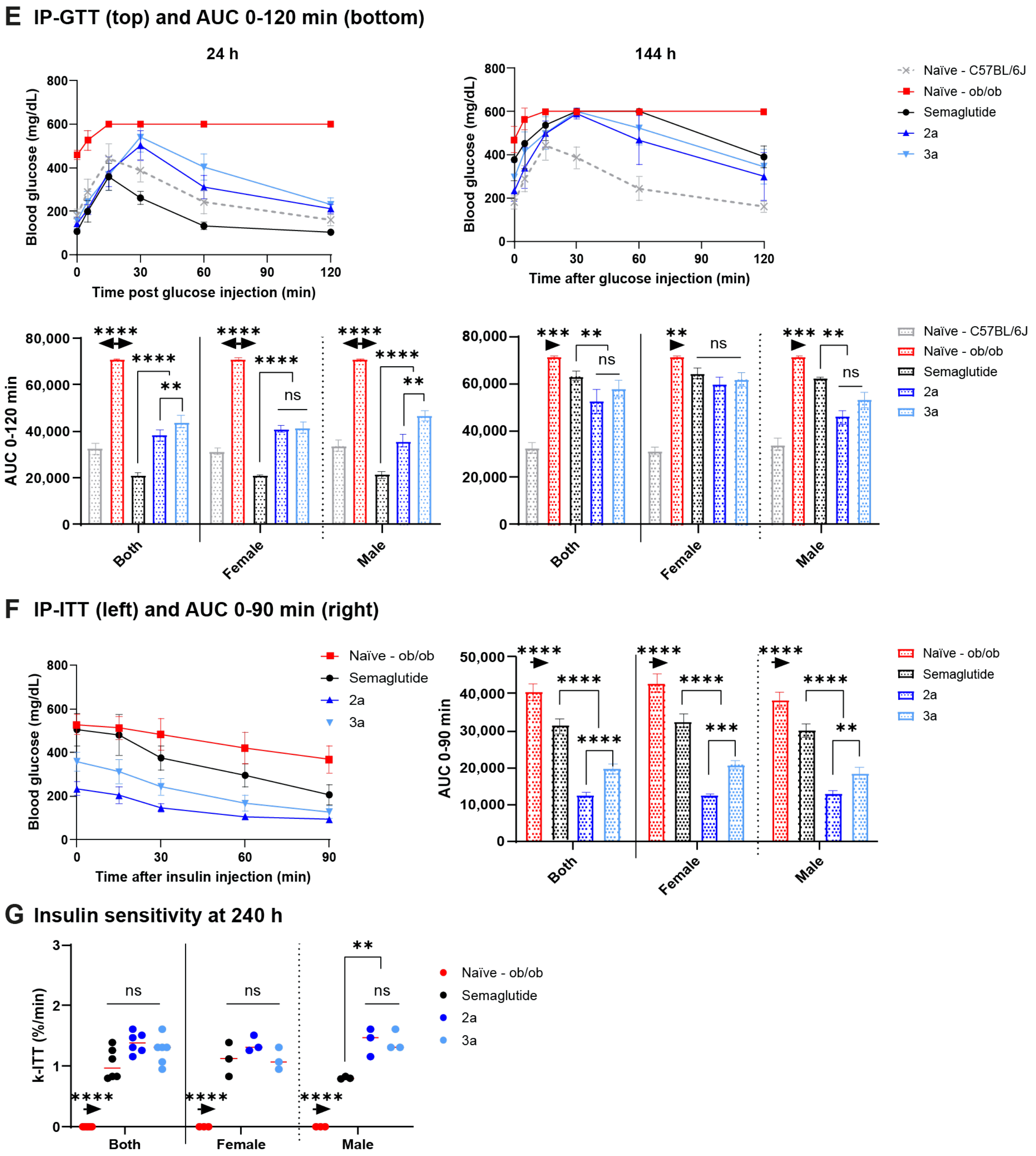

**Figure S16.** IgG painting with GLP-1**a** analogs in Lep$^{ob/ob}$ mice after intraperitoneal (IP) injection of a single 10 mg/kg dose extends GLP-1 efficacy. (A) Structure of electrophile affinity peptides **2a**,**3a**. (B) Experimental design: male and female Lep$^{ob/ob}$ mice were IP injected with 10 mg/kg (~170 nmol) of either Semaglutide (Ozempic®) or 10 mg/kg (~45 nmol) of **2a**,**3a**. IP-glucose tolerance tests (Ip-GTT) was performed after 24 h and 144 h, and Ip-insulin tolerance test (Ip-ITT) was performed at 240 post injection. (C) Body weight change from Day 0. (D) Fasting blood glucose measured at 24 h and 240 h. (E) Ip-GTT curves (top) and area under the curve AUC0-120 min (Bottom) measured at 24 h and 144 h post drug injection. (F) Ip-ITT curve (left) and AUC0-90 min (right) measured at 240 h post drug injection. (G) Plasma glucose rate disappearance (K-ITT) determined from Ip-ITT data, at 240 h post drug injection.

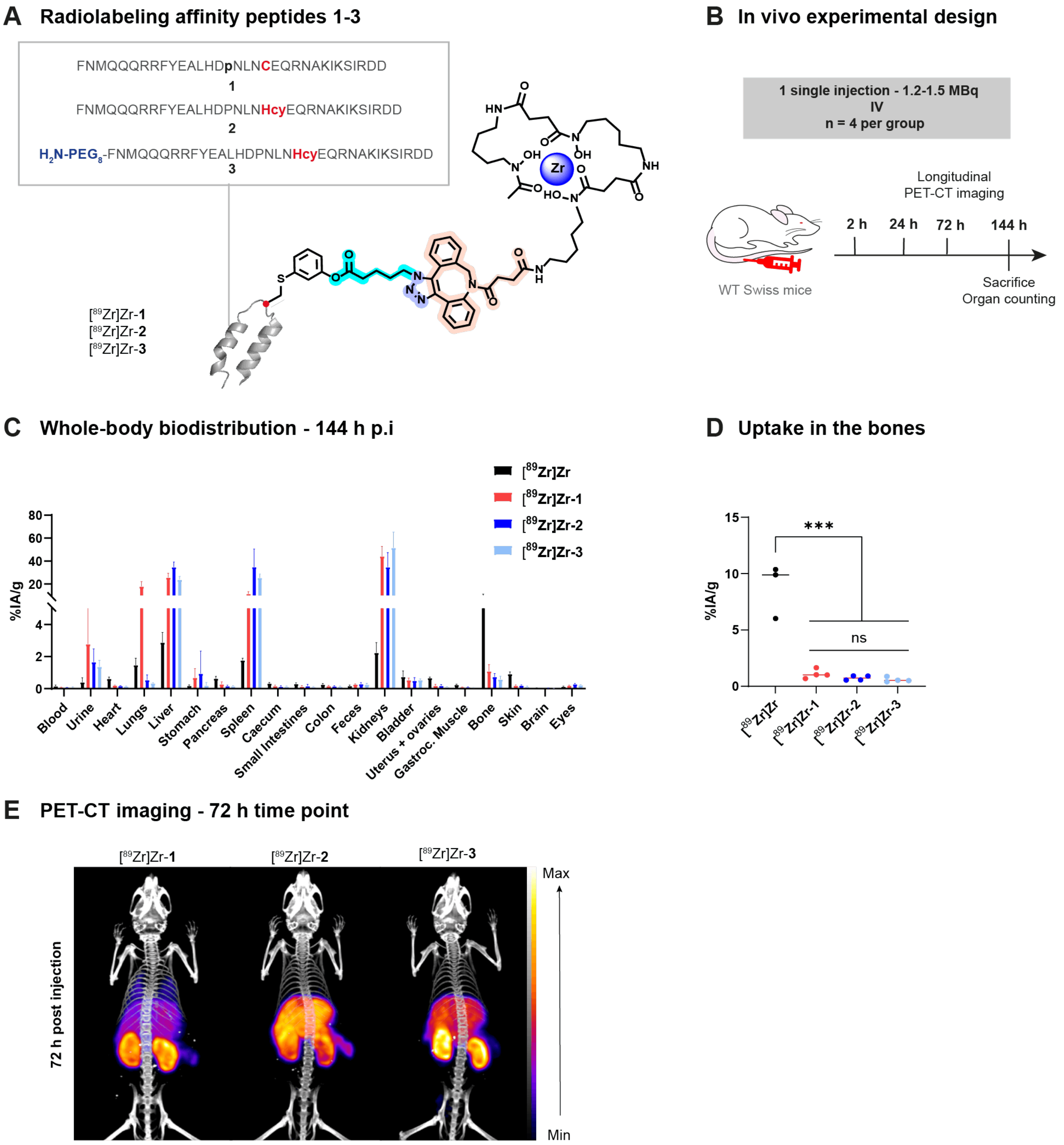

**Figure S67.** Complementary data set for IgG painting with radionuclides after intravenous (IV) injection. Complete data set from Figure 4 of main manuscript. (A) Structures of the radiolabeled affinity peptides [$^{89}$Zr]Zr-**1-3**. (B) Experimental design: female WT Swiss mice were IV injected with 1.2-1.5 MBq of [$^{89}$Zr]Zr-**1-3**. PET-CT imaging was performed over 6 days, then the organs were harvested for quantification. (C) Whole-body biodistribution of [$^{89}$Zr]Zr-**1-3** at 144 h post injection. (D) Uptake in the bones at 144 h post injection. (D) PET-CT imaging at the 72-h time-point post injection. Statistical analysis was performed using non-parametric T-tests: *P<0.05, **P<0.01, ***P<0.001, ****P<0.0001.

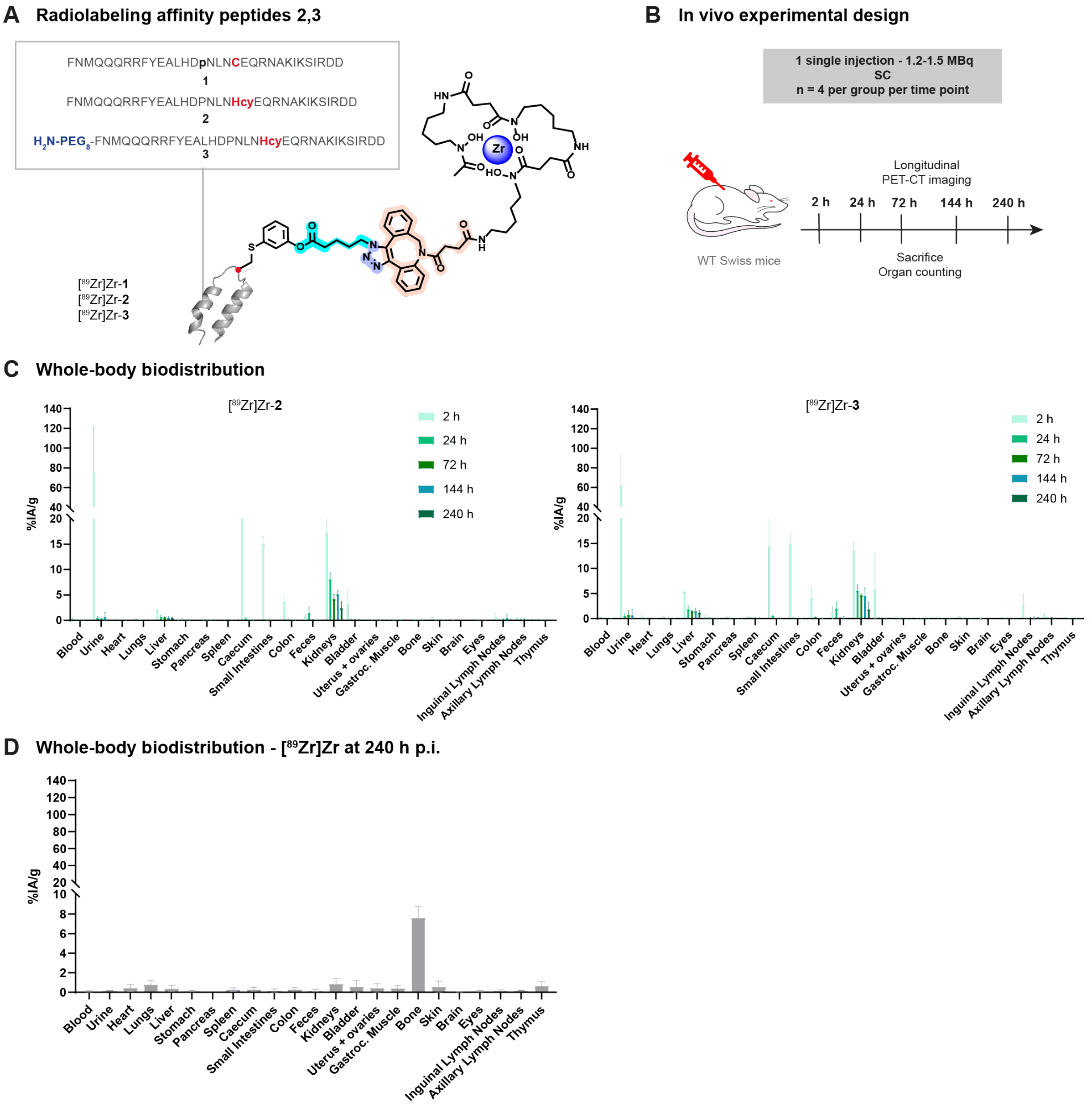

**Figure S78.** In vivo IgG painting with radionuclides after subcutaneous (SC) injection. (A) Structures of the radiolabeled affinity peptides [$^{89}$Zr]Zr-**2,3**. (B) Experimental design: female WT Swiss mice were SC injected with 1.2-1.5 MBq of peptides [$^{89}$Zr]Zr-**2,3** then sacrificed at different time points to quantify the uptake in the organs. (C) Whole-body biodistribution of [$^{89}$Zr]Zr-**2** (left) and [$^{89}$Zr]Zr-**3** (right). (D) Whole-body biodistribution of free [$^{89}$Zr]Zr at 240 h post injection (~1 MBq).

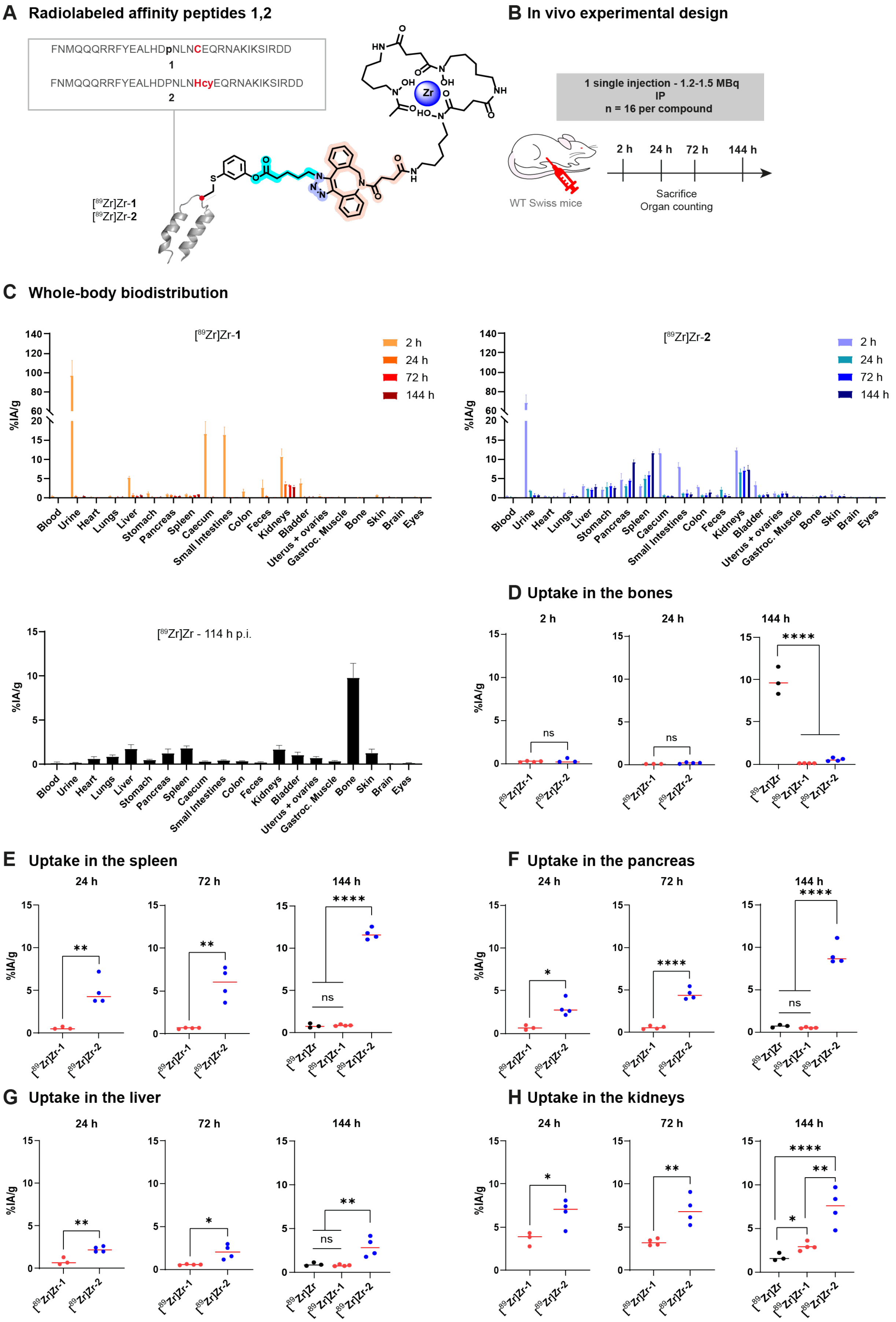
A Radiolabeled affinity peptides 1,2
FNMQQQRRFYEALHDpNLNCEQRNAKIKSIRDD
1
FNMQQQRRFYEALHDPNLNHcyEQRNAKIKSIRDD
2
[89Zr]Zr-1
[89Zr]Zr-2
B In vivo experimental design
1 single injection - 1.2-1.5 MBq
IP
n = 16 per compound
WT Swiss mice
2 h
24 h
72 h
144 h
Sacrifice
Organ counting
C Whole-body biodistribution
[89Zr]Zr-1
[89Zr]Zr-2
%IA/g
Blood
Urine
Heart
Lungs
Liver
Stomach
Pancreas
Spleen
Caecum
Small Intestines
Colon
Feces
Kidneys
Bladder
Uterus + ovaries
Gastroc. Muscle
Bone
Skin
Brain
Eyes
[89Zr]Zr - 114 h p.i.
D Uptake in the bones
E Uptake in the spleen
F Uptake in the pancreas
G Uptake in the liver
H Uptake in the kidneys
[89Zr]Zr
ns
*
**
****

**Figure S89.** In vivo IgG painting with radionuclides after intraperitoneal (IP) injection. (A) Structures of the radiolabeled affinity peptides [$^{89}$Zr]Zr-**1,2**. (B) Experimental design: female WT Swiss mice were IP injected with 1.2-1.5 MBq of [$^{89}$Zr]Zr-**1,2** then sacrificed at different time points to quantify the uptake in the organs. (C) Whole-body biodistribution of [$^{89}$Zr]Zr-**1** (left) and [$^{89}$Zr]Zr-**2** (right). Whole-body biodistribution of free [$^{89}$Zr]Zr at 144 h post injection (~1 MBq). (D) Uptake in the bones. (E) Percent of injected activity per gram (%IA/g) in the spleen. (F) %IA/g in the pancreas. (G) %IA/g in the liver. (G) %IA/g in the kidneys. Statistical analysis performed using non-parametric T-tests: *P<0.05, **P<0.01, ***P<0.001, ****P<0.0001.

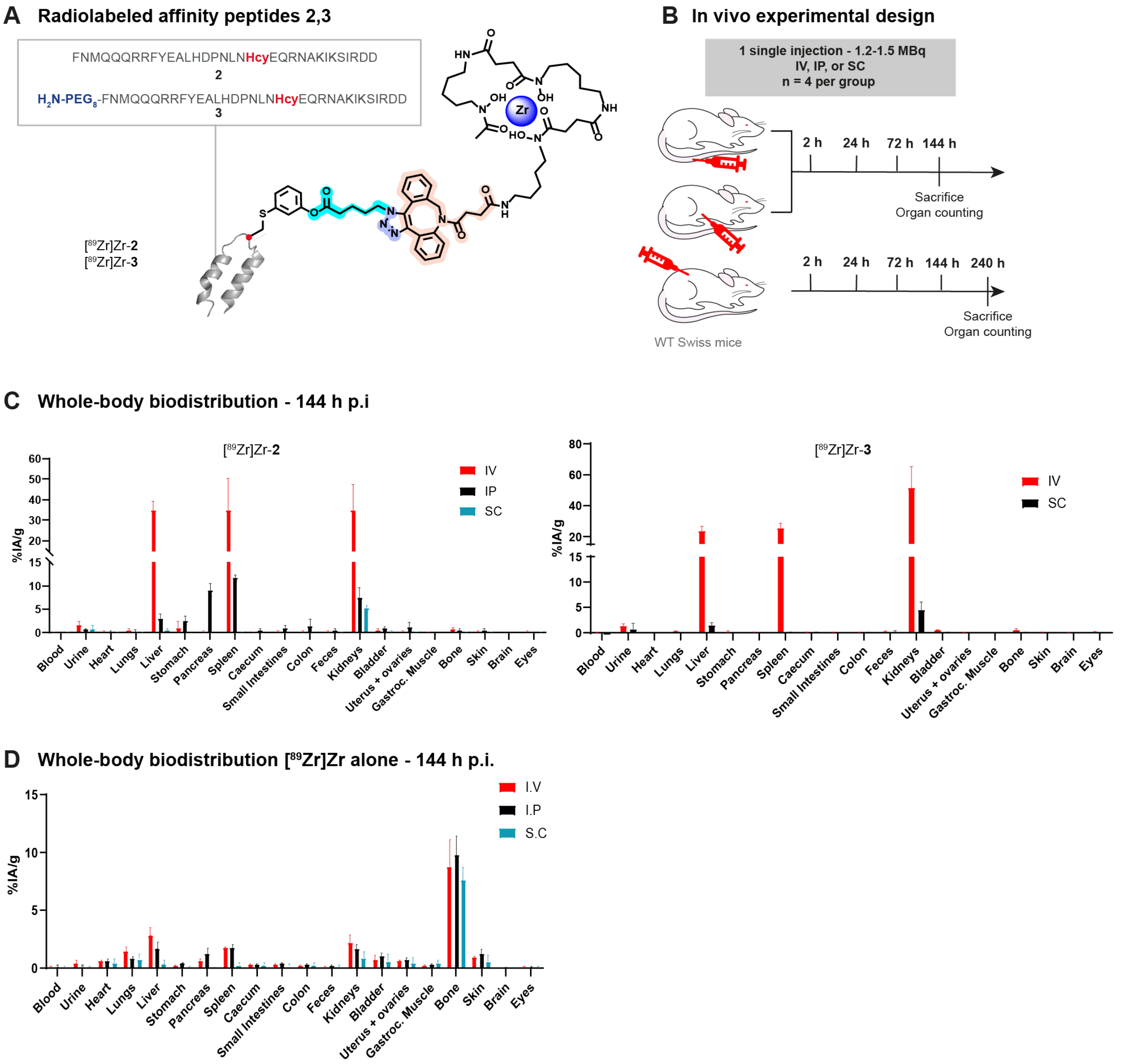

**Figure S20**. Comparison of the routes of injection for in vivo IgG painting using radionuclides, in WT Swiss mice. (A) Structures of the radiolabeled affinity peptides [$^{89}$Zr]Zr-**2**,**3**. (B) Experimental design: female WT Swiss mice were injected, either IV, IP, or SC, with 1.2-1.5 MBq of [$^{89}$Zr]Zr-**2**,**3**. (C) Whole-body biodistribution of [$^{89}$Zr]Zr-**2** (left) and [$^{89}$Zr]Zr-**3** (right) following IV, Ip, or SC, at 144 h post injection. (D) Whole-body biodistribution of free [$^{89}$Zr]Zr at 144 h following IV, IP, or SC injection (~1MBq).

A

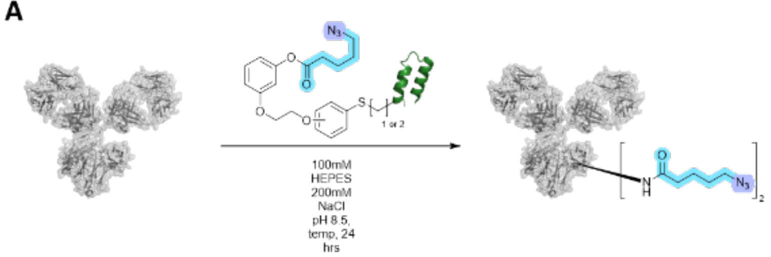

| Entry | Sequence of hIgG-fc Binding Peptide | Structure Attached on the C or Hcy | Eqiovalent of Electrophile | Reaction Temp. | Light Chain Reactivity | Heavy Chain Reactivity | Heavy Chain reaction selectivity |
|---|---|---|---|---|---|---|---|
| 1 | FN**C**QQQRRFYEALHDPNLNEEQRNAKIKSIRDD | VI | 10 | rt | <1% | 44% | >99% |
| 2 | FN**Hcy**QQQRRFYEALHDPNLNEEQRNAKIKSIRDD | VI | 20 | rt | <1% | 27% | 82% |
| 3 | FN**C**QQQRRFYEALHDPNLNEEQRNAKIKSIRDD | VII | 10 | 37°C | <1% | >99% | 85% |
| 4 | FN**Hcy**QQQRRFYEALHDPNLNEEQRNAKIKSIRDD | VII | 20 | rt | <1% | 38% | 88% |

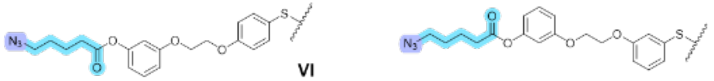

B

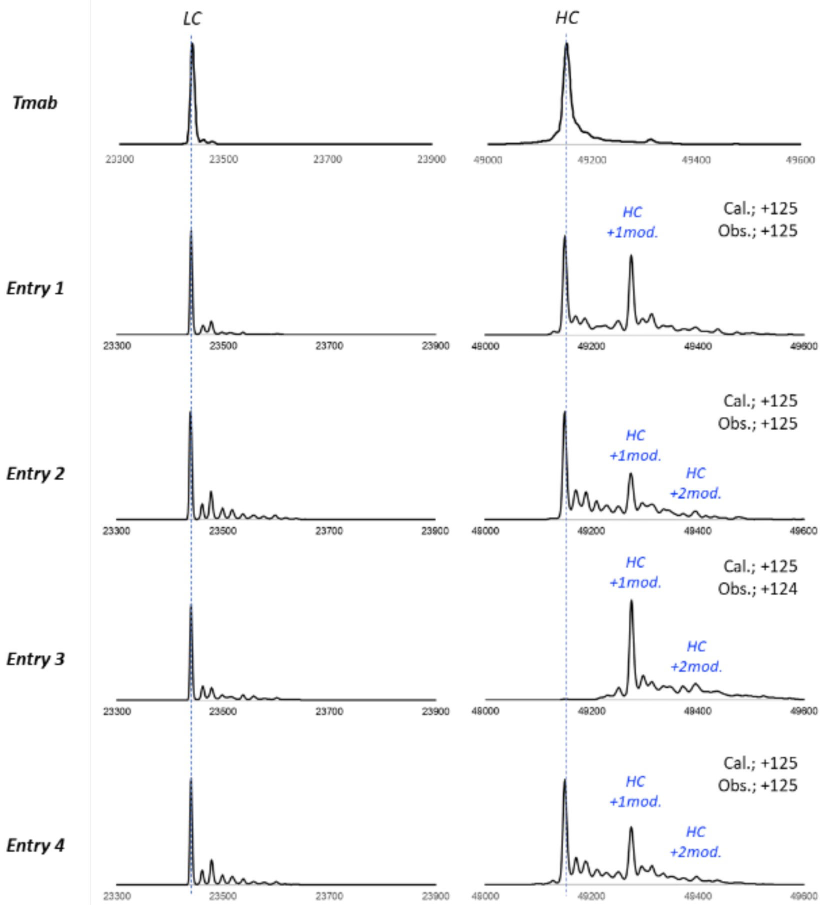

**Figure S21**. In vitro reactivity of selected electrophile affinity peptides for human IgG1 Trastuzumab (Tmab). (A) Percentage of modification on heavy and light chains of Tmab depending on the amino acid sequence of the affinity peptide and structure of the electrophiles. The conjugation site on the affinity peptide is highlighted in blue. "Heavy chain reaction selectivity" was calculated based on the percentage of lysine residues on the heavy chain with at least one modification. (B) Mass spectra (MS) post reaction. Tmab was deglycosylated and reduced beforehand. Left: MS for the light chain; Right: MS for the heavy chain.

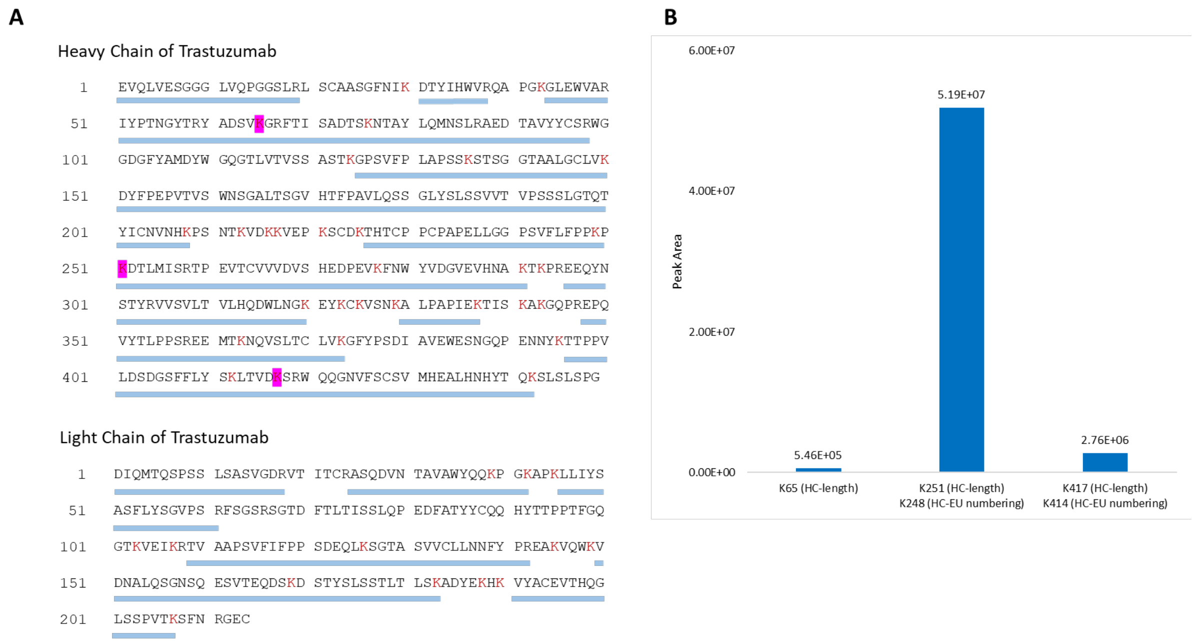

**Figure S22**. Identification of the sites of modification on IgG1 Trastuzumab (Tmab) after reaction with **N$_3$-8-VI**. (A) Sequencing of Tmab after digestion with trypsin and analysis by LC-MS/MS. Lysines are in red. Pale blue bars below the sequence indicate the observed fragments. The lysines exhibiting modifications are highlighted in magenta. (B) Peak areas corresponding to the lysines modified by the affinity peptides.

**A**

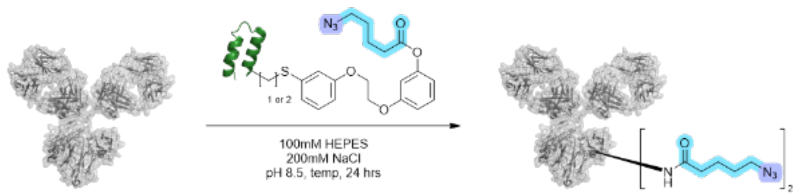

| Entry | Sequence of hIgG-fc Binding Peptide | Eqiovalent of Electrophile | Reaction Temp. | Light Chain Reactivity | Heavy Chain Reactivity | Heavy Chain reaction selectivity |
|---|---|---|---|---|---|---|
| 1 | FNMQQQRRFYEALHDPNLNEEQRNAKIKSICDD | 10 | rt | <1% | 21% | >99% |
| 2 | FNMQQQRRFYEALHDPNLNEEQRNAKIKSIRCD | 20 | rt | <1% | 32% | >99% |
| 3 | FNMQQQRRFYEALHDPNLNEEQRNAKIKSIRHcyD | 10 | 37°C | <1% | 27% | >99% |

**B**

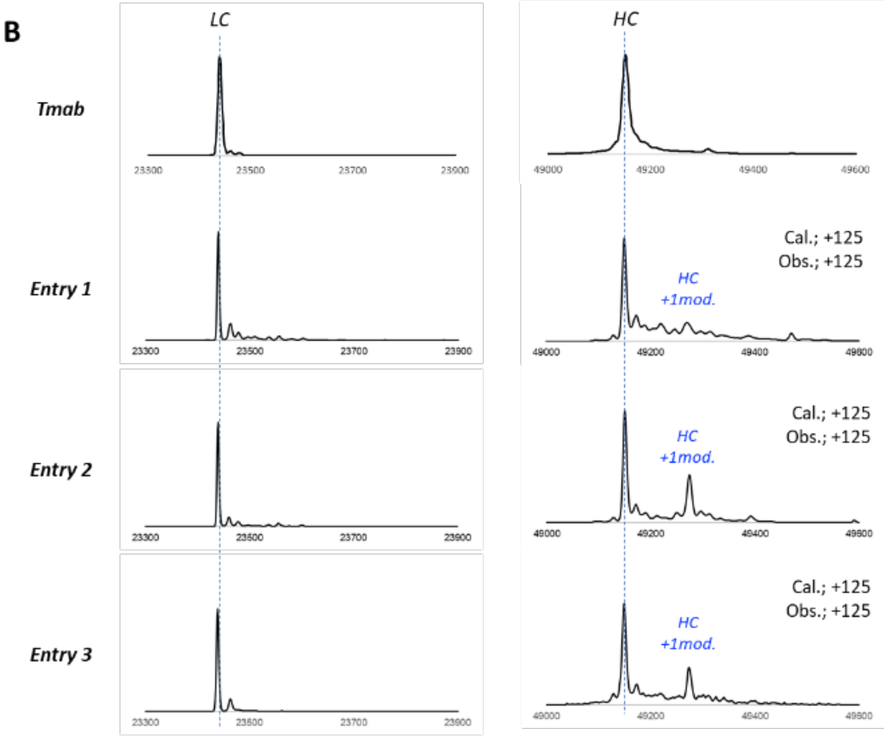

**Figure S23**. In vitro reactivity of selected electrophile affinity peptides for human IgG1 Trastuzumab (Tmab). (A) Percentage of modification on heavy and light chains of Tmab depending on the amino acid sequence of the affinity peptide and structure of the electrophiles. The conjugation site on the affinity peptide is highlighted in blue. “Heavy chain reaction selectivity” was calculated based on the percentage of lysine residues on the heavy chain with at least one modification. (B) Mass spectra (MS) post reaction. Tmab was deglycosylated and reduced beforehand. Left: MS for the light chain; Right: MS for the heavy chain.

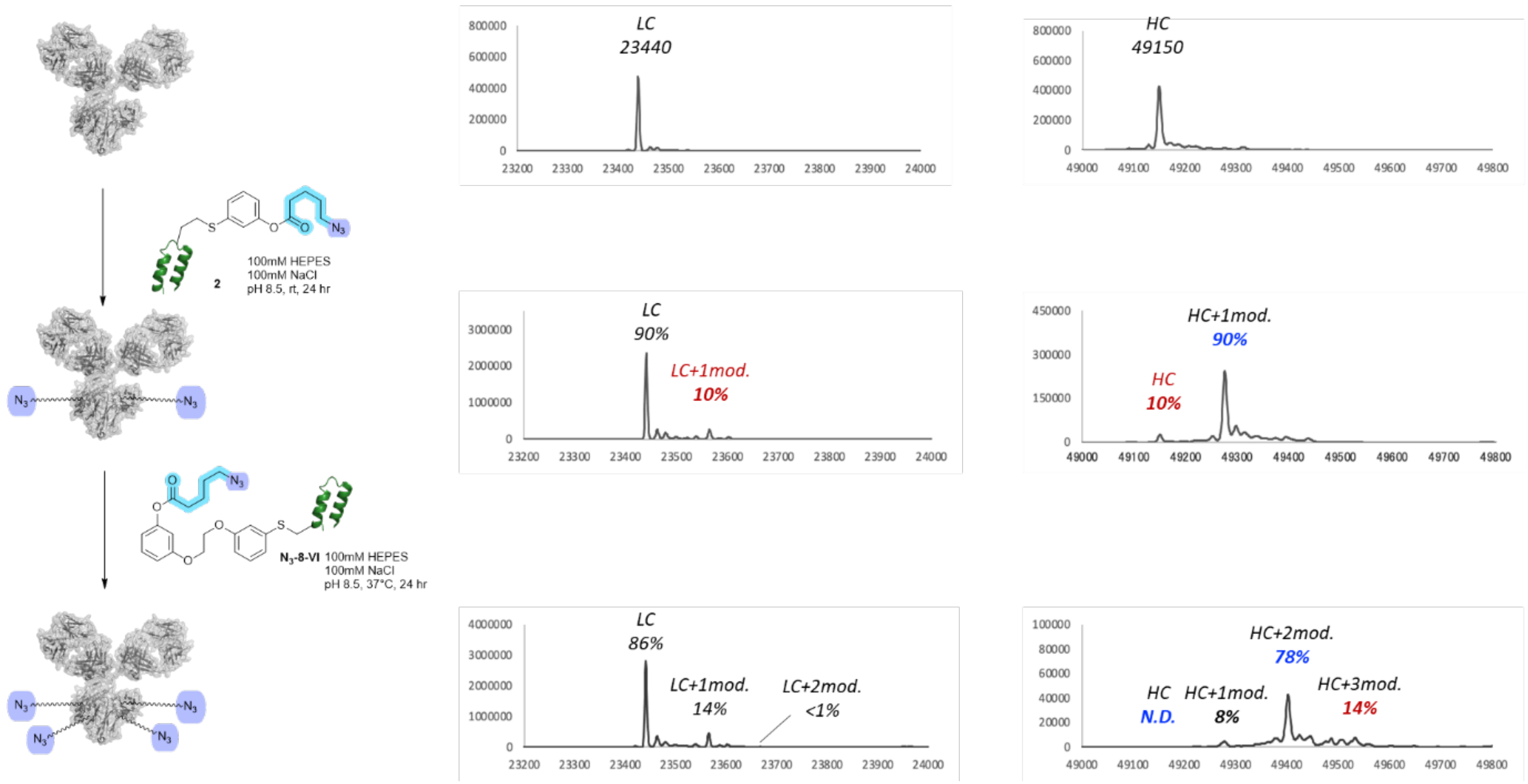

**Figure S24**. Dual modification of Trastuzumab heavy chain by stepwise addition of compound **2** and $N_3$-**8-VI**.

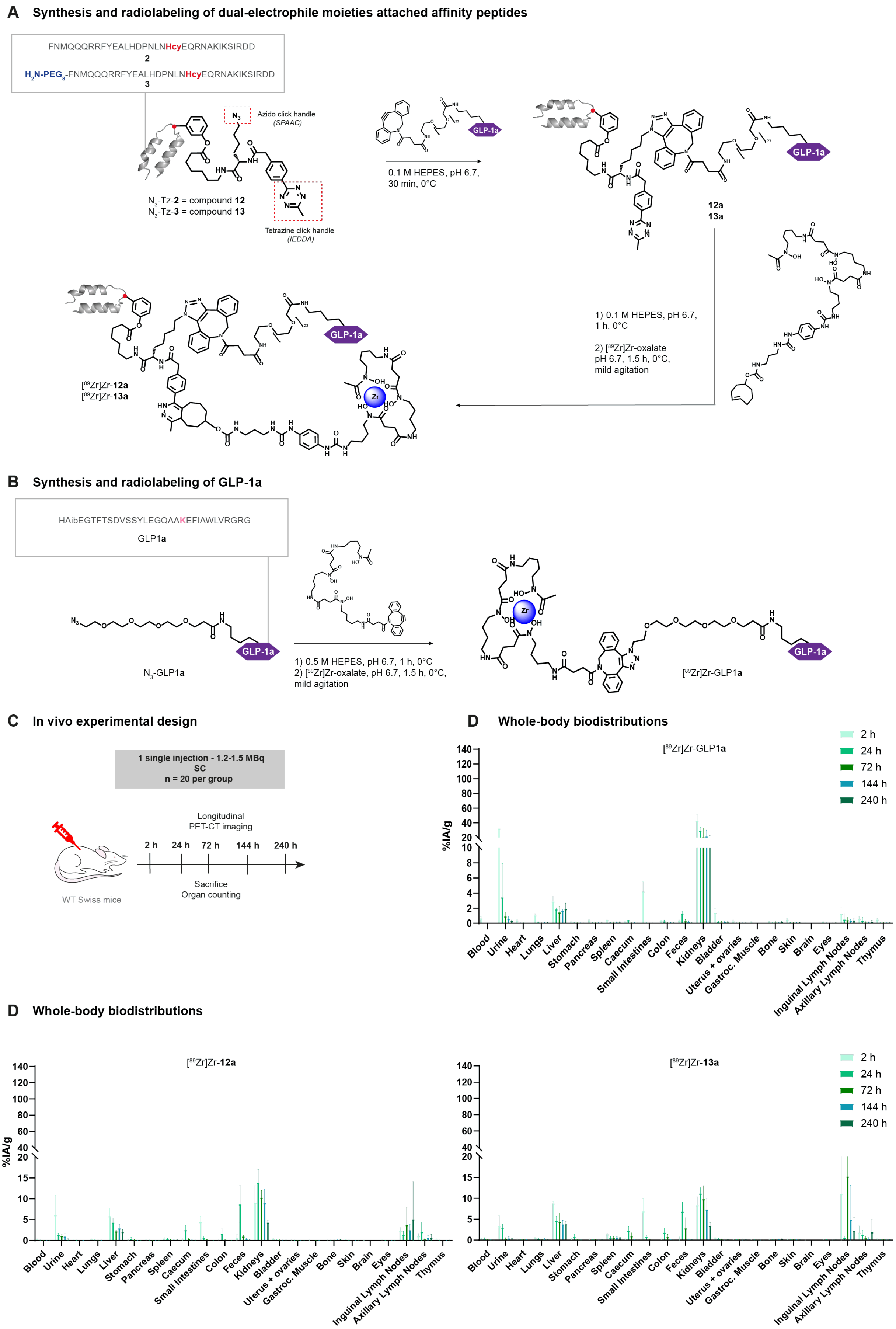

**Figure S25**. Dual in vivo IgG painting with radiolabeled GLP-1**a** analogs after SC injection.

(A) Synthesis and radiolabeling of [$^{89}$Zr]Zr-**12,13**. (B) Synthesis and radiolabeling of [$^{89}$Zr]Zr-GLP-1**a**. (C) Experimental design: female WT Swiss mice were SC injected with 1.2.-1.5 MBq of [$^{89}$Zr]Zr-**12a,13a** or [$^{89}$Zr]Zr-GLP-1**a** then sacrificed at several time points post injection. (D) Whole-body biodistribution of [$^{89}$Zr]Zr-GLP-1**a** (top right), [$^{89}$Zr]Zr-**12a** (bottom left), and [$^{89}$Zr]Zr-**13a** (bottom, right).

A

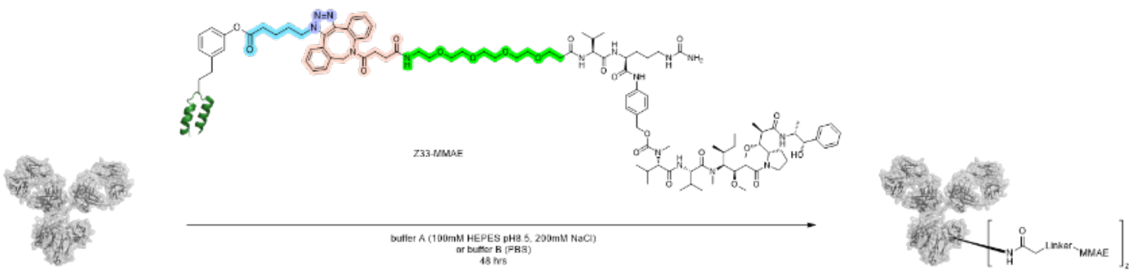

| Entry | Buffer | Temp. | Light Chain Reactivity | Heavy Chain Reactivity |
|---|---|---|---|---|
| 1 | A | rt | <1% | 46% |
| 2 | B | rt | <1% | 42% |
| 3 | A | 37°C | <1% | 54% |
| 4 | B | 37°C | <1% | 55% |

B

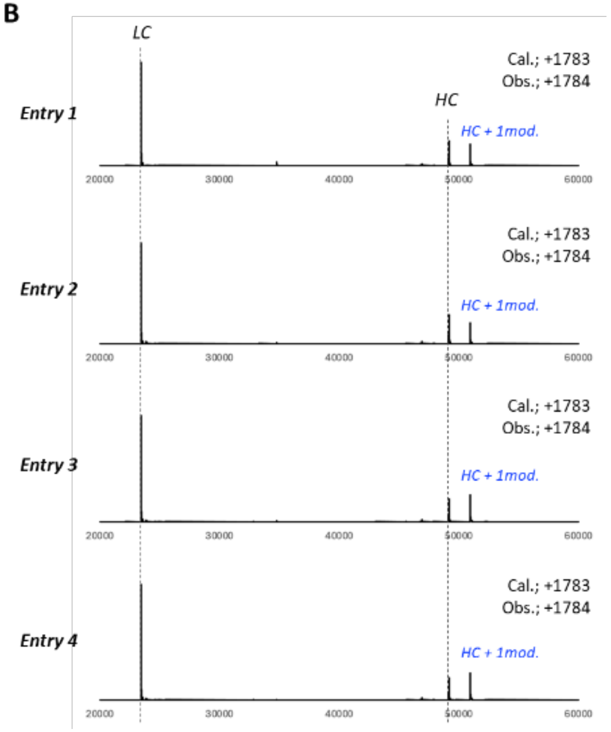

**Figure S26**. In vitro reactivity of MMAE-attached electrophile affinity peptides for human IgG1 Trastuzumab (Tmab). (A) Percentage of modification on heavy and light chains of Tmab depending on the amino acid sequence of the affinity peptide and structure of the electrophiles. The conjugation site on the affinity peptide is highlighted in blue. "Heavy chain reaction selectivity" was calculated based on the percentage of lysine residues on the heavy chain with at least one modification. (B) Mass spectra (MS) post reaction. Tmab was deglycosylated and reduced beforehand.
